# Plant mixed lineage kinase domain-like proteins limit biotrophic pathogen growth

**DOI:** 10.1101/681015

**Authors:** Lisa Mahdi, Menghang Huang, Xiaoxiao Zhang, Ryohei Thomas Nakano, Leïla Brulé Kopp, Isabel M.L. Saur, Florence Jacob, Viera Kovacova, Dmitry Lapin, Jane E. Parker, James M. Murphy, Kay Hofmann, Paul Schulze-Lefert, Jijie Chai, Takaki Maekawa

## Abstract

Mixed lineage kinase domain-like (MLKL) protein mediates necroptotic cell death in vertebrates. We report here the discovery of a conserved protein family across seed plants that is structurally homologous to vertebrate MLKL. The *Arabidopsis thaliana* genome encodes three MLKLs with overlapping functions in limiting growth of obligate biotrophic fungal and oomycete pathogens. Although displaying a cell death activity mediated by N-terminal helical bundles, termed HeLo domain, *At*MLKL-dependent immunity can be separated from host cell death. Cryo-electron microscopy structures of *At*MLKLs reveal a tetrameric configuration, in which the pseudokinase domain and brace region bury the HeLo-domains, indicative of an auto-repressed complex. We also show the association of two *At*MLKLs with microtubules. These findings, coupled with resistance-enhancing activity and altered microtubule association of a phosphomimetic mutation in the pseudokinase domain of *At*MLKL1, point to a cell death-independent immunity mechanism.

**One Sentence Summary:** Plants have a protein family that is structurally homologous to vertebrate mixed lineage kinase domain-like protein, which induces necroptotic cell death, but these plant proteins can confer immunity without host cell death.

## Main text

Regulated cell death (RCD) is intimately connected with innate immunity in plants and animals (*1–3*). A shared feature of several proteins involved in RCD in plants, animals and fungi is a four-helical bundle structure called the HeLo-domain (*4*). HeLo domain-containing MLKL (mixed lineage kinase domain-like protein) mediates necroptosis in animals (*5, 6*), a form of RCD that is proposed to combat pathogens by releasing pro-inflammatory molecules (*7, 8*). Necroptosis is initiated by the plasma membrane (PM)-resident death receptors, with downstream activation of receptor-interacting serine/threonine-protein kinases (RIPKs) leading to the phosphorylation of the pseudokinase domain of the terminal pathway effector, MLKL (*7, 8*). This process assembles monomeric MLKL into pro-necroptotic oligomers (*9*) that translocate to the PM where oligomerized HeLo-domains interfere with membrane integrity (*10–14*). The extent to which RCD in plants and animals is directly responsible for disease resistance is under debate (*3, 15–17*).

To identify novel immune regulators in plants, we searched for HeLo domain-containing proteins in plant genomes by comparing Hidden Markov Models. This analysis identified a protein family that is highly conserved across plants (Fig. 1, Table S1, Supplemental file 1), with a modular structure resembling MLKL (Fig. 1A). The kinase-like domain lacks canonical residues known to underlie phosphoryl transfer (*18*) (Fig. 1B), suggestive of a catalytically inactive pseudokinase (*8*). Hereafter, we refer to these proteins as plant MLKLs. Plant MLKLs additionally possess an extended serine-rich region of varying length after the pseudokinase domain without similarity to any known domain (Fig. 1A, Supplemental file 1). Plant MLKLs separate into two subgroups based on sequence similarity (Fig. 1D). *Arabidopsis thaliana* harbours three *MLKL* genes with *AtMLKL1* and *AtMLKL2* in subfamily I and *AtMLKL3* belonging to subfamily II (Fig. 1D).

**Fig. 1:**
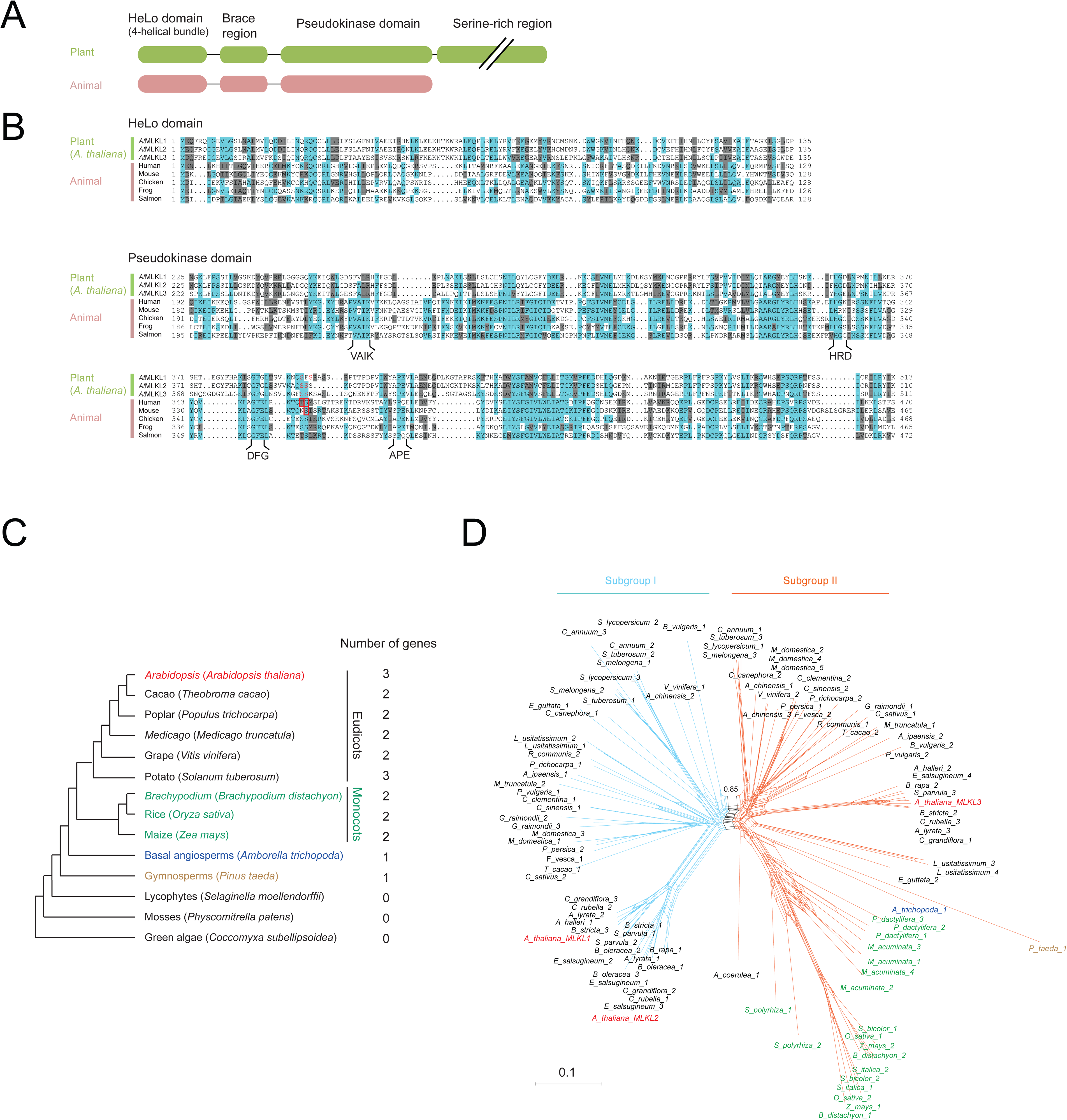
Plant genomes encode structural homologs of MLKL. **A**, A diagram of plant and animal MLKLs. A variably sized serine-rich region that has no similarity to known structures is directly C-terminal to the pseudokinase domain of plant MLKLs. **B**, Alignments of the four helical bundles (HeLo domain) and the pseudokinase domains of *A. thaliana* MLKLs (*At*MLKLs) and representative vertebrates. Invariant residues and conservative substitutions in > 50% of the sequences are shown in light blue and grey backgrounds, respectively. The residues responsible for the activation of human or mouse MLKL upon phosphorylation are indicated in the red box. The serine residues examined in this study are shown in red. **C**, Plant MLKL-encoding genes across representative plant species. A full list of MLKL-encoding genes in the genomes of 48 seed plants is shown in Table S1. **D**, Phylogenetic relationship of plant MLKLs. Neighbor-net analysis discriminates two subgroups of plant MLKLs colored in light blue and orange with bootstrap support of 0.85 (1,000 bootstrap replicates).

To explore similarities between plant and animal MLKLs, we expressed *At*MLKL2 and *A*tMLKL3 for structural analysis using cryo-electron microscopy (Fig. 2, Fig. S1 and S2). In gel filtration, both proteins eluted at an estimated molecular weight corresponding to tetramers (Fig. 2A, Fig. S2A). This contrasts with the vertebrate MLKL protein, which displayed heterogeneity in a similar assay (*13*). Representative 2D projection views indicate that both *At*MLKL oligomers form a triangle-like architecture with a 2-fold symmetry (Fig. S1AB, Fig. S2BC). Further 3D classification and refinement generated density maps of oligomeric *At*MLKL2 and *At*MLKL3 with a global resolution of 4.1Å (Fig. S2E) and 3.4Å (Fig. S1D) respectively, based on the gold Fourier Shell Correlation standard (Fig. S1C, Fig. S2D). The 3D reconstructions show that both *At*MLKL2 and *At*MLKL3 oligomers are composed of four MLKL molecules (Fig. 2C, Fig. S3A), confirming that these form tetramers. Tetramerization of *At*MLKL2 or *At*MLKL3 results in formation of a pyramid-like structure. Structure alignment of the two final models indicates that tetramers of two subfamily members are nearly identical (Fig. S3AB). As the quality of the density of *At*MLKL3 is superior for model building, we limited our structural analysis to *At*MLKL3.

**Fig. 2.**
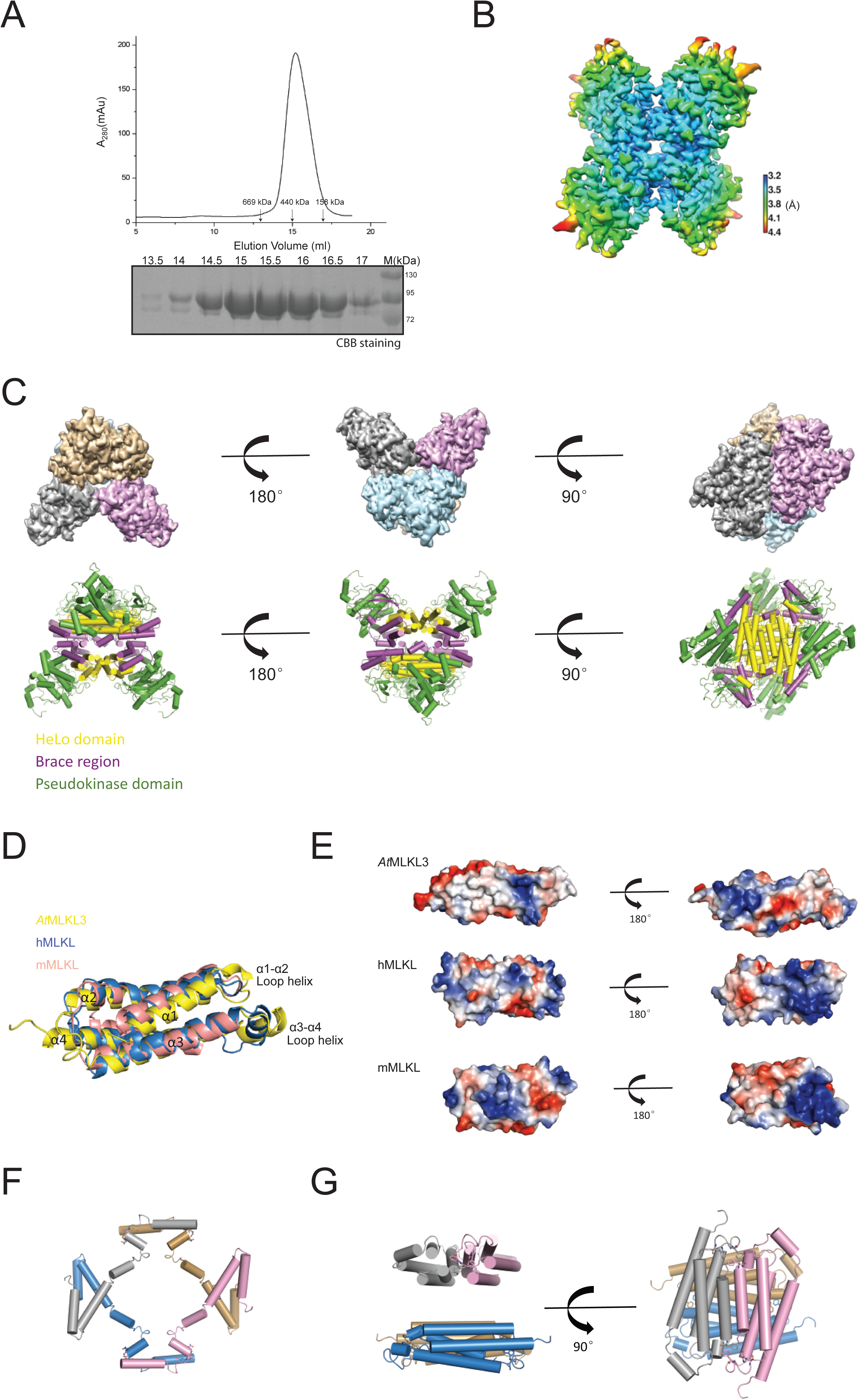
Structure of the *At*MLKL3 tetramer. **A**, Top: Gel filtration profiles of *At*MLKL3. Position of standard molecular weight is indicated by arrow. Bottom: Peak fractions on the top were verified by reducing SDS-PAGE with Coomassie Blue staining. **B**, Final EM density map of the *At*MLKL3 tetramer. Local resolution of the density map is shown on the right with scale indicated by colors, in angstroms. **C**, Top: Final 3D reconstruction of *At*MLKL3 tetramer shown in three orientations. Each monomer of *At*MLKL3 is shown in different colors. Bottom: Cartoon shows the overall structure of *At*MLKL3 tetramer in three orientations. Subdomains of *At*MLKL3 are shown in different colors. **D** Superposition of the HeLo domains of *At*MLKL3 (yellow, PDB ID code 6KA4), human MLKL (blue, PDB ID code 2MSV) and mouse MLKL (Pink, PDB ID code 4BTF). RMSD between *At*MLKL3 and mMLKL:3.745, RMSD between *At*MLKL3 and hMLKL:4.010. **E**, Electrostatic surface of HeLo domains of *At*MLKL3, human MLKL and mouse MLKL in two orientations. **F**, Cartoon shows the brace region of the *At*MLKL3 tetramer. Each monomer of *At*MLKL3 is shown in different colors. **G**, Cartoon shows the HeLo domains of the *At*MLKL3 tetramer in two orientations. Each monomer of *At*MLKL3 is shown in different colors.

The N-terminal HeLo domain of *At*MLKL3 forms a four-helix bundle (Fig. S3C) which superimposed well with the HeLo domains of mouse and human MLKL (mMLKL and hMLKL, respectively; Fig. 2D). This observation supports the idea that *At*MLKL3 is a bona fide homolog of vertebrate MLKLs. Nevertheless, compared to mMLKL, the HeLo domain of *At*MLKL3 packs tightly against its pseudokinase domain (Fig. S3D). The packing is further strengthened by the brace region of *At*MLKL3, which contains a string of five helices that simultaneously interact with the HeLo and pseudokinase domains (Fig. S3C). The HeLo domains and the brace regions form the core of the *At*MLKL3 tetramer, whereas the pseudokinase domains are presented at the apices of the pyramid-like structure (Fig. 2C). Hydrophobic packing of α1 helices (Fig. S3E) from two *At*MLKL3 molecules contributes to formation of a homodimeric *At*MLKL3. In the tetrameric *At*MLKL3, the two α1-mediated homodimers pack perpendicularly to each other (Fig. 2G). Four brace regions, which are positioned nearly in the same plane, are sandwiched between the two homodimers and exclusively mediate homodimer-homodimer interaction (Fig. 2F). The N-terminal halves of the four brace regions form two homodimer pairs, and the C-terminal halves another two pairs (Fig. S3E). The intermolecular interactions lead to further sequestering of the *At*MLKL3 N-terminal HeLo domain. Taken together, our observations indicate that *At*MLKL3 is a structural homolog of vertebrate MLKLs and its N-terminal HeLo domain is sequestered through both intra- and intermolecular interactions.

To determine the role of plant MLKLs in immunity, we challenged combinatorial loss-of-function mutants of *At*MLKLs (Fig. S4) with different microbial pathogens. *Atmlkl* single mutants exhibited increased susceptibility to the obligate biotrophic fungus *Golovinomyces orontii* compared to wild-type plants and, strikingly, the triple mutant was as susceptible as an *eds1* mutant (Fig. 3A), which is hyper-susceptible to a number of pathogens including the obligate biotrophic oomycete *Hyaloperonospora arabidopsidis* (*Hpa*). The immune response restricting *Golovinomyces* growth in Col-0 wild-type plants is not associated with host cell death (*19*). In *A. thaliana*, immunity to *Hpa* isolate Emwa1 is mediated by RPP4, an NLR that requires EDS1 for function (*20*). RPP4-mediated disease resistance was partially compromised in the *Atmlkl* triple mutant, as measured by increased *Hpa* sporangiophore formation on true leaves (Fig. 3B). We did not detect marked differences between the susceptibility of wild-type and mutant plants to hemi-biotrophic *Pseudomonas syringae* pv. *tomato* DC3000 bacteria or the fungal necrotroph *Botrytis cinerea* (Fig. S5AB). Furthermore, disease resistance mediated by RPS2 and cell death mediated by RPM1 NLRs were largely retained in the *Atmlkl123* mutant (Fig. S5CD).

**Fig. 3:**
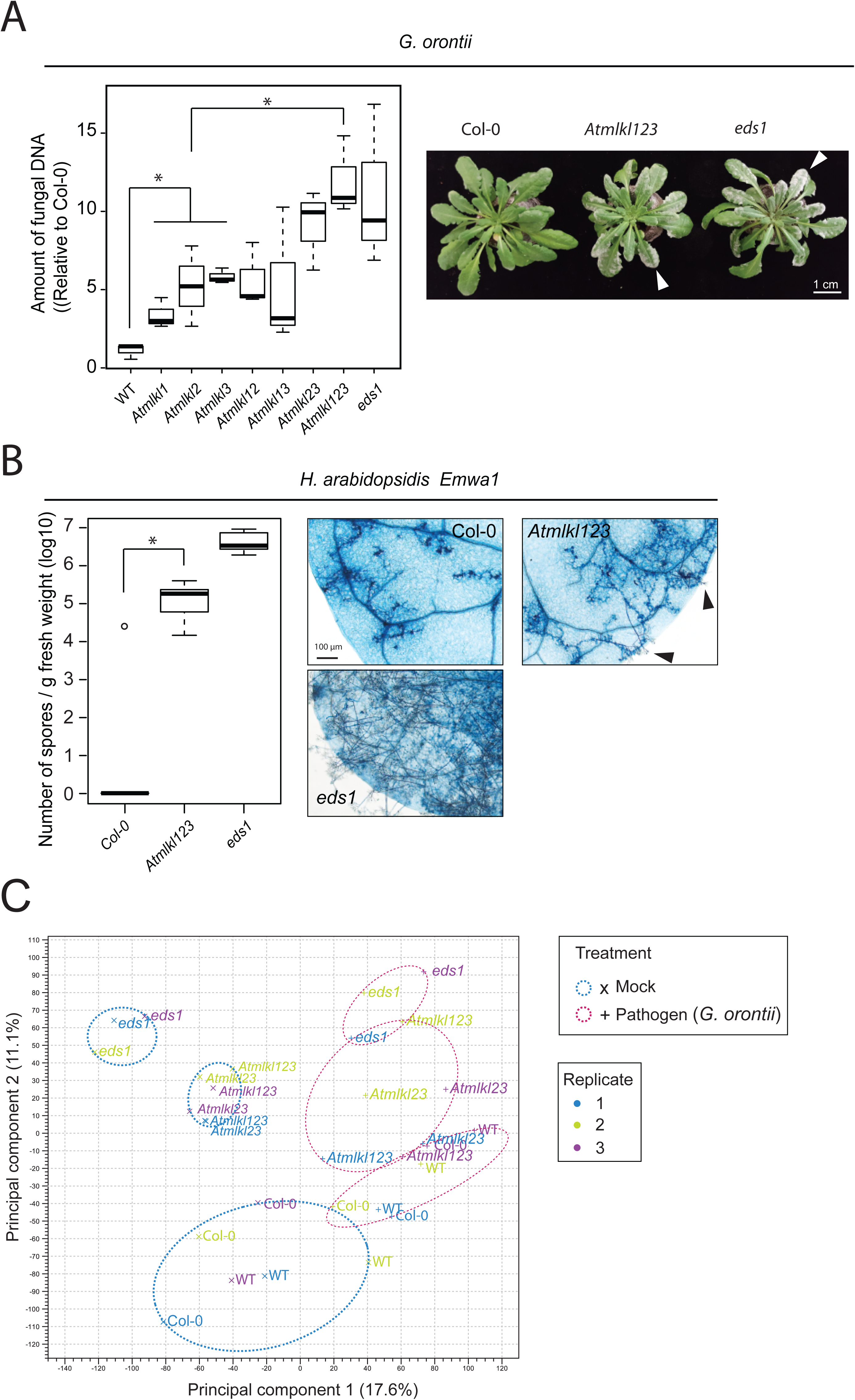
Arabidopsis MLKLs confer resistance to powdery and downy mildew pathogens. **A**, *Atmlkl* mutants are susceptible to the powdery mildew *Golovinomyces orontii* (arrowheads). Quantification of *G. orontii* DNA at seven days after inoculation is relative to the corresponding Col-0 samples. (n=3, Tukey HSD \**p* < 0.05). **B**, The *Atmlkl* triple mutant is susceptible to the downy mildew *Hyaloperonospora arabidopsidis*. The number of *Hpa* spores per gram fresh weight of plants is presented. Data were obtained in three independent experiments, each including two biological replicates (n=6). Asterisks indicate significant differences (Tukey HSD *p* < 0.05). The 1^st^ or 2^nd^ true leaves were stained with trypan blue to visualise hyphae growth. Sporangiophores were formed on the true leaves of the *Atmlkl* triple mutant (black arrowheads). **C**, Principal component analysis (PCA) of RNA-seq data from mock- and pathogen-challenged leaves collected at 48 hours post *G. orontii* inoculation. *WT is a segregant line derived from the cross between *Atmlkl1* and *Atmlkl3*. The *eds1* mutant was used as a susceptible control.

RNA-seq analysis using mock- and *G. orontii*-challenged leaves of wild-type, *Atmlkl23, Atmlkl123* and *eds1* lines showed that the *AtMLKL1* transcript was induced upon pathogen challenge (7.4 fold) and that induction was fully dependent on *EDS1* (Fig. 3C, Fig. S6). Further analysis revealed that *At*MLKLs regulate transcriptional reprogramming both under unchallenged conditions and upon pathogen attack (Fig. 3C). These data imply a specific and partly redundant role of *At*MLKLs in resistance to obligate biotrophic pathogens.

To test whether *At*MLKLs possess cell death activity, possibly regulated by phosphorylation, we introduced single phosphomimetic substitutions at serine residues in the activation loops of *At*MLKLs (Fig. 1B) and expressed in *A. thaliana* leaf protoplasts (see method). Upon overexpression, all wild-type *At*MLKLs elicited cell death, which was as potent as an N-terminal barley NLR cell death module (*21*)(Fig. S7). We found that a phosphomimetic substitution at serine^393^ but not serine^395^ in *At*MLKL1 enhanced its cell-killing activity (Fig. 4A). However, an alanine substitution at serine^393^ did not compromise the cell death activity, suggesting that other serine or threonine residues in the activation loop (Fig. 1B) have a compensatory function. Using transgenic *A. thaliana* lines, we observed that the enhanced susceptibility of *Atmlkl1* and *Atmlkl2* mutants to *G. orontii* was restored by stable transformations of genomic fragments encompassing wild-type *AtMLKL1* and *AtMLKL2*, respectively (Fig. 4B,C and Fig. S8-9). In line with the protoplast assay (Fig. 4A), *At*MLKL1 (S393D) transgenic plants exhibited enhanced resistance to *G. orontii* compared to Col-0 wild type or to plants expressing the transgenes *At*MLKL1 or *At*MLKL1 (S395D) (Fig. 4B,C). Unexpectedly, *At*MLKL1 (S393D) transgenic plants did not exhibit cell death lesions or apparent plant growth retardation indicative of autoimmunity (Fig. 4B, Fig. S8A). Collectively our results suggest that plant MLKL function is in part regulated by activation loop conformation induced by phosphorylation, similar to vertebrate MLKLs (*22*), but *At*MLKL1 activity needed to restrict *G. orontii* growth can be separated from host cell death.

**Fig. 4:**
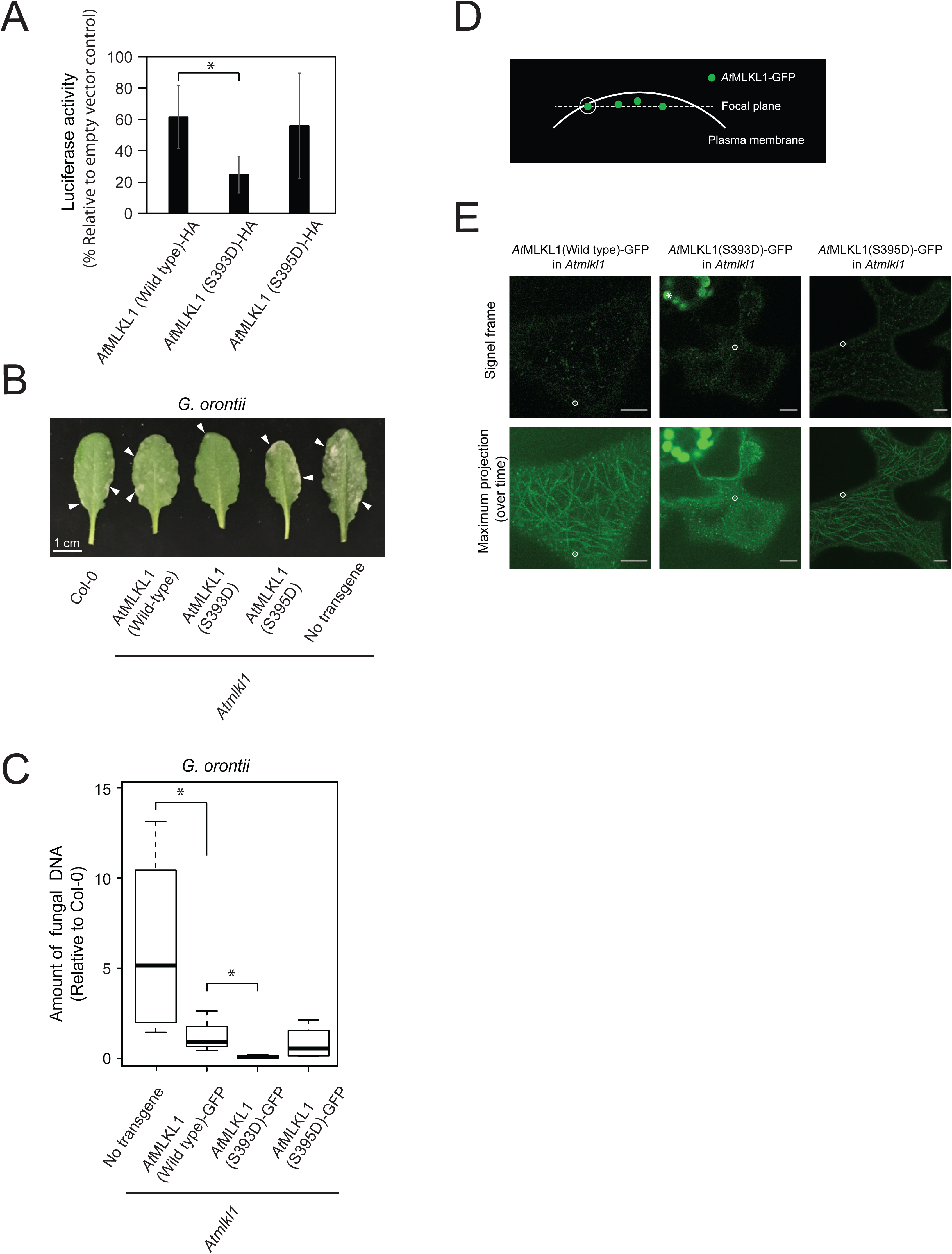
Cell death activity, disease resistance and subcellular localization of phosphomimetic *At*MLKL1 variants. **A**, A phosphomimetic substitution in *At*MLKL1 elicited enhanced cell death in *Arabidopsis* mesophyll protoplasts. Luciferase and *At*MLKL1 expression constructs were co-transfected into protoplasts and luciferase activity was measured as a proxy of cell viability at 16 hours post transfection. The positions of serine-to-aspartate residues are indicated in Fig. 1B. The C-terminally HA-tagged variants were expressed under the control of the constitutive cauliflower mosaic virus 35S promoter. (n=3, Tukey HSD \**p* < 0.01). **B**, Macroscopic phenotype of transgenic *Arabidopsis* expressing phosphomimetic variants of *At*MLKL1 in response to *G. orontii* (arrowheads). Plants were photographed seven days after pathogen challenge. **C**, Quantification of *G. orontii* DNA in infected leaves of transgenic plants at 14 days after pathogen challenge (n=4, Tukey HSD \**p* < 0.05). **D**, Schematic diagram of the confocal images shown in **B**. The white circle indicates GFP signal proximal to the PM. **E**, Confocal images of the abaxial epidermis of the *Arabidopsis* transgenic lines expressing phosphomimetic and wild-type variants of *At*MLKL1. GFP signals indicated by white circles were immobile in the examined time period. The corresponding movies are available as supplemental movies 1-3. The asterisk indicates plastidial autofluorescence. Scale bars = 5 µm.

We next examined whether the *At*MLKL N-terminal HeLo domain contributes to its cell death activity, as in animals (*8, 12, 14, 23*). Taking advantage of a chemically enforced oligomerization system (Fig. S10, see method), we found that expression of the HeLo domains with the brace region of *At*MLKL1 and *At*MLKL3 was sufficient for cytotoxic activity and this activity was further enhanced upon oligomerization (Fig. S10D). This finding mirrors the activity of the HeLo domains with the brace region of MLKL (*14*) and is consistent with our structure-based hypothesis that the full-length MLKL tetramer with buried HeLo domains represents an inactive form (Fig. 2).

As animal MLKL translocates to the PM upon activation (*10–14*) we examined whether GFP-tagged *At*MLKL variants associate with the PM using time-resolved confocal microscopy. We detected mobile punctate signals in the cytoplasmic space for all *At*MLKL1 variants (Fig. 4DE). Reduced mobility for a fraction of *At*MLKL1-GFP signals when proximal to the PM (Fig. 4E) is consistent with PM association. Intriguingly, maximum projection of the time-lapse images revealed that *At*MLKL1 (wild type)-GFP, the S395D mutant moved along filamentous structures (Fig. 4E, Supplemental movies 1 and 3; Fig. S11A). These filamentous structures specifically colocalized with the microtubule marker (*24*) mCherry-MAP4 (Fig. S11B-C), and association was detectable upon *G. orontii* invasion (Fig. S12). Intriguingly, no filamentous structures were detected in the S393D phosphomimetic line despite the presence of a microtubule array (Fig. S13). These data suggest that specific phosphorylation modulates the intracellular localization and activities of *At*MLKL1 via cytoskeletal association.

The plant MLKL family described here is a first example of a non-receptor immunity component consisting of multiple domains that is structurally shared between animal and plants. MLKLs in these two kingdoms are likely products of convergent evolution because their exon-intron structures are unrelated (∼4 and 11 exons in *At*MLKLs and hMLKL, respectively). Our work serves as a template to test structural predictions implicating the presence of a HeLo-domain fold in a number of plant modular proteins, including the ADR1 family of NLRs (*4, 25–27*). Despite the cell death activity of a putative HeLo-domain of ADR1 (*28*), this family confers disease resistance without apparent host cell death (*29*). This resembles *At*MLKL-mediated defense in wild-type plants to *G. orontii* without host cell death, although *At*MLKLs have the capacity to elicit cell death. These results raise the question of whether host defence and cell death functions can be further disentangled in animal necroptosis signalling.

## Acknowledgments

We thank the Max Planck Genome Centre Cologne for RNA-Seq and Petra Köchner, Sabine Haigis and Makoto Yoshikawa-Maekawa for technical assistance and Neysan Donnelly for editing the manuscript. We also thank Hirofumi Nakagami, Ton Timmers, Hamid Kashkar and Manolis Pasparakis for helpful suggestions. We also thank Takashi Hashimoto and Stefanie Sprunck for the mcherry-MAP4 and tagRFP-T-Lifeact vectors, respectively.

## Data and materials availability

All data underlying the study are deposited at Protein Data bank (PDB): https://www.rcsb.org/) or Gene Expression Omnibus (GEO) database (https://www.ncbi.nlm.nih.gov/geo/).

## Author contributions

T.M. conceptualized the project; L.M., M.H., X.Z., R.T.N., L.B.K., I.M.L.S., F.J., V.K., D.L., K.H., T.M. performed the investigations. J.P. J.M.M., P.S-L. J.C. T.M. validated the data, J.P. J.M.M., P.S-L. J.C. T.M. supervised the work, J.C. and T.M. wrote the original draft of the manuscript, J.P. P.S-L., J.M.M. reviewed and edited the manuscript.

## Data and materials availability

All data needed to replicate the work are deposited in the Protein Data Bank (PDB) or Gene Expression Omnibus (GEO) database. Plants and plasmids described in the manuscript are available upon request.

## Methods

### Plant material and growth conditions

The Arabidopsis thaliana (L.) Heynh. ecotype Columbia (Col-0) was used in this study. The T-DNA insertional mutants(1, 2) (SALK_041569c (AtMLKL1), SALK_124412c (AtMLKL2) and GABI_491E02 (AtMLKL3) were obtained from the Nottingham Arabidopsis Stock Centre (NASC). Double and triple mutants of Atmlkl were generated by crossing the T-DNA insertion lines. A segregant line derived from the cross between the Atmlkl1 and Atmlkl3 mutants were used as a wild-type line in addition to Col-0 wild-type. Each genotype was confirmed by PCR. The eds1-2 mutant was described previously (3).

*The transgenic lines expressing AtMLKL1 variants or AtMLKL2 fused to a monomeric green fluorescence protein* were established in *Atmlkl1* and *Atmlkl2* mutant backgrounds, respectively. Genomic fragments including coding region and native *cis*-regulatory sequence were amplified by PCR from Col-0 genomic DNA and cloned into pENTR/D-TOPO (Thermo Fisher Scientific, Waltham, MA, USA). The phosphomimetic substitutions were introduced using the QuikChange Lightning site-directed mutagenesis kit (Agilent Technologies, Santa Clara, CA, USA). Resulting entry vectors were transferred into pGWB550(*4*) using LR clonase II (Thermo Fisher Scientific). Plants were transformed by the floral dip method (*5*) with *Agrobacterium tumefaciens* strain GV3101 harbouring pMP90RK (*6*). Plant growth conditions were described previously (*7*). Primer sequences for genotyping and plasmid construction are listed in Table S3.

### Sequence analysis of plant and animal MLKLs

Sequence similarity within the animal and plant families was established by the generalized profile method (*8*). Sequences were aligned by the L-INS-I method of the MAFFT alignment software (*9*), followed by minor manual editing of ambiguously aligned regions. Sequence similarity between the animal and plant MLKL families was established by Hidden Markov Model (HMM)-to-HMM comparison using the HHSEARCH package (*10*). The 99,696 orthogroups (OGs) among 52 plant species have been established recently (*11*). An OG containing *At*MLKLs was used for the neighbor-net analysis of codon-aligned nucleotide sequence as descried previously (*12*). The sequence from the papaya genome was excluded in this study.

### Protein expression and purification

Full length *At*MLKL3 (residues 1-701) with an engineered C-terminal 6×His tag was generated by standard PCR-based cloning strategy and its identity was confirmed by sequencing. The protein was expressed in sf21 insect cells using the vector pFastBac 1 (Invitrogen). One litre of cells (2.5×10^6^ cells ml^−1^, medium from Expression Sysems) was infected with 20 ml baculovirus at 28°C. After growth at 28°C for 48 hours, the cells were harvested, re-suspended in the buffer containing 50 mM Tris-HCl pH 8.0 and 300 mM NaCl, and lysed by sonication. The soluble fraction was purified from the cell lysate using Ni2+-nitrilotriacetate affinity resin (Ni-NTA, Qiagen). The protein was then further purified by further purified by gel filtration (Superose 6, 10/30; GE Healthcare). For cryo-EM investigation, the purified protein was concentrated to about 0.3 mg/mL in buffer containing 50 mM Tris-HCl pH 8.0, 300 mM NaCl and 3 mM DTT.

The construct of full length *At*MLKL2 (residues 1-711) with N-terminal GST tag was cloned into the pGEX-6P-1 vector (GE Healthcare), and was expressed in *Escherichia coli* strain BL21(DE3; Novagen) at 16 °C. After isopropyl-β-D-thiogalactopyranoside (IPTG; Sigma) induction for 12 h, cells were harvested and re-suspended in buffer containing 50 mM Tris-HCl pH 8.0 and 300 mM NaCl, and lysed by sonication. The soluble fraction was purified from the cell lysate using Glutathione Sepharose 4B beads (Invitrogen). The proteins were then digested with PreScission protease (GE Healthcare) to remove the GST tag and further purified by gel filtration (Superose 6, 10/30; GE Healthcare). For cryo-EM investigation, the purified protein was concentrated to about 0.3 mg/mL in buffer containing 50 mM Tris-HCl pH 8.0, 300 mM NaCl and 3 mM DTT.

### Cryo-EM sample preparation and data collection

For cryo-EM analysis, an aliquot of 3.5 µl *At*MLKL2 or *At*MLKL3 protein was applied to a holey carbon grids (Quantifoil Cu 1.2/1.3, 200 mesh) glow-discharged (Harrick Plasma) with a middle force for 30 s after evaluating for 2 min. The grids were blotted by a pair of 55 mm filter papers (TED PELLA, INC.) for 3-3.5 s at 8 °C with 100% humidity and flash-frozen in liquid ethane using a FEI Vitrobot Marked IV. Cryo-EM data were collected on Titan Krios electron microscope operated at 300 kV and a Gatan K2 Summit direct electron detection camera (Gatan) using eTas. Micrographs were recorded in super-resolution mode at a nominal magnification of 22500 ×, resulting in a physical pixel size of 1.30654 Å per pixel. Defocus values varied from −1.7 µm to −2.3 µm for data set. The dose rate was 10.6 electron per pixel per second.

Exposures of 8.0 s were dose-fractionated into 32 sub-frames, leading to a total accumulated dose of 50 electrons per Å2. In total, two batches of data were collected, one for *At*MLKL3 and another for *At*MLKL2.

### Image processing and 3D reconstruction

A total of 1,434 and 1,828 raw images stacks of *At*MLKL3 and *At*MLKL2 acquired under super-resolution mode, were 2x binned processed using MotionCor2 (*13*), generating aligned, dose-weighted and summed micrographs in a pixel size of 1.30654 Å per pixel. CTFFIND4 (*14*) was used to estimate the contrast transfer function (CTF) parameters. After the removal of bad micrographs via the evaluation of CTF parameters, remaining images were processed in RELION (*15*). Approximate 2,000 particles were manually picked and 2D-classified to generate templates for auto-picking. 983,779 and 1,135,463 autopicked particles for *At*MLKL3 and *At*MLKL2 respectively were then used for reference-free 2D classification, to remove contaminants and bad particles. The left good particles were subjected to 3D classification using initial 3D reference model obtained by *ab initio* calculation from Relion3.0. Particles from good classes that possess density map with better overall structure features were selected for the 3D refinement. The final 3D refinement using D2 symmetry resulted in reconstructions of *At*MLKL3 and *At*MLKL2 tetramer at resolution of 3.4 Å and 4.1 Å, the resolutions were determined by gold-standard Fourier shell correlation. Local resolution distribution was evaluated using Relion.

### Model building and refinement

EM density map of *At*MLKL3 was used to build the model de novo, as the overall resolution of map density was efficient to display side chains. The model of *At*MLKL3 was manually built into the density in COOT(*16*), and was refined against the EM map by PHENIX(*17*) in real space with secondary structure and geometry restraints. The refined *At*MLKL3 model was docked into the density of *At*MLKL2. The sequence of the docked *At*MLKL3 model was changed to that of *At*MLKL2 under COOT and the *At*MLKL2 model with corrected sequence was subjected to refinement by PHENIX. The C-terminal serine-rich region of *At*MLKL2 or *At*MLKL3 is much less well defined in the density and is not included in the models. Final model of *At*MLKL3 and *At*MLKL2 was validated using MolProbity and EMRinger in PHENIX package. The structures of human MLKL (*18*) and mouse MLKL (*19*) were used for the superposition of the HeLo domains as shown in Fig. 2D.Table S2 summarized the model statistics.

### RNA sequencing

Total mRNA from leaves was obtained at 48 hours after challenge with conidia of *Golovinomyces orontii* using the RNeasy plant mini kit (Qiagen). RNA sequencing (RNA-Seq) libraries were prepared by the Max Planck Genome Centre Cologne (Cologne, Germany) using the Illumina TruSeq stranded RNA sample preparation kit (Illumina). The resulting libraries were subjected to 150-bp single-end sequencing using the Illumina HiSeq3000 (Illumina). Mapping of sequenced reads onto the *Arabidopsis thaliana* gene model (TAIR10), principal component analysis, and differential gene expression analysis were performed in the CLC Genomics Workbench (Qiagen, ver. 10.1.2) using the tool ‘RNA-Seq with the default parameter setting. The data derived from Col-0 and WT were pooled as data of wild-type lines in the analysis. The heat map of 93 genes differentially expressed between wild type lines and *Atmlkl* mutants (|log2FC|>1 and false discovery rate (FDR) < 0.05) were generated using the R package (ver. 1.08) with the pheatmap function. Gene ontology enrichment analysis was performed using the PANTHER classification system (http://pantherdb.org/) with default settings for *Arabidopsis thaliana*. The RNA-Seq data generated in this study have been deposited in the Gene Expression Omnibus (GEO) database under accession number GSE129011.

### Transient gene expression in *Arabidopsis* protoplasts

Isolation, transfection and luciferase activity measurement of Arabidopsis protoplasts were performed as described previously (*20*). Protoplasts were isolated from the leaves of two-week-old *Arabidopsis* plants grown in liquid 1 x Murashige and Skoog medium. Coding sequences (CDS) of *AtMLKL1* and *AtMLKL2* without stop codons were initially cloned into pENTR/D-TOPO (Thermo Fisher Scientific). The CDS of *AtMLKL3* without a stop codon was chemically synthesized and cloned into pENTR221 (Thermo Fisher Scientific). Two synonymous substitutions (G1371T and A1413G) were introduced into *AtMLKL3* CDS to remove restriction sites that hamper the DNA synthesis. Entry clones were transferred into the gateway cloning-compatible pAMPAT-GW-mYFP, pAMPAT-GW-3xHA, or pAMPAT-GW expression vectors (*21*), which are derivatives of pAMPAT-MCS (accession number: AY436765). Primers sequences of for plasmid construction are listed in Table S3. pENTR-tagRFP-T-Lifeact (*22*) was transferred into pAMPAT-GW. The expression vectors for *Hv*MLA(1-160aa), MAP4 and Tub6 were described previously (*21, 23, 24*).

### Protoplast viability assay

Following protoplast transfection and regeneration, Evans blue dye dissolved in water was added to the samples to a final concentration of 0.04% (w/v). The stained cells were examined under a standard microscope. For luciferase-based viability assay luciferase and *At*MLKL expression constructs were co-transfected into protoplasts and luciferase activity was measured as a proxy of cell viability.

### Induced oligomerization

Two domains of FKBP (F36V) tagged with HA without N-myristoylation signal were PCR amplified from pC4M-FV2E plasmid (ARIAD, Cambridge, MA, USA). *Nco*I and *Hind*III restriction sites were added to the 5’ end of forward and reverse primers, respectively. The digested PCR fragment with *Nco*I and *Hind*III were ligated into the same restriction sites present between the attR2 and the terminator sequences of pAMPAT-GW expression vector. The resulting vector is named pAMPAT-GW-FV2E-HA. Coding regions corresponding to the N**-** and C**-**terminal luciferase fragments (nLUC and cLUC) were PCR amplified from nLUC and cLUC expression vectors (*25*) and cloned into pENTR/D-TOPO. Respective entry clones were transferred into pAMPAT-GW-FV2E-HA. B/B homodimerizer (also known as AP20187 ligand) were purchased from TakaraBio, Japan. After transfection, protoplasts in incubation buffer (i.e. WI solution (*26*)) were separated into two tubes and added the same amount of incubation buffer supplemented with B/B homodimerizer (250 nM at the final concentration) or ethanol as solvent control. Primer sequences for the plasmid construction are listed in Table S2. We were able to reconstitute luciferase activity of co-expressed N- and C-terminal halves of luciferase fused to 2xDmrB domains in a ligand-specific manner (Fig. S8b) and the ligand itself did not affect the luciferase reporter assay in protoplasts (Fig. S8c).

### Pathogen infection assay

The *G. orontii* infection assay was performed as described previously (*27*). *G. orontii* DNA was quantified by qPCR at indicated time points after inoculation of conidia and normalised using the amount of plant specific gene (AT3G21215). The *Hpa* infection assay was performed as previously described (*28*). Lactophenol-trypan blue staining was described previously (*29*). The *B. cinerea* strain B05.10 was used in this study. Droplet inoculation of six-week-old plants was performed as described previously (*30*), except that 2 µl of the spore solution were used on each side of the leaf and two leaves of similar age were used per plant. *B. cinerea* DNA was quantified by qPCR as previously described (*31*). *Pseudomonas syringae* pv. *tomato* (*Pst*) DC3000 and *Pst* DC3000 expressing *AvrRpt2* or *AvrRpm1* were used in this study. *Pst* growth assays and ion leakage measurement following bacterial infiltration were performed as described previously (*7*).

### Immunoblot assays

Primary antibodies were monoclonal antibodies from mouse: α-GFP (JL-8, 1:5000, Takara, Shiga, Japan) or rat: α-HA (3F10, 1:1000, Sigma-Aldrich, St. Louis, MO, USA). Goat α-mouse IgG-HRP (1:10000, Santa Cruz Biotechnology, Dallas, TX, USA) or goat α-rat IgG-HRP (1:10000, Sigma-Aldrich) were used as secondary antibodies. The detailed procedure is described in (*7*).

### Biolistic transient gene expression

Biolistic delivery of plasmid DNA into the abaxial epidermis of leaves was essentially performed as described previously (*32*). Leaves were detached immediately before bombardment and the bombarded leaves were transferred to 1% agar plates supplemented with 85 µM benzimidazole and incubated at 20°C for 15 h before confocal microscopy.

### Confocal microscopy

Transfected protoplasts in a chamber slide (Nunc Lab-Tek, Thermo Fisher Scientific) with incubation buffer (i.e. WI solution, (*26*)) or 2-5 mm leaf discs prepared form rosette leave of 4-5-week-old plants were observed under a confocal microscope (LSM880, Carl Zeiss) equipped with a 40X water-immersion and a 63X oil-immersion objective. Lambda stack images were obtained for spectral imaging. Images were analyzed and processed with ZEN Software (Carl Zeiss) and ImageJ (NIH). In Fig. 4E, confocal images were acquired over time (for wild type, 124 seconds; for S393D, 194 seconds; for S395D, 166 seconds) and used for maximum intensity projection (bottom panels). Representative single frame images are shown (top panels).

## Supplemental data

**Supplemental Fig. 1:**
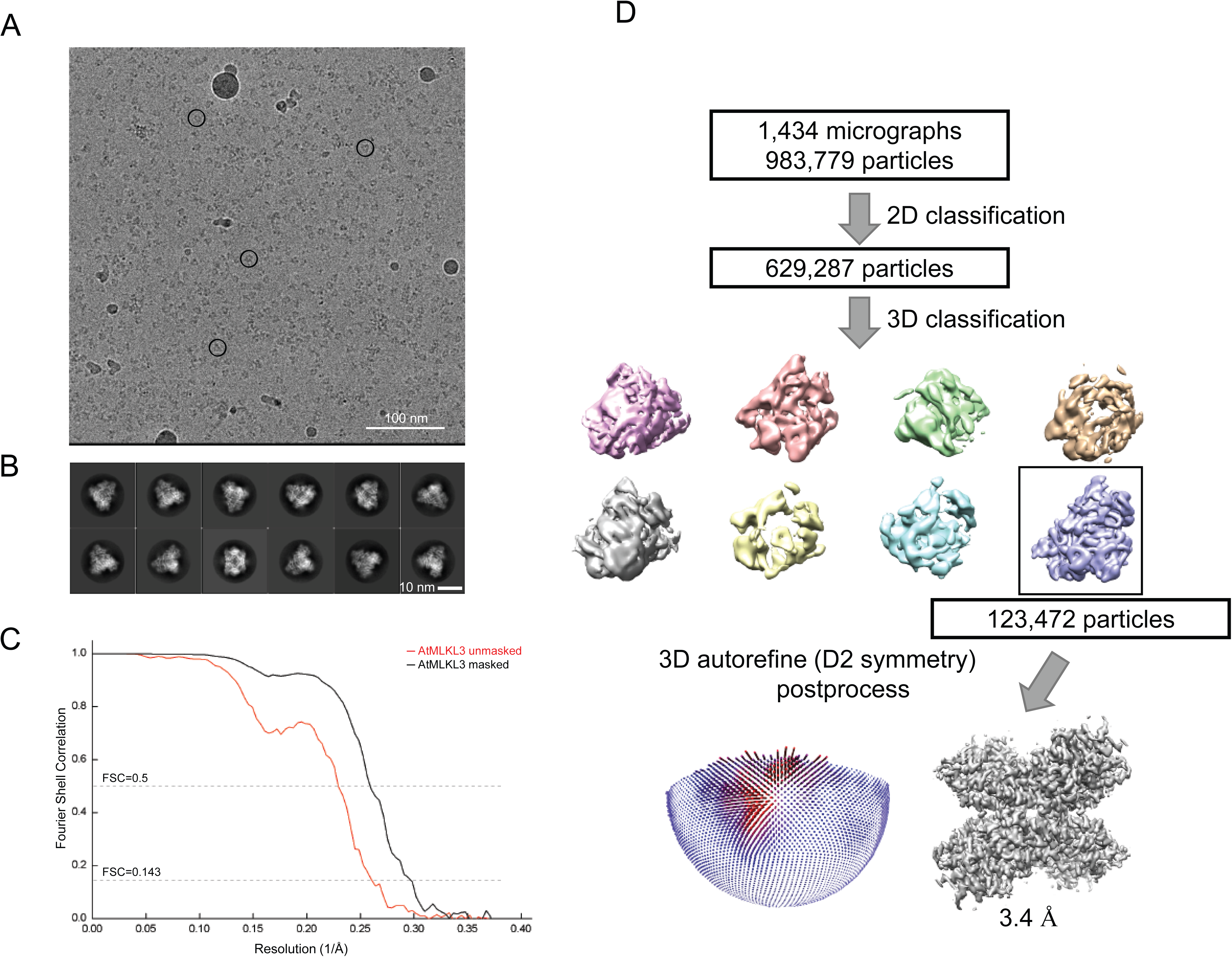
3D reconstruction of *At*MLKL3 tetramer. **A**, Representative cryo-EM image of *At*MLKL3 tetramer. Representative particles are indicated with black circles. **B**, Representative top and side views of 2D class averages of *At*MLKL3 tetramer. **C**, Fourier shell correlation (FSC) curves at 0.5 and 0.143 of the 3D reconstruction of *At*MLKL3 tetramer. **D**, Flowchart representing cryo-EM data processing and 3D reconstruction of *At*MLKL3 tetramer.

**Supplemental Fig. 2:**
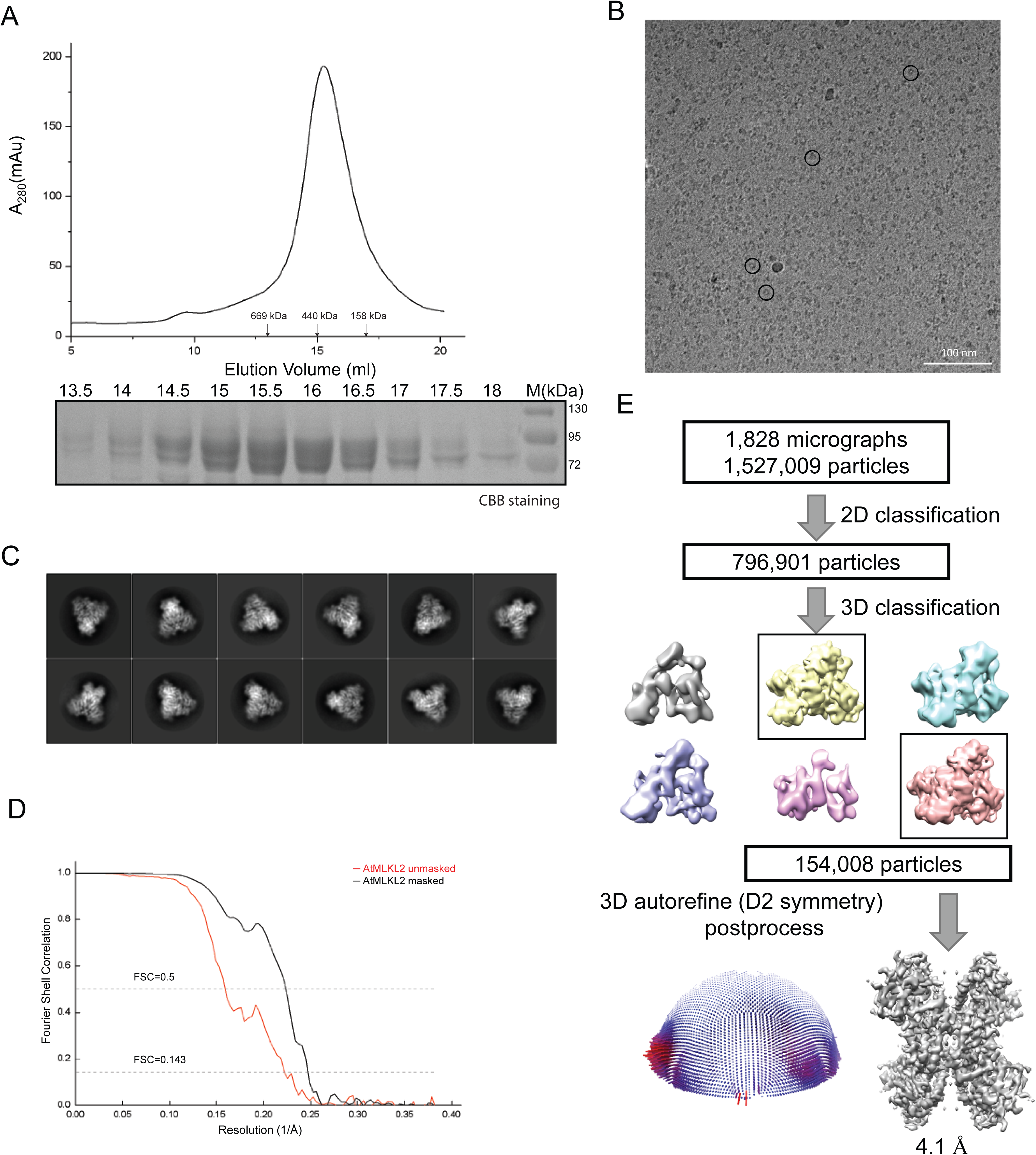
Purification and 3D reconstruction of *At*MLKL2 tetramer. **A**, Top: Gel filtration profile of *At*MLKL2 protein. Position of standard molecular weight is indicated by arrow. Bottom: Peak fractions in the top were verified by SDS-PAGE with Coomassie Blue staining. **B**, Representative cryo-EM images of the *At*MLKL2 tetramer. Representative particles are indicated with black circles. **C**, Representative top and side views of 2D class averages of *At*MLKL2 tetramer. **D**, Fourier shell correlation (FSC) curves at 0.5 and 0.143 of the 3D reconstruction of *At*MLKL2 tetramer. **E**, Flowchart representing cryo-EM data processing and 3D reconstruction of *At*MLKL2 tetramer.

**Supplemental Fig. 3:**
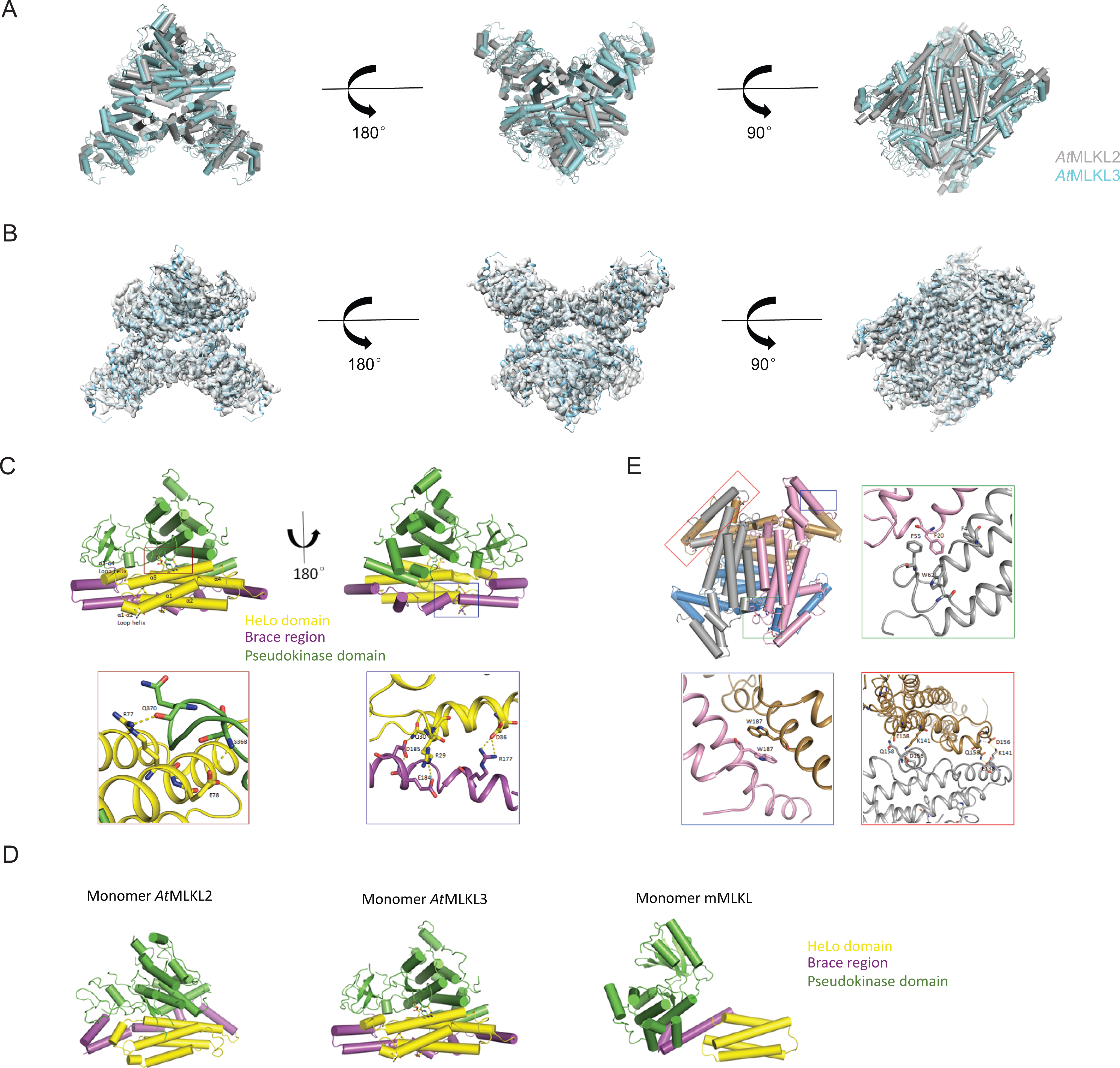
Inter- and intra-domain interactions of *At*MLKL3 tetramer and structural comparison of the *At*MLKL2 and *At*MLKL3. **A,** Superposition of *At*MLKL2 (grey) and *At*MLKL3 (blue). RMSD between monomer of *At*MLKL2 and *At*MLKL3:0.985. **B,** The sequence of *At*MLKL3 model (blue) was docked into *At*MLKL2 map (gray). **C**, Top: Cartoon representation of *At*MLKL3 monomer in two orientations. Subdomains of *At*MLKL3 are shown in different colors, the interacting regions between domains are highlighted with open frames. Bottom left: Detailed interactions of HeLo domain and pseudokinase domain for the red-framed region. Bottom right: Detailed interactions of HeLo domain and brace region for the blue-framed region. **D**, Structural comparison of the monomer *At*MLKL2 (left), *At*MLKL3 (middle, PDB ID code 6KA4) and *m*MLKL (right, PDB ID code 4BTF). **E,** Top left: Cartoon showing HeLo domain and brace region of *At*MLKL3 tetramer. Top right: Detailed interactions of HeLo domains for the green-framed region. Bottom left: Detailed C-terminal interactions of brace regions for the blue-framed region. Bottom right: Detailed N-terminal interactions of brace regions for the blue-framed region.

**Supplemental Fig. 4:**
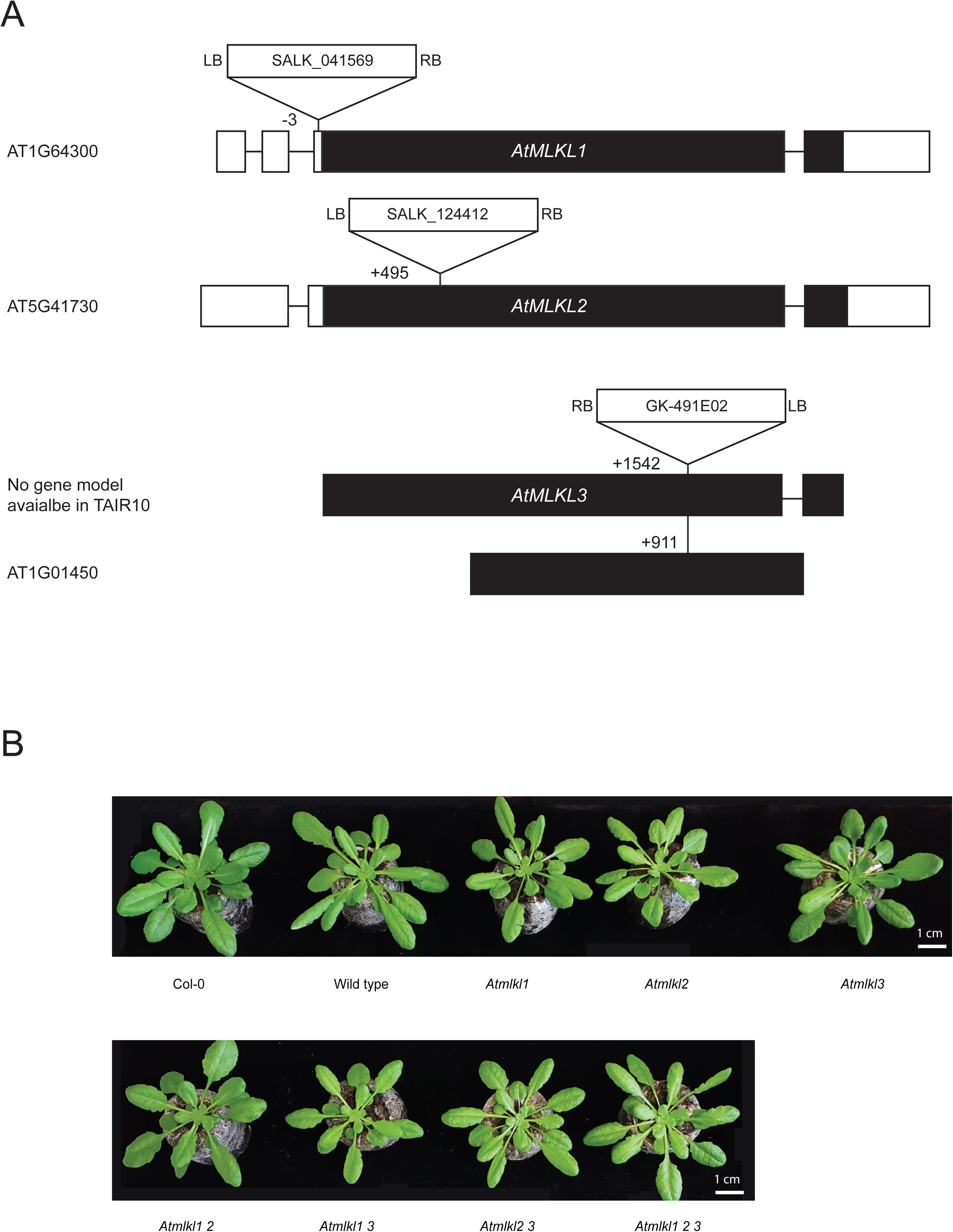
*Atmlkl* mutants exhibit no obvious growth defect. **A,** The Arabidopsis *MLKL* genes and T-DNA insertion sites of the *Atmlkl* mutants. White and black boxes indicate non-coding and coding exons, respectively. A gene model for *AtMLKL3* was deduced from other plant *MLKL* structures. **B**, Representative images of four-week-old plants of Col-0, wild type*, single mutants of *Atmlkl1*, *Atmlkl2*, *Atmlkl3*, double mutants *Atmlkl1 2*, *Atmlkl1 3*, *Atmlkl2 3*, and the triple mutant *Atmlkl1 2 3*. *The wild type is a segregant line derived from the cross between *Atmlkl1* and *Atmlkl3*. The plants were initially grown on Murashige and Skoog-agar plates for two weeks and subsequently transferred to Jiffy pots rehydrated in water with a fertilizer. Plants were grown for an additional two weeks under short-day conditions.

**Supplemental Fig. 5:**
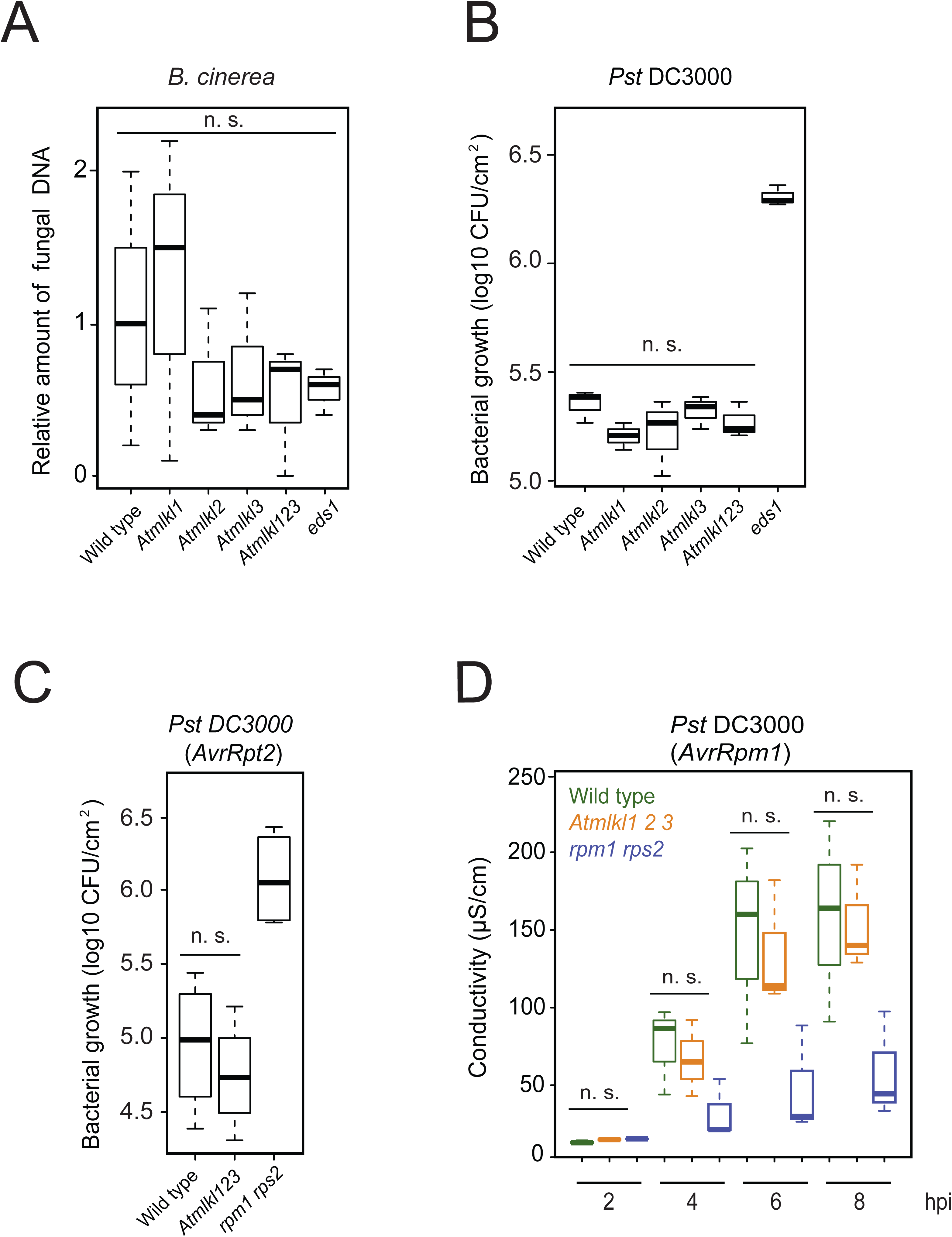
*Arabidopsis* MLKLs do not confer resistance to *Botrytis cinerea and Pseudomonas syringae* DC3000. **A**, *Botrytis cinerea* DNA was quantified by qPCR at three days after spore inoculation and normalized using the amount of plant specific gene (see method). Amounts are presented relative to the corresponding Col-0 samples. The wild type is a segregant line derived from the cross between *Atmlkl1* and *Atmlkl3* (see Fig. S4). Data were obtained in three independent experiments (n=3). **B**, Log_10_-transformed colony forming units of *Pseudomonas syringae* DC3000 per cm^2^ of *A. thaliana* leaves at two days after pathogen infiltration. The *eds1* mutant was used as a susceptible control. Data were obtained in three independent experiments (n=3). **C**, Log_10_-transformed colony forming units of *P. syringae* DC3000 expressing AvrRpt2 per cm^2^ of *A. thaliana* leaves at three days after pathogen infiltration. The *rpm1 rps2* mutant was used as a susceptible control. Data were obtained in three independent experiments (n=3). **D**, Ion leakage assay in *A. thaliana* leaves upon infiltration with *P. syringae* DC3000 expressing *AvrRpm1*. Samples were collected 30 min post infiltration with the bacterial suspension (OD600=0.1). The *rpm1 rps2* mutant was used as a negative control. Data were obtained in three independent experiments (n=3). n.s. not significant.

**Supplemental Fig. 6:**
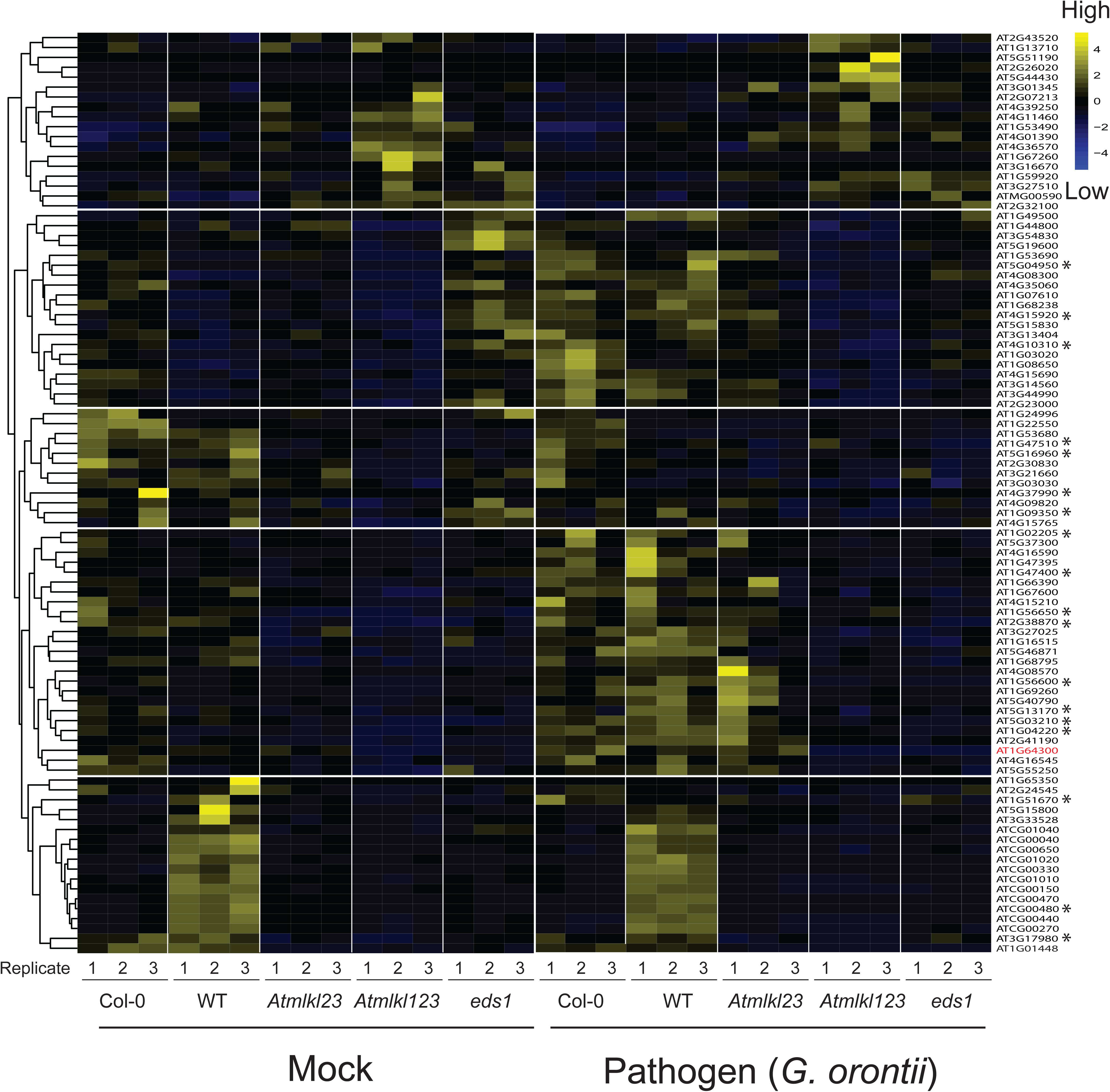
Heat map of 93 genes differentially expressed in *Atmlkl123* in comparison to wild type. The gene ontology (GO) term biotic or abiotic stresses (GO:0006950, asterisks) was overrepresented in the down-regulated transcripts in *Atmlkl123* compared to wild type lines. Mock- and pathogen-challenged leaves were collected at 48 hours post *G. orontii* inoculation. *WT is a segregant line derived from the cross between *Atmlkl1* and *Atmlkl3*.

**Supplemental Fig. 7:**
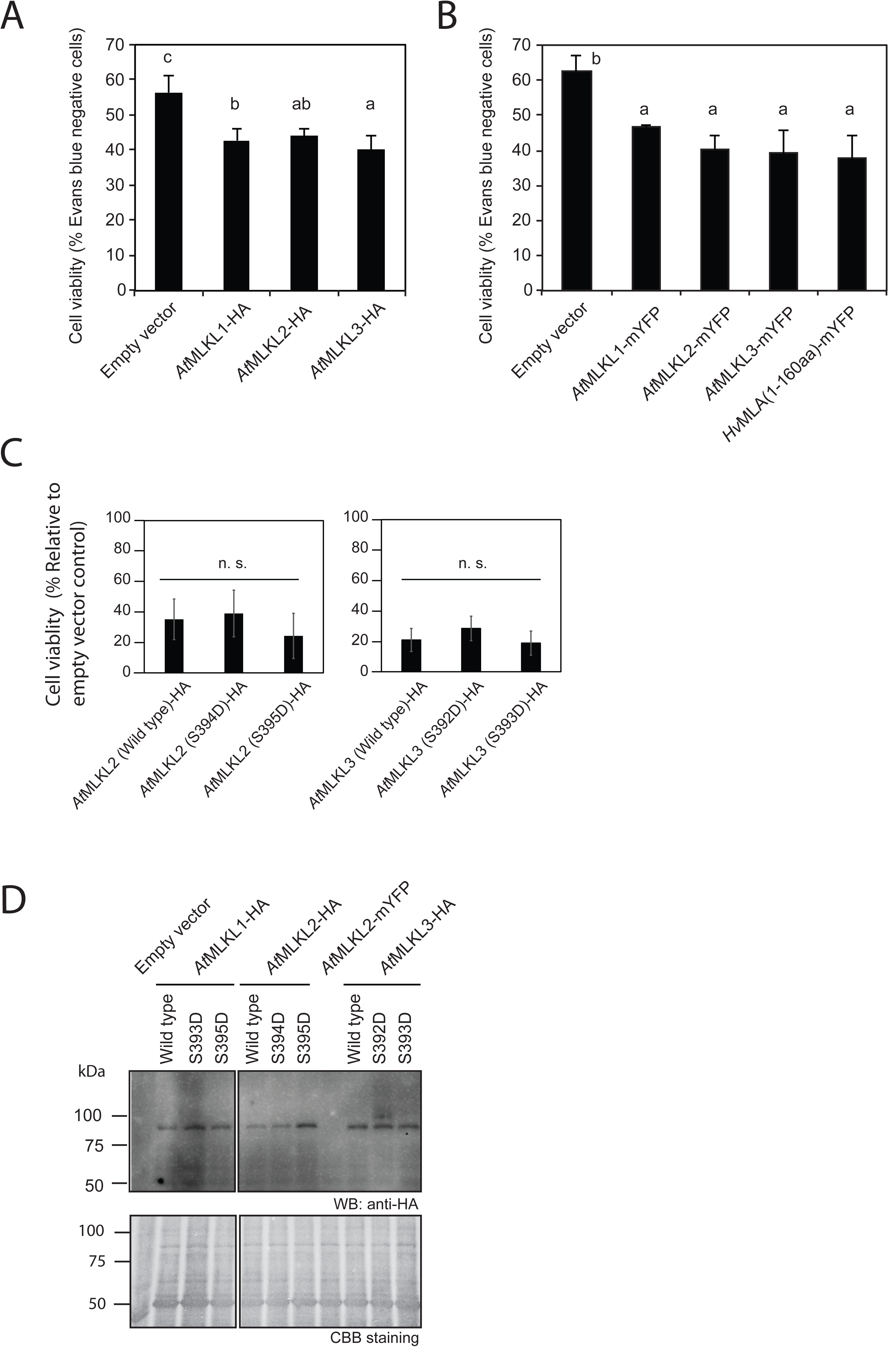
*At*MLKLs are capable of eliciting cell death in *Arabidopsis* mesophyll protoplasts. **A**, C-terminally HA-tagged and **B**, C-terminally mYFP-tagged *At*MLKLs were expressed under the constitutive cauliflower mosaic virus 35S promoter. Expression constructs for *At*MLKLs were transfected into protoplasts and cells were stained with Evans blue at 16 hours post transfection. Unstained cells (Evans blue-negative cells) were counted as living cells. For **B**, The N-terminal signalling region of barley NLR protein (*Hv*MLA_1-160_: (*7*) was used as a positive control. Data were obtained in three independent transfections (n=3) and different letters indicate statistically significant differences (Tukey HSD, *p* < 0.01). **C**. Phosphomimetic substitutions in *At*MLKL2 or *At*MLKL3 do not alter cell death in *Arabidopsis* mesophyll protoplasts. Expression constructs for luciferase and each of the *At*MLKL variants were co-transfected into protoplasts and luciferase activity was measured as a proxy for cell viability at 16 hours post transfection. The positions of serine-to-aspartate substitutions are indicated in Fig. 1B. The C-terminally HA-tagged variants were expressed under the 35S promoter. Data were obtained in three independent transfections (n=3). n.s. not significant. **D**, Western blot analysis of expression of C-terminally HA-tagged *At*MLKLs in *Arabidopsis* mesophyll protoplasts. Total protein extracts were collected at seven hours post transfection. Samples that were transfected with the corresponding empty vector or the expression construct for *At*MLKL2-mYFP were used as negative control.

**Supplemental Fig. 8:**
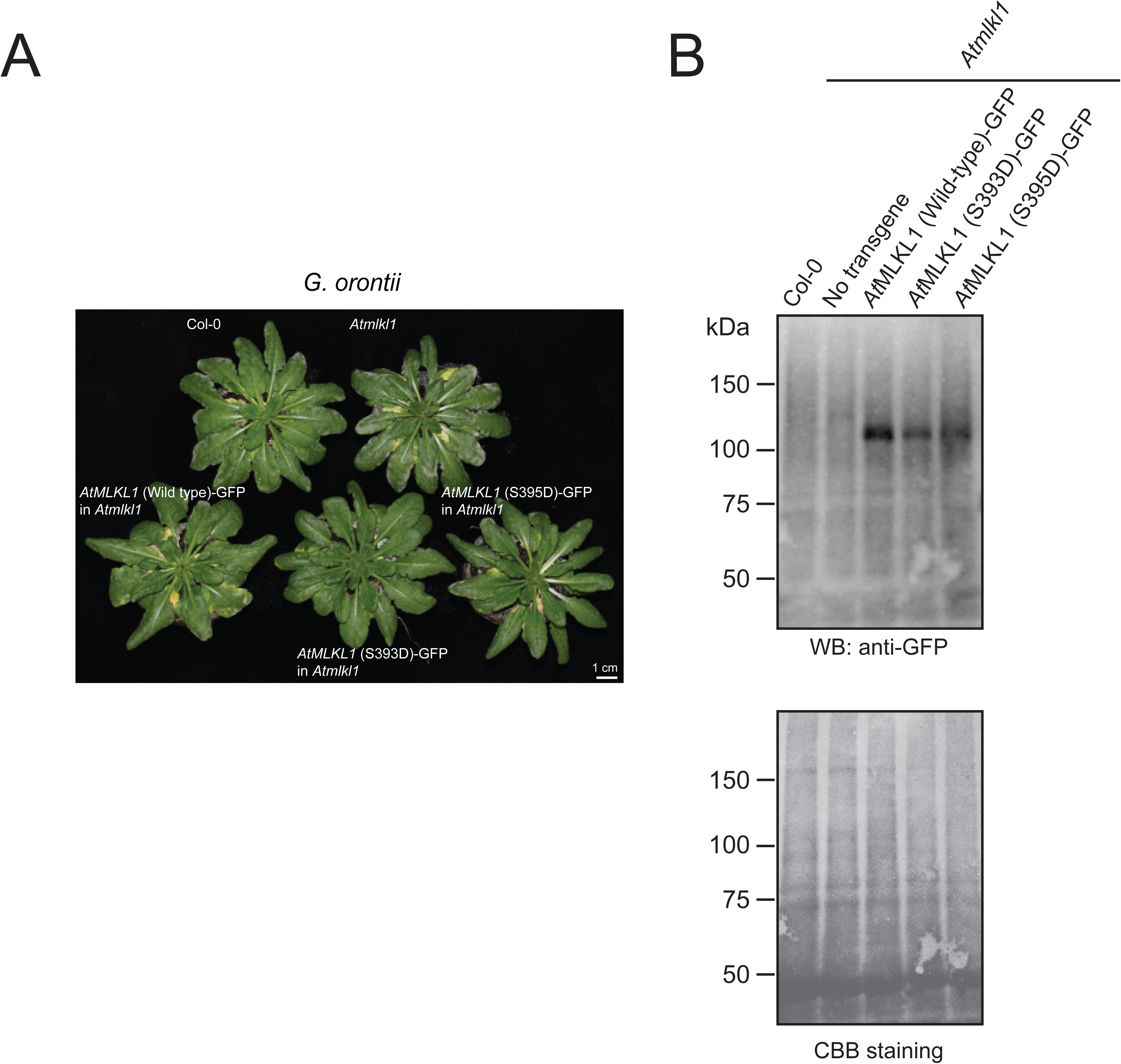
Genetic complementation of the *Atmlkl1* mutant. **A**, Macroscopic phenotype of transgenic *Arabidopsis* lines expressing wild-type or phosphomimetic variants of *At*MLKL1-GFP under the native *cis*-regulatory sequence in in *Atmlkl1* in response to *G. orontii*. Plants were photographed seven days after pathogen challenge. **B**, Western blot analysis of the C-terminally GFP-tagged *At*MLKL1 variants in the stable transgenic lines. Total protein extracts were collected from two-week-old plants of the transgenic lines grown on sterile Murashige and Skoog solid media.

**Supplemental Fig. 9:**
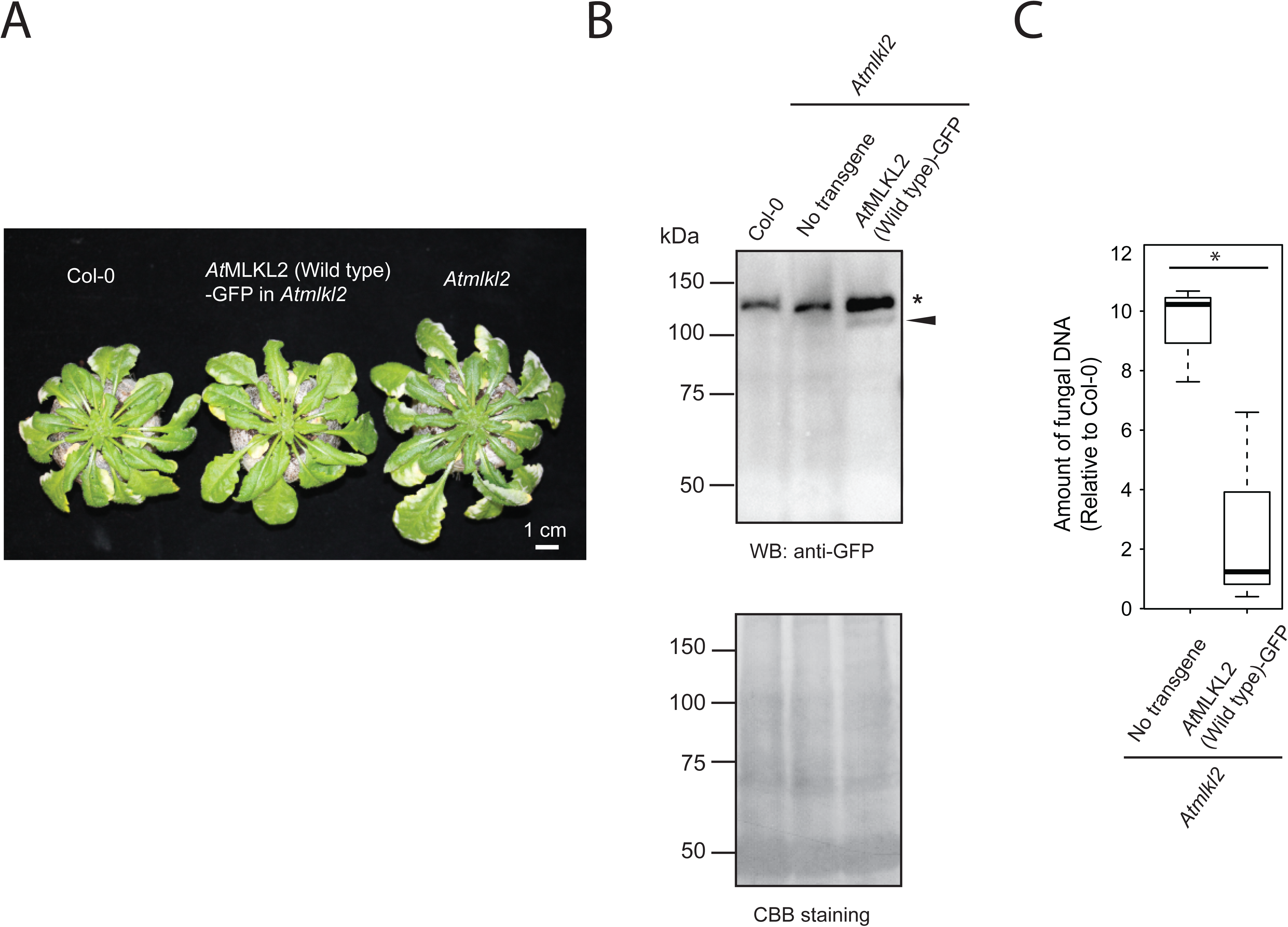
Genetic complementation of the *Atmlkl2* mutant. **A,** Macroscopic phenotype of transgenic *Arabidopsis* lines expressing *At*MLKL2-GFP under the native *cis*-regulatory sequence in *Atmlkl2*. Four-week-old plants were inoculated with *G. orontii* spores and plants were photographed at 14 days after pathogen challenge. **B**, Western blot analysis of C-terminally GFP-tagged *At*MLKL2. Total protein extracts were collected from two-week-old plants of the transgenic lines grown on sterile Murashige and Skoog solid media. The asterisk and arrow indicate non-specific and specific bands, respectively. **C,** Quantification of *G. orontii* DNA in infected leaves. Four-week-old plants were inoculated with spores of the powdery mildew and plants were examined at 14 days after the pathogen challenge. Fungal DNA was quantified by qPCR and normalised using the amount of plant specific gene (AT3G21215). The relative amounts to corresponding Col-0 wild-type samples were presented. Experiments were repeated three times (n=3) and the asterisk indicates a statistically significant difference (Tukey HSD, *p* < 0.05).

**Supplemental Fig. 10:**
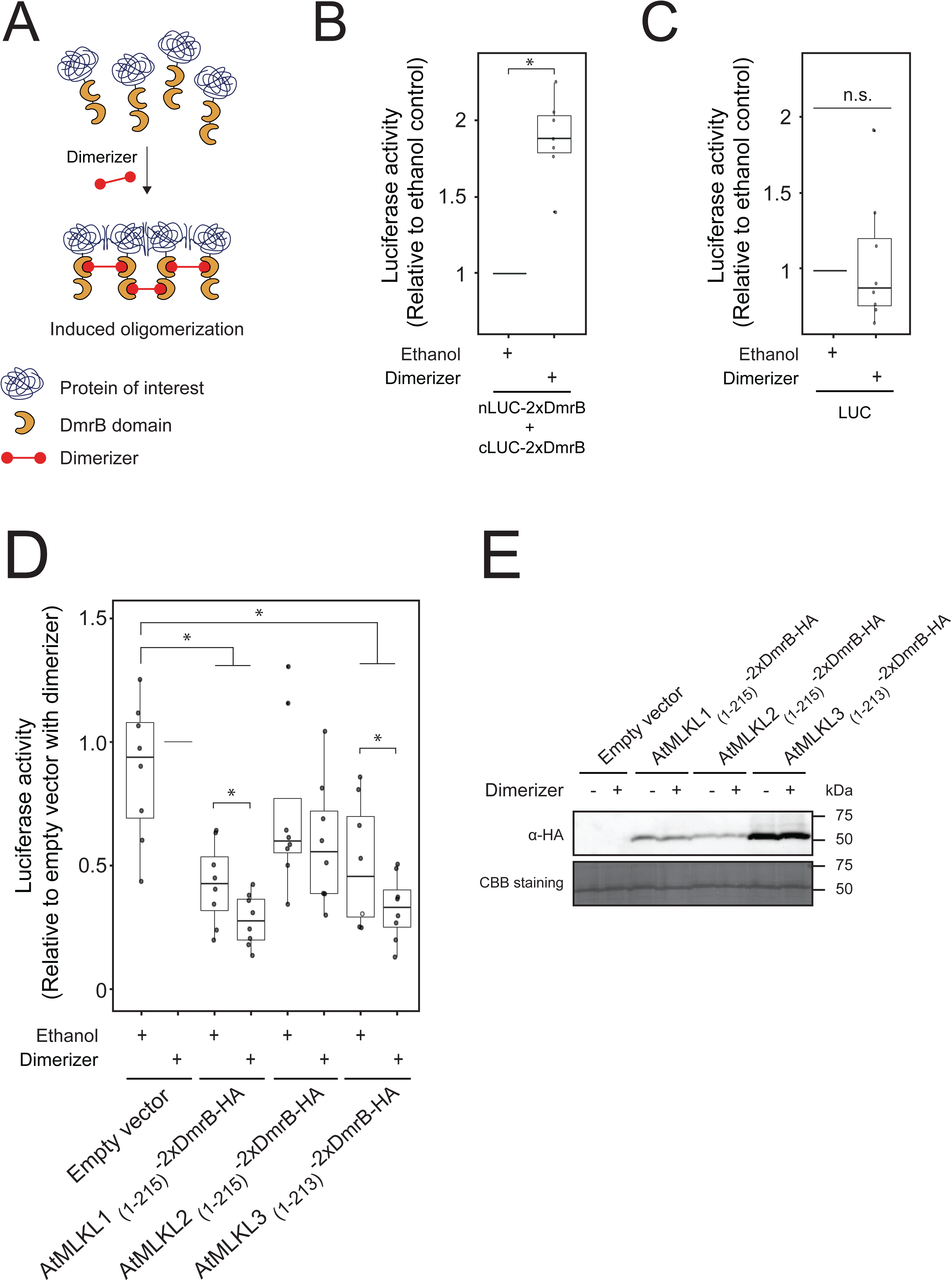
N-terminal domains of *At*MLKL1 and 3 were sufficient to elicit cell death and their enforced oligomerization potentiated the activities. **A,** Schematic of chemically induced oligomerization. AP20187 (Dimerizer), a synthetic cell-permeable ligand, induces homodimerization of fusion proteins containing the DmrB domain. Dimerizer–dependent oligomerization is facilitated by a tandem fusion of DmrB domains. **B**, Dimerizer–induced reconstitution of luciferase activity of the N- or C-terminal halves of luciferase fused with 2 x DmrB domains. Data were obtained with eight independent transfections (n=8). Luciferase activity was measured at 16 hours post transfection. The luciferase activity was statistically higher in the presence of dimerizer (one sample *t*-test, *p* < 0.01). **C,** Dimerizer does not influence the luciferase reporter assay. Data were obtained with seven independent transfections (n=7). Luciferase activity was measured at 16 hours post transfection. No statistically significant differences were detected (one sample *t*-test). **D**, HeLo domain plus brace region of *At*MLKL1 and *At*MLKL3 were sufficient to elicit cell death and their enforced oligomerization potentiated the activities. Expression constructs for luciferase and each of the *At*MLKL fusion proteins indicated in the figure were co-transfected into protoplasts and luciferase activity was measured as a proxy of cell viability at 16 hours post transfection. Relative luciferase activities compared to those of the empty vector control with dimerizer were plotted. Data were obtained with eight independent transfections (n=9). Asterisks indicate statistically significant differences (Tukey HSD, *p* < 0.01) **E,** Western blot analysis of the HeLo domain plus brace region constructs expressed in *Arabidopsis* mesophyll protoplasts. The fusion proteins were expressed as C-terminally HA-tagged proteins. Total protein extracts were collected at seven hours post-transfection. **B-E,** Mesophyll protoplasts were prepared from the triple mutant *Atmlkl123* and AP20187 (Dimerizer) was added after transfection (see method).

**Supplemental Fig. 11:**
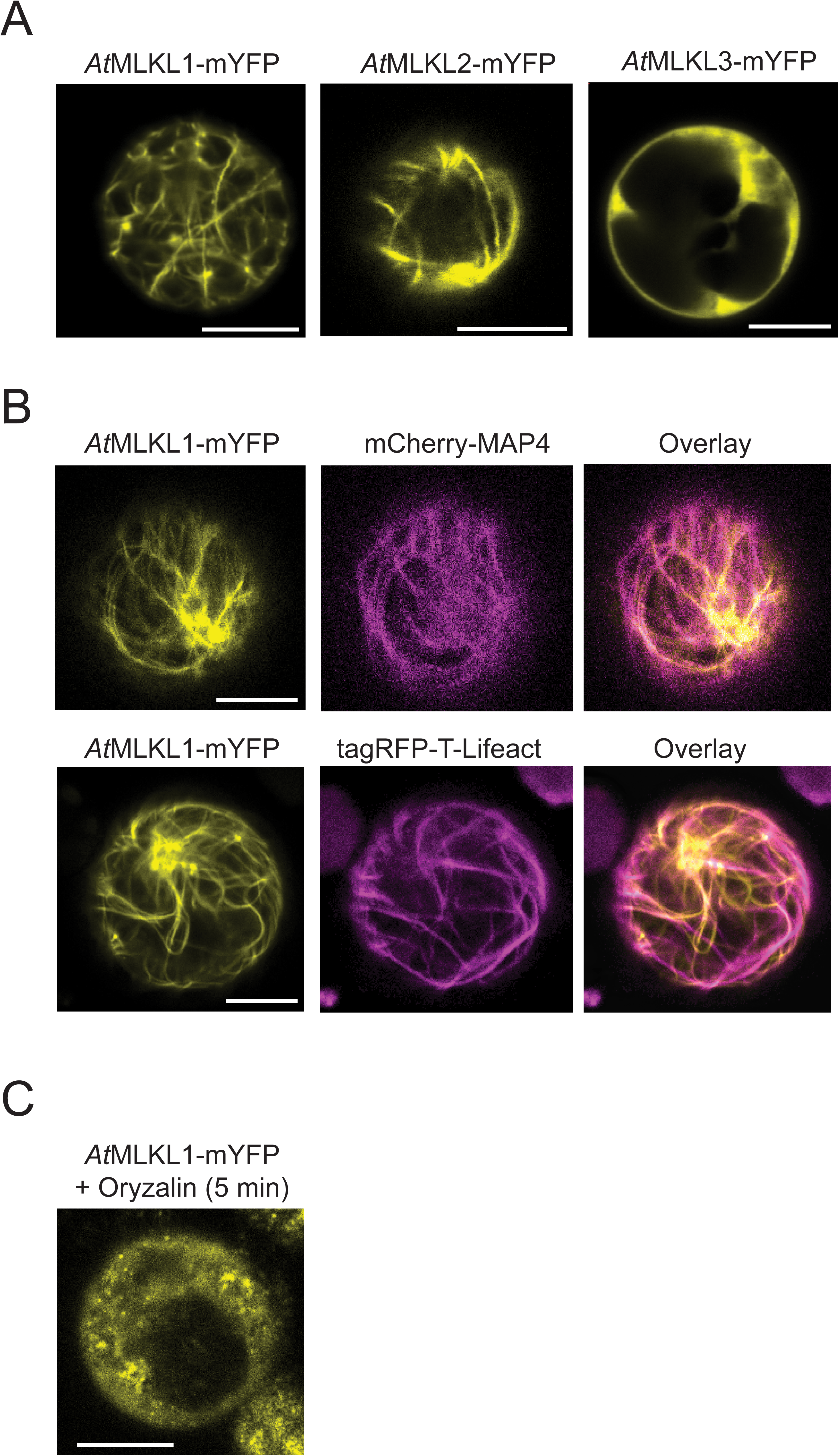
Subcellular localization of *At*MLKLs in protoplasts. **A,** Subcellular localization of *At*MLKLs in *Arabidopsis* mesophyll protoplasts. The C-terminally monomeric YFP (mYFP)-tagged variants were expressed under the 35S promoter. Representative confocal images were taken at ten hours post transfection. Scale bars = 10 μm. **B**, Co-expression of *At*MLKL1 and cytoskeleton markers in *Arabidopsis* mesophyll protoplasts. Co-expression of *At*MLKL1-mYFP and mcherry-MAP4 (microtubule marker: (*33*) top panels), or tagRFP-T-Lifeact (Actin marker: (*34*) middle panels). Representative confocal images were taken at 10 hours post transfection. Scale bars = 10 μm. **C**, The filamentous structures of *At*MLKL1-mYFP were undetectable at 5 minutes after application of 5 uM of the microtubule inhibitor, oryzalin. Representative confocal image was taken at 10 hours post transfection. Scale bar = 10 μm.

**Supplemental Fig. 12:**
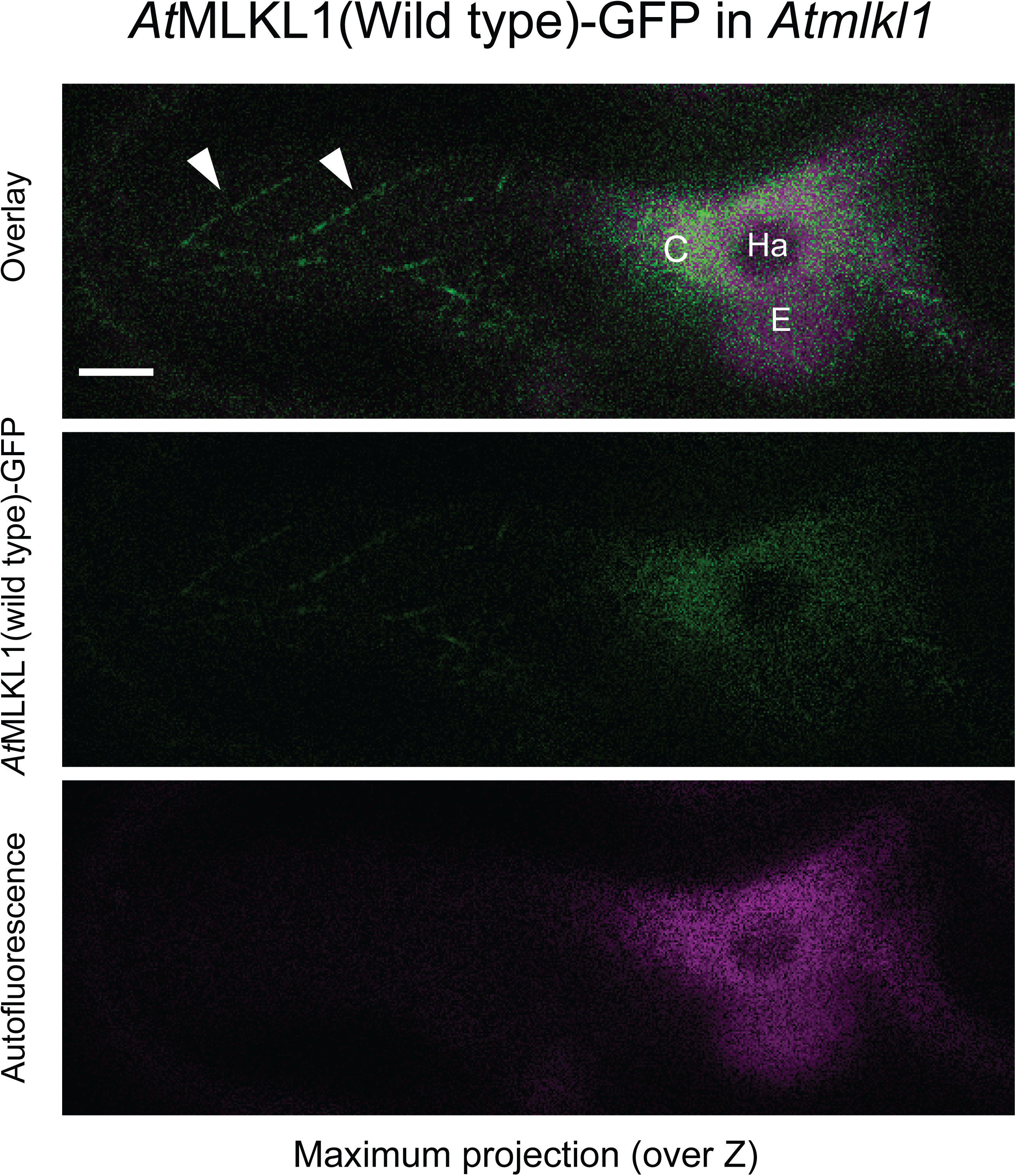
Confocal images of an epidermal cell that was infected by *G. orontii*. The maximum intensity projection was obtained from confocal Z-stack images and the autofluorescence of fungal structures were discriminated from *At*MLKL1-GFP signals by spectral imaging. Arrowheads indicate filamentous structures associated with *At*MLKL1-GFP. C; cytoplasm, Ha: haustorium, E, haustorial encasement. Scale bar = 5 µm.

**Supplemental Fig. 13:**
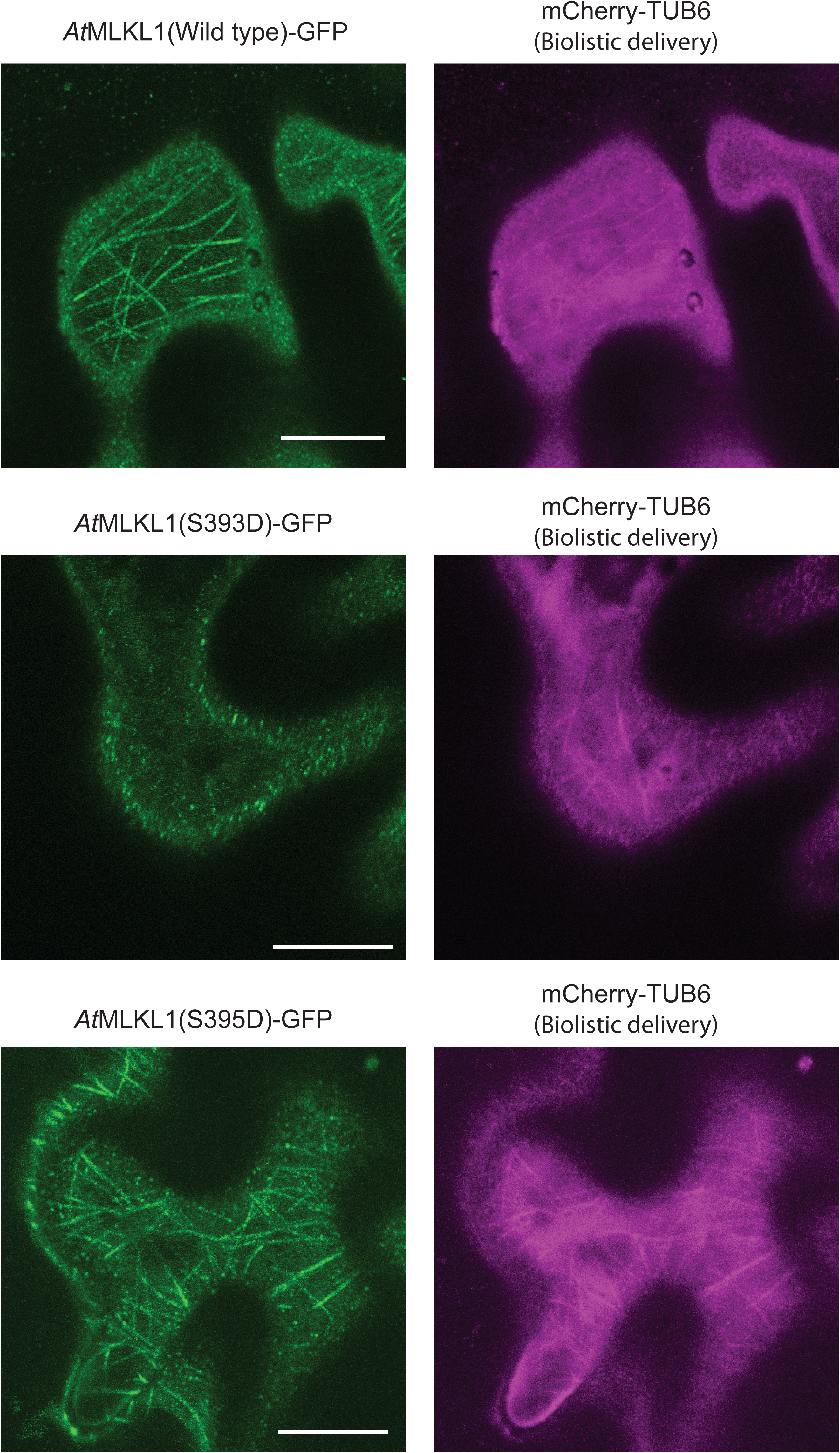
Cortical microtubule arrays in abaxial leaf epidermal cells of *transgenic A. thaliana lines expressing* phosphomimetic and wild type variants of *At*MLKL1. Representative images were obtained one day after biolistic delivery of the expression construct for the *At*ubiqutin10 promoter driven by mCherry-TUB6 (microtubule marker (*24*)) into the indicated transgenic lines. Scale bars = 10 μm.

**Supplemental movie 1**

Dynamics of *At*MLKL1(wild-type)-GFP in a non-pathogen-challenged abaxial leaf epidermal cell. The corresponding projection image is shown in Fig. 4E. The white circle indicates a GFP signal proximal to the PM (see Fig. 4D) and which was less mobile compared to other signals.

**Supplemental movie 2**

Dynamics of *At*MLKL1(S393D)-GFP in a non-pathogen-challenged abaxial leaf epidermal cell. The corresponding projection image is shown in Fig. 4E.

The white circle indicates a GFP signal proximal to the PM (see Fig. 4D) and which was less mobile compared to other signals.

**Supplemental movie 3**

Dynamics of *At*MLKL1(S395D)-GFP in a non-pathogen-challenged abaxial leaf epidermal cell. The corresponding projection image is shown in Fig. 4E. The white circle indicates a GFP signal proximal to the PM (see Fig. 4D) and which was less mobile compared to other signals.

**Supplemental file 1**

108 coding sequences (CDS) of plant *MLKLs* belong to the same orthogroups (*11*). A CDS derived from *Arabidopsis halleri* is incomplete due to the ambiguous assembly of the corresponding genomic region.

**Table S1.** Number of MLKL-like genes in plant genomes.

| Plant species | NCBI<br>Taxonomy<br>ID | Family | Number of *MLKL*-like<br>genes |
| --- | --- | --- | --- |
| *Coccomyxa subellipsoidea* | 248742 | Coccomyxaceae | 0 |
| *Ostreococcus lucimarinus* | 242159 | Bathycoccaceae | 0 |
| *Volvox carteri* | 3067 | Volvocaceae | 0 |
| *Physcomitrella patens* | 3218 | Funariaceae | 0 |
| *Selaginella moellendorffii* | 88036 | Selaginellaceae | 0 |
| *Actinidia chinensis* | 3625 | Actinidiaceae | 3 |
| *Amborella trichopoda* | 13333 | Amborellaceae | 1 |
| *Spirodela polyrhiza* | 29656 | Araceae | 2 |
| *Phoenix dactylifera* | 42345 | Arecaceae | 3 |
| *Arabidopsis halleri* | 81970 | Brassicaceae | 2 |
| *Arabidopsis lyrata* | 59689 | Brassicaceae | 3 |
| *Arabidopsis thaliana* | 3702 | Brassicaceae | 2 |
| *Brassica oleracea* | 3712 | Brassicaceae | 3 |
| *Brassica rapa* | 3711 | Brassicaceae | 2 |
| *Boechera stricta* | 72658 | Brassicaceae | 3 |
| *Capsella grandiflora* | 264402 | Brassicaceae | 3 |
| *Capsella rubella* | 81985 | Brassicaceae | 3 |
| *Eutrema salsugineum* | 72664 | Brassicaceae | 4 |
| *Schrenkiella parvula* | 98039 | Brassicaceae | 3 |
| *Beta vulgaris* | 161934 | Chenopodiaceae | 2 |
| *Cucumis sativus* | 3659 | Cucurbitaceae | 2 |
| *Ricinus communis* | 3988 | Euphorbiaceae | 2 |
| *Arachis ipaensis* | 130454 | Fabaceae | 2 |
| *Medicago truncatula* | 3880 | Fabaceae | 2 |
| *Phaseolus vulgaris* | 3885 | Fabaceae | 2 |
| *Linum usitatissimum* | 4006 | Linaceae | 4 |
| *Gossypium raimondii* | 29730 | Malvaceae | 3 |
| *Theobroma cacao* | 3641 | Malvaceae | 2 |
| *Musa acuminata* | 4641 | Musaceae | 4 |
| *Erythranthe guttata* | 4155 | Phrymaceae | 2 |
| *Pinus taeda* | 3352 | Pinaceae | 1 |
| *Brachypodium distachyon* | 15368 | Poaceae | 2 |
| *Oryza sativa* | 4530 | Poaceae | 2 |
| *Sorghum bicolor* | 4558 | Poaceae | 2 |
| *Setaria italica* | 4555 | Poaceae | 2 |
| *Zea mays* | 4577 | Poaceae | 2 |
| *Aquilegia coerulea* | 218851 | Ranunculaceae | 1 |
| *Fragaria vesca* | 57918 | Rosaceae | 2 |
| *Malus domestica* | 3750 | Rosaceae | 5 |
| *Prunus persica* | 3760 | Rosaceae | 2 |
| *Coffea canephora* | 49390 | Rubiaceae | 2 |
| *Citrus clementina* | 85681 | Rutaceae | 2 |
| *Citrus sinensis* | 2711 | Rutaceae | 2 |
| *Populus trichocarpa* | 3694 | Salicaceae | 2 |
| *Capsicum annuum* | 4072 | Solanaceae | 3 |
| *Solanum melongena* | 4111 | Solanaceae | 3 |
| *Solanum lycopersicum* | 4081 | Solanaceae | 3 |
| *Solanum tuberosum* | 4113 | Solanaceae | 3 |
| *Vitis vinifera* | 29760 | Vitaceae | 2 |

**Table S2.** Cryo-EM statistics and model refinement for AtMLKL tetramer.

|  | AtMLKL3 tetramer | AtMLKL2 tetramer |
| --- | --- | --- |
| **Data collection and processing** |  |  |
| Microscope | FEI Titan Krios | FEI Titan Krios |
| Detector | Gatan K2 Summit | Gatan K2 Summit |
| Voltage(kV) | 300 | 300 |
| Magnification | 22500x | 22500x |
| Pixel size (Å) | 1.30654 | 1.30654 |
| Total electron dose (e<sup>-</sup>/Å<sup>2</sup>) | 50 | 50 |
| Defocus range (μm) | -1.5 ~ -2.3 | -1.6~-2.2 |
| Micrographs collected | 1,434 | 1,828 |
| **Reconstruction** |  |  |
| Total extraced particles | 983,779 | 1,527,009 |
| Number of particles used for 3D reconstruction | 629,287 | 796,901 |
| Number of particles used for refinement | 123,472 | 154,008 |
| Symmetry | D2 | D2 |
| Resolution (Å) 0.143 after refinement | 3.84 | 4.51 |
| Resolution (Å) 0.143 after post-processing | 3.39 | 4.08 |
| Map sharpening B-factor (Å<sup>2</sup>) | -152 | -222.79 |
| **Refinement** |  |  |
| Estimated Resolution (Å) | 3.4 | 4.1 |
| Model composition |  |  |
| Number of protein atoms | 16584 | 14238 |
| Number of ligand atoms | 0 | 0 |
| MapCC (mask/box) | 0.83/0.79 | 0.81/0.81 |
| Rwork/Rfree (%) | 34.35/34.35 | 39.4/39.4 |
| R.M.S deviations |  |  |
| Bonds lengths (Å) | 0.011 | 0.010 |
| Bonds angles (°) | 0.941 | 0.797 |
| **Validation** |  |  |
| MolProbity overall score | 2.22 | 2.56 |
| All-atom clashscore | 15.96 | 18.07 |
| Rotamer outliers (%) | 0.67 | 1.57 |
| C-beta deviations | 0.0 | 0.0 |
| EMRinger score | 1.56 | 1.42 |
| Ramachandran plot statistics |  |  |
| Preferred (%) | 91.27 | 85.27 |
| Allowed (%) | 8.33 | 14.31 |
| Outlier (%) | 0.40 | 0.42 |

**Table S3.**
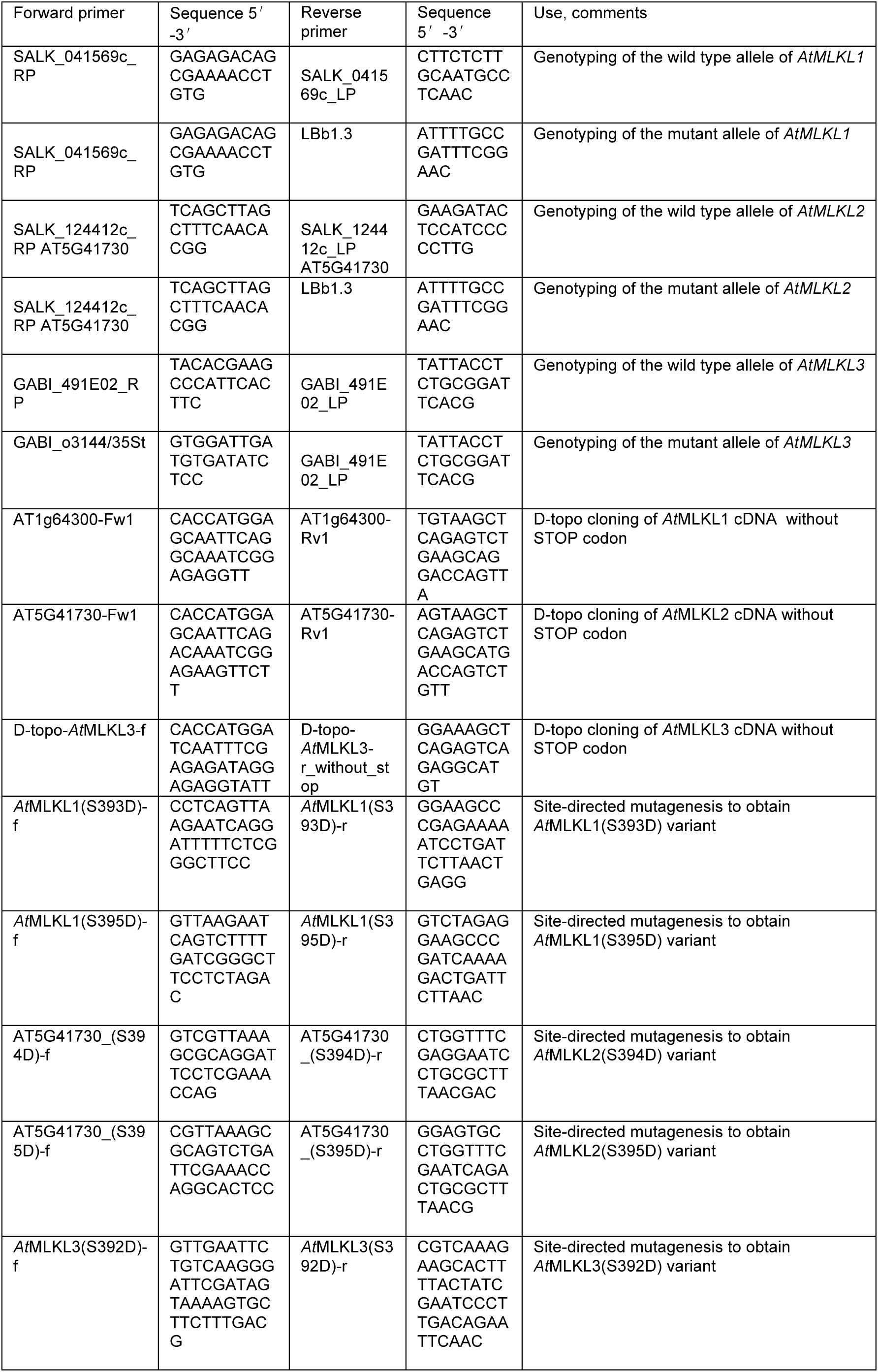
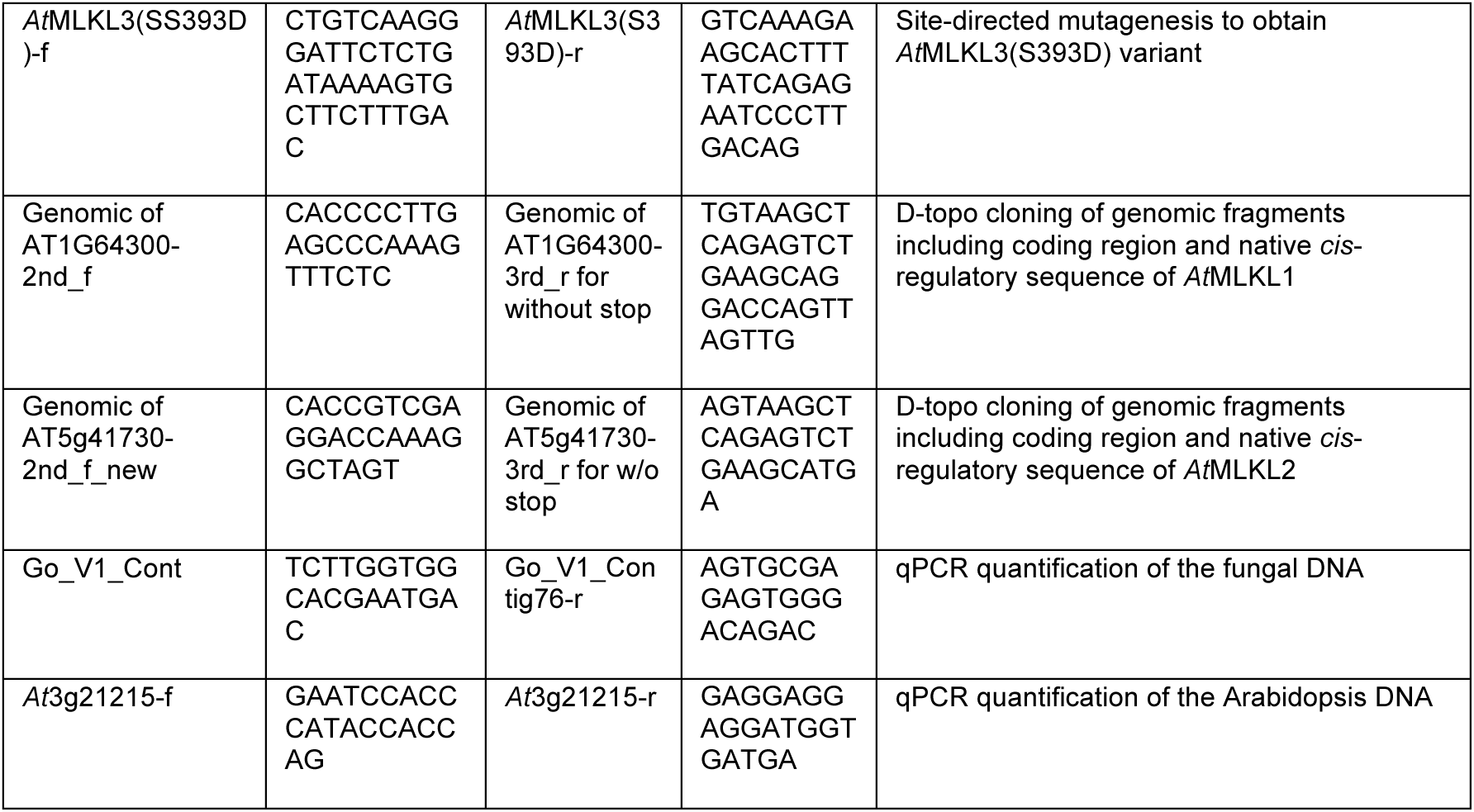
Primer pairs used in this study.

