## Supplemental file 1 for "Plant mixed lineage kinase domain-like proteins limit biotrophic pathogen growth"

> A_chinensis_1

ATGGTCTGCCACCTCCATTTTCATTTGGCCACCATTTTCTTGGCAAGACTTTTGAATTCCAATCTGGGAATGGAGCAATTCCGGCAGATCGGAGAGGTAGTGGGAAGCTTGAAGGCTCTAATGGTGTTCCGCGATGAAATACAGATCAATCAGAGGCAATGCATCTTGTTGCTCGACATGTTCAGCTTCGCCTACGAGTCAATCGCGGAAGAGATGAGACAGAACCTGAGATTCGAAGAGAAGCAAACGAAATGGAGAATTCTCGAGCAGCCATTGAAGGAGCTTCACAGGGTTTTTAAAGAAGGAGAGGTATACATAAGGCAGGGCTTGGAAACAAGAGATTGGTGGGTGAAAGCCATCACCCTCTATCAAGACACGGACTGCGTGGAGTTTCACATCCACAACTTGCTCTCTTGCATCCCAATTGTGGTAGAAGCAATTGAATCAGCTGGAGAAATATCCGGGTGGGACCACGACGAGATACAGAAGAAACGAATTATATGTTCGATGAAGTACCAAAGGGATTGCAAAGACCCGAAACTTTTCCAGTGGAGATTCGGGAAGCAGTACCTGGTTTCTCCGGATTTCTGCAACCGGTTTGCTGCTGTTTGGGAAGAGGATAGATGGATTCTTCTCAACAAAATCCGGGAGAATAGGAAGTTGGGTTTGACAAAGCACGAGAGGCAACTCGCGGACTTGCTTTGCAAGAACTTGAATGAATCAGAGCCGTTGAATGCTAGACTCTTGCCGTTTTCGATCTTGGTGGGTACGAAGGATTACCAGGTGAGGTGGCGGTTAGGGAGCGGAAGCCAATACAAGGAGATCCAATGGCTGGGTGAGAGCTTCTGTTTAAGGCACTTCTTTGGGGACATTGAATCAATAGTCCCTGAGATAGCCAAAGAAATCTCTCTCTCTCACCCAAATATAATGCAAATCCATTGTGGGTTCACCGACGAGGAAAAGAAAGAGTGCCTTCTGGTAATGGAACTCATGAATCGAGATCTCTCAAGCTATATCAAGGAGATCTGTGGCCCGAGGAAACGAATTCCAATTCCGTTCTCGCTCCCAGTTGCAGTTGATCTCATGCTTCAGATTGCAAGAGGAATGGAATATCTCCACTCGAAACAAATCTACCACAGAGATCTGAACCCTTCAAACATCCTCGTAAAGGCGAGAAATGCATCCACCGAAGGGTATTTGCATGTAAAATTTTCTGGGTTCGGCCTTTCATCCTCCATTAACCTCACCCAAAAGGCATCTCTGAATCAGAATGGGACACTCCCTTTCATCTGGCATGCTCCGGAGATTTTAGCAGAACAAGAACAATTGGGAAGTATAGGTAACTCCAGGTACACGGAAAAGGCTGATGTGTACAGCTTTGGGATGATCTGCTTCGAGCTTTTGACCGGGAAAGTCCCTTTCGAGGATGGTCATCTTCAAGGAGATAAGATGAGCCGAAACATTAGGACCGGAGAAAGGCCACTATTTTCATTCTCTTCACCTAAATATCTGACTACATTGACCAAAAGATGTTGGCATACCGACCCCAATCAACGGCCAAGTTTCCCATCGATTTGTAGGATTCTTCGCTACATTAAACGGTTCTTAGTGATGAACCCCGAGCTCAGCCAGACAAACCTACCCAATCCAACACTAGATTACTGTGAAACCGAGGCAGGGCTTTTAAAAAGCTTTCCTTCTTGGGGGAGTTCGAACTCATTGCCCGTCTCACAAATCCCATTTCAGATGTTTGCTTATAGAATTGTGGAGAAAGAAAAAACAAATACAAGTCAGAGAGATACCTCTGAATCGGGGAGTGATGGAGCCTCAGCGTGTGGGGATGAGAATGTGACTGCAGACGACCCATTCTCATCGGGAAGCGAGAAGAAGACATTTTTAACCTCTCCTGAAACTCTGCACATAAAATTGTCAAAATTGAAAAAATCTCCAGATGTGAAAAGCAGCAAACAGCCAGGGACACCGAAAGGACGGTCATTAAGGCCTCCACAAATGTCTCCTTGTGGACGCAGTTTAAGAATGAATTCTGAGAGCCAGCTAATGGCCATGAGTCCAAAACTACGAAGTACGTCTGGTCATGCCTCAGATTCAGAGCTCTCATAG

> A_chinensis_2

ATGAAGGCTCTTATGGTGTTGAAACATGATATTTTGATCAATCAAAGGCAGTGTTGTTTGCTTCTAAATATCTTCGATTTGGCCTTCGAAACTGTTTCGGACGAAATTAGACAGAACTTGAGACTAGAAGACAAGAACACAAAGTGGAAAGCTCTTGAACACCCTCTGAGAGAGCTTCACAGGGTCTTCAAAGAATGTGAACTCTACATAAGGCATTGTTTGGATGTCAAGGACTGGTGGGGCAAAGCAATGAGCCTCCACCAGAACAATGACTGTGTTGAGTTTCACATTCACAACTTGCTTTGCTGCTTTGCCATTATTTTCCAGTGGAAGTATGGGAAGCAGTACTTGGTTTCTCGAGAAATTTGCAATGAGTTGGGTAGTGCTTGGAGAGAAGACCGATGGATTCTCCTCGAGAAAATCAGGGAGAAGAGAAACACGGTATCTGTCACCGAGGCCAAGCATGAGCAGAGAATAGCAGACCTACTGATTAGGAAACTAAGTGGGTTAGAGCCGGTTAAGGGAAAGCTTTTGCCGGCTTGGATCTTAATAGGAGCGAAGGACTATCATGTGAAGCGACGGTTAGGATCTGGGGGAAGTCATTGTAAGGAGATTCAATGGCTTGGGGAAAGTTTCGCTTTGAAAAATTTCTTCGGGGAGATGAATGAACAACTACAAGATGAGATATCTTCGGTCCTCTCTCTTTCGCACCCCAACGTAACGCAATACCTTTGTGGGTTTTATGACGAGGAAAGAAAAGAGGGTTTTCTTGTTATGGAGCTTATGAACAAGTCTCTTGCCGGTCATATGAAAGAGAATTGTGGCCAGAGGAAACGGCTTCCATTCTCTATTCCAGTTGCAGTCGATATCATGCTTCAAATCGCAAGAGGTTTGGAATACCTACACTCTCGAAAGATCTATCACGGGGACCTGAACCCATCTAACATCCTCCTCAAAGCGAGAAATCCCATCCCGGAAGGTTATTTTTTTGCAAAAGTTACAGGTTTTGGCTTAACCTCCATTAAGAGTTACACTTCTCGAGCTCAGACAAGCCAAAATGGAGACAATCCCATTAGATGGTATGCCCCCGAAGTTCAGGCCGAGCAAGAACAGCAGGGCGGTGGTAAATGCAGCTCAAAATATGGGGAGAAAGCAGATGTCTATAGCTTTGGGATGCTTTGCTTCGAGATTCTAACCGGAAAAGTTCCGTTTGAAGAAGGGCATCTTCAAGAAGACAAGATGGGTCGTAACATAAGAGCTGGGGAGAGGCCTCTCTTCCCATATAATTCACCAAAGTACCTCGTGAACTTGACCAGAAAATGCTGGCAAGCCGAGCCAACTCAGCGCCCGAGCTTCTCGTCCATATGCAGGATTTTACGGTACATTAAAAAGTGCCTCGTGATAAACCCGGATCACGGTCAGCCAGAATCGCCTCCCCCACTCGTGGATTACTGTGAGATTGAAGCAGGGTACTCGAAATTGTTCCCTGGAGACGGAAGTCCTTATTTGGCACCGGTCTCACAAGTTCCGTTTCAAATGTTTGCTTATAAGCTTATTGAGAGAGAGAAGACTTCCGGGAGCTTTAAAGATAAGATTTGGGATTTGGCAATCGAAACGCGTTCAATTTGGGGGGGTGCTTCAGTTTTTGGGGATGATAACGGGGTTGTAGCCGACGATTCTTTTCTGGTTCCAATTGATCAGAGGTCAGTTTGTTCCGAGGTTCACGAGAGAAAAACCTTATCGAAAATAGGAGCCGACCAAAGGTCTGTTTGTTCCGAGACTCCAGGGAGGAGAGTTCAATCGTTTAACACAGCAGCCGATCAAATGTCGTTTCGTTCAGAGTTTTTATGGAGGAAAATGGACCAGAGATCGGTTGGTTCTGTAACTCCTGAGCGAAAATTTTTTCCAACAGCATCAACCGATCAAAAATCTATCGGTTACGTGACTCCAGAGAGGAAGATTTTCGCCCATGATCAAATATCCGTCGGTTCTGAGACTCCAGAGAGAAAATTTGTGCCCACAACAGAAACTGAGCAGATTTCAGGTCATTCGGAGAATGGAGAGAGGAAGGATTTACCGGTATCAGCAATGGATGGAAAAAATCAAAGTCCTGAATCCCCAGAGAAGAAACCAGATAAGAAACTGGCAACGACAATAGCAGCAAATCAAAATTCAGTTGGTTCTAAGCTTACAAAGAGAAGTACTTTTCCAGCAACAGAGGTCGATAAAGTTTCTCCCGAGAGTCATGAGACGAAGACTTTGTCGAGAGCAGTCTCAACTTGTTCCAAGTCTCTGGAGAGAAAAGTTTTGCCGCCAATAGCATCCGATTCAGATTTTTCCTCGATTTGTTCTGAGAGTCCGAAGAAGAAAAAAACATCGAAAAAATCAAGTGATGTATGGGTGGTTCGTCCTGAGGTTCGAGACAGAAAATGTTTGGTGAAACCGAATGTAGCGGTGGCAGAGACAAGCAAGGGTCCAGGTTAG

> A_chinensis_3

ATGGAACAATTCCGGCAGATCGGAGAGGTAGTGGGAAGCTTGAAGGCTCTAATGGTCTCCGCATACGAATCAATCACAGAAGAGATAAGACAGAACCTGAGATTCGAAGAGAAGCAAACGAAATGGAGAATTCTCGAGCAGCCATTGAAGGAGCTCCACAGGGTTTTTAAAGAAGGAGAGGGATACATAAGACAGAGCTTGGAAACCCGAGATTGGTGGGCGAAAGCCATCACCCTCTATCAAGACACGGACTGCGTAGAGTTTCACATCCACAACTTGCTCTCTTGCATCCCAATTGTGTGGAGATTCGGGAAGCAGTACCTGGTTTCTCCGGATTTCTGCAACCAGTTAGCTACTGTTTGGGAAGAAGATAGATGGATTCTTCTCAACAAAATAAGGGAGAAGAGGAAGTCGGGTTTGACAAAGCACGAGAGGCAACTTGCGGACTTGCTGTGCAAGAATTTGAACAAATCAGAGCCGTTGAATGCTAGACTCTTGCCGTTTTCCATCTTGGTGGGTACGAAGGATTACCAGGTGAGGAGGCGGTTAGGGAGCGGAAGCCAGTACAAGGAGATCCAATGGCTGGGTGAGAGCTTCTGTTTAAGGAACTTCTTTGGGGACATTGAATCAATAGTCCCTGAGATAGCCAAAGAAGTCTCTCTCTCTCACCCAAATATAATGCAAATCCATTGTGGGTTTGCTGACGAGGAAAAGAAAGAGTGCCTTCTGGTAATGGAACTCATGAATCGAGATCTCTTGAGCTATATCAAGGAGATCTGTGGCCCGAGGAAACGAATTCCGTTCTCGCTCCCAGTTGCAGTTGATCTCATGCTTCAGATTGCAAGAGGCATGGCATATCTCCACTCAAAACAAATCTACCATGGAGATCTGAACCCTTCAAACATCCTTGTAAAGGCGAGAAATGTATCTACCGAAGGGTACTTGCATGTAAAAGTTTCTGGGTTCGGCCTATCATCCTCCATTAACCTCACCCAAAAGGCATCTCCAAATCAGAATGGGACGCCCCCTTTCATCTGGCATGCTCCGGAGATTTTAGCAGAACAGGAACAATCGGGAAGTATAGGAAACTCCCGGTACACGGAAAAGGCTGATGTGTACAGCTTTGGGATGATCTACTTTGAACTTTTGACCGGGAAAGTCCCTTTCGAGGATGGTCATCTTCAAGGAGATAAGATGAGCCGAAACATTAGAACAGAAGAAAGGCCATTATTTTCGTTCTCTTCACCTAAATATCTGACTACTTTAAACAAAAGATGTTGGCATACCGACCCCAATCAACGGCCAAGTTTCCCATCAATTTGTAGGATTCTTCGCTACATTAAACGGTTCTTAGTGATGAACCCTGAGCTCAGCCAGACCGACCTACCCAGCCCAACACTAGATTACTGTGAAGCTGAGGCAGGGCTTTTAAGGAGCTTTCCTTCGTGGGGGAGTTCGGACTCCTTGCTTGTATCACAAATCCCATTTCAGATGTTTGCTTATAGGATTGTAGAGAAAGAAAAAACAAATACAAGTCAGAGAGATACCTCTGAATCGGGGAGTGATGGAGCCTCGGCTTGTGGGGATGATCATATGACTGCCGACGACTCATTCTCATCGGGAAGCGAAAAGAAGAACTTTTTAGCCTCTCCTGAAACTCTGCACAGGAAATTGCCATTATTGAAGAAATCTCCAGATGTGAAAACCAGCAAAAAACAAGGGACACCGAAAGGACGGTCATTAAGGCCTCCACAAATGTCTCCTTGTGGACGCAGTTTAAGGATGAATTCTGAAGGCCAGCTAATGGCCATGAGTCCAAAACTACGAAGAACGTCTGGTCATGCCTCAGATTCAGAGCTCTCATAG

>A_halleri_1

ATGGAGCAATTCAGGCAAATCGGAGAGGTTCTTGGAAGCTTAAACGCGCTAATGGTATTACAAGACGATATCTTAATCAACCAAAGACAATGTTGTTTGTTGTTAGATATTTTCAGTTTGGCTTTCAACACCGTAGCCGAAGAGATCAGACACAACTTGAAGCTTGAAGAGAAGCATACTAAATGGAGAGCTCTTGAACAGCCTTTGAGAGAGCTTTATCGAGTTTTTAAAGAAGGCGAAATGTATGTTCGGAATTGTATGTCCAATAAAGATTGGTGGGGTAAAGTTATCAACTTTCATCAGAACAAAGATTGTGTTGAGTTTCACATACACAATTTGCTTTGTTACTTCTCGGCTGTGATCGAAGCCATCGAGACAGCGGGTGAGATTTCGGGTCTCGACCCTGCTGAGATGGAGAGGAGGAGAGTTGTGTTTTCAAGAAAGTATGATAGGGAGTGGAATGATCCCAAGCTGTTCCAGTGGAGGTTTGGGAAACAGTATTTGGTACCGAGGGATATTTGCATTCGGTTCGAGCATTCTTGGAGAGAAGATAGGTGGAATCTAGTGGAGGCATTGCAAGAGAAGAGAAAGTCCAAGAGCGATGAGATTGGGAAGACTGAGAAGCGTTTAGCTGATTTTCTGTTGAAGAAACTAACCGGATTGGAACAGTTTAACGGTAAGCTCTTCCCAAATTCGATACTTGTTGGTTCTAAGGATTACCAAGTGAGGCGGAGATTAGGTGGTGGTGGTCAGTACAAGGAGATTCAATGGTTAGGGGATAGTTTTGTGTTGAGACATTTTTTCGGTGATCTTGAGCCGTTGAATGCAGAGATATCTTCTCTGTTATCGCTTTGTCATTCGAATATCCTTCAGTACCTTTGTGGATTCTATGATGAAGAAAGGAAAGAATGTTCTCTGGTTATGGAGTTAATGCACAAAGATTTGAAAAGCTACATGAAGGAGAATTGTGGACCTAGAAGAAGATATCTTTTCTCTGTTCCCGTTGTGATCGATATAATGCTTCAAATCGCGAGAGGGATGGAATATCTTCATTCTAATGAGATTTTTCATGGAGATTTGAATCCCATGAACATTCTTTTGAAAGAGAGAAGTCACACCGAAGGTTATTTCCACGCAAAAATTTCAGGCTTCGGTTTAACCTCAGTTAAAAATCAGTCTTTTTCTCGGGCTTCCTCTAGACCAACCACACCCGATCCTGTAATATGGTATGCACCTGAAGTTCTAGCAGAGATGGAACAAGATCTCAAAGGTACAGCTCCGAGATCAAAGTTTACACATAAAGCTGATGTCTATAGCTTTGCTATGGTGTGTTTCGAGCTTATAACCGGTAAAGTTCCATTTGAAGATGACCATCTTCAAGGCAAGAAAATGGCTAAGAACATAAGAACGGGAGAGAGACCTCTCTTCCCATTCCCTTCGCCAAAATACCTTGTTAGTCTAATTAAACGGTGTTGGCATTCAGAACCGAGTCAGCGTCCAACATTCTCTTCGATATGCCGGATCCTCCGCTACATCAAGAAGTTCCTAGTGGTGAATCCGGATCACGGTCACCTTCAAATCCAAAACCCACTTGTTGATTGCTGGGACTTAGAAGCTAGGTTCTTGAAAAAGTTCTCAATGGAGTCAGGTTCTCACGCAGGATCGGTGACACAAATACCATTCCAGCTTTACTCATACAGGATTGCAGAGAAAGAGAAGATGATGAGTCCAAACTTTAACAAAGAAGAGAATTCAGAAACAGGGGAATCTGTTTCTGAAAGCGTTTCAGTGGTTGAAGATCCACCAACAATGCCAATGTATACAAAGTCATTGTGCTTAGATGCAATATCTGAATACTCAGATACAAGATCAGTTTACTCAGAAGCTCCAATCAAAAAGACATCAGCTTTAAAGAAAAGCGGCGACACCATAAAAAACAGAAGAAACTCAATCTCAGGTTTACGGTCTCCAGGATCATCGCCTATAAAACCGAGATCAACACCGAAAGTGTCATCACCACTAAGTCCATTTGGAAGAAGCAGCAGCAAAGCAAGGAAGGATACAAGATTACCTTTGAGTCCTATGAGTCCTTTGAGCCATGGAAGACGTAGACAACTCACTGGTCCTGCTTCTGACTCTGAGCTTACATAG

> A_halleri_2	no_stop_codon

ATGGAGCAATTTCGAGAGATAGGAGAGGTATTAGGAAGCATAAGAGCGTTAATGGTGTTCAAGGACAACATTCAAATCAACCAGCGTCAATGCTCCCTATTACTCGACCTCTTCACCGCGTCCTACGAGTCGATTTCCGAATCGATGAGATCGAATCTACGCTTCAAAGAGAAGAACACTAAATGGAAGATTCTAGAACAGCCTTTGAGGGAGCTTCTATGGGTGGTAAGAGAAGGAGAGGCTTATGTTAGAATGTCTTTAGAGCCTAAACTTGGTTTTTGGGCTAAAGCTGTTGTGTTACATTGCAATAGAGATTGCACAGAGCTTCACATACATAACTTGCTCTCTTGTCTACCGATCATTGTCGAGGCTATTGAGACAGCGGGTGAGGTTTCAGGGTGGGATGAAGAAGAGATGAGCAAGAAGAGGTTGGTGCATTCTAACAAGTATATGAAGCAATGGAATGACTCTCAGATGTTTACCTGGAAGTTCGGGAGAGAGTATTTGGTGACCGAAGATTTCTGCAACCGGTTCGAGAGTGCTTTGACAGAGGACCGTTGGATTCTGATCAAGGAGCTTCAGGAGAAAAAGCAGCCTGGTTCGAGCAAACATGATTGGAAGATGGCTGATTTCCTTTTGAAGCATTTGGGAGATGGAAATGAGAGTCCCAAGCTATTTCCGTCTTCTCTTCTGGTTAACACGAAAGATTACCAAGTGAAAAAAAGATTAGGGAATGGAAGTCAGTACAAGGAGATTACTTGGTTAGGCGAGAGCTTCGCACTTAGGCATTTCTTTGGGGATATCGATGCTTTGCTTCCCAAAGTTACTCCATTGCTATCTCTTTCACACCCAAACATTGTTTATTACCTCTGCGGATTCACGGATGAGGAGAAGAAAGAATTATTCTTGGTCATGGAACTGATGAGTAAAACCCTTGGTATGCACATCAAAGAGGTATGTGGTCCGAGGAAGAAGAACACTCTCTCCCTCCCGGTCGCGGTTGATCTGATGCTTCAGATAGCTTTGGGTATGGAATATCTTCACTCGAAGAGAATATACCACGGAGAGTTGAATCCATCTAACATTCTTGTCAAACCAAGAAGTCACCAATCCGGAGATGGGTATTTGCTTGGGAAGATTTTTGGGTTCGGGTTGAATTCTGTCAAGGGCTTCTCCAGTAAAAGTGCTTCTTTGGCGAGTCAGAATGAGAATTTTCCATTCATTTGGTATTCTCCGGAAGTTCTAGAGGAGCAAGAACAGAGTGGAACTGCAAGAAGCCTTAAGTATACTGATAAATCTGACGTGTATAGCTTCGGGATGGTTTGCTTTGAGCTTTTGACTGGTAAAGTACCATTTGAAGACAGCCACCTGCAAGGAGATAAGATGAGTAGAAACATTAGGGCCGGGGAGAGACCGCTTTTTCCATTTAATTCACCTAAATTCATAACAAACCTGACAAAACGATGCTGGCATGCTGATCCCAATCAGCGTCCAACTTTCTCATCGATAAGCAGAATTCTCCGGTATATCAAACGGTTCCTAGCTTTGAACCCGGAATGTCACAGTAGTAGCCAACAAGACCAGCCCATTGGTCCTCCCGTGGATTACTGCGAAATCGAGACAAAGCTATTGCAGAAGCTTTCATGGGAATCTACAGAGTCAACTCAAGTTTCACAGGTTCCTTTTCAA

>A_ipaensis_1

ATGGAACAGTTCCGGCACATTGGAGAGACTCTTGGGAGTTTGAAAGCGCTTATGGTATTGCGAGATGAAATTCAAATCAACCAACGGCAGTGCTGTTTGATCCTTGAAATGTTTAGCTTGGCGTTCGAGACGATTGCAGAGGAGATCAGGCAGAATCTGAAGCTGGAAGAGAGAAGCAGCAAATGGAAGCCTCTGGAGTTGCCTCTGAGAGAGCTTTGCAGGGTTTTCAAGGAAGCCGAGTTGTACATCCGGCAATGTTTGGATTCCAAGGAGTGGTGGGGGAAAGCCATCACCATGTCTAACAATTCTGACTGTGTTGAGTTCCATATCCACAATCTCCTTTGCTGTTTCCCGGCCGTGATCGAGGCCATTGAGAATGCAGGGGAGGCCTGTTGTCTTGATCAGACCGAGAAGGAGAAGAAGAAGGTCATGGTGTCAAGGAAATATGACATGGAATGGAATGATTCCAAGCTTTTCCAATGGAGATTTGGGAAGCAGTACTTGGTCTCTCGGGAGATCATCAGCCAGTTGGAGAATGCATGGAAGGAAGATAGGTGGAGGCTTATAGAATCACTCAAGGAGAAGAAAACTACGTCGAAGAAGAACGAGCAGCGCCTCGCCGAGATACTCTTGAAGAAGCTTGTAAATGGATCTGACAGCAAAACGAATCAAGATCTTTGGCCGATCCACGTTCTCGTGGGAGCCAAGGATTACCAAGTGAGGCGAAGATTGGGAAAGGGGAAGGAGTATAAGGAAGTGACTTGGCTTGGACAGAGTTTTGCATTGAGGTACTTCAATGGAGAGAAGCAAGCCTATGAAGGTGAGATTTCAAAATTGTTATGTCTTTCTCACCCCAACATATTGCAATACCTTTGTGGATTCTACGATGAGGAGAAGAAAGAGTTTAATCTTGTGATGGAGTTGATGAGCAAGGATTTGTGGACCTACATGAAGGAGAATAGCGGTCCAAGGAGGCAGATCTTGTTCTCAATTCCAGTTGTGGTGGATCTCATGCTTCAAATTGCAAGAGGCATGGAGTATCTCCACTCCAAGAAAATCTACCATGGAGATCTTAACCCTTACGGTGTTCTACTCAAAGCAAGGAACTCCCAAGAAGGTTACTTCCATGCAAAAGTCTCAGGATTTGGCCTTCCTTCCCTGAGGAACCGCGATCCAGCACAGAACCCCGAGGACCTGAACCCTAGCATATGGTATGCTCCGGAAGTACTGCACGAGATAGAAAAAGGAGGGAAGGTTTCTACCTCGTGCAAGTACTCGGAGAAAGCAGATGCATACAGCTTTGGAATGATTTGCTTTGAGTTGCTGACAGGGAAAGTCCCTTTCGAAGATAACCACCTGCAAGGGGAGAGAACCACACAGAACATCAAAGCAGGGGAGAGGCCTCTCTTCCCTGGTCGCTCCCCAAAATACCTTGTTAGCTTAATCAAGAAGTGTTGGCAAACCGATCCTTCACAGCGTCCTACTTTCTCTTCTATTTGTAGGATCTTGCGTTACACCAAGAAGTTCCTCTCCACCAGTACTGAGTACTATATCATTAATCCCGAACTTAACAACCTTGAGCTTCAATCTCCACCTGTTGATTGCTGCGATATAGAGACCACTTTCCTTAAGAATCTACCGGAGGAGATGGCGCCACATGTTTCCGCTGTTTCTCAAATCCCTTACCAGATCTATGCTTACAAGGTTGTGGAGAAAGGCAAGATCAGCCTCTTGAATTCAAGAGATAAAGGCAGCGAGACAGAGTTGCCAAAGGAAGATTCTCCTCAAGGAAGCGATGAATTCAGCTTCGCTTTTGAAGATGATCAAGTTTCCGCGGCTGAACCCTTTCAGTTTATAACTTCTCCCAAGTCAGTTCATGGCGGCGAAACACCGTCTTTTTTCTCCGAAGCACCGGCGAAGAAGACTGTTAGAATAAAGAGCCCAGTGCGAAAACTTGACAAGCTCAAGAAAGACCAAGGAGCGAAATCAATGCCGGCATCTCGCATTACTTTAAAGGCAAGTAAGGCATATCTGGCGTCACCGGCAGCTAGTCCATTGCACCCTGCACTCAGAAAAAAACCAACACAAGTCTCAGAGACAGAGCGCAACACTTCTAAGAACAACAAAGCTGTGACCACTACTCCTTCGACCCCAATAAGGAGAAGAAAATGCGAAGGGCAACACGTGCCAGAATTCATGGGAAGCCCGAGGGTGAAACCAAGGGATCAGTCACCATTAATGGCTAACAAGCTGAAGAAGACACCGCTGACAATGACCCCAGGAAGGATGATCACACGTACAAGCGACGCGCATTTGATACTGCAGAAGAGGCTCTTGATGGGTTCAAGCACAAGCAAAGGAGGAAGATGGGACCATGGAACTGAACACAGGACATCGCTGTCATCATTGGCATTGAGTCCTTTGAGTCCATATGTGATGAGAAGAAGAAGATCAACGGCGACAGCAGCAACATCATGTGGCCACGCCTCTGACTCAGAGAGCACTTGTTCTAAGCACAAAAGAGAGAGCATATCACCATTCGCAATGAGTCCTTTGAGCCCCTACGCTTCCAGAACAACAGCCTGTGGCCATGTCTCCGATTAA

>A_ipaensis_2

ATGGAACAATTTAGACAGATTGGTAAGGCATTGGGGAGTTTGAAGGCTTTGATGTTATTCAAAGAGAATATTCAAATCAATCAAAGACAGTGTTGCCTTCTTCTTGAAGTGTTCACCTTTGCATATGATACTATTGCAGATGAGATCAAACAGAACCTCAAATTTGAAGAAAGGAATGTGAAATGGAAAGGATTAGAACAACCTTTGAAAGAGATTCACAAGATCTTCAAAGAAGGTGAATCATATGTTAGGCATTGTTTGGAAACAAAGGATTGGTGGGCTAAGGCTATTATATTGTGTCACAACACAGATTCTGTTGAGTTGCATATACATAACTTGATTTGTTGCATGCCAGTTGTGATTGAAGCTATCGAATTGGCCGGAGAAACTTCAGGGATCGATCAGGACGACATGAATAAGAGGAAGTTAATAAATTCTCATAAGTTCAGAACAGAGTTTAGAGATATGAACCTTTTTCAATGGAAATTTGGGAAACAGTATTTAATTACACCTGAGTTTTGCAGCCGATACGAAACGGTTTGGAAGGAGGACAGGTGGTTTCTTCAACAGAAAATGCATGAGAAGAAGATGGCAGGAGCAACAAAGCAAGAGAGAAGATTGATAGAACTTATTTTGAAGAATTTGCAAGAATCATGGGAAGGGAAGCTTATTCCAAGTTCAACCTTGGTTGGATCTAAGGACTTCCAGGTTAAGAGAAGGATGGGGAATGGACAAGACAAGGAAATTGCGTGGTTAGGCGAAAGCTTCGTGATCAAGCATTATACCGGAGAGATCGAAGCTTGGGAGCCAGAGATCAGAGAAGTTTTGTCTCTGTCTCATCCAAACATAATGGATTGTCTTTGTGGTTTCACGGATGATGACAAGAAAGAATGTTTCTTGCTTACGGAAGTTATGAGCAAAACATTAAGCACTTACATCAAGGAGATCCATGGCCCCAAGAAGCAGAAACCATTCTTACTCCATGTCGCGATCGATCTTATGCTTCAGATTGCGCGAGGAATGGAATACCTGCATTCAAAGAAAATTTATCATGGAGAGCTAAATCCTTCAAACATTCTTGTTAAGCCAAGAGGTAACTTGCCTGATGATTACTTGCATGCCAAGGTAACTGGTTTCGGCCTATCTTCGGCCATGGAATCAAACCAGAAAGGGGGTACAAATCAAAATCAGAATGGAACTTCACCATTCATCTGGTATGCTCCAGAAGTACTTGAAGAGCAAGAAAATTCCGGGGGTGCCTTGCAATGCAAGTATTCAGAGAAATCTGATGTTTATAGCTTCGGAATGGTTTGCTTTGAGCTTCTAACAGGTAAAGTCCCTTTTGAAGATAGTCATCTACAAGGAGAGAAGATGAGTAGGAACATAAGGGCAGGGGAGAGGCCACTTTTCTCGCTAAATTCGCCGAAATATGTCATCAACTTGACAAAGAAATGTTGGCATACAGACCTGAATCAGCGTCCGAGCTTCTCTTCCATCTGTAGAATTCTCCGGTACATAAAACGGTTTCTTGCCATGAATCATAGTCATATCACTCAATTAGAGGCACCGGCGCCAACGCCGGCTATAGATTACTGTGACATAGAGACTGCACTTTTGAAGAAGTTTCCTACTTGGGGGAATTCTGAATCTACATCGATATCACAAATTCCTTTTCAAATGTTTTCCTATCGAGTCGCGGAACGAGAGAAAACAGGTACATACTCTAAGGAGAACTCTGAATCTGGAAGCGACGCTTCGGCGTGTGGAGATGATCTTGCTACCTCCGGAGATGAGGCATTCATATCATCCACAAATGAAAAGAAGATTTGCCCTGCAAATGAAAGCATGAGTACCAAGAAGCATTCACTAGCAAGGAAATCATTAGATTTTAATAAGCCAATAAAACAGCAAGTTACACCAAGATCAGCAAGGCCTCCGCAGATGTCGTCTTTTGGGCGGCCGAAGAGGACTAGTTCGGACCAGCAAACATCGAATTCAAGAACAAGAAGAACAGCTTCGGGTCATGTCTCGGATTCCGAGCTCTCTTAG

>A_ lyrata_1

ATGGAGCAATTCAGACAAATCGGCGAAGTTCTTGGAAGTCTTAATGCACTAATGGTATTACAAGATGATATCTTGATCAACCAAAGACAATGTTGTTTGTTGTTAGAACTCTTCAGTTTAGCTTTCAACACGGTCGCAGAAGAGATCAGACAGAATCTAAAGCTTGAAGAGAAGCATACTAAATGGAGAGCTCTTGAACAACCTTTAAGGGAGCTTTATAGAGTGTTTAAAGAAGGAGAGTTATATGTTAAGCATTGTATGGACAATAGTGATTGGTGGGGTAAAGTTATCAATCTTCATCAGAATAAAGATTGTGTTGAGTTTCATATACACAACTTGTTCTGTTATTTCTCTGCGGTTGTTGAAGCGATCGAGGCTGCTGGTGAGATTTCGGGTCTTGACCCTTCTGAGATGGAGAGAAGGAGAGTTGTGTTTTCAAGAAAGTATGATAGAGAGTGGAATGATCCTAAGCTGTTTCAATGGAGGTTTGGGAAACAGTATTTGGTATCGCGGGATATTTGCAGCCGGTTTGAGCATTCTTGGAGAGAAGATAGGTGGAATCTAGTGGAGGCATTACAAGAAAAGCGGAAATCGGATAGCGATGATATTGGTAAAACCGAGAAGCGTTTAGCGGATTTGCTGTTGAAGAAGTTAACCGGTTTGGAGCAGTTTAACGGTAAACTGTTTCCGAGTTCGATACTTCTTGGTTCAAAGGATTACCTGGTGAAAAGACGGTTAGATGCGGATGGACAATACAAGGAGATTCAATGGTTAGGTGATAGCTTTGCGGTGAGGCACTTTTTCAGCGATCTTGAGCCGCTAAGTTCTGAAATTTCTTCTCTGTTATCGCTTTGTCATTCGAATATACTTCAGTACCTTTGTGGATTCTATGATGAAGAAAGGAAAGAATGCTTTTTGGTTATGGAGTTAATGCACAAAGATTTGCAGAGCTACATGAAGGAGAATTGTGGACCAAGAAGAAGATATCTGTTCTCGATTCCCGTCGTGATCGATATCATGCTGCAAATTGCAAGGGGAATGGAATATCTTCATGGAAATGATATCTTCCATGGAGATTTAAACCCCATGAACATTCATTTGAAAGAGAGGAGTCACACCGAAGGTTATTTCCACGCGAAAATATCGGGATTCGGTTTGTCTTCAGTCGTTAAAGCGCAGTCTTCTCGGTCTTCCTCGAAACCAGGCACTCCTGATCCTGTAATCTGGTATGCACCTGAAGTTCTAGCAGAGATGGAACAAGACCTGAATGGTAAAACTCCGAAATCAAAGTTGACGCATAAAGCTGATGTGTATAGCTTTGCAATGGTGTGTTTTGAGCTTATAACAGGTAAAGTTCCATTTGAAGATAGTCATCTTCAAGGTGAACCAATGGCTATCAACATTAGAATGGGAGAGAGACCTCTTTTCCCTTTCCCTTCACCAAAATACCTCGTTAGTCTGATTAAACGGTGTTGGCACTCAGAACCGAGTCAGCGTCCTAACTTCTCTTCGATATGTCGGATACTTCGCTACATCAAGAAGTTCCTAGTTGTGAATCCAGATCATGGTCATCCTCAAATGCAAACTCCATTAGTTGATTGTTGGGACTTAGAAGCAAGGTTCTTGAGAAAGTTTCCAGGAGATGCAGGGTCTCACACCGCGTCCGTGAATCAAATCCCGTTCCAACTATACTCATACCGGGTTTCGGAAAGAGAGAAGATGAATCCAAACTCAAAAGAAAGCTCAGAAGCAAGTGATCAATCTGAAAGCGTTTCAGTGGTTGAAGATCCACCTAATGCCATTATTGCAAGAGATACAAAATCATTGTGTCTAGATACAATATCTGAATACTCAGATACAAGATCAGTCTACTCAGAAGCTCCCATCAAAAAGGTCTCAGCTTTAAAGAAAAGCGGCGAAATGGCAAAGCTCAGAAGAAGTCCAAGCTTAGGCTCAGAAAAGTTGAGATCAACAGGAACTTCACCGGTAAAAGCAAGATCATCGCCAAAAGTATCAGCATTGAGTCCATTTGGAAGAAGCATTAAAGCAAGGAAAGATAATCGATTGCCTTTGAGTCCAATGAGTCCTTTAAGCCCAGGAATACGTAGACAACATACTGGTCATGCTTCAGACTCTGAGCTTACTTAG

>A_ lyrata_2

ATGGAGCAATTCAGGCAAATCGGAGAGGTTCTTGGAAGTTTAAACGCGCTAATGGTATTACAAGACGATATCTTAATCAACCAAAGACAATGTTGTTTGTTGTTAGATATTTTCAGTTTGGCTTTCAACACCGTAGCCGAAGAGATCAGACACCACTTGAAGCTTGAAGAGAAGCATACTAAATGGAGAGCTCTTGAACAGCCTTTGAGAGAGCTTTATCGAGTTTTTAAAGAAGGCGAAATGTATGTTCGGAATTGTATGTCTAATAAAGATTGGTGGGGTAAAGTTATCAACTTTCATCAGAACAAAGATTGTGTTGAGTTTCACATTCACAATTTGCTTTGTTACTTCTCGGCTGTGATCGAAGCCATCGAGACAGCGGGTGAGATTTCGGGTCTCGACCCTGCTGAGATGGAGAGGAGGAGAGTTGTGTTTTCAAGAAAGTATGATAGGGAGTGGAATGATCCCAAGCTGTTTCAGTGGAGGTATGGGAAACAGTATTTGGTACCGAGGGATATTTGCATTCGGTTCGAGCATTCTTGGAGGGAAGATAGGTGGAATCTAGTGGAGGCATTGCAAGAGAAGAGGAAGTCCAAGAGCGATGAGATTGGGAAGACTGAGAAGCGTTTAGCTGATTTTCTGTTGAAGAAACTAACTGGATTGGAACAGTTTAACGGTAAGCTCTTCCCGAGTTCGATACTTGTTGGTTCTAAGGATTACCAAGTGAGGCGGAGATTAGGTGGTGGTGGTCAGTACAAGGAGATTCAATGGTTAGGGGATAGTTTTGTGTTGAGACATTTTTTCGGGGATCTTGAGCCGTTGAATGCAGAGATTTCTTCTCTGTTATCGCTTTGTCATTCGAATATACTTCAGTACCTTTGTGGATTCTATGATGAAGAAAGGAAAGAATGTTCTCTGGTTATGGAGTTAATGCACAAAGATTTGAAAAGCTACATGAAGGAGAATTGTGGACCTAGAAGAAGGTATCTTTTCTCTGTTCCCGTTGTGATCGATATAATGCTTCAAATCGCGAGAGGGATGGAATATCTTCATTCTAATGAGATTTTCCATGGAGATTTGAATCCCATGAACATTCTTTTGAAAGAGAGAAGTCACACCGAAGGTTATTTCCACGCGAAAATCTCAGGTTTCGGCTTGACCTCAGTTAAAAATCAGTCATTTTCTCGGGCTTCCTCTAGACCAACCACACCCGATCCTGTAATATGGTATGCACCTGAAGTTCTAGCAGAGATGGAACAAGATCTCAAAGGTACAGCTCCGAGATCAAAGTTTACACATAAAGCTGATGTCTATAGCTTTGCTATGGTGTGTTTCGAGCTTATAACCGGTAAAGTTCCATTTGAAGATGACCATCTTCAAGGCGAGAAAATGGCTAAGAACATAAGAACGGGAGAGAGACCTCTCTTCCCATTCCCTTCGCCAAAATACCTCGTTAGTCTAATTAAACGGTGTTGGCATTCAGAACCGAGTCAGCGTCCAACATTCTCTTCGATATGCCGGATCCTCCGCTACATCAAGAAATTCCTAGTTGTGAATCCGGATCACGGCCACCTCCAAATCCAAAACCCACTTGTTGATTGCTGGGACTTAGAAGCAAGATTCTTGAAAAAGTTCTCAATGGAGTCAGGTTCTCACGCAGGATCCGTGACACAAATACCATTCCAGCTCTACTCATACAGGATTGCAGAGAAAGAGAAGATGAGTCCAAACTTTAACAAAGAAGAGAATTCAGAAACAGGGGAATCTGTTTCTGAAGGTGTTTCAGTGGTTGAAGATCCACCAACAATGCCAATGTATACAAAGTCATTGTGCTTAGATGCAATATCTGAATACTCAGATATAAGATCAGTTTACTCAGAAGCTCCAATCAAAAAGACATCAGCTTTAAAGAAAAGTGGTGACACCATAAAAAACAGAAGAAACTCAATCTCAGGTTTACGGTCTCCAGGATCATCGCCTATAAAACCGAGATCAACACCGAAAGTGTCATCACCACTAAGTCCATTTGGAAGAAGCAGCAGCAAAGCAAGGAAGGATACAAGATTGCCTTTGAGTCCTATGAGTCCTTTGAGCCATGGAAGACGTAGACAACTCACTGGTCCTGCTTCAGACTCTGAGCTTACATAG

>A_ lyrata_3

ATGGATCAAAAGCGGCCAACCATCGCTGGCGATGAAGAAATGGCGGCTGAGAAACCGTTAAAACCAGCACTCCAAAAACCACCGGGGTTTCGTGATCAACAGAATCAACCATCAGCTCCGCCTTCCGGAACAGGTCGCCGTCCAAGGCCTATCCATCCGGCGTCACTCTACCCGGAGAAGAAACGGCGGTGTAGTTTCTGCCGCGTTTTCTGTTGCTGCGTGTGTATCTTCTTGGCCGTGATTCTCCTCATCTTTCTCATCGCTGTGGCGGTTTTCTTCCTTTGGTACAGCCCGAAACTACCCGTCGTCCGCCTAGCCTCATTCAAAATCAGCAATTTCAACTTTTCCGACGGAAACTCCGACGACGGATGGTCATTTTTGACGGCGGATACAACGGCGGTGCTCGATTTCAGAAACCCTAACGGAAAGCTAACGTTTTATTACAGAGATGCTGACGTGGCTGTGATTCTCGGAGAGAAAGACTTTGAGACTAACCTAAGGTCAACGAAAGTCAAAGGCTTTATTGAGAAGCCAGGAAATCGGACGGCTGTGATTGTCCCAACGAGGGTGACGAAACGACAGGTGGATGATCCAACGGCAAAGAGGCTACAGGCAGAGCTGAAGGGTAAGAAGCTGCTGGTGACAGTGACAGCTAAAACGAAGGTAGGATTGGCTGTGGGCAGTCGCAAGATTGTTGCTGTGGGCGTTAGTCTCAGATGCGGAGGCGTTAGATTACAGACCCTTGATTCGCAAATGGCCAAATGCACCATCAAATTGCTGAAATGGCCCTGTGTAAAATTGGAGAGGTGTGAAATGGAGCAATTTCGAGAGATAGGAGAGGTATTAGGAAGCATAAGAGCGTTAATGGTGTTCAAGGACGACATTCAAATCAACCAGCGTCAATGCTCTCTATTACTCGACCTCTTCACCGCGGCCTACGAGTCGATTTCCGAATCGATGAGATCGAATCTACGCTTCAAAGAGAAGAACACTAAATGGAAGATTCTAGAACAGCCTTTGAGGGAGCTTCTATGGGTGGTAAGAGAAGGAGAGGCTTATGTCAGAATGTCTTTAGAGCCTAAACTTGGGTTTTGGGCTAAGGCTATAGTGTTGCATTGCAATAGAGATTGCACAGAGCTTCACATACATAACTTGCTCTCTTGTCTACCGATCATTGTCGAGGCTATTGAGACAGCGGGTGAGGTTTCAGGGTGGGATGAAGAAGAGATGAGCAAGAAGAGGTTGGTGCATTCTAACAAGTATATGAAGCAATGGAATGACTCTCAGATGTTTACCTGGAAGTTCGGGAGAGAGTATTTGGTGACCGAAGATTTCTGCAACCGGTTCGAGAGTGCTTGGACAGAGGACCGTTGGATTCTAATCAAGGAGCTTCAGGAGAAAAAGCAGCCCAGTTTGAGCAAACATGATTGGAAGATGGCTGATTTCCTTTTGAAGCATTTGGGAGATGGTAATGAGAGTCCCAAGCTATTTCCGTCTTCTCTTCTGGTAAACACGAAAGATTACCAAGTGAAAAAAAGATTAGGGAATGGAAGTCAGTACAAGGAGATTACTTGGTTAGGCGAGAGCTTTGCGCTTAGGCATTTCTTTGGGGATATTGATACTTTGCTTCCCAAGGTTACTCCATTGCTATCTCTTTCACACCCAAACATTGTTTATTACCTCTGCGGATTCACGGATGAGGAGAAGAAAGAATGTTTCTTGGTCATGGAACTGATGAGTAAAACCCTTGGTATGCACATCAAAGAGGTATGTGGTCCGAGGAAGAAGAACACTCTCTCCCTCCCGGTCGCGGTTGATCTGATGCTTCAGATAGCTTTGGGTATGGAATATCTTCACTCGAAGAGAATATACCACGGAGAGCTGAATCCATCTAACATTCTTGTCAAACCAAGAAGTCACCAATCCGGAGATGGGTATCTGCTTGGGAAGATTTTTGGGTTCGGGTTGAATTCTGTCAAGGGCTTCTCTAGTAAAAGTGCTTCTTTGACGAGTCAGAATGAGAATTTTCCATTCATATGGTATTCTCCCGAAGTTCTAGAGGAGCAAGAACAGAGCGGAACAGCAGGAAGCCTCAAGTATACTGATAAATCTGACGTGTATAGCTTCGGGATGGTTTGCTTTGAGCTTCTGACAGGTAAAGTACCATTTGAAGATAGCCACCTGCAAGGAGATAAGATGAGTAGAAACATTAGGGCTGGGGAGAGACCGCTTTTTCCATTTAATTCACCTAAATTCATAACAAACCTGACAAAACGATGCTGGCATGCTGATCCGAATCAGCGTCCAACTTTCTCATCGATAAGCAGAGTTCTCCGGTATATCAAGCGATTTCTAGCTTTGAACCCAGAGTATCACAGTAGTAGCCAACAAGACCCGTCCATTGCTCCTCCTGTGGATTACTGGGAGATCGAGTCTAAGCTATTGCAGAAGCTTTCATGGGAATCTACAGAGTTAACTCAAGTTTCACAGGTTCCTTTTCAAATGTTTGCATATAGGGTGGTGGAACGGGCACAAACTTGTAAAAAGGATAATCTCCGGGAAACATCGGAATCAGGTAGTGAATGGGCTTCCTGTAGCGAGGATGAGGGTGGGGCAGGATCAGATGAGCAGGTGTCATATGAGAAGGAGAGAAGATTGTCATGCTCAAGTGATGTTGGTATGAGCAAGAAGCATGTTTCAAATCTTCTGAAAAGAGCTTCTAGCCTGAAACCAATTCAAAAACCAGCCTTTGGATTTAGTTCAGGAACAACTCCGAGAGGACGATCAAGGCATCCGCCACTAAGCCCATGTGGGCAAAGCATGAGAGCAAACTCTGAGAGTCAACTGATATTGATAAGCCCAAGGATACGACGATCAAACTCTGGCCATGCCTCTGACTCTGAACTTTCCTAG

>A_ thaliana_MLKL1_AT1G64300

ATGGAGCAATTCAGGCAAATCGGAGAGGTTCTAGGAAGCTTAAACGCGTTAATGGTATTACAAGACGATATCTTAATCAACCAAAGACAATGTTGTTTGTTGTTAGATATTTTCAGTTTGGGTTTCAACACCGTAGCCGAAGAGATCAGACACAACTTGAAGCTTGAAGAGAAGCATACTAAATGGAGAGCTCTTGAACAGCCTTTGAGAGAGCTTTATCGAGTTTTTAAAGAAGGCGAAATGTATGTTCGGAATTGTATGTCTAATAAAGATTGGTGGGGTAAAGTTATCAACTTTCATCAGAACAAAGATTGTGTTGAGTTTCATATACATAATCTGCTTTGTTACTTTTCGGCTGTGATCGAAGCCATCGAGACCGCGGGTGAGATTTCGGGTCTTGACCCTTCTGAGATGGAGAGGAGGAGAGTTGTGTTTTCAAGAAAGTATGATAGGGAGTGGAATGATCCCAAGCTGTTTCAGTGGAGGTTTGGGAAACAGTATTTGGTACCGAGGGATATTTGCTTACGGTTCGAGCATTCGTGGAGAGAAGATAGGTGGAATCTCGTTGAGGCATTGCAAGAGAAGAGGAAGTCCAAGAGCGATGAGATTGGCAAGACTGAGAAGCGTTTAGCTGATTTTCTGTTGAAGAAACTAACGGGACTAGAACAGTTTAATGGTAAGCTCTTCCCCAGTTCGATACTTGTTGGTTCTAAGGATTACCAAGTGAGGAGGAGATTAGGTGGTGGTGGTCAGTACAAGGAGATTCAATGGTTAGGGGATAGTTTTGTGTTGAGACATTTTTTCGGTGATCTTGAGCCGTTGAATGCAGAGATTTCTTCTCTGTTATCGCTTTGTCATTCGAATATACTTCAGTACCTTTGTGGATTCTATGATGAAGAAAGGAAAGAATGTTCTCTTGTTATGGAGTTAATGCACAAAGATTTGAAAAGCTACATGAAGGAGAATTGTGGACCTAGAAGAAGATATCTTTTCTCTGTTCCCGTTGTTATCGATATAATGCTTCAAATCGCGAGAGGGATGGAGTATCTTCATTCAAATGAGATTTTCCATGGAGATTTGAATCCAATGAACATTCTTTTGAAAGAGAGAAGTCACACCGAAGGTTACTTCCACGCGAAAATCTCGGGTTTCGGTTTAACCTCAGTTAAGAATCAGTCTTTTTCTCGGGCTTCCTCTAGACCAACCACACCTGATCCTGTAATATGGTATGCACCTGAAGTTCTAGCAGAGATGGAACAAGATCTCAAAGGTACAGCTCCAAGATCAAAGTTTACACATAAAGCTGATGTCTATAGCTTTGCTATGGTGTGTTTTGAGCTTATAACCGGTAAAGTTCCATTTGAAGATGACCATCTTCAAGGCGATAAAATGGCTAAGAACATAAGAACGGGAGAGAGACCTCTCTTCCCATTCCCTTCACCAAAATACCTTGTTAGTCTAATCAAACGGTGTTGGCATTCAGAACCGAGTCAGCGTCCAACATTCTCTTCGATTTGCCGGATCCTCCGCTACATCAAGAAGTTCCTAGTGGTGAATCCGGACCACGGTCACCTTCAAATCCAAAACCCTCTAGTTGATTGCTGGGATTTAGAAGCAAGATTCTTGAAAAAGTTCTCAATGGAGTCAGGTTCTCACGCAGGATCAGTGACACAAATACCATTCCAGCTTTACTCATACAGGATTGCAGAGAAAGAGAAGATGAGTCCAAACTTTAACAAAGAAGAGAATTCAGATGCAGGGGAATCTGTTTCTGAAAGTGTTTCAGTAGTTGAAGAGCCACCAACAACAATGCCAATGTATACTAAGTCATTATGCTTAGATGCAATATCTGAATACTCAGATACAAGATCAGTCTACTCAGAAGCTCCAATCAAAAAGACATCAGCTTTAAAGAAAAGTGGTGACACCATCAAAAACAGAAGAAATTCAATCTCAGGTTTAAGGTCTCCAGGATCATCGCCTATAAAACCGAGATCAACACCTAAAGTATCATCACCACTAAGTCCATTTGGAAGAAGCAGCAGCAACAAAGCAAACAAGGATACAAGATTGCCTTTGAGTCCTATGAGTCCACTGAGCCATGGAAGACGTAGACAACTAACTGGTCCTGCTTCAGACTCTGAGCTTACATAG

>A_ thaliana_MLKL2_AT5G41730

ATGGAGCAATTCAGACAAATCGGAGAAGTTCTTGGAAGTCTAAATGCACTAATGGTATTACAAGATGATATCTTGATCAACCAAAGACAATGTTGTTTGTTGTTAGAACTCTTCAGCTTAGCTTTCAACACGGTCGCAGAAGAGATCAGACAGAATCTAAAGCTTGAAGAGAAGCATACTAAATGGAGAGCTCTTGAACAACCTTTAAGGGAGCTTTATAGAGTTTTTAAAGAAGGAGAGTTATATGTTAAGCATTGTATGGATAATAGTGATTGGTGGGGTAAAGTAATCAATCTTCATCAGAACAAAGATTGTGTTGAGTTTCATATACACAACTTGTTCTGTTATTTCTCTGCGGTTGTTGAAGCCATCGAGGCTGCTGGTGAGATTTCGGGTCTTGACCCTTCTGAGATGGAGAGAAGGAGAGTTGTGTTTTCAAGAAAGTATGATAGAGAGTGGAATGATCCTAAGATGTTTCAATGGAGGTTTGGGAAACAGTATTTGCTCTCGAGAGATATTTGCAGCCGGTTTGAGCATTCTTGGAGAGAAGATAGATGGAATCTAGTGGAGGCATTGCAAGAGAAGAGGAAATCGGATAGCGATGATATTGGTAAAACCGAGAAGCGTTTAGCTGATTTGCTGTTGAAGAAGTTAACCGGTTTGGAGCAGTTTAACGGTAAACTGTTTCCTAGTTCGATACTTCTTGGTTCAAAGGATTACCAGGTGAAGAAACGGTTAGATGCAGATGGACAATACAAGGAGATTCAATGGTTAGGTGATAGTTTTGCACTAAGGCACTTTTTCAGCGATCTTGAGCCGCTAAGTTCTGAAATCTCGTCTCTTTTAGCGCTTTGTCATTCGAATATACTTCAGTACCTTTGTGGATTCTATGATGAAGAAAGGAAAGAATGTTTTTTGGTTATGGAGTTGATGCACAAAGATTTGCAGAGCTACATGAAGGAGAATTGTGGACCAAGAAGAAGATATTTGTTCTCGATCCCTGTGGTGATTGATATCATGTTGCAAATTGCAAGGGGGATGGAGTATCTTCATGGAAATGATATCTTCCATGGAGATTTAAACCCAATGAACATTCATCTGAAAGAGAGGAGTCACACCGAAGGTTACTTCCATGCGAAGATTTGTGGATTCGGTTTATCTTCAGTCGTTAAAGCGCAGTCTTCCTCGAAACCAGGCACTCCTGATCCTGTAATATGGTATGCACCTGAAGTTCTAGCAGAGATGGAACAAGACCTGAATGGTAAAACTCCGAAATCTAAGTTGACGCATAAAGCTGATGTGTATAGCTTTGCAATGGTGTGTTTTGAGCTTATAACAGGTAAAGTTCCATTTGAAGATAGTCATCTTCAAGGTGAACCAATGACTATCAATATTAGAATGGGAGAGAGACCTCTTTTCCCTTTCCCTTCACCAAAATACCTCGTTAGTCTGATCAAACGATGTTGGCACTCAGAACCGAGTCAGCGTCCTAACTTCTCTTCGATATGTCGGATACTACGCTACATCAAGAAGTTCCTAGTTGTGAATCCAGATCATGGTCATCCTCAAATGCAAACTCCACTTGTTGATTGTTGGGACTTAGAAGCAAGGTTCTTGAGAAAGTTTCCAGGTGATGCAGGGTCTCACACCGCGTCAGTGAATCAAATACCGTTCCAACTATACTCATACCGGGTTTTGGAAAAAGAGAAGATGAATCCAAACTCAAAAGAAAGCTCAGAGACTAGTGAATCTGAAAGCGTTTCAGTGGTTGAAGATCCACCTAATGCCATGATTACAAGAGATACAAAGTCATTGTGTCTTGATACAATATCTGAATACTCAGATACAAGATCAGTCTACTCGGAAGCTCCCATGAAAAAGGTCTCAGCTCTAAAGAAAAGCGGCGAAATGGCAAAGCTCAGAAGAAGTCCAAGCTTAGGTTCAGAAAAGTTGAGATCGGCAGGAACTTCAACGGTAAAAGCAAGATCATCGCCAAAAGTATCACCACTGAGTCCATTTGGAAGAAGCATTAAAGCAAGGAAGGATAATCGATTGCCTTTGAGTCCAATGAGTCCTTTAACCCCTGGAATACGTAGACAACAGACTGGTCATGCTTCAGACTCTGAGCTTACTTAG

>AtMLKL3

ATGGATCAATTTCGAGAGATAGGAGAGGTATTAGGAAGCATAAGAGCGTTAATGGTGTTCAAGGACAGCATTCAAATCAACCAGCGTCAATGCTCTCTATTACTCGACCTCTTCACCGCAGCCTACGAGTCAATTTCTGTGTCGATGCGATCCAATCTACGCTTCAAAGAGAAGAACACTAAATGGAAGATACTAGAACAGCCCTTAAGGGAGCTTCTATGGGTGGTTAGAGAAGGAGAGGCTTATGTTAGAATGTCTTTAGAGCCTAAACTTGGGTTTTGGGCTAAGGCTATTGTGTTGCATAGCAATAGAGATTGCACAGAGCTTCACATACATAACTTGCTCTCTTGTCTACCGATCATCGTCGAGGCTATTGAGACAGCGAGTGAGGTTTCAGGGTGGGATGAAGAAGAGATGAGCAAGAAGAGGTTGGTGCATTCTAACAAGTATATGAAGCAATGGAATGACTCTCAGATGTTTACCTGGAAGTTTGGGAGAGAGTATTTGGTGACCGAGGATTTCTGCAACCGGTTCGAGAGTGCTTGGACAGAGGACCGTTGGATTCTGATCAAGGAGCTTCAGGAGAAAAAGCAGTCCGGTTCGAGCAAACATGAACGGAAGATGGCTGATTTCCTTTTGAAGCATTTGGGAGATGGAAATGAGAGCCCCAAGCTATTTCCTTCTTCTCTTCTGGATAACACAAAAGATTACCAAGTGAAAAAAAGATTAGGGAATGGAAGTCAGTACAAGGAGATTACCTGGTTAGGCGAGAGCTTTGCGCTTAGGCATTTCTTTGGGGATATTGATGCTTTGCTTCCTCAGATTACTCCATTGCTATCTCTTTCACACCCAAACATTGTATATTACCTCTGCGGATTCACGGATGAGGAGAAGAAAGAGTGTTTCTTGGTCATGGAACTGATGAGGAAAACCCTTGGAATGCACATCAAAGAGGTATGTGGTCCTAGGAAGAAGAACACTCTCTCCCTCCCGGTCGCGGTTGATCTGATGCTTCAGATAGCTTTGGGTATGGAATATCTTCACTCGAAGAGAATATACCACGGAGAGTTGAATCCATCAAACATTCTTGTCAAACCAAGAAGTAACCAATCCGGAGATGGGTATCTGCTTGGGAAGATTTTTGGGTTCGGGTTGAATTCTGTCAAGGGATTCTCTAGTAAAAGTGCTTCTTTGACGAGTCAGAATGAGAATTTTCCATTCATATGGTATTCTCCAGAAGTTCTAGAGGAGCAAGAACAGAGTGGAACTGCAGGAAGCCTTAAGTATAGTGATAAATCTGACGTGTATAGCTTTGGGATGGTTAGCTTTGAGCTTCTGACAGGTAAAGTACCATTTGAAGATAGCCACCTGCAAGGAGATAAGATGAGTAGAAACATTAGGGCCGGGGAGAGACCGCTTTTTCCATTTAATTCACCAAAGTTCATAACAAACCTGACAAAACGATGCTGGCATGCTGATCCCAATCAGCGTCCAACTTTCTCATCGATAAGCAGAATTCTCCGGTATATCAAACGGTTTCTAGCTTTGAACCCGGAATGTTACAGTAGTAGCCAACAAGACCCGTCCATTGCGCCTACTGTGGATTACTGCGAGATCGAGACAAAGCTATTGCAGAAGCTTTCTTGGGAATCTACAGAGCTAACCAAAGTTTCACAGGTTCCTTTTCAAATGTTTGCATATAGGGTTGTGGAACGGGCAAAAACGTGTGAAAAGGATAATCTCCGGGAACCATCGGAATCAGGAAGTGAATGGGCTTCGTGTAGCGAGGATGAGGGTGGGGCAGGATCAGATGAGCAGTTGTCATATGCCAAGGAAAGAAGATTGTCATGCTCCTCAAATGATGTTGGTATGAGCAAGAAGCAGGTTTCAAATCTTCTGAAAAGAGCTTCTAGCCTGAAACCAATTCAAAAACCAGCCTTTGGATTCTGTTCAGGGACAACTCCGAGAGGACGATCAAGGCATCCGCCACTAAGCCCTTGTGGTGGGCAAAGCATGAGAGCAAACTCTGAGAGTCAGCTGATATTGATAAGCCCAAGGATACGACGATCAAACTCTGGACATGCCTCTGACTCTGAGCTTTCCTAG

>B_distachyon_1

ATGGAACAGCTCCGTCAGGTCGGCGAGGCCCTGGGCGGCATCACGGCGCTCATGGCGTTCCACCACGAGCTCCGCGTGAACCCGCGCCAATGCCGCCTCCTCGCCGACGCCTGCGCGCTCGCCTTCGACGCCGTGGCCGCCGAGGTGAGCGCCCACCTCCGCCTTGAGGACAGATGGAGGCCCCTCGAGCACCCTCTCCGTGAGCTCCACCGCGCCGTCCGCGACGCCGAGCTCTACATCCGCCACTGCCTCCTCGGAGGCGCCGGCGCATCGTGGTGGGCGCGCGTGGCCGCGGCCACGCATGGCGACGAGTGCGTCGAGCACCACCTCCATGCCATCCTCTGGTGCGTCGCCGTCGTCCTTGAGGCCATCGAGACCGCCGCCACGGAGACCACGGCATCCTCGTGCGACGACGATCTTGCCCGGAGCAGCCGGCTGATGTTCGCCAGGGATTACGACAAGGAGCTCCTCGACCCGGCCCTGTTCCGGCGAAGCAGAGTCGGGAAAGCCTACATGGCCACCAAGGACCTCGCTGCCAGGATGGACATGGCGTGGAAGGAGGACCGCTGGCTCCTCTCGCAGCTGCTCGACGAAATGAAGCACGGCAGCTCGGCATCTTCTTCTTCCTCGAAGCCGTTGTCCCGGCACGAGCACCGGCTGGCCGACCTCCTCGCCGCGCCACACGGGAAGCTGCACCCGGCGTCGGTGCTCGCCCACGACTTCCACGTGCGGCGCCGCCTCGGAGGCTGCCTGAAGGAGGCGCATTGGATGGGGGAGCCATTCGCCGTGAAGCACTACGTCGGCATCGACGCCGACGCGGCGGACGCCGAGGCCGGCACGCTCATGGCGGCCGCGGCGCACCCGAACGTGGCGGGATGCCGGTTCTGCTTCCAAGACGAGGAGAAGCGGGAGCTGTTCGTCGTCATGGACGACCAGCTGATGACAAAAGATCTCGGAAGCTACGTCAAGGAGCAGGTCAGCAAGCGGCGGGCCACGCCGATGCCGCTGGTCGTCGTCGTGGACGCCATGCTCCAGATCGCGCGCGGCATGGAGCACCTCCACGCCAAGAAGATCTTCCATGGCGAGCTCAGCCCGGCCAACGTGCTCGTCAAGCCACGGCACGGCAATTCCGCCGACGCCGGCTACCTGCTCGTCAAGGTCGCTGGCTTTAACCGTGAGCCCGCCGTGCCTGCCAGCCCCACCCGGAAGACATCGGCGCCGGCCAACAATGCCAATGCCAATGCCAGTGTCAACCCGTGCATCTGGTACGCGCCGGAGGTGCTGGAGCAGGAGACGGAGAAGCGCACGGAGAAGGCGGACGTGTACAGCTTCGCCATGATCTGCTTCGAGCTGATCACAGGGAAGATTCCCTTCGAGGACCATCACCTGCAAGGGGAGCACATGAGCAAGAACATCCGCGCGGGCGAGCGGCCGCTGTTCCCGTTTAATTCGCACAAGCAGCTCACGGGGCTCACGAGGCGCTGCTGGCAAGCCGACCCGGCGCAGCGGCCGGCATTCGGCTCCATCTGCCGCGTGCTCCGCTACGTCAAGCGGTTCCTCGTCATGAACCCGCAGCCGGACCAGCAGCAGCAGCCTGACTCGCCGACAATGCCGATGCCGATGCCGGCGGTGGACTACCTCGACATTGAGGCGCTGCTGCTGAGGAAGTTCCCGGCGTGGGAGGGCGCGGCGCCGAGGGTGGCTGACGTGCCGTTCCAGATGTACGCGTACAGGGTGATGGAGAAGGACAGGAGCAGCAGCAAGACGACACCGACGATGGCGGCATCGGCGGCGGCGATGCTGCACATTGGGAAGATGGACAGGAGCTCTGACTCCGGCAGCGACGGCAACTCGCTGTGCGGCGACGAGAGCGTCCACAGCGCGCCCGACGGCGTCGTCGAGGCCGGCTCGATGACGTCGAGCCGCGCCACCACGACGCCGCGGTCGCTGTCGGGCCGGAGCAGCGGCAGCAGCAGGATGGCGGCGGTGTCGACGTCGTCCACGACGTCCCCGCCGCGCAGGCCGGCAGCCAGGGTCGCCTTCAGCAAAGGAGGGTCGCCGCAGAAGTCCAAGTCGATGGTGATCCGGGCGCCGGCGCAGCCAAGCACGCCCCGACGGACTCCGAGGATAAAGTCCGACGGGCAGCTGCAGGCAGCTCTAATTCCGCCCTCCCGACGGCGTGTGTCCAGTGGGCATGCTTCAGACTCCGAATTAGCGTAG

>B_distachyon_2

ATGGAGCAGCTCCGTCAGCTCGGCGAGGCGGTGGGAAGCATCAACGCGCTCATGGCGTTCGAGCCGGACATCCGCATCAACCCTCGGCAGTGCCGCCTCCTGGCGGACGTCTGCGCGCACGCGCTGGACGCCGTGACCGGCGAGGTCCGCGCCCACCTCCGTTTCGACGAGCGCGGCACCACCAAGTGGCGCGCCCTCGACGCCCCGCTCCGGGAGCTCCACCGCGTGCTCCGCGACGCCGATGGCTACGTCCGGCAGTGCCTCGACCCGCGCCCAGGCAGCTGGTGGGGCCGGGCCGCGGCCATGGCGCACGGCACGGACTGCGTCGAACACCTCCTCCACAACGTCCTGTGGTGCGTCTCGGTCGCGATCGAGGCCGTCGAGACCGCCGCGGAGGTCACCGCGGGCTCCGACGCGGACGATCTCGCGCAGCGGACGCGGGTGCTGCTCGCCAAGAAGTACGACGGGGACATGCTCGAGCCCAAGACGTTCCAGCACGCGCACGGTAAGCTGTATCTGGTCTCCCGAGAACTCGTCGCTCGGATGGATGCCGCGTGGAAGGAGGACAGGTGGGTGCTGGCGCAGCTGTTCGACGAAATGACGGGCCCGGCGGCGCCGAACAAGCGCCTGACGAAGAACGAGCACCGCCTCGCCGAGGTCCTGGCCGCGCCGAGGGGGACGCTGCACCCGGCGTCCATTCTGCTTGGCGGCGACTACAGCGTCCGGAGGCGGCTCGGCGGCCGGCTCAAGGAGGCGCAGTGGATGGGGGAGAGCTTCGCGGTGAACCATTTCATCGGGGGCGGCGAGGCGGTCAGCGCCGAGGTCGCGCTCCTGTCCTCGGTGGCGCACCCTAACGTGGCACACGCCTCGTACTGCTTCCACGACGAGGAGCGGAAGGAGTACTTTGTCGTGATGGACCAGCTCATGGCAAAGGACCTGGGGAGCTACATCAAGGAAATGAGCTCCCCACGCCGGCGGACGCCGTTCACTCTGGCCGTCGCCGTGGACATCATGCTGCAGATCGCTCGCGGGATGGAGTACCTGCACGGCAAGAAGATTTACCACGGCGAGCTGAACCCGTCCAATGTGCTCGTCAAGCCGCGGCAACCCGACGGCGGCTACGTGCACGTCAAGGTCGCCGGGTTCGGGCAGTCGGATGGCACAAAAGCGTCTGCTAACGCCAACGCCAACGGCGACGACAACACCTGCATCTGGTACGCGCCGGAAGTGCTCAAGCCGGAGGGCGTACCCGTGGCGGACGCGGAGGCCAGGTGCACGGAGAAGGCCGACGTGTACAGCTTCTCGATGATCTGCTTCGAGCTGCTGACCGGCAAGGTCCCGTTCGAGGACAACCACCTGCAGGGCGACAAGACGAGCAAGAACATCCGCGCCGGCGAGCGGCCGCTGTTCCCGTTCCAGACGCCCAAGTACCTCACCGCGCTGACCAAGCGGTGCTGGCACGCCGACCCGGCGCAGCGGCCGGGCTTCTCCTCCGTCTGCCGTGTCCTCCGGTACGTCAAGCGGTTCCTTGTCATGAACCCGGAGAAGGAGAACCAGCAACAGCAGCAGCAGGCTGGCCAGCAGGCCGACGCGCCCGTGGCGCCGCCCGTGGACTACCTCGACGTCGAGATGCAGCTGCTGAGGAGGCTCCCGGCGTGGCAGCGAGGCGAGGGCGCGCGCGTCTCGGACGTCCCGTTCCAGATGTTCGCGTACAGGGTCGTGGAGAGGGAGAAGACTGCGGCCACCGTGCACGCCAAGGACAGGGCATCCGACTCAGGCAGCGAAGGGAACTCGCTGTACGGCGACGAGAACGGCGTCGTGGCGATGTCTCCGGACCACGCGGCGGCCCCCGTGTCGAGCGTCACCGTTCGGTCGGTGCCGGATAGCAGCGACGGCAAGAAGCTGCCGTCGGCCAAGAAGGCGGACGGCAGCAACAGCAAGGCGTCCAAGCAAGCAGGGTCTGCGCAGAAGGTGAAGGCCGCAAATCAGGCGAAGGCTCTGCAGCCACCAAGACGAACGCTCGGCGTGAAGACCGAAGGCATGTAA

>B_oleracea_1

ATGGAACAGTTCAGACAAATTGGCGAAGTTCTTGGAAGTCTAAATGCTCTTATGGTATTACAAGACGATATCTTGATCAACCAAAGACAATGTTGTCTGCTGTTAGAGATCTTCAGCTTAGCTTTCACCACGGTCTCCGAAGAGATCAGACAGAATCTAAAACTGGAAGAGAAGCACACCAAATGGAAAGCCCTTGAACAGCCCTTGAGAGAGCTTTACAGAGTGTTTAAAGAAGGAGAGTTATACGTCAAGCATTGCATGGACAATAGTGACTGGTGGGGCAAAGTCATCAATCTTCATCAGAACAAAGATTGCGTCGAGTTTCACATACACAACTTGTTCTGTTATTTCCCGGCCGTTGTTGAAGCGATCGAGGCTGCGGGAGAAATCTCGGGCCTCGACCCTTCTGAGATGGAGAGAAGGAGAGTTGTGATCTCGAGAAAGTACGATAGAGAGTGGAATGATCCTAAGCTGTTTCAATGGAGGTTTGGGAAACAGTATCTTGTTCCAAAGGATGTCTGCAGCCGGTTCGAGAATTCGTGGAGAGAAGACAGATGGAAACTAGTGGAGGCATTGCAAGAGAAGAGGAAATCAAACACCGACGAGATCGGAAAGACGGAGAAGCGTTTAGCTGATCTGCTTTTGAAGAAGCTAACCGGTTTGGAGCAGTTTAACGGTAAACTGTTTCCGAGTTCGATACTTCTCGGCTCAAAGGATTACCAGGTGAAAAGGAGGTTAGATGGAGACGGACATTACAAGGAGATTCAATGGTTTGGTGATAGTTTTGTTGTGAGACACCTCTTCAGCGATCTTGAGCCGTTGAGTTCAGAGATTTCATCTCTTTTAGCACTTTGTCATTCGAATATACATCAGTACCTGTGTGGATTCTATGACGAAGAGAGGAAAGAATGTTTTTTGGTTATGGAGTTGATGCACAAAGATTTGCAGAGCTACATGAAGGAGAATTGCGGACCGAGAAGAAGGTATCTCTTCTCCGTTTCCGTGGTGGTTGATATCATGATGCAGATAGCGAGAGGGATGGAGTATCTCCATGGGAATGATATCTTCCATGGAGATTTAAACCCGACGAATATTCTCTTGAAAGAGAGATGTCACACCGAAGGTTATTTCCACGTGAAAATCTCAGGTTTCGGTTTATCTTCGGTCAAATCTCAGTCTTCTTCTTCCTCGAGAGCAAGCACTCATGATCCTGTGATATGGTATGCACCTGAAGTTCTAGCGGAGATGGAACAAGGCACAAAAACTCCGAAATCGAAGCTGACGCATAAAGCCGATGTGTATAGTTTTGCGATGGTTTGTTTTGAGCTGATAACAGGTAAAGTACCTTTTGAAGATAGTCATCTCCAAGGTGAGCAAATGGCTATCAACATTAGAATGGGAGAGAGACCTCTCTTCCCTTTCCCTTCACCAAAGTACCTCGTTTGTCTGATCAAACGGTGTTGGCACTCGGAACCGAGCCAGCGTCCGAACTTCTCTTCCATCTGCCGGATACTACGTTACATCAAGAAGTTCCTAGTTGTGAATCCGGACCATGGTCACCCTCAGATTCAAACTCCACTAGTTGACTGTTGGGATTTGGAAGCAAGGTACTTGAGAAAGTTCCCAGGTGAGTTAGGGTCCCACGTGGCGTCGGTGACTCAAATTCCTTTCCAGCTGTACTCATACCGGGTTTTGGAAAGAGAGAAGATGAATCCAACAAACTCAAAAGAAAGCAGTAGTACAGAGGCAAGCGAATCAGAAGTTGAAGATCCTTCACCTAATGCAGGGATAATAAGAGATACAAAGTCTTTGTGTTTAGATACAATATCTGAATACTCAGATACAAGATCAGTCTACTCAGAAGCTCCCATCAAAAAGGTCTCAGCTTCAAAGAACAGTGAAGAATTGGTCAAGCTAAGAAAAAGTCCAAGCTTAGGGTCGTCAACAAAGTTGAGATCAACCGGAACTTCGCCGGTAAAAGCAAGATCATCACCAAAAGCATTACCCTTGAGTTCATTTGGAAGGAGCATAAAAACGAGGAAGGATAGTCGATTGCCTTTGAGTCCAATGAGTCCACTAAGTCCAAGGAGACGTAGACTACAAACTGGTCATGCTTCTGACTCGGAGCTTCCTTAG

>B_oleracea_2

ATGGAGCAATTCAGGCAAATCGGCGAGGTTCTTGGAAGCTTAAACGCGCTTATGGTGTTACAAGACGATATCTTGATCAACCAAAGACAATGCTGTTTGTTGTTAGAGATTTTCAGTTTGGCGTTCAACACCGTAGCGGAAGATATCAGAGAAAACTTGAAGCTCGAAGAGAAGCATTCCAAATGGAGAGCCCTTGAACAGCCTTTGAGAGAGCTCTACAGAGTGTTTAAAGAAGGTGAAACCTATGTTCGAAGCTGCATGTCTAATAAAGATTGGTGGGGCAAAGTAATCAACTTTCATCAGAACAAAGATTGTGTTGAGTTTCACATACACAACTTGTTCTGTTACTTTCCCGCCGTGATCGAGGCGATAGAGACAGCTGGAGAGATTTCTGGTCTCGACCCTTCTGAGATGGATAGAAGGAGAGTTGTCTTCTCTAGAAAGTACGATAGAGAGTGGAATGATCCTAAGCTGTTTCAGTGGAGGTTTGGGAAACAGTATCTGGTTCCCAAGGATATTTGTAGCCGCTTTGAGCATTCATGGAGAGAAGACAGATGGAATCTAGTTGAGGCGTTACAAGAGAAGAGAAAGTCCAAGAGCGATGAGATTGGGAAAACCGAGAAGCGTTTAGCTGACTTTCTGTTGAAGAAACTAACCGGTTTGGAACAGTTTAACGGTAGGCTCTTCCCGAGTTCGATATTCGTTGGCTCTAAAGATTACCAAGTGAGGAGGAGACTAGGCGGTGGTGGTCAGTACAAGGAGATTCAATGGTTAGGTGATTCTTTTGTCCTGAGACATTTTTTCGGAGATCTTGAGCCGTTGGATGCAGAGATATCTTCTCTGTTATCGCTATGTCACTCGAATATACTTCAGTACCTATGTGGATTCTATGACGAAGAGAAGAAAGAATGTTCTCTGGTAATGGAGTTAATGCACAAAGACTTGAAAAGCTACATGAAGGAGAACTGTGGACCTCGAAGAAGATATCTCTTCTCTGTTCCCGTTGTAATTGATATAATGCTTCAGATCGCGCGAGGCATGGAGTATCTTCATTCAAACGAGATCTTCCATGGAGACTTGAATCCAATGAATATTCTTTTGAAAGAGAGAAGTCACACCGAAGGTTACTTCCACGCCAAGATATCCGGTTTCGGTTTGAACTCAGTCAAAACTTTTACTCGAGCTTCCTCTAGACCAACCACTCCTGCTCCTGTGATTTGGTATGCACCTGAGGTTCTAACAGAGATGGAACAAGATCTCAAAGGTATAACAGTTCCGAGATCAAAGTTTACTCATAAAGCTGATGTGTATAGCTTTGCTATGGTGTGTTTTGAGCTTATAACCGGTAAGGTACCATTTGAAGATAGTCATCTACAAGGAGATGAGATGGGTAAGAACATAAGAAGGGGAGATAGACCTCTCTTCCCATTCCCTTCTCCTAAATACCTCGTTAGTCTTATCAGACGGTGTTGGCACTCAGAACCTAGCCAGCGTCCGACTTTCTCTTCCATTTGCCGGATACTCCGCTACGTAAAGAAGTTCTTAGTTGTGAATCCGGATCAGGGTCACATCCAAATCCAAACTCCGCTTGTTGATTGCTGGGACTTGGAAGCAAGATTCTTGAAAAAGTTCTCAATCGAGACAGGGTCTCACGCGGAGTCCGTGATGCAAATACCGTTCCAGCTTTACTCGTATAGAGTCGCAGAGAAGGAGAAGATGAGTCCAAACTTGAGCAAAGAAGAGAGTTCGGACACAGGAGGTGAGTCTGCTTCAGAAAGCGTTTCGGATCCACCAACAACAACGCCAAAGTATACAAAGTCATTGTGCTTAGATGCAATATCTGAATACTCAGAGTCAGATACAAGATCAATTTACTCAGAAGCTCCTAAGAAAAAGATCTCACCAGCTTCAAAGAAAAGTGGTGACATGGCCAAACTCAGGAGAAACTCAAGCGCAGGTTTACGGTCAACAGGATCATCGCCAGTAAAGCCAAGACCAGCACCGAAAGTGACATTGCCGTTAAGTCCGTTTGGAAGAAACAGTAAAGCAAGGAAGGATACAAGATTGCCTTTGAGTCCTATGAGTCCTTTAGGCCATGGAAGACGCAGACATCTCTCTGGTCCTGCTTCGGACTCTGAGCTAACTTAG

>B_oleracea_3

ATGGAGCAGTTCAGGCAAATCGGCGAGGTTCTCGGAAGCTTAAACGCGCTTATGGTATTACAAGACGATATCTTGATCAACCAAAGACAATGCTGTTTGTTATTAGACATTTTCAGTTTGGGTTTCAACACCGTAGCCGAAGAGATCAGACAAAACTTGAAGCTCGAAGAGAAGCATACCAAATGGAGAGCCCTTGAACAGCCTTTGAAAGAGCTCTACAGAGTGTTTAGAGAAGGTGAAACCTATGTTCGAAGCTGCATGTCTAATAAAGATTGGTGGGGCAAAGTAATCAACTTTCATCAGAACAAAGATTGTGTTGAGTTTCACATACACAACTTGTTCTGTTACTTTCCCGCCGTGATCGAGGCGATAGAGACAGCTGGAGAGATTTCTGGTCTCGACCCTTCTGAGATGGATAGAAGGAGAGTTGTCTTCTCTAGAAAGTACGATAGAGAGTGGAATGATCCTAAGCTGTTTCAGTGGAGGTTTGGGAAACAGTATCTGGTTCCCAAGGATATTTGTAGCCGCTTTGAGCATTCATGGAGAGAAGACAGATGGAATCTAGTTGAGGCGTTACAAGAGAAGAGAAAGTCCAAGAGCGATGAGATTGGGAAAACCGAGAAGCGTTTAGCTGACTTTCTGTTGAAGAAACTAACCGGTTTGGAACAGTTTAACGGTAGGCTCTTCCCGAGTTCGATACTCGTTGGCTCTAAAGATTACCAAGTGAGGAGGAGACTAGGCCGTGGTGGTCAGTACAAGGAGATTCAATGGTTAGGTGATTCTTTTGTCCTGAGACATTTTTTCGGAGATCTTGAGCCGTTGGATGCAGAGATATCTTCTCTGTTATCGCTATGTCACTCGAATATACTTCAGTACCTATGTGGATTCTATGACGAAGAGAAGAAAGAATGTTCTCTGGTAATGGAGTTAATGCACAAAGACTTGAAAAGCTACATGAAGGAGAACTGTGGACCTCGAAGAAGATATCTCTTCTCTGTTCCCGTTGTAATTGATATAATGCTTCAGATCGCGCGAGGCATGGAGTATCTTCATTCAAACGAGATCTTCCATGGAGACTTGAATCCAATGAATATTCTTTTGAAAGAGAGAAGTCACACCGAAGGTTACTTCCACGCCAAGATATCCGGTTTCGGTTTGAACTCAGTCAAAACTTTTACTCGAGCTTCCTCTAGACCAACCACTCCTGCTCCTGTGATTTGGTATGCACCTGAGGTTCTAACAGAGATGGAACAAGATCTCAGAGGTATAACAGTTCCGAGATCAAAGTTTACTCATAAAGCTGATGTGTATAGCTTTGCTATGGTGTGCTTTGAGCTTATAACCGGTAAGGTACCATTTGAAGATAGTCATCTACAAGGAGATGAGATGGGTAAGAACATAAGAAGGGGAGATAGACCTCTCTTCCCATTCCCTTCTCCTAAATACCTCGTTAGTCTTATCAGACGGTGTTGGCACTCAGAACCTAGCCAGCGTCCGACTTTCTCTTCCATTTGCCGGATACTCCGCTACGTAAAGAAGTTCTTAGTTGTGAATCCGGATCAGGGTCACATCCAAATCCAAACTCCGCTTGTTGATTGCTGGGACTTGGAAGCAAGATTCTTGAAAAAGTTCTCAATCGAGACAGGGTCTCACGCGGAGTCCGTGATGCAAATACCGTTCCAGCTTTACTCGTATAGAGTCGCAGAGAAGGAGAAGATGAGTCCAAACTTGAGCAAAGAAGAGAGTTCGGACACAGGAGGTGAGTCTGCTTCAGAAAGCGTTTCGGATCCACCAACAACAACGCCAAAGTATACAAAGTCATTGTGCTTAGATGCAATATCTGAATACTCAGAGTCAGATACAAGATCAATTTACTCAGAAGCTCCTAAGAAAAAGATCTCACCAGCTTCAAAGAAAAGTGGTGACATGGCCAAACTCAGGAGAAACTCAAGCGCAGGTTTACGGTCAACAGGATCATCGCCAGTAAAGCCAAGACCAGCACCGAAAGTGACATTGCCGTTAAGTCCGTTTGGAAGAAACAGCAAAGCCAGGAAGGATACAAGGTTGCCTTTGAGTCCTATGAGTCCTTTGGGACATACAAGACGTAGTAGACATCTCTCTGGCCCTGCTTCAGACTCTGAGCTAACTTAG

>B_rapa_1

ATGGAGCAGTTCAGACAAATCGGCGAAGTTCTTGGAAGTCTAAATGCTCTCATGGTATTACAAGACGATATCTTGATCAACCAAAGACAATGTTGTTTGCTGTTAGAAATCTTCAGCTTAGCTTTCACCACCATTGCCGAAGAAATCAGACAGAATCTAAAACTCGAGGAGAAACACACCAAATGGAAAGCCCTTGAACAGCCGTTAAGAGAGCTTTACAGAGTGTTTAAAGAAGGAGAGTTATACGTCAAGCATTGCATGGACAACAGTGACTGGTGGGGCAAAGTAATCAATCTTCATCAGAACAAAGACTGTGTTGAGTTTCACATACACAACTTGTTCTGCTACTTCCCGGCGGTCATCGAAGCGATCGAGGCTGCGGGAGAGATCTCGGGCCTCGACCCGAGCGAGATGGAGAGAAGGAGAGTTGTGATCTCGAGAAAGTACGATAGAGAGTGGAATGATCCTAAGCTCTTTCAGTGGAGGTTTGGGAAACAGTATCTTGTTCCAAAGGATATCTGCAGCAGGTTCGAGAATTCGTGGAGAGAAGACAGATGGAAACTAGTGGAGGCATTGCAAGAAAAGAGGAAATCAAACAGCAACGAGATCGGAAAGACGGAGAAGCGCTTAGCTGATCTGCTTTTGAAGAAGCTAACCGGTTTGGAGCAGTTTAACGGTAAACTGTTTCCGAGCTGGATACTTGTCGTCTCAAAGGATTACCAGGTGAAAAGGAGGTTAGATGGAGACGGACATTACAAGGAGATTCAATGGTTTGGTGATAGTTTTGTTGTGAGACACTTCTTCAACGATCTTGAGCCGTTGAGTTCTGAGATTTCATCTCTTTTAGCGCTTTGTCATCCGAATATACATCAGTACCTGTGTGGATTCTATGACGAAGAGAGGAAAGAATGTTTTCTGGTTATGGAGTTAATGCACAAAGATTTGCAGAGCTACATGAAGGAGAATTGCGGACCAAGAAGAAGATATCTCTTCTCTGTTTCCGTCGTGGTTGATATCATGATGCAGATTGCGAGAGGAATGGAGTATCTCCATGGGAATGATATCTTCCATGGAGATTTAAACCCGACGAATATTCTATTGAAAGAGAGATGCCACACCGAAGGTTATTTCCACGCGAAAATCTCCTGTTTTGGTTTATCTTCGGTCAAATCTCAGTCTTCTTCATCCTCAAGAGCAAGCACTCATGATCCTGTGATATGGTATGCACCTGAAGTTCTAGCAGAAATGGAGCAAGGAACAAAAACTCTGAAATCGAAGCTGACGCATAAAGCTGATGTGTACAGCTTTGGGATGGTTTGTTTTGAGCTGATAACAGGTAAAGTACCTTTTGAAGATAGTCATCTTCAAGGTGAGCAAATGGCTATCAACATTAGAATGGGAGAGAGACCTCTTTTCCCTTTCCCTTCACCCAAGTACCTCGTTAGTCTGATCAAACGGTGTTGGCACTCAGAACCGAGCCAGCGTCCAAACTTCTCTTCCATCTGCCGGATACTACGCTACATCAAAAAGTTCCTAGTTGTGAATCCGGACCATGGTCACCCTCAGATTCAAATTCCACTAGTTGACTGTTGGGATTTGGAAGCAAGGTACTTGAGAAAGTTCCCAGGTGAGTTAGGGTCCCACATGGCGTCGGTGAGTCAGATCCCTTTCCAGCTATACTCATACCGAGTTTTGGAAAGAGAGAAAATGAATCCAACAAACTCAAAAGAAAGCAGTAGTACAGAGGCAAGCGAATCAGAAGTTGAAGATCCACCTAATGCCGTGATAATAAGAGATACAAAGTCTTTGTGTTTAGATACAATATCTGAATACTCGGATACAAGATCAGTTTACTCAGAAGCTCCCATCAAAAAGGTCTCAGCTTCAAAGAACAGTGACGAATTGGTAAAGCTCAGAAAAAGTCCAAGCTTAGGGTCGTCAACGAAGTTGAGATCAACCGGAACTTCGCCTGTAAAAACAAGATCATCACCAAAAGCATTACCCTTGAGTTCATTTGGAAGAAGCATAAAAACGAGGAAGGATAGTCGATTGCCTTTGAGTCCAATGAGTCCACTAAGTCCAAGGAGACGTAGAGTACAAACTGGTCATGCTTCTGACTCGGAGCTTACTTAG

>B_rapa_2

ATGGAGCAACTTCGAGAGATAGGAGAGGTATTAGGAGGCATAAGAGCCTTAATGGTGTTCAAGGACAACATTCATATCAACCAGCGTCAATGCACTCTATTACTCGACCTCTTCATCGCCACATACGACTCAGTTTCCGAATCCATGAGACTAAACCTACGTTTCGGAGAGAAGAACACGTCCAAATGGAAGATTCTAGAACAGCCCTTGAGGGAGCTTCTATGTGTGGTACGAGAAGGGGAGGCTTACGTTAGGTTCTCTTTAGAGCCTAAACTAGGCTTTTGGGCTAAAGCTGTTTTCTTACAACACAACATAGATTGCATGGAGCTTCACGTTCATAACCTTCTCTCTTCTGTACCCATCATCATCGAGGCTATAGAGATGGCAAGCGAGCTTTCAGGGTGGGACGAACAAGAGATGAACAAGAAGAGGCTGGTGCATTCGAACAAGTACATGAAGCAATGGAACGACTCTCAGATGTTTACATGGAAGTTCGGGAGAGAGTATTTGGTGACCGAGGACTTGTGCAGCCGGTACGAGAGTGCTTGGAGAGAGGACATGTGGCTTCTAACGCAAGAGCTTCAGGAAAAAAGGCGTCCGGGTTCAAGCAAACAAGACAGGAAGATGGCAGAGTTCCTTTTGAAGAATCTAGGAGATGGTAACGAGCTGTTTCCATCTTCTATTCTGGTCAGCTCGAAAGATTACCAAGTGAAGAAGAGATTAGGTAATGGGAGTCAGTACAAGGAGATTACATGGTTAGGTGAAAGCTTTGCGCTTAGGCATTTCTTTGGAGACATCGATGCTCTGCTTCCTCAGGTCACTTCATTGCTATCTCTTTCACACCCAAACATTGTGTATTACCTTTGTGGATTCGCTGATGAGGAGAAGAAAGAGTGTTTCCTTGTTATGGAACTAATGAGTAAAAGCCTTGGGACGCACGTCAAAGAGGTGTGTGGTCCGAGGAGGAAGAACACACTCTCCCTCCCGGTCGCGGTTGATCTGATGCTTCAGGTATCACGTGGTATGGAATATCTTCACTCCAAGGGAGTGTACCACGGAGAGTTGAATCCATCTAACATTCTTGTCAAACCAAGAAGAAGTAACCAAACCGGAGATGGGCATTATCTACACGGCAAGATTTGTGGCTTTGGTTTGACTTCTGTCAAGGGCTTCTCCTTTAAAAGTGCTTCTTCGAATGAGAGTTTTCCATTCATATGGTATTCCCCTGAAGTGCTAGAGGAGCAGGAACAGGGCGGAAGCAGCCTTAAGTACACCGAGAAATCTGATGTGTACAGCTTTGGAATGGTTTGCTTTGAGCTTTTAACAGGTAAAGTACCGTTTGAGGATAGCCACCTTCAAGGAGATAAAATGAGTAGAAACATTAGGGCTGGGGAGAGACCGCTTTTCCCGTTTCATTCACCTAAATTCATAACAAACCTGACCAAAAGATGCTGGCATGCTGATCCGAACCAGCGTCCATCTTTCTCGTCCATAAGCAGAGTTCTGCGGTACATCAAACGGTTTTTAGCTTTGAACCCGGAATGTCAACAAGACACGCTTGTCTCTCCTCCTGTGGATTACTGTGAGATAGAGACTAAGCTGTTGAAAAAGCTCTCATGGGAAAAAACAACAGAGTTAGTTCAAGTTTCACAGGTTCCTTTTCAGATGTTTGCATATAGGGTGGTGGAACAGGCAAAGACTTGTAAGAAAAACAATCTCCGAGAGGCTTCAGAATCGGGTAGTGAATGGGCATCGTGTAGAACAAGTAGAGGACGATCAAGGCATCCCCCACTGAGCCCATGTGGGCAAAGTATGAGAACGCACTCTGAGAGTCAACTGATAGTGATGAGTCCAAAGATACGGCGATCAAGCTCCGGTCATGTCTCTGACTCTGAGCTTTCCTAG

>B_stricta_1

ATGGAGCAATTCAGACAAATCGGCGAAGTTCTTGGAAGTCTAAATGCACTTATGGTATTACAAGATGATATCTTGATCAACCAAAGACAATGTTGTTTGTTGTTAGAAATCTTCAGTTTAGCTTTCAACACCGTAGCAGAAGAGATCAGACAGAATCTGAAGCTTGAAGAGAAGCACACTAAATGGAGAGCTCTTGAACAACCTTTAAGGGAGCTTTATAGAGTGTTTAAAGAAGGTGAATTGTATGTTAAGCATTGTATGGACAATAGTGATTGGTGGGGCAAAGTAATCAATCTTCATCAGAACAAAGATTGTGTTGAGTTTCATATACACAACTTGTTCTGTTATTTCTCTGCGGTTGTTGAAGCGATCGAGGCCGCTGGAGAGATTTCGGGTCTTGACCCTTCTGAGATGGAGAGGAGGAGAGTTGTGTTCTCAAGAAAGTATGATAGAGAGTGGAATGATCCTAAGCTATTTCAATGGAGGTTTGGGAAGCAGTATTTGGTACCGAGGGATATTTGCAGTCGGTTTGAGCATTCGTGGAGAGAAGATAGATGGAATCTAGTGGAGGCATTACAACAGAAGAGGAAATCGGACAGCGATGATATTGGTAAAACCGAGAAGCGTTTAGCTGATTTGCTTTTGAAGAAGTTAACCGGTTTGGAGCAGTTTAACGGGAAACTGTTTCCGAGCTCGATACTTGTTGGTTCAAAGGATTACCAGGTGAAAAGAAGGTTAGATGCAGATGGACAATACAAGGAGATTCAATGGCTAGGTGATAGTTTTGCAGTGAGGCACTTTTTCAGCGATCTTGAGCCGCTGAGTTCTGAAATTTCATCTCTTTTAGCTCTTTGTCATTCGAATATACTTCAGTACCTTTGTGGATTCTATGATGAAGAAAGGAAAGAATGCTTTTTGGTTATGGAGCTAATGCACAAAGATTTGCAGAGTTACATGAAGGAGAACTGCGGACCAAGAAGAAGATATCTGTTTTCGGTTCCAGTCGTGATCGATATCATGCTGCAAATTGCAAGGGGAATGGAATTTCTCCATGGAAATGATATCTTCCATGGAGATTTAAACCCCATGAACATTCTTTTGAAAGAGAGGAGTCATACCGAAGGTTATTTCCATGCGAAAATTTCTGGATTCGGTTTATCTTCCGTCAAAGCTCAGTCTTCTCGGTCTTCATCGAGACCAGGCACTCCTGATCCTGTGATCTGGTATGCACCTGAAGTTCTAGCAGAGATGGAACAAGACCTGAATGGTACAACTCCGAAGTCGAAGTTGACGCATAAAGCTGATGTGTATAGCTTTGCAATGGTGTGTTTTGAGCTCATAACCGGTAAAGTTCCATTTGAAGATAGTCATCTTCAAGGTGAACAAATGGCGATCAACATTAGGATGGGTGAGAGACCTCTTTTCCCTTTCCCTTCACCAAAATATCTCGTTAGTCTCATTAAACGATGTTGGCACTCAGAACCGAGTCAGCGTCCTAACTTCTCTTCGATTTGTCGGATACTACGCTACATCAAGAAGTTCCTAGTTGTGAATCCAGATCATGGTCATCCTCAAATGCAAACTCCACTAGTTGATTGTTGGGATTTAGAAGCAAGGTTCTTGAGAAAGTTCCCAGGTGATGCAGGGTCTCACACCGCATCCGTGAATCAAATCCCTTTCCAACTATACTCATACCGGGTTTCGGAAAGAGAGAAGATGAATCCAAACTCAAACGAAAGCTCAGAGGCAAGTGAATCTGAAAGCGTTTCAGTGGTTGAAGATCCACCTAATGCCATGATTACAAGAGATACAAAATCATTGTGTCTAGATACAATATCTGAATACTCAGATACAAGATCAGTCTACTCAGAAGCTCCCATCAAAAAGGTCTCAGCTTTAAAGAAAAGCGGCGAAATGGCAAAGCTCAGAAAAAGTCCAAGCTTAGGTTCAGAAAAGTTGAGATCTACAGGAACTTCTCCGGTAAAAGCAAGATCATCGCCAAAAGTATCACCGTTGAGTCCATTTGGAAGAAGCATTAAAGCAAGGAAAGATAATCGATTGCCTTTGAGTCCAATGAGTCCTTTAAGCCCAGGATTACGTAGACAACAAAACTAG

>B_stricta_2

ATGGAGCAATTTCGAGAGATTGGAGAGGTATTAGGAAGCATAAGAGCGTTAATGGTCTTCCAGGACAACATTCAAATCAACCAGCGTCAATGCACTCTACTACTCGACCTCTACACCGCGGCCTACGATTCAATTTCCGAATCCATGCGATCTAATCTACGCTTCAAAGAGAAGAACACTAAATGGAAGATTCTAGAACAGCCCTTAAGGGAGCTTCTATGGGTGGTAAGAGAAGGGGAGGCTTATGTTAGAATAGATTGCACAGAGCTTCACATACATAACTTGCTTTCTTGTCTTCCGATCATCGTCGAGGCTATTGAGACAGCGGGTGAAGTTTCAGGGTGGGATGAAGAAGAGATGAGCAAGAAGAAGTTGGTGCATTCTAACAAGTATATGAAGCAATGGAATGACTCTCAGATGTTTACCTGGAAGTTCGGGAGAGAGTATTTGGTGACAGAGGATTTCTGCAACCGGTTCGAGAGTGCTTGGACAGAGGACCGTTGGATTTTAACCAAGGAGCTTCAGGGGAAAAAGCAGCCCGGTTCAAGCAAACATGATCGGAAGATGACTGATTTCCTTTTGCAAAATTTGGGAGATGGAAACAAGAGTCCCAAGCTATTTCCATCTTTTCTTCTGGTTAGCACGAAAGATTACCAAGTGAAAAAAAGATTAGGGAATGGAAGTCTGTACAAGGAGATTACCTGGTTAGGCGAGAGCTTTGCGCTTAGGCATTTCTTTGGGGATATCGATGCTTTGCTTCCTCAAGTTACTCCACTGCTATCTCTTTCACACCCAAACATTGTATATTACCTCTGCGGATTCACAGATGAGGAGAAGAAAGAGTGTTTCTTGGTCACAGAACTGATGAGTAAAACACTAGGTATGCACATAAAAGAGGTATGTGGTCCGAGGAAAAAGAACACTCTCTCCCTCCCGGTAGCGGTTGATCTAATGCTTCAGATAGCTTTGGGTATGGAATATCTTCACTCGAAGAAAATATACCACGGAGAGTTGAATCCATCAAACATTCTTGTCAAACCAAGAAGAAGTAACCAATCCGGAGATGGGTATCTGCACGGGAAGATTTTTGGGTTCGGGTTGAATTCTGTCAAAGGATTCTCTAGTAAAAGTGCTTCTTTGACAGATCAAAATGAGAATTGTCCATTCATATGGTATTCCCCGGAAGTTCTAGAGGAGCAAGAACAGAGCGGAACGGCAGGAAGCCTCAAGTATACTGATAAATCTGATGTGTATAGCTTCGGGATGGTTTGCTTTGAGCTTCTGACAGGTAAGGTGCCATTTGAAGATAGCCACCTGCAAGGAGATAAGATGAATAGAAACATCAGAGCCGGGGAGAGACCGCTTTTTCCATTTCATTCACCTAAATTCATAACAAACCTGACAAAACGATGCTGGCATGCTGATCCTAATCAGCGTCCAATTTTTTCATCGATAAGCAGAATTCTCCGGTACATCAAACGGTTTCTAGCTTTGAACCCGGAATGTCATAGTAGTAGCCAACAAGACCCGCCCATTGCTACTACTGTGGATTACTGCGAGATCGAGACTAAACTATTGCAGAAGCTTTCATGGGAATCGACAGAGTTAACTCAAGTTTCACAAGTTCCTTTTCAAATGTTTGCATATAGGGAACTCCCAGAGGGCGATCACGGCATCCCCCCGCTAAGCCCTTGTGGGCAAAGCATGAGAACAAACTCTGAGAGTCAACTGATGTTGATAAGCCCAAAGATACGACGATCACACTCTGGTCATGTCTCTAACTCTGAGCTTTCCTAG

>B_stricta_3

ATGGAGCAATTTAGGCAAATCGGAGAGGTTCTAGGAAGCTTAAACGCGCTTATGGTATTACAAGACGATATCTTAATCAACCAAAGACAATGTTGTTTGCTGTTAGATATTTTCAGTTTGGGTTTCAACACCGTAGCCGAAGAGATCAGACACAACTTGAAGCTTGAAGAGAAGCATACTAAATGGAGAGCCCTTGAACAACCTTTAAGAGAGCTTTATCGAGTGTTTAAAGAAGGCGAAATGTATGTTCGGAATTGTATGTCCAATAAAGATTGGTGGGGCAAAGTTATCAACTTTCATCAGAACAAAGATTGTGTTGAGTTTCATATACACAACTTGCTTTGTTACTTTCCGGCTGTGATCGAAGCCATCGAGACAGCTGGAGAGATTTCGGGTCTCGACCCTGCTGAGATGGAGAGGAGGAGAGTTGTGTTCTCAAGAAAGTATGATAGAGAGTGGAATGATCCCAAGCTTTTTCAATGGAGGTTTGGGAAACAGTATTTGGTTCCAAGGGATCTTTGCAGCCGCTTTGAGCATTCATGGAGAGAAGATAGATGGAATCTGGTGGAAGCATTACAAGAGAAGAGAAAGTCCAAGAGCGATGAAGTTGGGAAGACTGAGAAGCGTTTAGCTGATTTTCTGTTGAAGAAACTAACTGGATTGGAACAGTTTAACGGTAAGCTCTTCCCGAGTTCGATACTTGTTGGTTCTAAGGATTACCAAGTGAGGCGGAGATTAGGTGGTGGTGGTCAGTACAAGGAGATTCAATGGTTAGGGGATAGTTTCGTACTGAGACATTTTTTCGGGGATCTTGAGCCGTTAAATGCAGAGATTTCTTCTCTGTTATCGCTTTGTCACTCGAATATACTTCAGTACCTTTGTGGATTCTATGATGAAGAAAGGAAAGAATGTTCTCTGGTTATGGAGTTAATGCACAAAGACTTGAAAAGCTACATGAAGGAGAATTGTGGACCTAGAAGAAGATATCTCTTCTCTGTTCCTGTTGTGATCGATATAATGCTTCAAATCGCGAGAGGGATGGAGTATCTTCATTCAAATGAGATTTTCCATGGAGATTTGAATCCCATGAACATTCTTTTGAAAGAGAGAAGTCACACCGAAGGTTATTTCCACGCGAAAATCTCAGGTTTCGGTTTGACATCAGTCAAAAATCAATCTTTTTCTCGGGCTTCTTCTAGACCAACCACACCTAATCCTGTAATCTGGTATGCACCTGAAGTTCTCACAGAGATGGAACAAGATCTCAAAGGTACAGCTCCGAGATCAAAGTTTACACATAAAGCTGATGTCTATAGCTTTGCAATGGTGTGTTTTGAGCTTATAACCGGTAAAGTTCCATTTGAAGATGGTCATCTAGATCAAGGCGATAAAATGGCTAAGAACATTAGAATGGGAGAGAGACCTCTCTTCCCTTTCCCTTCGCCGAAGTACCTCGTCAGTCTAATTAAACGGTGCTGGCATTCAGAACCAAGTCAGCGTCCGACTTTCTCTTCGATTTGCCGGATCCTCCGCTACATCAAGAAGTTCCTAGTTGTGAATCCGGATCACGGTCACCTTCAAATCCAAAATCCACTTGTTGATTGCTGGGACTTAGAAGCAAGATTCTTGAAAAAGTTCTCAATAGAGTCAGGTTCTCACGCGGGATCCGTGACACAAATACCGTTCCAGCTTTACTCATACAGGATTACAGAGAAAGAGAAGATGAGTCCAAACTTGAACAAAGAAGAGAATTCAGAAACAGGTGAATCTGTTTCCGAAAGTGTTTCAGTAGTTGAAGATCCACCAACAATGCCAACGTATACAAAGTCATTGTGCTTAGATGCAATATCTGAATACTCAGATACGAGATCAGTTTACTCAGAAGCTCCAATCAAAAAGACCTCAGCTTTAAAGAAAAGCAGTGACACCATTAAACAAAGAAGAAATTCAATCTCAGGTTTACGATCTTCAGGATCATCGCCTATAAAACCAAGATCAGCACCGAAATTGTCATCACCGCTAAGTCCATTTGGAAGAAGCAGCAGCAAAGCAAGGAAGGATACTATATTGCCTTTGAGTCCTATGAGTCCTTTGACCCGTGGAAGACGTAGACAACTCTCTGGTCCTGCTTCCGACTCTGAGCTTACATAG

>B_vulgaris_1

ATGAAAGCTTTAATGATATTACAAGACAATATTCAGATAAACAAGAGGCAGTGCAGTTTTTTGTTTGACATGTTTAATTTAGCCTTTGAGACATTGTCAGAAGAGATTAAGCAAAACCTTAGATTAGAAGAGAAGAACACAAAATGGAGAGCATTAGATCAACCTTTAAAGGAGCTTCATAGAGTATTTAAAGAAGCAGAATTGTATATGAAACAATGCATGGATTCTAATAAAAATTGGTGGGCGAAAGCGATTTACTTTCATAACAGTCAAGAGTGTATTGAATTTCATATGCATGACTTGCTGTGTTATTTTGCAATCGTTATCGAAGCGATTGAAACAGCTGGGGAGATATCAGGGCTTGACATAGATGAGATGAAGAAGAAGAGAATCATACTCTCTAACAAGTATGATATGACTTGGAATGATCCAAAACTATTCTCTTGGAAATATGGGAAACAATATTTGATTTCCAATGATATTTTAAGTAAAATGCATAATGCTCCGAAAGAGGATAAGTGGCTTCTGATAGGAATGATCAAGGAGAAGAAGGTTTCTGGTTCGTCTATCATGAGTAAACAACTCAGTGATTTACTTGTGAAGAAACTGAATGAGGAAGAATTGCTCTGCCATAGTACTTCGGTTTTAACTGGAGCAAGAGATTATCAAGTGAGAAGGCGGTTAGGTGGGGGAATAATGCAGTGTAAAGAGATACAATGGTTAGGGGAGAGTTTTGTTTTGAGACACTTTTTTGGAGAGTCTTCATTGTTAAATTCGGAAATTTCTTCATTGATGTCTCTTTCACACCCTAATGTAGGTCATTATCTTTGTGGATTTCATGACGAGGAAAAGAAAGAGTATTTCGTTGTGATGGAGTTGATGAGCAAAGATCTTAACTCCTATATTAAAGAGGTTTGTGGTTCAAAAAGGCGGATGCCCTTCTCTTTCCCAGCTGCAGTTGATATCATGCTCCAAATCGCGAGAGGTATGGAATATTTACACTCAAAGAAGATATATCATGGAGATTTGAACCCTTCGAATGTTCTTATCAAGCCTAGAAATTCATCTTCAGAGGGTTACTTTCATGCTAAAATCTCTGGTTTTGGTTTAAAGGGTGTTAAAACTGTGGCTTCTAGGACCTCCTCGCCGAACCAAAATGGAATACAACCATCTTTTATTTGGCATGCCCCTGAAGTCCTGATCGAGCAAGAAAAGGCAGGAAGTAGTAGTGGAAAAGACTTCAAGTACTACTCTGAGAAAGCTGATGTTTATAGCTTTGGGATGCTTTGTTTTGAGGTTTTGACAGGAAAAGTTCCTTTCGAAGATGGGCATCTTCAGGGAGATAAAATGAGTAGAAATATAAAGGCAGGAGAGCGGCCTTTATTTCCATTTGCTTTGCCAAAGTATTTTGCAAATCTGACTAAGAGGTGCTGGCAAACAGACCCAACTCAGAGACCAAATTTCTCATCAATTTGTAGGATACTACGCTATATCAAGAGATTGATTGTGATAAATCCTCATCTAGCTCACTCTGAAATTCCGTCGATAGTTCAAGTAGATTATTGTGATATAGAGGCCGGTTTCATGAAGAAGTTCGTGTCAGAGGCTAATGTTGCAGTAACTCCTGTATCACAGATACCATTTCAGTTGTTTTCTTATAGATTAGCTGAGAAAGAGAAGCTTAATGACGGGATCAGTAAAAAAAAGAGTTGGGAATCACATAGCGAGGTTGTTATGTCATCTCATATGGATGATATGGCTAATGGTATTTTTGAAGGCTCATTTTCACCAGCTCATGACACCAAGTCTATGTGCTTAGAGATTCCTGAGAAGAGACTTTCAGTTTTCAGGAGGATTAATGGCCTGACCACAACCACAAATAAGCAGTTCTTGCAAAGACGCAATACAATGTCCGTGTTTCAGGAAATCCCGGAGAATAATCTTCCGATCCTAAGGAAATCAAAAGGTGTAGCAGCTGGTGACTCGTCTTTGCTAGTGCACACTGCAAAGTCAGGCTGCTCAGAAATTCCTGAGAAGAAATCTTCACCTTTGAAGAAATGCAACACACTTTCTTCTCTCAAAGTTCCTGATGAGACACCACCACTACCTGAATGTAGTGAGATCCCTGCAAAGAAACTTCCAGATAAGAAGGTTTTGTTTCAGAGGAAATCTAATAGTGCCAGAACGAGCAAAGATCCAGGGAAAGCTAAAGGCAGAATGAAAGGACCATCATTGTTTAGTCCGCGAGGTCATAGTACCCGGATGACTGCAGAGAGACCACAACAACAATCCACTCCGCTGAAGCCATTTAGATATCAACAGAAACAAGCTGGTCATTCACCAGATCACTCAAATCATTAG

>B_vulgaris_2

ATGGAGCAATTCCGGCAGGTTGGCGAAGTTCTCGGAGGTTTAAAGGCATTGATGGTATTCCAGCAGAATATTCAGATCAATCAAAGGCAATGCTGTTTTCTCCTTGATATTCTTACCTTGGCTTATGAGACAATTGCTGATGAAATGAGAATCAACTTAAAATTTGAAGAAAAGAACAGAAATTGGAGAGTTCTAGAACAACCCTTGCAGGAGCTCCACAAGATATTCCAGGAAACTGAGACATACGTTCGAAACTGCTTCGAAAACAAGGATTGGTGGGCCAAAGCCATAATTTTGTACCAAAATCGAGATTGTGTCGAATTCCATATTCATAACCTGTTTTCCTGCATTCCGATTGTGATTGAAGCAATTGAAACTGCAGCTCAGTTATCCAGTGGGGATGAGGATCAAATCAATAAAAGGAGACTCGTGTGGTCAAGGAAATACCAGAATGAATGGGTTGACCAGAAACTCTTCCAATGGAAATTCGGGAAACAATATTTGATTACAGCAGAAATATGTAATCGGACTGATGAAATATGGAAAGAAGATCGATGGATTTTACTAGATAAAATCCGGGGAAAAAGAATACCTGGATCAACAAAGTATGATCAACAATTAATGAATCTTCTTTTGAAGAACTTGGATGAGGCAGGGTTCTTGAATGAAAAGCTTTTACCAAGCTCAGTCTTGCAAACTTCAAAAGACTACCAAGTTAGAAAGAGGCTGGAGAATGGAGGGGAATACAAAGAAATCTCCTGGTTAGGCGAAACCTTTGTTCTTCGACACATCAAAGGGGATATTGAGTCTTTGATTCCCGAGATATCTCAACTATTATCCCTTTCCCACCCAAATATACTACAGTTTCTTTGTGGTTTCGTGGATGATGAAAAGAAAGAGTGTTTCTTGGTTACAGAGCAAATGAGCAGGAATCTCGGGATCCATGTTAAGGAATTTTATAACCCGAGGAAGCGACTTCCACTTTCGCTTCCTGTTGCAGTTGATGTAATGCTTCAGATTGCAAGAGGGATGGAGTATCTTCACTCGAAGAAAATATATCATGGGGATTTGAAACCGTCAAACATTCTTACTAAACCAAGAAACAATTCTTCAGAGGGGTTTGTTCATGTTAAGGTTACAAGATTTGGGTTATCAACACTGACGAGCAAGAATCAAAACCTAAGTGAAGAAGTTTCATACATCTGGCATTCCCCAGAAGTATTAACGGAGCAAGAGGAGTTAAAAGGTGTACCAGAGGCAAAGTACACAGAAAAATCTGATGTGTATAGCTTTGGTATGATATGTTTTGAGCTTCTGACAGGAAAAGTCCCTTTTGAAGATGCACATCTTCAAGGAGACAAGATGAGCAGAAACATTAGGGCCGGGGAAAGGCCACTTTTCCCGCATCATTGCCCGAAATATGTAACAAACTTAACTAGAAGATGTTGGCATTCTGACCCGGATCAACGTCCCACTTTCCCGTCCATTTGCCGGATACTCCGGTATATAAAACGGTTTCTTATGATGAACCCAGAGTACAGCCACCCAAATAATCCTGTTCCTCTCGTTGATTACTGTGATATCGAGTCAGGGTTTCTAAAGAGATTTCCTTCTTGGATAAACAGCGAACCATATCCTCCTGTTTCAGAAATCCCATTTCAGATATTTGCATATCGAGTCATGGAAAGGGAAAAATGTATATCATACATCAAAGATACCTCAGAATCTGGCAGTGAAGTAGCTTATACGTCTGGAGATGAGCACATGACGACAGCAAATGAGACTCTTATATCCGTCGTGGAAAAGAAGAATCCAAAACCTGATGAGTATACGAAAAGGAGGCTTTCATTAAATAATAGATCACTTGATGCAAGAGCAAATAAATATCCAGGAACACCAAAAGGACGATCTCCAAGGCCACCTCAACTTCTACGCAGTTCACGTAGCATGCACTTAATACCCGATAACCACGCGCTTCTTGCGAGTCCACCACGGATCCGAAGGAAGTCTGGTCATGCATCAGACTCTGAGCTATCATAA

>C_annuum_1

ATGCCAGCGTTAGAGGAATTCAGGCAAATAGGAGAGATAATTGGGAGTCTTAAAGCTCTAATGGTTTTCCAAGATGATATACAAATAAATCAAAAACAATGTTGTTTGTTGGTGGACATGCTTAAATGTGCCTACAGGACAATTGCTGAAACCATGAAACAGAACTTGAGATTTGAAGAGAGGAATGCGAAATGGAAAATTCTTGAGAACCCCTTAAGGGAGCTTCTCAGGGTGTTCAAAGAGGCAGAGCAGTATATCAAGCAATCCCTGGAAACTAAGGATTTTTGGGCAAAAGCCATAGTTGTTTACAAGAATACAGATTGTGTTGAATTTCACATCCACAATTTGCTCTCTTGTGTGCCAATTGTCATTGAGGCCATAGAAATAGCAGGAGAGATTTCAGGTAGTGACCATGATGAGATACAGAAGAAGAGGTACATTTACGCGATGAAGTATCAGAAAGAGTGTAAAGACCCAAAAATCTTCCAATGGAAATTCGGAGACCAGTACTTGGTTTCCGAGAAGTTCTGCGATAGGGTATGCTTAGTTTGGAATGAAGACAAGTGGACTTTGCAGAATAAAATTCGAGAGAAGAAGAATTCAGGTTCATGTAGTTTGAAAAAGCATGAGCAGCGACTTGCTGATCTCCTCTTGAAGAACTTAAACGAGATGGAGACAGAGAGCGATCATAAACTTTCACCTAGCTCAGTTCTTATAAATTCTAAGGATTACCAGGTAAGAAGGCGATTAGGTAGTGGAAGTCAATACAAGGAGATACAATGGTTGGGGGAGACCTTCTGTTTACGGCATTTGTTTGGGGATATAAAGCCCCTGATTGCAGATATTTCTCGAGAACTGCATCTCTCTCATCCAAATATAATGCACATCTCCTGTGGTTTTACTGACGAAGAAAAGAGAGAATGTTTCTTAATCATGGAACTTATGAACAAAGATTTATCGAGTTACATCAAAGAGATCTGTGGACCAAGAAAGCGCGTACCATTTTCTCTTCCTGTTGCAATTGATTTGCTCCTTCAGATTGCAAGAGGCATGGAATACCTCCACTCAAAGAAAATCTACCATGGTGAGTTGAGTCCTTCCAACATCCTTGTAAAGACAAGGAATGTATCTACAGAAGGGCATTTACATGCAAAGGTTTATGGATATGGCTCATCACGTTCTATCAACGTTCCTCAGAAAGCTAATGTGAATCAAAATAATGGGACGCTCCCAATCATTTGGTTTGCCCCAGAAGTCCTAGCTGAACAAGAGCACTCGGGGAATGGAGGGAACTTTCAGTATACTGAGAAATCTGATGTGTACAGTTTTGGGATGATATGTTTTGAGGTTTTAACAGGGAAAGTTCCATTTGAAGATAGTCATCTACAAGGAGACAAAATGAGTAGAAATGTAAGGGCAGGAGAGAGGCCATTATTCCCATTTCACTCACCAAAATATTTGACTAGCTTGACAAAGAAGTGTTGGCATACTGATCCATATCAACGACCAAGTTTCTCATCCATCTGTAGGGTTCTCCGCTATGTAAAGAGATTCTTGGTGATGAATCCGGAACACAGTCAACAAGACTCACTATTGCCACCAGTAGACTATGGTGAAATTGAAGCTGCGATCTTAAGGAGCTTCCCCTTTATGGGAAATTCTGAATCTGATCCTCTTCCTGTCACACAAATACCTTTCCAAATGTTTGCATACAGGGTCACCGAGAAAGAAAAGTCGAGCACAATTCACAAAGACATAAATTCTGAATCAGGAAGTGATGGAAATTCAGGTTGTGGGGATGATCCTGTAACGGCAGAAGATTCACTTCCATCACCAACCGAAAAGAAGATTGCATCCCCCGAGATTTTGACCAAGAGACTTTCTGTTAAGAAACCTACCGATATCAAAGTCACCAAGCAACCAGGTAGTGTTTCTTTGCTCAAAAGCCTTATCATGCAGCATTTTCACCTAATCTTCCTTTGCCTTCCTAAAGGCTACTTTACCTTATTCCAAAATTTTAAAGAAAGTCGACTGCCTAACAGAACTTCTGGGACACCAGAAACAACATATACTCCAAAATTTTATTGTTATTTGTTCCTTACTAACAACAAATTCCAAAAGTAA

>C_annuum_2

ATGGATCAATTCAGGCAGATTGGTGAGGTTGTTGGGAGCTTAAAGGCTCTAATGATACTGAAACATGATATTCAAATAAACCAGAAACAATGTTGTCTTCTGTTTGACATGTATGTACAAGCTTTCGACACGATTTCCGAGGGGATCAAGCATAATTTAAGGTTAAATGAGAGGTATTCAAAGTGGAAAGCACTTGAACATCCAATGAAAGAGCTTCATAGGATCTTTAAAGAAGGTGAAATGTACATAAAAAGTTGTTTGGATGTGAAGGATTTGTGGGGCAAAGCTATAAGTTTACATCTCAACCAGGATTGTGTTGAGTTTCATGTTCACAATTTGCTTTGTTGCTTCCCTGTTGTGATTGAGGCTATTGAAACAGCTGCAGAAATCTCAGGATTCGACGAGGAACTTATGCAGAAGAGGAGGACTGCGCTCGCTAAAAAGTACGAGGGAGAGTCAATGTGTGATCCAATATTCTTCCAGTGGATGTTTGGGAAACAGTACTTAGTAACTAGAGAGATATGTAGTGAATTGGAGAGTTGTTGCAAAGAAGATAGATGGTATCTTGTTGAAATGATAAGGCAAAAGATGAATGCTGCAGAAAATTTGGCGATAAACGAGCATGGTATAGCTGAGATATTGCTCAAGAAACTAGATGGATCAGAAAAACTCAAACAAATAATGCTCTTGCCTAGTTCAATCTTGGTTGGAGCAAGTGATTATCATGTGAAGCGGCGTTTAAGCTCGCGAGGAGGACATGTTAAGGAGATTCAATGGTTAGGAGAAACGTTCGCGCTGAAGAACTTTTTTGGTGAACTGAATGAATCAGTAGTTGCTCAACTTTCTCTAGTGTTTTCACTTTCACATCCCAACATTCTGCAGTATCATTGTGGTTTTTACGACGAAGTAAAGAAAGAAGGGTACCTTGTTATGGAGCTGATGAACAAAAGTTTGTCGGCTTCCATAAAAGAGCATTCCGGTCAGAGGAAGAAAGGGCCGTTTACAGTCCAAGTTGCTGTCGATATAATGCTTCAGATTGCTCGAGGGATGGAGTACCTGCACTCGAGAAAGATCTATCACGGAGAGTTGAATCCATCTAATGTGCTCCTTAAACCAAGAAACTCCTCCGCGGAGAATTATTTTCATGCGAAAGTAAGAGGCTTTGGTTTAACATCTATGACGAGTAATCGCAAGGTTGATGATGACAATGCTGCAGAATCATTCATATGGTATGCACCAGAAGTACTGACTGAAAAGGAAAAACAGGAAAGCAAATGCACTCACAAGTATACGGAGAAAGCTGATGTTTACAGTTTCGGAATGATATGCTTTCAACTTTTAACGGGGAAAGCTCCTTTCGATGAACAGCTTCAAGGGGAGAAAATGGCGCGGAATATTAGAACAGGAGAGAGACCGCTCTTCCCGTATCCTTCACCTAAATATCTCATAAACTTGACAAAAAAATGCTGGCAAACGAATCCAGAACTTCGCCCGAGTTTCTCTTCTTTGTGTAGGATACTGCATTATATCAAGAAAGTACTCGTCATAAATCCAGGACATGGCCAGCCGGAATGTCCTCCTCCACTCGTCGACTATTGTGAAATCGAGGCTGGCTACTCAAAGAAGTTCCTGGGAGAGGAAAGCGCTGGTTTGGCCCCGGTATCACAAATTCCTTTCCAGATGTTTGCTTATAGACTTGTTGAGAAAGAGAGAATTTCGGGCAACTCTAAAGACGAGCGCTGGGATTCATCACATTATGGTTTTTCGAGCCACAGGACAGCATCAATGCGGAGCGATGATGAGCATATGGCTGCAATAGATGATCTCTTTTTCGCTCCAACTGATCGAAGGTCAGTTTGTTCAGAGATAATAGAGAGTAAAGAACCAAGATTCTGGGATCAGAGGTCAGTCATCTCCGAGACACCTCTTCGAAAAGTTTCATCATTTGATCAAATGTCAGTTAGTTCTGAGAGTTCTAAAGAGAAATTTCCACCAGCAGCATCAGCAAATGAAAAGACAATTTATACTGATTTTCCGGAAAGGAAAGCAATACCAACAGCATCAGGCGATCAGTTCTCACAGCGTGTTGATGCACCAGGGAAGACGATTTCATCAAAGAGAGTGAATCAAAAGCTGAATCTTTCGGAAACCCAAGAGAATGCAATGTCTCAGAAAACGGATGACAGAAAACCTTCATCGACTGAGCAGAAATCGGTGCCTCCTGTGATTTCAGAGAAGAAAAATTCATCAGGCGATCAGATATCAACGTATTTCAACATTCAAGAAAAGAAACATGCACCAGATGATCAGAAACCATCAAGTTCTGCGACTCAAGAGGAGGAAAACTCATCTGATAACCAAAAACAAAAAGATTCTGGAAGTCCAGAAAAGAAGCAGTCATCAACAGTTGCCAATGTCAAGTCCGTTCGTGGTGATTCTACGCAAAGGAAAGCTATGTCAAGAAGAACACAATCCAAGAATCCAGAGAAGAAAGTTCTATCAAGTCCGGAAGCCAATCAAAACTCAATGTCATCTGAGATAACAGCATCAAAACCAAGCAGTTCAAAGTTATCAGTAAAGAAAACGCCATTTCGCAAAAAAATACCGGACAAAGAGAGTAGCACTACACCAGAGAAACTGACAGACTCGTCGCCAAGATCTTCACCAGCTAGAGTTAAGAGAATACAGTCATCAACTATGTCTTCTCCAACACAAAGCCTAAAAGCTTCATCTCCAAGATCACATGCAGTCACTACATATCGACATGGATATCTATCGCCAACTGCATCACCATTAAATCCGTGCCCTCGCTATGCCAGAACAAACCGAGAACTCCATTTGTCCTCAGCTATGAGCCCACATAGACCATGGAAACCTAATAGCAATAGCAATAGCTAA

>C_annuum_3

ATGCTGGTAGAAGGTAGCAGGGGAGCATTGGAGGAACTGGTAAAGTTGCTGCCATGTGACCAGGAGGTCACGGGTTCAAGCTTTGGAAATAGCCTCTTGCAGAAATGTAAGGAGGTCACGGGTTCAAGCCTTGGAAACAACCTCTTGCAGAAATGCAAGGTAAGGCTGCGTTCAATACACACTTGTGGTGGGGCCCTAAACCCGGACTCTGCGCATAGCGGGAGCTTTACTGCACCGGACTGCCCTTTTAGTACTGGTAGAAGTTGTTGCCATGTGACCAAGAGGTCACGGGTTCAAGCCTTGGAAACAGCCTCTGGCAGAAATGCAAGGTTAAATGAGATGAATATAAAGTGGAAAGCTCTTGAATTGCCAATGAAAGAGCTTCATAGGATCTTTAAAGAAGGCGAATTGTACATAAAATATTGTCTTGATGTTAAAGATTTGTGGGGGAAAGCTATAAGTCTTCATATGAACAAAGATTGTGTTGAGTTTCATGTTCATAACTTGCTGTGTTGTTTTCCTGTTGTTATCGAGGCTATTGAGACTGCTGCAGAAATTTCCGGGTTCGATGAGGAGGAAATGCAGTATAGGAGAAGCGCGTTGATGAGGAAGTACGATAGGGAGTTGATTGATCCGAAGTATTTTCAGTGGATGTATGGGAAGCAGTATTTGGTTACACGCGAAATGTGTTGCCGGTTGGAGAGTTGTTGGAAAGAGGATAAATGGTTGCTCGTCGAGGCGATAAGGAGAAAGAAAACTGAAACTTTGTTGAAGTATGAACAAAGGCTTGGTGAGTTCTTGTTAAGGAAGTTAGACGGTGTTGAATTGGTGAACAAGTTATTGCTGTTGCCTAGTTCGATCTTAGTTGGATCGAGTGATTATCATGTGAAGAGACGGTTAGGTTCGTGGGGTGGACATGTTAAGGAGATACAATGGCTAGGCGAAAATTTTGCTTTAAGGAACTTTTTCGGAGAGGTTGAGCCATTACGCGATGAGATATCACTGGTGTTTTCGTTATCTCATCCAAACATTCTGCAATATCACTGCGGTTTTTATGACGAAGAGAGGAAAGAAGGGTACCTTGTTATGGAGTTGATGAATGATACTCTTGCGACGTATATAAAAGATCATTCCGGTCAGAGGAAGAGACCGCCTTTTACAACGTCTGCTGCTGTTGATGTCATGCTTCAGATCGCGAAAGGGATGGAGTATCTGCATTCGAGGAAGATGTATCACGGTGAGTTGAATCCGTCTCATGTGCTTCTTAAATCGAGGAGTTACTCATCGGAGAGCTATTTTCAGATGAAACTCAAAGGGTTCGGGTTGAGTTCGATCAGAAGTACTTACAAGACTGCTAGTCAGAAAACAGCTGACTCAATCATCTGGTACGCCCCGGAGGTTCTTGCTGAACAGGAAAAGCCAGGAAGCAAATGTGCCTTCAAGTACACGGAGAAAGCCGATGTGTATAGCTTCGGAATGATCTGTTTCCAGATTCTAACGGGGAAAGTTCCGTTTCATGAAGGACATTTGCAAGGGGAGAAAGTTGCGCGGAACATCAGAGCCGGGGAGAGGCCTCTCTTTCCTTATCCTTCGCCTAAATACCTTGTGAACTTGACAAGAAGATGTTGGCATACCAATCCAAATTTTCGGCCGCGCTTCTCCTCTATTTGCAGGATTTTACGTTACATCAAGAAGGTACTTGTTATAAATCCCGAACATGGTCAGCCTGAAACTGCTCCGCCACTTGTTGACTATTGCGATATTGAGTCTGGTTACTCGAAAAAATTTGTTGAAGAGGAGAGGAGCACGAGTTTAGCCCCGGTATCACAGGTTCCTTTTCAGATGTTTGCTTATCGGCTCATTGAGAAAGAGAAGATTTTCGGAAAGAAATGGGATCGAGGATCAATGCAGAGTGAATACGGTCAATTGTCTGCGATGGATGATCTCTTTCTTGCCCCGAGTGATGGAAGGTCAGTTTGCTCCGAGATAATCGACAGGAAAGATACAAGATTGTTTGATCAACGGTCAGCGATCTCAGAGATACCACATAGAAAGTTATTGTTATTCGATCAAATGTCAGTTGGTTCTGAGAGTCCGGAGACAAGATTCTCGTCTGTTGCAGCAGCAGCAGATGAATCGTTTTTCTTTGCTGATAGTCCGGACAGGAAAGCAGTATCGACACCAATAGTTAATAGGTCACCACGCGTCGTCACACTGGAGAAAAAGATTGTATCTTCGATGAGCAAGAACCAAAGACGTAGATTTTTGGATAACCAGGAAAATGCAATGTCTCCGAAAGCAGATGAAGAAAGACCTTCATTAAACGTAGCTGCTTCGCAAACTTTAGTATCTGCACAGACATCGGAGAACACGAAACCAGCTGAGCAAACTTTAGTATCTCCACCGATTTCAGAGAAGATGAAACCAGCCGAGCAGAATTTAGCATCTCCGCGGACTTCAGAGATGGAGAAATCACCCGGGCAAAAGGTAGCATCTCCTCCGATGTCAGAGAAGAAAATTTCATCTTATGATAAGAAATTGACTAGTCCTGAGATTCAAGAAAAGAAACATGAACCAGATGATCGAAAACTAATCAATTCCGATGCTCCTGTGAAGAAACATTCACCTGATGATCAGAAATTACGACGCTCCAAGAATCCAGATAAGGCGAAGTCATTAGCTGCCAATGACAAACCACTTCATGGCGACCGCGTGCAAAGGAAAAGTTCGAGGACCAGGAGACTAAACCAAAAGTTAACCAAGACTGTTGAGAAGAAGGTTCCTGAAGCTGAATCCGTGAATCCCGAGACATCTGCATCCAAGCCAAGCAATCAAAAGATAGCAGTCAAGAAAGATTCATCGCGCAACAAAGTGCACGACACAAAAACTAGCACCGTACCAGAAAAACTGATTGACGACTCACCAAGATCTCCACCAGCTAGAGTAAACAGAATACATTCATCGCCAATGTTTTCTCCAACGCGAACTCCAAAGACACATTCAAGATCACCCGCAATTGCCTCAAAGGGATACGGATATCATTCCCCAAGTTCATCACCTTTGAATCCATGTTCGCGATGTGCAAGATTAAACCGAGAGTGCCAACCTTCAGGGATGAGCCCACATATACACAGGAAAGCTCATACCTCTCATTCCGAGATAGCTTAG

>C_clementina_1

ATGGAGCAATTCAGGCAGGTGGGCGAGGTTCTGGGAAGTTTAAAGGCTCTCATGGTGTTGAAAGATGATATTCAAATCAATCAGCGACAGTGTTGTTTGCTTCTTGACATCTTTACTTTAGCCTTTTCCACAGTTGCTGAGGAAATCAGGCTTTTCTTAAAATCGGAAGAGAAGAACACAAAGTGGAAAGCTCTTGAACAACCTTTGAGAGAGCTTCAGAAGGTCTTTAGAGAAGGAGAGCATTATGTTAAGCAATGCTTGGATTATAAAGATTGGTGGGGTAAAGCAATTAGTCTCCATCAAAACAAAGATTGTGTTGAATTTCATATTCACAATTTGCTTTGCTACTTCCCCGCTGTGATTGAGGCCATTGAGGCTGCTGGTGAAATCTCAGGGCTTGACCCAGATGAGATGCAAAGAAGGAGAGTAGCATTTGCAAGAAAATATGACAGGGAATGGAATGACCCTAAACTTTTCCAATTGAGATTTGGGAAGGAGTATTTGATCCCTCGAGAAGTCTGTAATGAGTTTGAGAGTGCTTATAAGGAAGATAAATGGCTTCTTATTGACGCACTGAAGGAGAAGAAAAGATTAGGATCTGTTGTTTTGACAAAGAACGAGCAGCGGCTTGTAGACATGCTGCTCAAGAAACTCAATGGTACCGATCCAAGTAACGTTAAACTTTTTCCTAGTTCAATTTTGTTAGGAGGTAAGGATTACCAGGTGAGAAGAAGATTAGGTGCAAGCAGCCAGTTTAAGGAAATTCAATGGTTAGGAGATAGCTTTGTCCTGAGGCATTTCTACGGGGAACTGGAATCGCTAAATGCCGAGATTTCGACTATGCTCTCCCTTTCACACCCCAACATAGTGCAATACCTATGTGGATTTTGTGATGAGGAGAAAAAGGAATTTTTTCTCGTAATGGAGTTAATGAGCAAGGATTTGTCCTGCTACATGAGGGAGACATTTGGTTCAAGAAGGCGGAATTCATTCTCTCTACCTGTTGTTGTTGATATCATGCTTCAGATTGCAAGAGGTATGGAATTTCTCCATGCACAAAAGATCTATCACGGAGAATTGAACCCTTCAAATATTTACCTCAAGGGCAGGAGTATGGAAGGCTATTTTCATGTAAAAGTTTCTGGGTTTGGTTTATCCACTGCCAGAACTTACGCATCTCGAAACACACCACCAGCATCACCACAAAACCAGACTGCGCCAAACCCTTATATATGGTATGCCCCGGAAGTTCTGGCGGAGCAAGAAGGGACTGGAAGTACTTCTACTTCCAAGTGCTCAGAGAAGGCAGATGTCTACAGCTTTGGAATGCTTTGCTTTGAGCTTTTAACTGGGAAAGTTCCTTTTGAAGATGGGCATCTTCAAGGAGACAAGATGACCAAAAATATAAGAGCAGGGGAGAGGCCTCTCTTCCCATCAGGTTCTCCGAAATACCTCGTGAACTTAACCAAAAAATGTTGGCACACAAATCCATCTCAGCGTCCAAGTTTCTCGTCCATTTGTCGAATTTTACGTTACATCAAGAAGTTCATGGCAAATAACCCTGACATCGCACGGTCCGAGTTTCAATCACCGCTAGCAGATTACTGCGACATAGAGGCAGGATTTGTAAGGAAATTTGTAGGAGAGGGCTGCCCTGATGTAGCGCCAGTATCACAAATTCCATTTCAAATGTTTGCATATAAAATAGCTGAAAAAGAGAAGGCTAATTCGAGTAATAAAGGTAAGCATTGGGAGTTAGGTAGTGATGTAGCCTCAACTTGTAAGGATGAGGTTTTTTCCGCAGGAGATGAGGGAATGGTACCAGTGACCGAGACAAGATCAGTTTGCTCTGACGTGAAATCTGTTACTTTTGATATGAGATCAGTTTATTCTGAGGCTTTTGGCAAGAAAGTTTCAAATTTTGATACAAGGTCAGTAAGATCCGAGTCCCCGAACAAGAAAATGCAGGGTAGTGTTTCAAGGTTGATTCAATCCATGGCTCTGGAGAAGAATGTTTCCAGTTCTGATACTAAACCAGCGGTTCCGGAGAACAGTATTTCTAATTCAGAAGTTAAATCCGAGGTTCCAGAGAAGAATATTTCTAATTCTGATACTAAACTTGTGGTTTCAGAGAATAACATCTACAATACTGACACTAAATCTGCCTGTTCTGACAATATGCCGGAGAAAAATATCTGTAATGTTGATACAAGATCAGTTTGTTCCAAGAATTCGGGCAAGATCAAGCAATCCACTGGTGTCAAGGTCAAAAAAGGTCCAGGGACAACAAAAATGAGATCAGGAAGAGCAGCACCAAAAGCACGATCACCAAGAACATCAACATCTCAATCACTGCGATCATCAACTTCTCAATCACCAAGGCCACCTCCCAAATCACCACCGTTTATTCCGTCTGGACGCTGTTTGAAGAAGGGCCAAGACCTTAGAAGCCCAAGCAGAAGAAGATCATCAGTCAATGTCTCAAATTCTGAGGTAGCATAG

>C_clementina_2

ATGGAGCAATTTGGACGAATTGGAGAGGTCATGGGAAGTATGAAGGCGCTAATGGTATTCAGAGACAGTATACAAATAAATCAAAGGCAATGCTGTTTGTTGCTAGACATTTTCAGTTTTGCGTATGATTCAATAGCGGAAGAGATGAGACAAAACCTGAGATTCGAAGAGAGACGCGCGAAATGGAAAATTCTTGAACAACCATTGAAAGAGCTCTTCAGGATATTTAAAGAAGGAGAAAATTACATTAAACAGTGCTTGGAAATTAGGGACTGGTGGGCTAAAGCCATTACCCTTTATCAGAACACGGACTGTGTTGAGTTTTACATCCATAATTTACTTAGTTGCATTCCGATCTTGATCGAAGCAATAGAGACGGCAGCAGAATTCTCTGGATGGGATCCTGATCAGATGCACAAGAAGAAACTTGTACACTCCGGCAAGTATAAAAAAGAATGGAAGGACCACAAACTTTTCCAATGGAGATACGGGAAGCAGTATCTTATTACCCCAGATTTTTGCTATAGGATAGACACCGTATGGAAAGAAGATAGGTGGATCCTTTTCAACAAAATCCAGCAAAAGAAAATATCAGGTTCAACTAAGCAAGAGCAGGGACTCATAGATGTCCTTTTCAAAAACTTGGATGGTTCAGGATCTTTGAGTGGGAAACTCTTACCAAGTAGAATCCTAATTAAGTCTGAGGATTACCAGGTGCGGCGAAGATTAGGGAGTGGGAGTCAATACAAGGAGATCCTGTGGTTGGGTGAAAGTTTTGCTTTGAGACACTTTTTTGGGGACATTGAACCCTTAGTTCCTGAGATTTCTTCTCTATTATCTCTTTCCCACCCGAATATCATGCATTTCCTGTGCGGGTTTACCGATGAGGAAAAGAAAGAGTGTTTTCTGATTATGGAACTAATGAGTAGAGACCTTTGCAGCTACATCAAAGAGATTTGTTGCCCAAGAAAGAGAATTCCATTTTCTCTTCCGGTTGCTGTTGATCTTATGCTTCAAATTGCTAGAGGAATGGAGTATCTCCACTCAAAGAAAATCTATCATGGTAATCTAAATCCTTCTAACATCCTCCTTAAACCCAGAGGTGCCTCAACAGAAGGGTACCTGCATGCAAAGATTTCAGGATTTGGCCTGTCTTCGGTGAAAAACTTTGGCCCAAAAAGTCCATCTCAGAGTGGGACTACCCATCCATTTATATGGCATGCTCCAGAAGTTCTAGAAGAGAATGAGCAGACAGAAAGTGCATCAAATTCAAAGTACAGCGAAAAGTCTGATGTTTACAGTTTTGGAATGATCTGCTTTGAGATTCTCACGGGGAAAGTTCCATTTGAAGATGCTCACCTTCAAGGAGACAAAATGAGTCGAAACATTAGGGCAGGAGAGAGGCCATTGTTCCCATTTCATTCACCAAAGTATGTGACAAATTTGACAAAGAGATGTTGGCACGCTGACCCAAATCAACGGCCAAGCTTCTCATCCATCTGTAGAATTCTTCGCTATATAAAACGGTTCATTATGATGAACCCTCATTATAATAGCCAGCCAGACCCGCCAATGCCTTTGGTAGATTATAGCGACATTGAGTCAAGGCTTTTGAGAAAATTCCCTTCTTGGGAGACTCACAATGTTTTGCCAATATCAGAGATCCCCTTTCAAATGTTTGTTTATAGAGTTGTGGAGAAAGAGAAAATAAGTTCAAGCCCTAAAGACACATCAGACTCAGGAAGTGATAAAGCTTCTGTAAGTGGGGACGAAAATATGACCACACCAGAGGATCCATCTCCACCTGTGACTGAAAGAAAGTTTTTACCTTCACCCGAAGCTGTGAACAAGAAACTCTCCGGTGTCAAGAAACTGTCAGAATCGAAAGTTATTAAACAAACAGGGACACCAAAAGGGCGAGCTGTAAGACCTCCGCAACTAATTCCTTGCGGTCGTAGTTTAAGAATGAGCTCAGAAGGGCAGCTAATGGCGATGAGTCCAAGAATACAGAGAACATCCTCTGGCCATGTGTCGGATTCTGAGCTCTCCTAG

>C_grandiflora_1

ATGGAGCAATTCCGAGAGATAGGAGAGGTGTTAGGAAGCATAAGAGCGTTAATGGTGTTCAAGGACGACATTCAAATCAACCAGCGTCAATGCGCTCTACTACTCGACCTCTTCATCGCGGCCTATGATTCGATTTCCGGATCCATGCGATCTAATCTACGCTTCAAAGAGAAGAACACCAAATGGAAGATTCTAGAACAGCCCTTTAGGGAGCTTCTATGGGTGGTAAGAGAAGGGGAGGCTTATGTTAGAATGTCTTTAGAGCCTAAACTTGGGTTTTGGGCTAAGGCTATCCTGTTGCATTCCAATAGAGATTGCAACGAGCTTCACATACATAACTTACTCTCTTGTCTACCGATCATTGTCGAGGCTATCGAGACAGCGGGTGAGGTTTCAGGCTGGGATGAAGAAGAAATGAATAAGAAGAAGTTGGTACATTCTAACAAGTATATGAAGCAATGGAATGACTCTCAGATGTTTACCTGGAAGTTCGGGAGAGAGTATTTGGTGACAGAGGATTTCTGCAACAGGTTCCAGAGTGCTTGGACAGAGGACCGTTGGATTTTAATCAAGGAGCTTCAGGAGAAAAGGCAGCCCGGTTCAAGCAAACATGATCGAAAAATGGCTGATTTCCTTTTCAAGAAACTGGGAGATGGAAACGAGTGTCCCAAGCTATTTCCGTCTTCTCTTCTGGTTAGCGCGAAAGATTACCAGGTGAAGAAAAGATTAGGGAATGGAAGTCAGTACAAGGAGATTACATGGTTAGGCGAGAGCTTTGCGCTTAGGCATTTCTTTGGGGATATCGATGCTTTGCTTCCTCAAATTACTCCACTGCTATCTCTCTCCCACCCAAACATTGTATATTACCTCTGCGGATTCACAGATGAGGAGAAGAAAGAATGTTTCTTGGTTACAGAACTGATGAGTAAAACACTAGGTATGCACATCAAAGAGGTATGTGGTCCAAGGAAAAAGAACACTCTCTCCCTCCCCGTAGCGGTTGATCTGATGCTTCAGATAGCTTTGGGTATGGAATATCTTCACTCGAAGAAAATATACCATGGAGAGTTGAATCCATCAAACATCCTTGTCAAATCAAGAAAACTTAACCAATCTGGAGATGGGTATCTGCACGGGAAGATTTTTGGGTTCGGGCTGAATTCTGTCAAAGGGTTCCCTGGTAAAAGTGCTTCTTTCACGGGTCAAAATGAGAATTTTCCATTCATATGGTATTCCCCAGAAGTTCTAGAGGAGCAAGAACAGAGCGGAACTGTAGGAAGCTTCAAGTATACTGATAAATCTGATGTATATAGCTTCGGTATGGTTTGCTTTGAGCTTTTGACTGGTAAAGTGCCATTCGAAGATAGCCACCTGCAAGGAGATAAGATGAGTAGAAACATTAGGGCAGGGGAGAGACCGCTTTTTCCATTTCATTCACCTAGATTGATAACAAACCTGACGAAACGATGTTGGCATGCTGATCCGAATCAGCGTCCCACTTTCTCATCGATAAGCAGAATTCTCCGGTACATCAAACGTTTTCTAGCTTTGAACCAAGAATGCCATAGTAGTAGCCAACAAGACCCGCCCATTGCTCCCCCTGTGGATTACTGTGAGATCGAGACTAAACTATTGCAGAAGCTTTCATGGGAATCGACAGAGTTAACTCAAGTTTCCCAAGTTCCTTTTCAAATGTTTGCATATAGGGTTGTGGAACGAGCAAAAACTTGTAAGAAACATAATCTCGAGACATCTGAATCGGGTAGTGAATGGGCTTCATGTAGCGAGGATGAGGGTGGAACAGGATCAGATGAGCAAGTGTCATTTGAGAAGGAAAGAAGACTACCATGCTCAATTGATGTTGGTATGAGCAAGAAGCATGCTTCAAAACTTATGAAAAGAGCTTCTGGCCTGAAACCAATTCAAAAAACAGGAACTCCCAGAGGACGATCACGACATCCACCACTAAGCCCTTGTGGGCAAAGCATGAGAACGAACTCTGAGAGCCAACTAATATTGATAAGCCCAAAGATACGAAGATCAAACTCTGGTCATGTCTCTGACTCTGAGCTTTCCTAG

>C_grandiflora_2

ATGGAGCAATTCAGACAAATCGGTGAAGTTCTTGGAAGTCTTAATGCACTTATGGTATTACGAGATGATATCTTGATCAACCAAAGACAGTGTTGTCTCTTGTTAGAAATCTTCAGTTTAGCTTTCAACACCCTAGCAGAAGAGATCAGACAGAATCTCAAGCTTGAGGAGAAGCAAACTAAATGGAGAGCTCTTGAGCAACCTCTGAGAGAGCTTTACAGAGTTTTTAAAGAAGGTGAGTCTTATGTTAAGCAGTGTATGGACAGTAGTGACTGGTGGGGTAAAGTTATCAATCTTCATCAGAACAAAGATTGTGTTGAGTTTCATATACACAACTTGTTCTGTTATTTCTCTGCTGTTGTTGAAGCCATCGAAGATGCTGGGGAGATTTCGGGTCTTGACCCTTCTGAGATGGAGAGGAGAAGAGTTGTGTTTTCAAGAAAGTATGATAGAGAGTGGAATGATCCTAAGCTGTTTCAATGGAGGTTTGGGAAACAGTATTTGGTGTCTAGAGATATTTGCAGCCGGTTTGAGCATTCTTGGAGAGAAGATAGATGGAATCTTGTTGAGGCATTACAAGAGAAGAGGAAATCGGACAGCGATGATATTGGTAAAACCGAGAAGCGTTTAGCTGATTTGCTTTTGAAGAAGTTAACCGGTTTGGAGCAGTTTAACGGGAAACTCTTTCCGAGCTCTATACTTCTTGGTTCAAAGGATTACCAAGTGAAAAGAAGGTTAGATGCAGATGGACAATACAAGGAGATACAATGGCTAGGTGATAGTTTTACAGTGAGGCACTTCTTCAGCGATCTTGAGCCGCTAAGTTCTGAAATTTCATCTCTTTTAGCGCTTTGTCATTCTAATATACTTCAGTACCTCTGTGGATTCTATGATGAAGAAAGGAAAGAATGTTTTTTGGTTATGGAGTTGATGCACAAAGATCTGCAGAGCTACATGAAAGAGAACTGCGGACCAAGACGAAGATATCTGTTTTCGGTTCCAGTCGTGATTGATATCATGCTGCAAATTGCAAGAGGAATGGAGTATCTCCATGGGAATGATATCTTCCATGGAGATTTAAACCCCTTGAACATTCTTTTGAAAGAGAGGAGTCATACTGAAGGTTATTTCCATGCGAAAATTTCTGGATTCGGTTTATCTTCTGTCAAAGCTCAGTCTTCTCGGTCTTCCTCGAGACCAGGCACTCCTGATCCTGTGATTTGGTATGCACCTGAAGTTCTAGCAGAGATGGAACAAGACCTTAATGGTACAACTCCCAAGTCAAAGTTGACTCATAAAGCTGATGTGTATAGTTTTGCAATGGTGTGTTTTGAGCTCATAACAGGTAAAGTTCCATTTGAAGATAGTCATCTTCAAGGTGAACAAATGGCAATCAACATTAGGATGGGAGAGAGACCTCTTTTCCCTTTCCCTTCACCAAAATACCTCGTTAGTCTGATCAAACGTTGCTGGCACTCAGAACCGAGTCAGCGTCCTAACTTCTCTTCGATTTGCCGGATACTACGCTACATCAAGAAGTTCCTAGTTGTGAATCCAGATCATGGTCATCCACAGATGCAAACTCCACTAGTTGATTGTTGGGATTTAGAAGCAAGGTTCTTGAGAAAGTTTCCAAGTGATGCAGGGTCTCACACCGCATCCGTGAACCAAATCCCTTTCCAACTCTTCTCATACCGGGTTTCGGAAAGAGAGAAAATGAATCCAAACTCAAAAGAAAGCTCAGAAGGAAGTAGTGAATCTGAAAGCGTTTCAGTGGTTGAAGATCCACCTAATGCCATGAGTACAAGAGATACAAAATCATTGTGTCAAGATACAATATCTGAATACTCAGATACAAGATCA

>C_grandiflora_3

ATGGAGCAATTCAGGCAAATCGGAGAGGTTCTTGGAAGTTTAAACGCGCTTATGGTATTACAAGACGATATCTTAATCAACCAGAGACAATGCTGTTTGCTGTTAGATATTTTCAGTTTGGGTTTCAACACCGTTGCCGAAGAGATCAGGCACAACTTGAAGCTTGAAGAGAAGCACTCTAAATGGAGAGCCCTTGAGCAGCCTTTGAGAGAGCTCTATCGAGTCTTTAAAGAAGGCGAAATGTATGTTCGGAACTGTATGTCCAATAAAGATTGGTGGGGCAAAGTTATCAACTTTCATCAGAACAAAGATTGTGTTGAGTTTCACATACACAACTTGCTATCTTACTTTCCGGCTGTGATCGAAGCTATCGAGACGGCTGGAGAGATTTCTGGTCTCGACCCTGCAGAGATGGAGAGGAGGAGAGTTGTGTTTTCAAGAAAGTATGATAAAGAGTGGAATGATCCTAAGCTTTTCCAGTGGAGGTTTGGGAAACAGTATTTGGTTCCAAGGGATCTTTGCAGCCGGTTTGAGCATTCATGGAGAGAAGATAGGTGGAATCTGGTGGAAGCATTGCAAGAGAAGAGAAAGTCCAAGAGCGATGAGATTGGGAAGACTGAGAAGCGTTTAGCTGATTTTCTGTTGAAGAAGCTAACCGGGTTGGAACAGTTTAACGGTAAGCTTTTCCCGAGTTCGATACTCGTTGGTTCTAAGGATTACCAAGTGAGGCGGAGATTAGGTGCTGGTGGTCAGTACAAGGAGATTCAATGGTTAGGGGATAGTTTCGTATTGAGACATTTTTTCGGGGATCTTGAGCCGTTAAACGCAGAGATTTCTTCTCTGTTATCGCTTTGTCACTCGAATATACTTCAGTACCTTTGTGGATTCTATGATGAAGAAAGGAAAGAATGTTCTCTGGTTATGGAGTTAATGCACAAAGACTTGAAAAGCTACATGAAGGAGAATTGTGGGCCTAGAAGAAGATATCTCTTCTCTGTTCCTGTTGTGATTGATATAATGCTTCAAATCGCGAGAGGGATGGAGTATCTTCATTCAAATGAGATTTTCCATGGAGATTTGAATCCCATGAACATTCTTTTGAAAGAGAGAAGCCACACCGAAGGCTATTTCCACGCGAAAATTTCAGGATTTGGTTTGACATCAGTCAAAAATCAGTCTTTTTCTCGGGCTTCTTCTAGACCAACCACACCTGATCCTATAATCTGGTATGCACCTGAAGTTCTCGCAGAGATGGAAAAAGATCTCAAAGGTACAGCTCCGAGATCAAAGTTTACACATAAAGCTGATGTCTATAGCTTTGCAATGGTGTGTTTCGAGCTTATAACCGGTAAAGTTCCATTTGAAGATAGCCATCTTCAAGGCGATAACATGGCTAAGAACATTAGAACGGGAGAGAGACCTCTCTTCCCTTTCCCTTCGCCGAAGTACCTAGTCAGTCTAATTAAACGGTGCTGGCATTCAGAACCAAATCAGCGTCCGACTTTCTCTTCCATTTGCCGGATCCTGCGCTACATCAAGAAGTTTCTGGTTGTGAATCCGGATCAAGGTCACCTCCAAATCCAAAATCCACTTATCGATTGCTGCGACTTGGAAGCAAGATTCTTGAAAAAGTTCTCTCTGGAGTCAGGTTCTCACGCGGGATCCGTGACACAGATACCGTTCCAGCTTTACTCATACAGGATTACAGAGAAAGAGAAGATGAGTCCAAACTTCAACAAAGAAGAGAATTCAGAAACAGGCGAATCAATATCCGAAAGTGTTTCAGTAGTTGAAGATCCACCAACAATGCCAACGTATACAAAGTCATTGTGCTTTGATGCAAAATCTGAATACTCAGATACAAGATCAGTTTACTCAGAAGCTCCAATCAAAAAGACCTCAGCTTTAAAGCAAAGCACTGACGCCGTAAAACTCAGAAGAAACTCAATCTCAGGTTTACGATCTTCAGGATCGTCGCCTATAAAACCAAGATCAGCAACAAAAGCATCATCACCGCTAAGTCCATTTGGAAGAAGCAGCAGCAAAGCAAGGAAGGATGCTAGATTGCCTTTGAGTCCTATGAGTCCTTTGAGCCATGGGAGACGTAGACAACTCTCTGGTCCTGCTTCTGACTCTGAGCTAACATAG

>C_rubella_1

ATGGAGCAATTCAGACAAATCGGTGAAGTTCTTGGAAGTCTTAATGCACTTATGGTATTACGAGATGATATCTTGATCAACCAAAGACAGTGTTGTCTCTTGTTAGAAATCTTCAGTTTAGCTTTCAACACCCTAGCAGAAGAGATCAGACAGAATCTCAAGCTTGAGGAGAAGCAAACTAAATGGAGAGCTCTTGAGCAACCTTTAAGAGAGCTTTACAGAGTTTTTAAAGAAGGCGAGTCTTATGTTAAGCAGTGTATGGACAGTAGTGACTGGTGGGGTAAAGTTATCAATCTTCATCAGAACAAAGATTGTGTTGAGTTTCATATACACAACTTGTTCTGTTATTTCTCTGCTGTTGTTGAAGCCATCGAAGATGCTGGGGAGATTTCGGGTCTTGACCCTTCTGAGATGGAGAGGAGGAGAGTTGTGTTTTCAAGAAAGTATGATAGAGAGTGGAATGATCCTAAGCTGTTTCAATGGAGGTTTGGGAAACAGTATCTGGTGTCTAGAGATATTTGCAGCCGGTTTGAGCATTCTTGGAGAGAAGATAGATGGAATCTAGTGGAGGCATTACAAGAGAAGAGGAAATCGGACAGCGATGATATTGGTAAAACCGAGAAGCGTTTAGCTGATTTGCTTTTGAAGAAGTTAACCGGTTTGGAGCAGTTTAACGGGAAGCTGTTTCCGAGCTCTATACTTCTAGGTTCAAAGGATTACCAAGTGAAAAGAAGGTTAGATGCAGATGGACAATACAAGGAGATACAATGGCTAGGTGATAGTTTTACAGTGAGGCACTTCTTCAGCGATCTTGAGCCGCTAAGTTCTGAAATTTCATCTCTTTTAGCGCTTTGTCATTCGAACATACTTCAGTACCTCTGTGGATTCTATGATGAAGAAAGGAAAGAATGTTTTTTGGTTATGGAGTTGATGCACAAAGATCTGCAGAGCTACATGAAAGAGAACTGCGGACCAAGACGAAGATATCTGTTCTCGGTTCCAGTCGTGATTGATATCATGCTGCAAATTGCAAGAGGAATGGAGTATCTCCATGGAAATGATATCTTCCATGGAGATTTAAACCCCTTGAACATTCTTTTGAAAGAGAGGAGTCATACTGAAGGTTATTTCCATGCGAAAATTTCTGGATTCGGTTTATCTTCTGTCAAAGCTCAGTCTTCTCGGTCTTCCTCGAGACCAGGCACTCCTGATCCTGTGATCTGGTATGCACCTGAAGTTCTAGCAGAGATGGAACAAGACCTGAATGGTACAACTCCGAAGTCCAAGTTGACTCATAAAGCTGATGTGTATAGTTTTGCAATGGTGTGTTTTGAGCTCATAACAGGTAAAGTTCCATTTGAAGATAGTCATCTTCAAGGTGAACAAATGGCCATCAACATTAGGATGGGAGAGAGACCTCTTTTCCCTTTCCCTTCACCAAAATACCTCGTTAGTCTGATCAAACGTTGCTGGCACTCAGAACCGAGCCAGCGTCCTAACTTCTCTTCGATTTGCCGGATACTACGCTACATCAAGAAGTTCCTAGTTGTGAATCCAGATCATGGTCATCCTCAAATGCAAACTCCACTAGTTGATTGTTGGGATTTAGAAGCAAGGTTCTTGAGAAAGTTTCCAAGTGATGCAGGGTCTCACACCGCATCCGTGAACCAAATCCCTTTCCAACTCTTCTCATACCGGGTTTCGGAAAGAGAGAAGATGAATCCAAACTCAAAAGAAAGCTCAGAAGGAAGTAGTGAATCTGAAAGCGTTTCAGTGGTTGAAGATCCACCCAATGCCATGAGTACAAGAGATACAAAATCATTGTGTCAAGATACAGTATCTGAATACTCAGATACAAGATCAGTCTACTCAGAAGCTCCCATCAAAAAGGTCTCAGCTTTAAAGAAAAGTGGTGAAATGGTAAAGCTCAGAAAAAGTCCAAGCTTAGGTTCAGAAAAGTTGAGATCTATGGGAACTTCACCGGTGAAAGCAAGATCATCGCCAAAAGCATCACCATTGAATCCATTTGGAAGAAGCATAAAAGCAAGGAAAGATAATAGATTGCCTTTGAGTCCAATGAGTCCTTTAAGTCCAGCAATACGCAGAAAACAAACTGGTCATGCTTCAGATTCTGAGCTTACTTAG

>C_rubella_2

ATGGAGCAATTCAGGCAAATCGGAGAGGTTCTTGGAAGCTTAAACGCGCTTATGGTATTACAAGACGATATCTTAATCAACCAGAGACAATGCTGTTTGCTGTTAGATATTTTCAGTTTGGGTTTCAACACCGTTGCCGAAGAGATCAGGCATAACTTGAAGCTTGAAGAGAAGCACTCTAAATGGAGAGCCCTTGAGCAGCCTTTGAGAGAGCTCTATCGAGTCTTTAAAGAAGGCGAAATGTATGTTCGGAATTGTATGTCCAATAAAGATTGGTGGGGCAAAGTTATCAACTTTCATCAGAACAAAGATTGTGTTGAGTTTCACATACACAACTTGCTATCTTACTTTCCGGCTGTGATCGAAGCTATCGAGACGGCTGGAGAGATTTCTGGTCTCGACCCTGCAGAGATGGAGAGGAGGAGAGTTGTGTTTTCAAGAAAGTATGATAAAGAGTGGAATGATCCTAAGCTTTTCCAGTGGAGGTTTGGGAAACAGTATTTGGTTCCAAGGGATCTTTGCAGCCGGTTTGAGCATTCATGGAGAGAAGATAGGTGGAATCTGGTGGAAGCATTGCAAGAGAAGAGAAAGTCCAAGAGCGATGAGATTGGGAAGACTGAGAAGCGTTTAGCTGATTTTCTGTTGAAGAAGCTAACCGGGTTGGAACAGTTTAACGGTAAGCTTTTCCCGAGTTCGATACTCGTTGGTTCTAAGGATTACCAAGTGAGGCGGAGATTAGGTGCTGGTGGTCAGTACAAGGAGATTCAATGGTTAGGGGATAGTTTCGTATTGAGACATTTTTTCGGGGATCTTGAGCCATTAAACACAGAGATTTCTTCTCTGTTATCGCTTTGTCACTCGAATATACTTCAGTACCTTTGTGGATTCTATGATGAAGAAAGGAAAGAATGTTCTCTGGTTATGGAGTTAATGCACAAAGACTTGAAAAGCTACATGAAGGAGAATTGTGGACCTAGAAGAAGATATCTCTTCTCTGTTACTGTTGTGATCGATATAATGCTTCAAATCGCGAGAGGGATGGAGTATCTTCATTCAAATGAGATTTTCCATGGAGATTTGAATCCCATGAACATTCTTTTGAAAGAGAGAAGCCACACCGAAGGCTATTTCCACGCGAAAATTTCAGGATTTGGTTTGACATCAGTCAAAAATCAGTCTTTTTCTCGGGCTTCTTCTAGACCAACCACACCTGATCCTATAATCTGGTATGCACCTGAAGTTCTCGCAGAGATGGAAAAAGATCTCAAAGGTACAGCTCCGAGATCAAAGTTTACACATAAAGCTGATGTCTATAGCTTTGCAATGGTGTGTTTCGAGCTTATAACCGGTAAAGTTCCATTTGAAGATAGCCATCTTCAAGGCGATAACATGGCTAAGAACATTAGAACGGGAGAGAGACCTCTCTTCCCTTTCCCTTCGCCGAAGTACCTAGTCAGTCTAATTAAACGGTGCTGGCATTCAGAACCAAATCAGCGTCCGACTTTCTCTTCCATTTGCCGGATCCTGCGCTACATCAAGAAGTTTCTAGTTGTGAATCCGGATCAAGGTCACCTCCAAATCCAAAATCCACTTATCGATTGCTGCGACTTGGAAGCAAGATTCTTGAAAAAATTCTCTCTGGAGTCAGGTTCTCATGCGGGATCCGTGACACAGATACCGTTCCAGCTATACTCATACAGGATTACAGAGAAAGAGAAGATGAGTCCAAACTTCAACAAAGAAGAGAATTCAGAAACAGGCGAGTCAATTTCCGAAAGTGTTTCAGTGGTTGAAGATCCACCAACAATGCCAACGTATACAAAGTCATTGTGCTTTGATGCAAAATCTGAATACTCAGATACAAGATCAGTTTACTCAGAAGCTCCAATCAAAAAGACCTCAGCTTTAAAGCAAAGCACTGACGCCGTAAAACTCAGAAGAAACTCAATCTCAGGTTTACGATCTTCAGGATCGTCGCCTATAAAACCAAGATCAGCAACAAAAGCATCATCACCGCTAAGTCCATTTGGAAGAAGCAGCAGCAAAGCAAGGAAGGATGCTAGATTGCCTTTGAGTCCTATGAGTCCTTTGAGCCATGGGAGACGTAGACAACTCTCTGGTCCTGCTTCTGACTCTGAGCTAACATAG

>C_rubella_3

ATGGAGCAATTCCGAGAGATAGGAGAGGTGTTAGGAAGCATAAGAGCGTTAATGGTGTTCAAGGACGACATTCAAATCAACCAGCGTCAATGCGCTCTACTACTCGACCTCTTCATCGCGGCCTACGATTCGATTTCCCGATCCATGCGATCTAATCTACGCTTCAAAGAGAAGAACACCAAATGGAAGATTCTAGAACAGCCCTTTAGGGAGCTTCTATGGGTGGTAAGAGAAGGGGAGGCTTATGTTAGAATGTCTTTAGAGCCTAAACTTGGGTTTTGGGCTAAGGCTATCCTGTTGCATTCCAATAGAGATTGCAACGAGCTTCACATACATAACTTACTCTCTTGTCTACCGATCATTGTCGAGGCTATCGAGACAGCGGGTGAGGTTTCTGGCTGGGATGAAGAAGAAATGAATAAGAAGAAGTTGGTACATTCTAACAAGTATATGAAGCAATGGAATGACTCTCAGATGTTTACCTGGAAGTTCGGGAGAGAGTATTTGGTGACAGAGGATTTCTGCAACCGGTTCCAGAGTGCTTGGACAGAGGACCTTTGGATTTTAATCAAGGAGCTTCAGGAGAAAAGGCAGCCCGGTTCAAGCAAACATGATCGAAAAATGGCTGATTTCCTTTTCAAGAAACTGGGAGATGGAAACGAGTGTCCCAAGCTATTTCCGTCTTCTCTTCTGGTTAGCGCGAAAGATTACCAGGTGAAGAAAAGATTAGGGAATGGAAGTCAGTACAAGGAGATTACATGGTTAGGCGAGAGCTTTGCGCTTAGGCATTTCTTTGGGGATATCGATGCTTTGCTTCCTCAAATTACTCCACTGCTATCTCTCTCCCACCCAAACATTGTATATTACCTCTGCGGATTCACAGATGAGGAGAAGAAAGAATGTTTCTTGGTTACAGAACTGATGAGTAAAACACTAGGTATGCACATCAAAGAGGTATGTGGTCCAAGGAAAAAGAACACTCTCTCCCTCCCCGTAGCGGTTGATCTGATGCTTCAGATAGCTTTGGGTATGGAATATCTTCACTCGAAGAAAATATACCATGGAGAGTTGAATCCATCAAACATCCTTGTCAAATCAAGAAAACTTAACCAATCTGGAGATGGGTATCTGCACGGGAAGATTTTTGGGTTCGGGCTGAATTCTGTCAAAGGGTTCCCTGGTAAAAGTGCTTCTTTGACGGGTCAAAATGAGAATTTTCCATTCATATGGTATTCCCCAGAAGTTCTAGAGGAGCAAGAACAGAGCGGAACTGTAGGAAGCTTCAAGTATACTGATAAATCTGATGTATATAGCTTCGGTATGGTTTGCTTTGAGCTTTTGACTGGTAAAGTGCCATTCGAAGATAGCCACCTGCAAGGAGATAAGATGAGTAGAAACATTAGGGCAGGGGAGAGACCGCTTTTTCCATTTCATTCACCTAGATTGATAACAAACCTGACGAAACGATGTTGGCATGCTGATCCGAATCAGCGTCCCACTTTCTCATCGATAAGCAGAATTCTCCGGTACATCAAACGGTTTCTAGCTTTGAACCAAGAATGCCATAGTAGTAGCCAACAAGACCCGCCCATTGCTCCCCCTGTGGATTACTGTGAGATCGAGACTAAACTATTGCAGAAGCTTTCATGGGAATCGACAGAGTTAACTCAAGTTTCCCAAGTTCCTTTTCAAATGTTTGCATATAGGGTTGTGGAACGAGCAAAAACTTGTAAGAAACATAATCTCGAGACATCTGAATCGGGTAGTGAATGGGCTTCATGTAGCGAGGATGAGGGTGGAACAGGATCAGATGAGCAAGTGTCATTTGAGAAGGAAAGAAGACTACCATGCTCAATTGATGTTGGTATGAGCAAGAAGCATGCTTCAAAACTTATGAAAAGAGCTTCTGGCCTGAAACCAATTCAAAAAACAGGTGATTCTCAGTAG

>C_sativus_1

ATGGAGCAATTTCGGCAGCTTGGAGAGGCATTAGGAAGTGTAAAAGCGCTCATGCTATTCAAAGATAGCGTTCATATAAATCAGAGGCAATGCTGTTTGTTGCTTGATGTTTTGAGCTTTGCTTATGATTCAGTGGCAGAGGAGATGAAACAGAATCTTCGATTTGAAGAGAAGCATACTAGATGGAAGGTTTTGGATCAGCCTTTGAGAGACCTAAACAGGGTATTCAAAGAAGCAGAATGGTATATTAGGCAATGTTTGGAAACAAAAGATTGGTGGGCAAAAGTGATTATGCTATATCAGAATACTGATTGTATTGAATTTCACATTCATAATCTGTTGTACAGCATTACAGTTGTCGTTGAAGCCATTGAAATGGCTGGGGAGAGTTCTGGTAGTGATCATGATGAGTTGCTGAAGAAGAAACTAATCAACTCCATCAAATACAGGAGAGAACACAAGGATTTGAAGATTTTCAAGTGGAAATTTGGGAAACAATACCTTGTTACTCAAGATTTCTGTAATCGGATTGAGGCGGTTTGGAATGAGGATAGATGGTTTCTGCTCAATAAGATCCGGGAAAAGAAATTAATGGCTTCGTCCAAGTACGAACAACGATTAACAGATCATCTTCTGAAAAACATTAATGGGTCAGAATCTTTTAATGGAAAGCTATTGCCAAGTTTAATGCTAGTGGGATCAAAGGATTACCAGGTAAGGAGAAGATTAGGGGTTGGGAGTCAATATAAAGAGATTCTCTGGTTAGGGGAAAGTTTTGCTATGAGACATTTTTTTGGGGAAATTGAATCCTTAATTCCAGAGATATCAATGCTGTTATCTCTTTCTCATCCAAACATCACAAGATTTCTTTGTGGGTTTACTGATGAGGAAAAGAAAGAGTGTTTCTTGATTATGGAACTCATGAGTAGAGATTTGTCAGGCTATGTAAAGGAGATTTGTGGCCCTCGCAAGAGAATTCCATTTACTCTTCCTGTAGCTCTAGATTTGATGCTTCAGATTGCAAGAGGAATGGAATATCTTCACTCAAAGAAGGTCTACCATGGTGATTTAAACCCTTGCAACATTCTGGTTAAGCCGAGAGCTTATTCTACAGATGGCTATGTACATGGTCAGGTCTCAGGGTTTGGCCTACCTGCTGTCAAGTTTAAAAACTCTTCCAATCAGAATGAATCTCTCCCGTTCATATGGTATGCTCCAGAAGTTTTAGAAGAACAGGACCAATCAGGAAGTGCTGAGAGTTGTAAGTATACAGAGAAGTCTGATGTGTACAGCTTTGGCATGGTTTGTTTTGAAGTTCTGACTGGGAAAGTCCCTTTTGAGGATAGCCATCTTCAAGGGGATAAAATGAGTCGAAACATTAGAGCAGGAGAGAGGCCACTTTTTCCTCATAGCATGCCAAAATATGTGACGAACTTGACCAAGAGGTGTTGGCAAACCGACCCGAATCAAAGGCCGAGCTTCACGTCTATTTGTAGGATTCTTCGATATACAAAGCGATTTGTTGCAATGAATCCTGATTACAATAGCCAGACAGATCCTGCTATGCCAACAGTAGATTACTGCGACATTGAATCAGGTTTATTGAGGAGGTTGCCATCTTGTGGAATAAGCGATGCTGCATCACCAATTACAGATATTCCCTTTCAGATGTTTGCATACAGAGTAGTTGAAAAAGAGAGAGCTGGTGCCACCTTTAAAGATACCTCGGAATCAGGAAGTGATGCCTCAGCATGTGGGGATGAAACAGCATCATCAATAGACGATCCTTTCCCGACGCCGGTTGAGAGAAAATTACCAGCTCCGCGTGAAGGTAGTAGGAGGCTTTCATTGACCAAGAATAATTCTGATGTCAGACTGAACAAGCTACCAGGCACACCAAAAGGGCGATTTTCAAGGCCTCCGCAAATCAGTCCTCGTGGGCGAAGTATGCGAATGAATTCTGAAAGCCAGCTTATGGCAATAAGTCCAAAGATTAGAAGATTATCTGGTCATGCCTCAGATTCTGAGCTACCCTAA

>C_sativus_2

ATGGAACAATTCCGGCGTATTGGAGAGGTATTGGGAAGCTTAAAGGCTCTTATGGTGTTGCAAGATGATATTCAATTCAACCAGCGTCAATGTTGTTTACTTCATGATATGTTTAGGTTGGCTTTTGATACCATTGCCGGAGAGATTAGGGATAATCTTAAGCTTGAAGAGAAGAACACCAAGTGGAAAGCTCTTGAACAGCCTTTGAGGGAATTGCATAGAGTTTTTAAAGAAGGGGAGCTTTACATTAAGCAATGCATAGATGGCAAAGACTGGTGGGCTAAAGTAATCAGCTTCCATCACAGCAAAGACTGCATTGAATTCCATGTCCATAACTTGCTTTCATGCTTTCCTGCTGTCATTGAAGCGATTGAGACGGCTGGAGAGATATCAGGGCTCGATCAGGACGAGATGCAGAAGAGGAGGCTTGTACTTATGAGGAAATATGATATGGAATGGAATGACTTGAAACTGTTCCACTGGAGATTTGGGAAACAGTATTTGGTTCCTCGAGAGATACGTAATCGGATGCAGAGCGTTTTGAGAGAAGATCGATGGCTGCTAGTTGAAGCACTCAAAGAGAAGATAAGTTCACCAGGAACAGCTGTTTCAAAGAACGAGCAGCAGCTTGGTGAATTGTTGATAAAGAAATTAAACAACTCAGAACCGTCGAAGGCAAAGCTGTTCCCGAGTTCGATTTTAGTTGGAACAAAGGATTATCAGGTGAGGAGACGGTTGGACGGAGGGCAGTCTAAGGAAGTTCAATGGTTTGGAGAGAACTTTGGAATGAGACAGTTCACGGCGGAAACTGAAGAAACAGAATCCGAGGTTCCAATTCTTTTATCACTTTTACATCCTAATATATTGCAATATCTCTGTGGCTTTTTGGATGAGGAGAAGAAAGAGTACTTTCTTGTAACTGAGTTGATGTCCAAGGATCTTTCTTCCTACATGAAGGATAACAACGGAGCGAGGAGAAGACTCTTGTTTCCTCTCCATGTTTCGGTTGATATCATGCTTCAAATTGCTAGAGGCATGGAATATCTTCACTCTCAAATGATCTATCACGGAGACTTAAACCCTTCCAATGTATTTATGAAGCCAAGAAACTCTTCAGAAGGCTCTTATCTAGTAAAAGTCTCCGGTTTCGGTTTATCATCCGTCAAGAATTCACCTCCCAGAAACTCAACAAACCAACTAGAAACCAACCCTTTTATCTGGCATGCCCCAGAAGTGATGGCAGAGCAAGAACAACAAGCTCCAGGGACTGTTTCTTTTTTCCGAAAGACAGAGAAAGCAGATGTTTACAGTTTTGGAATGCTTTGTTTTGAGCTTTTGACTGGAAAGGTTCCATTTGAGGACAGCCATTTACAAGGGGAGAAGATGAGTCGCAATATCCGAGCAGGAGAGAGACCCCTCTTCCCGTTTCCAACTCCTAAATACTTGGTCAGCCTTACGAAAAGATGCTGGCACTCTGACCCGTCTCAGCGGTTATCTTTCTCTTCCATTTGTCGAATCCTTCGCCAGGTGAAGAAATTCCTCGCTATGAACCCTGCTGAAAGCAATCAGCCCGAACTGCAAATGCCTACCGTTGATTACTGCGACGTTGAAGCAGGAGTTGCAAGGAAGTTCTCCTCAGATGGGGTTGGTGATTTGTGTTCAGTTTCACAGATTCCATTCCAAATGTTTGCTTATAGACTTGCAGAGAAAGAGAAGACAAATCCAAGCAAAATCAAGACTTGGGATTCTGCAAGTGATGTGGTTTCCATTAGTAAGGATGATTGTGCATCCATTTACAGGGATGACACAGTTTCTGTAATAGAAGATCCATTCACCATCCCAGCAAGTGATACAAGATCCTTTTATTCCGATATGAGATCTGTCTATTCTGAAGCTCCATCCAAGAAAATGCCGATTACAAAGAAAGTTCCAGATACAAAGATCAAAAGAGGCACAGGGATCCCAGAAACAAAAACACGAATGACTTCACGAACACCATCAAGAACACCGACACGAACAACTGCACGACCTCGAGCATTGAAGACAAACAGAGATATCCCTTTACCATTTTCTAGCCCTCTGAGCAAGGGAAGAAGGAGACTGAATGGTCATAGATTTGATACAACATGGCAAACGAATGGACGATTAGAGAAAGTGACAAGTGCAGCAGCATACTCTCGATACTACTTTCTGTTTGAG

>C_sinensis_1

ATGGAGCAATTCAGGCAGGTGGGCGAGGTTCTGGGAAGTTTAAAGGCTCTCATGGTGTTGAAAGATGATATTCAAATCAATCAGCGACAGTGTTGTTTGCTTCTTGACATCTTTACTTTAGCCTTTTCCACAGTTGCTGAGGAAATCAGGCTTTTCTTAAAATCGGAAGAGAAGAACACAAAGTGGAGAGCTCTTGAACAACCTTTGAGAGAGCTTCAGAAGGTGTTTAGAGAAGGAGAACATTATGTTAAGCAATGCTTGGATTATAAAGATTGGTGGGGTAAAGCAATTAGTCTCCATCAAAACAAAGATTGTGTTGAATTTCATATTCACAATTTGCTTTGCTACTTCCCCGCTGTGATTGAGGCCATTGAGGCTGCTGGTGAAATCTCAGGGCTTGACCCAGATGAGATGCAAAGAAGGAGAGTAGCATTTGCAAGAAAATATGACAGGGAATGGAATGACCCTAAACTTTTCCAATTGAGATTTGGGAAGGAGTATTTGATCCCTCGAGAAGTCTGTAATGAGTTTGAGAGTGCTTATAAGGAAGATAAATGGCTTCTTATTGACGCACTGAAGGAGAAGAAAAGATTAGGATCTGTTGTTTTGACAAAGAACGAGCAGCGGCTTGTAGACATGCTGCTCAAGAAACTAATGGTGAGAAGAAGATTAGGTGCAAGCAGCCAGTTTAAGGAAATTCAATGGTTAGGAGATAGCTTTGTCCTGAGGCATTTCTACGGGGAACTGGAATCGCTAAATGCCGAGATTTCGACTATGCTCTCCCTTTCACACCCCAACATAGTGCAATACCTATGTGGATTTTGTGATGAGGAGAAAAAGGAATTTTTTCTCGTAATGGAGTTGATGAGCAAGGATTTGTCCTGCTACATGAGGGAGACATTTGGTTCAAGAAGGCGGAATTCATTCTCTCTACCTGTTGTTGTTGATATCATGCTTCAGATTGCAAGAGGTATGGAATTTCTTCATGCACAAAAGATCTATCACGGAGAATTGAACCCTTCAAATATTTACCTCAAGGCCAGGAGTATGGAAGGCTATTTTCATGTAAAAGTTTCTGGGTTTGGTTTATCCACTGCCAGAACTTACGCATCCCGAAACACACCACCAGCATCACCACAAAACCAGACTGCGCCAAACCCTTATATATGGTATGCCCCGGAAGTTCTGGCGGAGCAAGAAGGGACTGGAAGTACTTCTACTTCCAAGTGCTCAGAGAAGGCAGATGTCTACAGCTTTGGAATGCTTTGCTTTGAGCTTTTAACTGGGAAAGTTCCTTTTGAAGATGGGCATCTTCAAGGAGACAAGATGACCAAAAATATAAGAGCAGGGGAGAGGCCTCTCTTCCCATCAGGTTCTCCGAAATACCTCGTGAACTTAACCAAAAAATGTTGGCACACAAATCCATCTCAGCGTCCAAGTTTCTCGTCCATTTGTCGAATTTTACGTTACATCAAGAAGTTCATGGCAAATAACCCTGACATCGCACGGTCTGAGTTTCAATCACCGCTAGCAGATTACTGCGACATAGAGGCAGGATTTGTAAGGAAATTTGTAGGAGAGGGCTGCCCTGATGTAGCGCCAGTATCACAAATTCCATTTCAAATGTTTGCATATAAAATAGCTGAAAAAGAGAAGGCTAATTCGAGTAATAAAGGTAAGCATTGGGAGTTAGGTAGTGATGTAGCCTCAACTTGTAAGGATGAGGTTTTTTCCGTAGGAGATGAGGGAATGGTACCAGTGACCGAGACAAGATCAGTTTGCTCTGACGTGAAATCTGTTACTTTTGATATGAGATCAGTTTATTCTGAGGCTTTTGGCAAGAAAGTTTCAAATTTTGATACAAGGTCAGTAAGATCCGAGTCCCCGAACAAGAAAATGCAGGGTAGTGTTTCAAGGTTGATTCAATCCATGGTTCTGGAGAAGAATGTTTCCAGTTCTGATACTAAACCAGCGGTTCCGGAGAACAGTATTTCTAATTCAGAAGTTAAATCCGAGGTTCCAGAGACGAATATATCTAATTCTGATACTAAACTTGTGGTTTCAGAGAATAACATCTACAATACTGACACTAAATCTGCCTGTTCTGACAATATGCCGGAGAAAAATATCTGTAATGGTGATACAAGATCTGTTTGTTCCAAGAATTCGGGCAAGATCAAGCAATCCACTGGTGTCAAGGTCAAAAAAGGTCCAGGGACAACAAAAATGAGATCAGGAAGAGCAGCACCAAAAGCACGATCACCAAGAACATCAACATCTCAATCACTGCGATCATCAACTTCTCAATCACCAAGGCCACCTCCCAAATCACCACCGTTTATTCCGTCTGGACGCTGTTTGAAGAAGGGCCAAGACCTTAGAAGCCCAAGCAGAAGAAGATCATCAGTCAATGTCTCAAATTCTGAGGTAGCATAG

>C_sinensis_2

ATGGAGCAATTTGGACGAATTGGAGAGGTCATGGGAAGTATGAAGGCGCTAATGGTATTCAGAGACAGTATACAAATAAATCAAAGGCAATGCTGTTTGTTGCTAGACATTTTCAGTTTTGCGTATGATTCAATAGCGGAAGAGATGAGACAAAACCTGAGATTCGAAGAGAGACGCGCGAAATGGAAAATTCTTGAACAACCATTGAAAGAGCTCTTCAGGATATTTAAAGAAGGAGAAAATTACATTAAACAGTGCTTGGAAATTAGGGACTGGTGGGCTAAAGCCATTACCCTTTATCAGAACACGGACTGTGTTGAGTTTTACATCCATAATTTACTTAGTTGCATTCCGATCTTGATCGAAGCAATAGAGACGGCAGCAGAATTCTCTGGATGGGATCCTGATCAGATGCACAAGAAGAAACTTGTACACTCCGGCAAGTATAAAAAAGAATGGAAGGACCACAAACTTTTCCAATGGAGATACGGGAAGCAGTATCTTATTACCCCAGATTTTTGCTATAGGATAGACACCGTATGGAAAGAAGATAGGTGGATCCTTTTCAACAAAATCCAGCAAAAGAAAATATCAGGTTCAACTAAGCAAGAGCAGGGACTCATAGATGTCCTTTTCAAAAACTTGGATGGTTCAGGATCTTTGAGTGGGAAACTCTTACCAAGTAGAATCCTAATTAAGTCTGAGGATTACCAGGTGCGGCGAAGATTAGGGAGTGGGAGTCAATACAAGGAGATCCTGTGGTTGGGTGAAAGTTTTGCTTTGAGACACTTTTTTGGGGACATTGAACCCTTAGTTCCTGAGATTTCTTCTCTATTATCTCTTTCCCACCCGAATATCATGCATTTCCTGTGCGGGTTTACCGATGAGGAAAAGAAAGAGTGTTTTCTGATTATGGAACTAATGAGTAGAGACCTTTGCAGCTACATCAAAGAGATTTGTTGCCCAAGAAAGAGAATTCCATTTTCTCTTCCGGTTGCTGTTGATCTTATGCTTCAAATTGCTAGAGGAATGGAGTATCTCCACTCAAAGAAAATCTATCATGGTAATCTAAATCCTTCTAACATCCTCCTTAAACCCAGAGGTGCCTCAACAGAAGGGTACCTGCATGCAAAGATTTCAGGATTTGGCCTGTCTTCGGTGAAAAACTTTGGCCCAAAAAGTCCATCTCAGAGTGGGACTACCCATCCATTTATATGGCATGCTCCAGAAGTTCTAGAAGAGAATGAGCAGACAGAAAGTGCATCAAATTCAAAGTACAGCGAAAAGTCTGATGTTTACAGTTTTGGAATGATCTGCTTTGAGATTCTCACGGGGAAAGTTCCATTTGAAGATGCTCACCTTCAAGGAGACAAAATGAGTCGAAACATTAGGGCAGGAGAGAGGCCATTGTTCCCATTTCATTCACCAAAGTATGTGACAAATTTGACAAAGAGATGTTGGCACGCTGACCCAAATCAACGGCCAAGCTTCTCATCCATCTGTAGAATTCTTCGCTATATAAAACGGTTCATTATGATGAACCCTCATTATAATAGCCAGCCAGACCCGCCAATGCCTTTGGTAGATTATAGCGACATTGAGTCAAGGCTTTTGAGAAAATTCCCTTCTTGGGAGACTCACAATGTTTTGCCAATATCAGAGATCCCCTTTCAAATGTTTGTTTATAGAGTTGTGGAGAAAGAGAAAATAAGTTCAAGCCCTAAAGACACATCAGACTCAGGAAGTGATAAAGCTTCTGTAAGTGGGGACGAAAATATGACCACACCAGAGGATCCATCTCCACCTGTGACTGAAAGAAAGTTTTTACCTTCACCCGAAGCTGTGAACAAGAAACTCTCCGGTGTCAAGAAACTGTCAGAATCGAAAGTTATTAAACAAACAGGGACACCAAAAGGGCGAGCTGTAAGACCTCCGCAACTAATTCCTTGCGGTCGTAGTTTAAGAATGAGCTCAGAAGGGCAGCTAATGGCGATGAGTCCAAGAATACAGAGAACATCCTCTGGCCATGTGTCGGATTCTGAGCTCTCCTAG

>S_melongena_1

ATGGATCAATTCAGGCAAATTGGTGAGGTAGTAGGGAGCTTAAATGCTCTAATGATATTAAAACATGACATTCAAATAAACCAGAAGCAATGTTGTCTGCTGTTTGACATGTATGTACAAGCCTTCGACACGATTTCCGAGGGGATCAAGCATAATTTAAGGTTGAATGAGAGGAATACAAAGTGGAAAGCACTTGAACATCCAATGAGGGAGCTTCATAGGATCTTTAAAGAAGGTGAAATGTACATAAAAAGTTGTTTGGATGTGAAAGATTTTTGGGGTAAAGCTATAAGTTTACATCTCAATAAGGATTGTGTTGAGTTTCATGTTCATAATTTGCTCTGTTGTTTCCCTGTGGTGATTGAGGCTATTGAGACAGCTGCAGAAATCTCGGGATTCGATGAGGAAGATATGCAGAAGAGGAGGAGTGCGCTCGCTAGAAAGTACGAGGGAGAAACAGTGTGTGATCCGAGATTCTTTCAATGGATGTTTGGGAAACAATACTTGGTTACTAGAGAGATATGTAGTGAATTGGAAAGTTGTTGGAAAGAAGATAGATGGTATCTTGTTGAAACGATAAGTCGAAAGATGAATGCTGCAGAAAATTTGGCGATAAACGAGCATAGGCTAGCTGAGATATTGCTTAAGAAATTAGATGGATCAAAACAAATCAAGCAAATAATGCTCTTGCCTAGTTCAATCTTGATTGGAGCAAGTGATTATCATGTGAAGAGACGTTTAGGCTCGCGGGGAGGACATGTTAAGGAGATTCAATGGTTAGGAGAAACGTTTGCGTTGAGGAACTTTTTTGGTGAGCTGATTGAACCAGTAGTTGCTGAAATTTCTCTAGTGTTTTCACTTTCACATCCCAACATTCTGCAATATCATTGTGGTTTTTACGACGAAGAAAAGAAAGAAGGGTACCTTGTTATGGAGCTAATGAACAAAAGTTTAGCGGCTTACATAAAAGAGCATTCTGGTCAGAGGAAGAAAGGACCGTTTACTGTCCAAGTTGCTGTCGATATTATGCTTCAGATTGCTCGAGGGATGGAGTATCTGCACTCGAGAAAGATCTATCATGGAGAGTTGAATCCATCTGATGTGCTCCTGAAACCAAGAAATTCCTCCGCGGATAGTTATTTTCATGCAAAAGTTAAAGGATTTGGCTTAACATCTATTAAGAGTAGCTATAAGGCTTGTGGTGACAATGCTGCTGAGTCTTTCGTATGGTATGCACCAGAAAGCAAATGCACTTACAAGTACACAGAGAAAGCTGATGTTTATAGTTATGGAATGATATGCTTTCAACTTCTAACGGGGAAAGCTCCGTTTGATGAATATCCCCAAGGGGAGAATATGGCGCGGAATATCAGAACAGGCGAGAGACCTCTCTTTCCGCATCCTTCACCTAAATATCTCGTTAACTTGACAAGAAAATGCTGGCAAACGAATCCAGAACTTCGCCCAAGTTTCTCTTCTCTGTGTAGGATTCTGCATTATATCAAGAAAGTACTTGTCATAAACCCAGGACATGGCCAACCTGAATGTCCCCCTCCTCTCGTAGACTACTGTGAAATCGAGGCTGGTTACTCTAAGAAGTTCCCGGGAGAGGATAGCACTGGTTTGGCCCCAGTATCACAAATTCCTTTCCAGATGTTTGCTTATAGACTTGTTGAGAAAGAGAGAATTTCTGGGAACTCTAAAGAAAAGCACTGGGATTCATCACATTATGGTTATTCAAGACACAGGACAGCATCAATGCAGAGCGCCGAAGAGCAAATGGATGCAGTAGATGATCTCTTTTTTGCTCCGAGCGATAGAAGGTCAGTTTGCTCAGAGATAATAGAGAGTAAAGATTCAAGATTCTGGGATCAGAGGTCAGTCATTTCTGAGACACCACTTCAAAAAGTCTCCTCATTTGATCAAATGTCAACTAGTTCTGAGGGTCCAAAAGAGAAATTTCCAGAAGCAGCAACAGGAGCAAATGAACAGACAATCTATGCTGATATTCCGGAAAGGAAAGCAAAACTAACAACATCAGGCAATCAGCGCTCATCGCGTGTTGATACAACACCAGGGTCGAAGAACGTTTCATCAATGAGAAAGAACCAAAAGCTGGATGAAATCCAGAAGAATGTAATGTCTCCGAAAACAGATGACAGAAAACCTTCATCGAGTAAGCAGAAAATAGCGGAGTCTCCTGTGATATCAGAGAAGAAAGATTCATCAGGTGATGATATATTAACTTATTCCGAGATTCAAGAAAAGAAACATGCACCAGAAGATCAGAAACCAACAAGTTCTGCAACTCCAGAGGAGAAAAACTCATCTGATAACCAAAAACTAAGAGATTCTCAAAGTCCAGAAAAGCAGCAGCCATCAACAGCTGCCAATGTCAAGCACATTCGTGGTGATTCTACGCAAAGGAAAGCTATGTCAAGAAGAACAAAACCCAAGAATCCAGAGAAGAAAGTTCTATCAAGTCCGGAAGCCAATCAAAACTCAATGTCATCTGAGACAACAGCGTCGAAACCAAGCAATTCAAAGGGATCAGTGAAGAAACCACCATTTCACAAAAAACTACAGGACAAAGGGAGTAGCACTACACCAGAGAAACTGACAGACCCATCGCCAAGATCTTCACCAGCTAGAGTTAAGAGAATACAGTCATCAACTATGTCTTCTCCAACACGAAGCCAAAAAGCTCCGTCTCCAAGATCACCTGTGATCACCTCGAATAGAAATGGATATCAATCTGCAAGTGCATCACCATTAAACCCGTGTTCTCGCTATAGCAGAGTAAACCGAGAGCTTCCTTTGTCCTTAGCTATGAGCCCACATAGACCAAAGAAACCTCATAGCTCTAGCAATAGCCAAAGTGTACACAACAGTTAA

>S_melongena_2

ATGGATCAATTTATGCAAATTGGTGAAGTAGTTGGTAGTTTGAAGGCTCTAATGGTATTGAAAAATGATATCCAAATCAATCAAAGGCAATGTTCTTTTCTAGTTGATATGTTTGTTCATTCTTTTGATACCATTTCTGAAGAAATCAAACATAATTTAAGGTTAAATGAGATGAATATAAAGTGGAAAGCTCTTGAGTTGCCAATGAAAGAGCTTCATAGGATCTTTAAAGAAGGCGAAGCGTACGTAATATACTGTTTGGATGTTAAAGATTTGTGGGGGAAAGCCATAAGTCTCCATATGAACAGAGATTGTGTTGAGTTTCATATTCATAACTTGCTGTGTTGCTTCCCTGTTGTTATTGAAGCTATTGAGACTGCTGCAGAAATCTCGGGGTTCGATGAGGAGGAAATGCAGAAGAGGAGAACCGCGTTAATGAGGAAATATGATAGGGAATTCATTGATCCGAGAATTTTTCAGTGGATTTCTGGGAAGCAGTATATGGTAACTCGCGAAATATGTAGCCGGTTGGAGAGTTGTTGGAAAGAGGATAGATGGTTGCTTATTGATATCATTAGGCAAAAGAAAACAGAAACTTTGTTGAAGTATGAACAAAGGCTTGGAGACTTATTGTTAAGGAAATTAGATGGTGTTGAGTCAACTAACAAGATCATGTTGCCGAGTTCAGTCTTAGTCGGATCAAATGACTATCATGTGAAGAGACGTTTAGGGTCGTGGGGCGGACATGTTAAGGAGATACAATGGTTAGGAGAGAGTTTTGCTCTGAGGAACTTTTTCGGAGAGGTTGAACCGTTACATGATGAGATTTCTTTGGTGTTTTCTTTGTCTCATCCCAACATTCTGCAATATCACTGTGGTTTTTATGATGAAGAGAGGAAAGAAGGGTACCTTGTTATGGAGCTGATGAACGATACGCTTGCAACGTACATAAAAGAGCATTCCGGCCAGAGGAAGAGACTGCCTTTTTCAACGTCTGCTGCAGTCGATATTATGCTTCAGATTGCAAAAGGGATGGAGTATCTGCACTCGAGAAAGATCTATCACGGAGAGTTGAACCCGTCTCATGTACTTCTTAGAGCAAGGAATTCCTCAGCTGAGAGCTATTTTCACGCGAAAGTCAAAGGGTTTGGCTTAACTTCAATCAAAAGTACTTACAAGACTGCTAACCACAACGCGGCTGATTCCATCATATGGTATGCCCCGGAAGTTCTAGCTGAACAAGAAAAACCAGGAAACAAGTGTGTCTACAAGTACACAGAAAAAGCTGATGTTTATAGCTTCGGAATGATTTGTTTCCAAATTTTAACGGGGAAGGTTCCGTTTGATGAAGGCCATTTGCAAGGGGAGAAAGTTGTGCGTAACGTCAGAGCTGGGGAGAGGCCTCTATTTCCCTATCCTTCACCTAAATACCTTGTCAACTTAACAAGAAGATGTTGGAACACAAATCCAAATCTTCGTCCACATTTCTCGTCTATATGCAGGATTCTTCGTTACATCAAGAAGGTCCTTGTCATAAATCCGGAACATGGCCAGCCTGAAACTGCCCCTCCACTTGTAGAGTACTGTGACATCGAGGCTGCCTACTCGAAGAAGTTTGCTGAAGAGGAGAGCAAGACGAGTTTGACCCCGGTATCACAGATTCCTTTTCAAATGATTGCTTACCGGCTCATTGAGAAAGAGAAGATTCTTGGAAAGAGCTGGGATCCATCAAATGATGGTTTTTCGGTCCATAGGAGAGAATCAATGCTGAGTGATGACGGGCGATTGTCTGTGATGGATGATCTCTTTCTTGCACCGAGTGATAGAAGGTCAGTTTGCTCCGAGATAATTGACAGGAAAGATTCAAGATTGTTTGATCAGAGGTCAGCTATTTCCGAGATACCACATAGAAGACTTTTCTTATTTGATCAAGCATCAGTTGGTTCTGAGAGTTCGGAGAGGAGATTCTTAGTTAACTTAAGGAAAGGAGCAGTATCATCACCAATAGTTAATAGGTCACCGCGTATTGACACACTGGAGAAAAGGATCATCTCAACGATGAGCAAGAACCAACGACTCAAATTTTTGGCCGACCAAGAGAAGGCAATGTCTCCAAAAGCAGGTGATGAAAAACCTTCATCAAGCGTAGCAGCTCAGCAAAATTTAGCATCTCCTCAGACTCCAGAGCAGATGAAATCAGCTGAAGAAAATTTACTATCCCCTCCGACTTCAGAGGAGATGAAACCAGCTGAGCAAAATTTAGTATCTCCACGGACTACAGAGACGAAGAAACCAGCTGAGCCAAATTTAGTATCTCCACGGACTACAGAGACGAAGAAACCAGCTGAGCCAAATTTAGTATCTCCACAGACTACAGAAACGAAGAAATCAGCTGAGCCAAATTTAGCATCTCCACAGACTACGGAGACGAAGAAATCAGCTGAGAAAAATTTAGTATCTCCTCCGACTACAGAGAAGAAAATTCTGTCTTATAATCAGAAACTGACGAGTTCTGAGACTCGAGCAAAGAAACACAAATCAGATGATCAGAAACTAATCAGTTCCGAGGCTCATGAGAAGAAACATGCATCTGATGATCAGAAGCTAATCACTTCCGAGGCTCATGAGAAGAAGCTTTCATCTGATGATCAGAAATTACGACGTTCCAAGACTTTAGATCAGAAAAAGTCATTAGCTGCCAATGACAAACCACTTCATGCCAATGACACGCAAAGTAAAAGTTTGAGGATCAGAAGAATAAACCAAAAGATATTTAACCTTGTAGAGAAGAAAGTTCCAGAAGCTAATCAAATTTCAATGTCATCCGAGAATCACGAGACAACTGCATCCAAGCCCATCAATCAAAAGACAGCAGCCAAGAAAGATTCATCGCGCAACAAAGTGCAGGACGCAAGGACTAGCACTATATCAGGTTAG

>S_melongena_3

ATGCCGGCGTTGGAGGAATTCAGGCAGATGGGAGAGGTAATTGGGAGTATCAAAGCTCTAATGGTATTCCAAGATGAGATACAAATAAATCCAAGACAATGTTGCTTGTTGGTGGATATGCTCAAATGTGCCTACAAGACAATTGCAGAAATGATGAAACAGAACTTGAGATTTGAAGAGAAGAATATGAAATGGAAAATTCTTGAGAACCCCTTAAGAGAGCTTCTCAGGGTGTTCAAAGAAGCAGAACAGTACATCAAGCAATCCCTGGAACATAAGGATTTTTGGGCAAAAGCCATACTTCTCTATAAGAATACAGATTGTGTTGAATTTCACATCCACAATTTGCTCTCTTGCGTGCCAATTGTCATTGAGGCCATAGAAATAGCAGGAGAGATTTCAGGTGGTGACCATGACGAGATACAAAAGAAAAGATTCATTTACTCGATGAAGTATCAGAAAGAGTGTAAGGACCCAAGAATCTTCCAATGGAAATTCGGAGAACAGTACATGGTTTCCCAGAAGTTCTGCGAAAGGGTAGGCTTAGTTTGGAATGAAGATAAGTGGATTTTGAAGAACAAAATTCGAGAGAAGAAGGATTCAGGTTCATGTACGTTGACAAAGCATGAGAAGCGACTTGCTGATCTCCTCTTGAAGAACTTAAATGACATAGAGACAGAGAGTGATCATAAGCTTTCACCTAGCTCAGTTCTTGTAAATTCTAAGGATTACCATATAAGAAGGCGATTAGGTAGTGGAAGCCAATACAAGGAGATCCAATGGTTGGGGGAGACCTTCTGTTTAAGGCACTTGTTTGGGGATATAAAGCCCTTAATTCCAGATATTTCTCAAGAACTGCATCTCTCGCATCCAAATATAATGCACATCTCCTGTGGTTTTACTGACGAAGAAAAGAGAGAATGTTTCTTAATCATAGAACTTATGAACAAAGATTTATCTAGTTACATCAAAGAGATCTGTGGACCAAGAAAGCGCGTACCATTTTCTCTTCCAGTTGCAGTTGATTTACTCCTTCAGATTGCAAGAGGCATGGAATACCTGCACTCGAAGAAAATCTATCATGGTGAATTGAGTCCTTCCAACATCCTTATCAAGGCTAGGAACGTATCGACAGAAGGGTATTTACATGCAAAGGTTTGTGGATTTGGCTCATCATCTTCTATCAACCTTCCTCAGAAAGCTAATGTGAATCAAAATAATGGGACACTCCCATTCATTTGGTTTGCCCCAGAAGTCCTAGCTGAACAAGAGCAGTCAGGGAATGGAGGAAACATCAAGTATACTGAGAAATCTGATGTGTACAGTTTTGGAATGATATGTTTTGAGGTTTTAACAGGGAAAGTTCCATTTGAAGATAGCCATCTACAAGGAGACAAAATGAGTAGAAATATAAGGGCAGGAGAGAGGCCATTATTCCCATTTCACTCACCAAAATATGTGACTAGCTTGACAAAGAGGTGTTGGCATCCTGATCCATATCAACGGCCAAGTTTTTCATCCATCTGCAGGGTTCTCCGCTATGTGAAGAGATTCTTGGTGATGAATCCTGAACACAGCCAACAAGACTCACCATTGCCACCAGTCGACTATTGCGAGATTGAAGCTGCGATCTTAAGAAGCTTCCCCTTCCTAGGAAATTCTGAAGCTGATCCTCTACCAGTCACACACATACCTTCCCACATGTTTGCATACAGGGTGACCGAGAAAGAAAAGTCAAGCACAATTCACAGAGAAATAAATTCTGAATCAGGAAGTGATGGAACTTCAGCATGTGGGGATGATTTTGTAACGGCAGATGATGCACTTCCATCACCAACCGATAGGAAGAATACTGCATCTCCCGATAATTTAACGAAAAGACTTTCAATTAAGAAACCTGCAGATATCAAAGTCAGCAAGCAACCAGGAACACCAAGAGGACGGACGGTAAGACCTCCAAGCATACGTACTGTAAGGCAGAATTCTGAAAGTCAGTTAATGATGATGAACAGCCCAAGAACAAGAAGATCATCTGGTCATACATCAGATTCAGATCTTCCTTAA

>E_salsugineum_1

ATGTCTAATGAAATGAGGTCTCTCTACAGAGTGTTCAAAGAAGGAGAATTGTATGTCAAGCATTGTATGGACAGTACCGATTGGTGGGGCAAAGTCATAAACCTTCATCAGAACAAAGACTCTGTTGAGTTTCACATACACAACCTGTTCTGTTATTTCCCGGCGGTTGTTGAAGCCATCGAGGCAGCCGGTGAGATTTCTGGGCTTGACCCATCAGAGATGGAGAGAAAGAGAGTTGTCTTCTCAAGAAAGTATGACAGAGAGTGGAATAATTCTAAGCTTTTCCAATGGAGGTTTGGGAAACAGTATCTGGTACCTAGAGACATTTGCAGCCGGTTTGAGCATTCTTGGAGAAAAGATAGATGGAATCTTGTGGAGGCATTGCAAGAGAAGAGGAAATCAGACAGCGATGGTGTTGGAAAGACAGAGAATCGTCTAGCTGATCTGCTTCTGAAGAAACTAACCGGTTTGGAGCAGTTTAACGGGAAGCTGTTCTCGAGCTCGATCTTGCTTGGCTCAAAGGATTACCAGGTGAGAAAGTGGTTAGTTGCAGATGGACAGTACAAGGACATTCAATGCGATCTTGAGCCTCTAAGCTCTGAGATCTCTTCTCTGTTGGCTCTTTTTCATTCCAACATACTTCAGTATCTCTGCGGATTATATGATGAAGAGATGAAAGAATGCTTCTTGGTTATGGAGTTGATGCACAAATATCTGCAGAGCTACATGAAGGAGAATTGTGGACCAAGAAGAAGATATCTCTTCTCGGTTTCCGTCTTGAGAAGAAGTCACACAGATGGTTATTTCCATGCCAAAATCTCTGGATTCGGTTTATCTTCTGTCAAGTCTTGTTCCTCTAGTAGTCGACCAAACACTCCTGATCCTGTGATCTGGTACGCACCTGAGGTTCTAGCAGAGATGGAACAACAAGGTAAAACTGATCCGAAATCGAAGCTGACACACAAGGCTGATGTGTATAGCTTTGCAATGGTGTGTTTTGAGCTTATAACAGGTAAAGTCCCTTTTGAAGATAGTCACCTTCAAGGTGAGAAAATGGCTATCAACATTAGGATGGGAGAGAGACCACTGTTCTCTTTCCCTTCACCGAAATACTTGGTTAGTCTGATCAAACGGTGTTGGCACACAGAACCGAGCCAGCGTCCAAACTTCTCTTCGATTTGCAGGATACTACGCTACATCAAGAAGTTTCTTGTTGTGAATCAAGATCACGGTCACCCTCAGATGCAAACTCTGCTTGTTGATTGTTGGGACTTGGAAGCAAGGTTCTTGAGAAAGTTTCCAGGTGATCACTCATCAGGATGTCACGCAGCTTCCGTGACTCAGATCCCTTTCCAGCTATACTCTTACAGGGTTTTCGAAAGAGAGAAGATGAATCCAAAAAAACTCCAAAGAAACCGCGTTTCAGTGGTTGAAGATCCACCTATTGCCATGATGATAAGAGATACAAGATCCTTGTGTCTAGATACAATATCTGAATACTCAAATACAAGATCAGTCTACTCAGAAGCTCCCCATAAAAAGGTGTCAGCACTAAAGAAAACTCCAAGCTTAGGTTCAGGAAAGTTGAGATCTACAGGGAACTCGCCGGTAAAAGCAAGATCATCACCAAAAGCATCACCGTTGAGTCCATTTGGAAGAAGCATTAAAGCCAGGAAAGATAATCGATTGCCTTTGAGTCCAATGAGTCCTCTAAGCCCAGGGAGACATAGACAACATACTGGTCATGCTTCATACTCTGAGCTTACTTAG

>E_salsugineum_2

ATGGAGCAATTCAGGCAAATCGGGGAGGTTCTAGGAAGCTTAAACGCGCTTATGGTATTACAAGACGACATCTTGATCAACCAAAGACAATGCTGTTTACTGTTGGATATCTTCAGTTTGGGTTTCAACACCGTAGCCGAAGAGATCAGACAAAACTTGAAGCTTGAAGAGAAGCATACCAAATGGAGAGCCCTTGAACAGCCTCTAAGGGAGCTCTATAGAGTGTTCAAAGAAGGAGAAAGGTATGTCCGGAACTGTATGTCCAACAAAGATTGGTGGGGCAAAGTAATCAACTTTCATCAGAACAAAGATTGTGTTGAGTTTCACATACACAACTTGTTCTGTTACTTTCCGGCTGTCATCGAAGCCATCGAGACAGCCGGAGAGATTTCGGGTCTCGACCCTTCTGAGATGCAGAGGAGGAGAGTTGTGTTTTCGAGAAAGTATGACAGAGAATGGAATGATCCGAAGCTGTTTCAATGGAGGTTTGGGAAACAGTATTTGGTTCCAAAGGATATTTGCAGCCGGTTTGAGCATTCATGGAGAGAAGATAGGTGGAATCTAGTGGAGGCATTGCAAGAGAAGAGAAAGTCCAAGAGCGATGATATTGGCAAGACAGAGAAGCGTTTAGCTGATTTTCTGTTGAAGAAGCTGACCGGATTAGAACAGTTTAACGGTAAGCTCTTCCCGAGTTCGATACTTGTTGGTTCTAAAGATTACCAAGTGAGGAGGAGATTAGGTGGTGGTGGTCAGTACAAGGAGATTCAATGGTTAGGGGATAGTTTCGTGTTGAGACATTTCTTCGGTGATCTTGAGCCGTTAGCTGAAGAGATCTCTTCTCTGTTATCGCTTTGTCACTCGAATATACTTCAGTACTTATGTGGATTCTATGATGAAGAAAGGAAAGAGATTTCTCTGGTTATGGAGTTGATGCACAAGGACTTGAAAAGCTACATGAAGGAGAATTGTGGACCAAGAAGAAGATATCTCTTCTCTGTTCCCGTTGTGATCGATATAATGCTCCAAATCGCTCGAGGGATGGAATATCTTCATTCGAATGAGATCTTCCATGGAGACTTGAATCCAATGAACATTCTTTTGAAAGAGAGAAGTCACACCGAAGGTTACTTCCACGCCAAGATCTCAGGTTTCGGTTTGATCTCGGTCAAAACTCAGTCTTTTACTCGGGCTTCCTCAAGACCAACCACTCCTGATCCTGTGATTTGGTATGCACCTGAAGTTCTAGCAGAGATGGACCAAGATCTCAAAGGCAGAGCTCCGAGATCAAAGTTTACACATAAAGCTGATGTTTATAGCTTTGCTATGGTGTGTTTTGAGCTCATAACCGGCAAAGTTCCGTTTGAGGATAGTCATCTTCAAGGAGATAAGATGGCTAAGAACATTAGAAATGGAGAGAGACCTCTCTTCCCATTCCCTTCGCCGAAATATCTTGTTAGTTTAATCAAACGGTGTTGGCATACAGAGCCGAGCCAGCGTCCGACTTTCTCTTCGATTTGCAGGATCCTCCGCTACATCAAGAAGTTCCTTGTTGTGAATCCGGATCACGGTCACATTCAAATCCAAAATCCACTTGTTGATTGTTGGGACTTGGAAGCAAGATTCTTGAGAAAGTTCTCACTAGAGTCAGGGTCTCACGCGGAGTCCGTGACACAGATACCGTTCCAGCTTTACTCATACAGGATTGCAGAGAAAGAGAAGATGAGTCCGGAGATTAACAAAGAAGAGAATTCAGAGACAGGTGAGTCTGCTTCAGAGAGCGTTTCAGTGGTTGAAGATCCACCAACAATGCCAATGTACACAAAGTCGTTGTGCTTAGATGCAATATCTGAATACTCAGATACAAGATCAGTTTACTCAGAAGCTCCCATGAAAAAGATCTCAGCTTTGAAGAAAAGTGGCGACATGGCAAAGCTCAGAAGGAATTCAAGCGCAGGTTTAAGGTCTCCAGGATCATCGCCTATAAAGCCAAGATCAGCACCGAAAGTGTCATCGCCGCTAAGTCCATTTGGAAGAAACAGCAAAGCAAGGAAGGATACTAGATTGCCTTTGAGCCCTATGAGTCCGTTGAGCCATGGAAGACGTAGACATCTCTCTGGTCCTGCTTCAGACTCTGAGCTAACATAG

>E_salsugineum_3

ATGGAGCAATTCAGACAAATCGGCGAGGTTTTGGGAAGTCTAAATGCACTAATGGTATTACAAGATGATATATTGATCAACCAAAGACAATGTTGCTTGTTGTTAGAGATCTTCAGCTTAGCTTTCAACACTGTGGCAGAAGAGATCAGACAGAATCTGAAGCTTGAAGAGAAGCACACTAAATGGAGAGCCCTTGAACAGCCTCTAAGGGAGCTCTACAGAGTGTTCAAAGAAGGAGAATTGTATGTCAAGCATTGTATGGACAGTAGCGATTGGTGGGGCAAAGTCATAAACCTTCATCAGAACAAAGAATCTGTTGAGTTTCACATACACAACCTCTTCTGTTATTTCCCGGCGGTTGTTGAAGCCATCGAGGCAGCCAGTGAGATTTCTGGTCTTGACCCATCTGAGATGGAGAGAAGGAGAGTTGTGATCTCAAGAAAGTATGACAGAGAGTGGAATGATCCTAAGCTTTTCCAATGGAGGTTTGGGAAACAGTATCTGGTACCTAGGGACATTTGCAGCCGGTTTGAGCATTCTTGGAGAGAAGATAGATGGAATCTTGTGGAGGCATTGCAAGAGAAGAGGAAATCTGACAGCGACGATGTTGGAAAGACCGAGAAGCGTCTAGCTGATCTGCTTCTTAAGAAACTAACCGGTTTGGAACAGTTTAACGGGAAGCTGTTCCCGAGCTCGATCTTGCTTGGCTCCAAAGATTACCAGGTGAGAAAGCGGTTAGTTGCAGATGGACAGTACAAGGAGATTCAATGGTTAGGTGATAGTTTTGCGGTGAGGCATTTTTTCAGCGATCTTGAGCCTCTAAGCTCTGAGATCTCTTCTCTTTTGTCTCTTTGTCATTCCAACATACTTCAGTACCTCTGCGGATTCTATGATGAAGAGAGGAAAGAATGCTTCTTGGTTATGGAGTTGATGCACAAAGATCTGCAGAGCTACATGAAGGAGAATTGTGGACCAAGAAGAAGATATCTGTTCTCGGTTTCCGTAGTGGTTGATATCATGCTGCAGATCGCAAGGGGAATGGAATATCTTCATGGAAACGATATCTTCCATGGAGATTTAAACCCCATGAACGTTCTTTTGAAAGAGAGGAGTCACACAGATGGTTATTTCCATGCAAAAATCTCTGGATTCGGTTTATCTTCTGTCAAGTCTTGCTCCTCTAGTACACGACCAAACACTCCTGATCCTGTGATCTGGTACGCACCTGAGGTTCTAGCAGAGATGGAAACAGGTAAAAATGATCAGAAATCGAAGTTGACACACAAGGCTGATGTGTATAGCTTTGCAATGGTGTGTTTTGAGCTTATAACAGGTAAAGTCCCCTTTGAAGATAGTCACCTTCAAGGTGAGAAAATGGCTATCAACATTAGGATGGGAGAGAGACCACTTTTCCCTTTCCCTTCACCGAAATACCTGGTTAGTCTGATCAAACGGTGTTGGCACTCAGAACCGAGCCAGCGTCCAAACTTCTCTTCGGTTTGCAGGATACTACGCTACATCAAGAAGTTTCTTGTTGTGAATCAAGATCATGGTCACCCTCAGATGCAAACTCCGCTTGTTGATTGTTGGGACTTGGAAGCAAAGTTCTTGAGAAAGTTTCCAGGTGATCACTCATCAGGGTGTCACATAGCATCCGTGACTCAGATCCCTTTCCAGCTATACTCTTACAGAGTTTCAGAAAGAGAGAAGATGAATCCAAAAAACTCAAAAGAAAACGGTAACTCAGAGGCAAGCGAATCTTTAGAAAGCGTTTCAGTGGTTGAAGATCCACCTAGTGCCATGATGATAAGAGATACAAAATCTTTGTGTCTAGATACAATATCTGAATACTCAGATACAAGATCAGTCTACTCAGAAGCTCCCATTAAAAAGGTGTCAACATTAAAGAAAAGCGGCGAATTGGTAAAGCTAAAAAAACCTCCAAGCTTAGGTTCAGGAAAGTTGAGATCTACAGGGACATCGCCGGTAAAAGCAAGATCATCACCAAAAGCATCACCGTTGAGTCCATTTGGAAGAAGCATTAAAGCGAGGAAAGATAATCGATTGCCTTTGAGTCCAATGAGTCCTCTAAGCCCAGGGAGACTTAGACAACATACTGGTCATGCTTCAGACTCTGAGCTTACTTAG

>E_salsugineum_4

ATGGAGCAGTTTCGAGAGATAGGAGAGGTATTAGGAAGCATAAAATCGTTTATGGTGTTCAAGGACGACATTCAAATCAACCAGCGTCAATGCACTCTTTTACTCGACCTCTTTACCGCAGCCTACGATTCGATTTCCGAATCTATGCGATCGAATCTACGCTTCTCAGAGAAGAGCACTAAATGGAAGATTCTAGAACAACCCTTGAGGGAGCTTCTATGGGTGGTACGAGAAGGGGAGGCTTACGTTCGAATGTCTTTAGAGCCTAAACTCGGGTTTTGGGCTAAGGCTATTGTGTTACAACACAATCGAGATTGCACAGAGCTTCACATACATAACTTGCTCTCTTGTGTACCGACCATCATCGAAGCTGTTGAGATGGCGAGTGAGGTCTCAGGGTGGGATGAAGAAGAGATGAACAAGAAGAGACTGGTGCATTGTAACAAGTATATGAAGCAATGGAATGACCCTCAGATGTTTACCTGGAAGTTCGGGAGAGAGTATTTGGTGACCGCGGATTTCTGCAGTCGGTATGAGAGCGCCTGGAGAGAAGACAGGTGGATTCTGATGAAGGAGCTTGAGGAGAAAAGACGTCCCGGTACAAGCAAACACGACAGGAAGATGGCTGATTTTCTTTTGAAGAATCTAGGAGATGGAACCGAGAGTCCCAAGCTATTTCCGTCTTCTATTCTGGTTAGCAATAAAGATTACCAAGTGAAAAAGAGATTAGGGAATGGAAGTCAGTATAAGGAGATTACCTGGTTAGGCGAGAGCTTTGCGCTTAGGCACTTCTTTGGGGATATCGATGCTTTGCTTCCTCAAGTTACTCCGTTGCTCTCTCTTTCCCACCCAAATATTGTATATTACCTCTGCGGATTCGCGGATGAGGAGAAGAAAGAGTGTTTCTTGGTCATGGAACTGATGAGTAAAACCCTCGGGACGCACATCAAAGAGGTATGTGGTCCGAGGAAGAAGAACACTCTCTCCCTCCCGGTCGCGGTTGATCTGATGCTTCAGATAGCCCGGGGTATGGAATATCTTCACTCCAAGAAAATATACCACGGAGAGTTGAACCCTTCTAACATTCTTGTCCAACCAAGAAGAAATAACCAATCCGGAGATGGGTATCTGCAGGGGAAGATTTCTGGGTTCGGATTGAATTCTGTCAAGGGTTTCTCTAGTAAAAATGCTTCTTCGACGGGTCAGAGTGAGAGTTTTCCATTCATATGGTATTCCCCGGAATTGCTAGACGAGCAAGAACAGAGCGGAACTGCAGGAAGCATCAAGTATACTGAAAAATCTGATGTGTATAGTTTTGGGATGGTCTGCTTTGAGCTTCTAACAGGTAAAGTACCATTTGAGGATAGCCACCTGCAAGGAGATAAGATGAGTAGAAACATTAGGGCCGGGGAGAGACCGCTTTTCCCGTTCCATTCACCTAAATTCATAACAAACCTGACAAAAAGATGCTGGCATGCTGACCCTAATCAGCGTCCAACTTTCTCGTCGATAAGCAGAATTCTACGATACATCAAGCGGTTTCTAGCTTTGAACCCGGAATGTCACAGTATAAGCCAACAAGACCCGCCAGTTTCTCCTCCTGTGGATTACTGCGAGATCGAGACTAAGCTGTTGCAGAAGCTTTCATGGGAATCCACAGAGTTACATCAAGTGTCACAGGTTCCGTTTCAAATGTTTACATATAGAGTGGTGGAACGGGCAAAGACTTGTAAGAAAAATAATCGCCGAGACACATCTGAATCGGGCAGTGAATGGGCTTCGTGTAGCGAGGATGAGGGAACCCCCAGAGGAGGACGATCAAGGCATCCCCCACTGAGCCCTTGTGGGCAAAGCATGAGAACGCACTCTGAGAGTCAACTGATATTGATGAGCCCAAAGATACGGCGATCAAACTCTGGTCATGTCTCTGACTCTGAGCTTTCCTAG

>F_vesca_1

ATGGAACAATTCCGGCATATTGGAGAGGTTTTGGGTAGTATTAAGGCCCTGATGGTCTTACAAGATGGTATCAAGATCAATCAAGGGCAGTGCTGTTTATTGCTTGATATTTTCACGTCGGCGTTTGAGACGATAGGGGAGGAGATCAGGATGAACCTGAAACTGGAGGAGAAAAAGACCAAATGGAAGGGTCTAGATCAGCCTTTGAGGGAGCTCTACACAGTTTTCAGAGAAGGCGAGATTTACATCAGGAACTGCATGGATACCAAAGACTGGTGGGGCAAAGCAATTGCCCTCTATCAGAACAAGGACTGTGTCGAATTTCACATACACAACTTGTTCTGCTACTTCCCGGCTGTGATTGAGGCGATTGAGAACGCTGGAGAGATTGCGGGGCTCGATCAGGATGAGATGAAGAAGAAGAGGATTGTGCTTAAGAGGAAGTATGATACGGAGTGGAATGATCCGACACTGTTCCAGTGGAGATTTGGGAAGCAGTATTTGGTGCCGAAGACGATGTGCAAAAGGTTGGAGAGTGCTTACAGGGAAGATAGATGGAGGCTCGTTGAAGCTCTAAATGAGAAGAAAATTGCAGGTGGTTTGACAAAGAATGAGCAGCAGCTTGGAGACTTGCTGCTCAAGAAACTACACGGCGCAGAGTTTTCAAGTGGGAAACTGTTTCCGAGCTCAATATTAGTAGGAGCTGAGGAGTACAACATCAAGCGGCGGTTAGGAAATGGACGGCAGTACAAGGAAATCCACTGGTTGGGGCACAACTTTGCCATGAGACACTTCTTTGGGGAACTTGAACCTATGAGGTCTGAGATTTCGACTCTCCTCTCACTCTCCCACCCCAATGTACTGCAATACCTTTGCGGCTTTTATGACGAAGATAAGAAGGAATGCTTTCTTGTTATGGAGTTGATGAGTAAGGATCTCCGCTGCTATATGAAGGAGAATTGCGGCGCAAGAAGGCAGATCTTGTTCTCTCTCCCAGTAGTTGTTGATATCATGCTTCAGATTGCAAGAGGCATGGAGTACCTCCACTCCAGGAAGATCTACCATGGAGACTTGAACCCCTTCAATATCTTTCTCAAGGCAAGGAGCTCCACAGAAGGTTTCTTTCAAGCAAAAGTCTCAGGTTTTGGTTTGTCATCTTTACAAAACAAACCTACCTATCGAAACTCACAGCAACAACAGCAGCAGAAAAATGAAATTGACCCTCTGATTTGGTGTGCCCCGGAAGTCCTAGCTGAGCAAGAACAAACAGGAAATACCAATGTCCGCTCCAAATTCACAGAGAACGCAGATGTATACAGCTTTGGGATGCTTTGCTTTGAGCTCTTGACTGGGAAGATTCCTTTCGAAGATTCACATCTCCAAGGGGACAAGATGAGTCAGACTATAAGAGCAGGAGGGAGGCCTCTATTTCCCTTCCCTTCACCAAAGTACCTTGTAAATCTAACCAAGAGGTGCTGGCAGACTGACCCTTCTCAGCGCCTGAGTTTCGCATCCATTTGCAGGATCTTGCGCTACATCAAGAAATTCCTATCCATGAATCCTGATGTTGATCAGCCAATACTGCAGTCCCCTCCCACGGACTATTGTGAAATAGAGGCATGGTTCCAGAAGCAGTATAAGGCAGGAGGCCATGCTGATGCAGCCTCCGTATCACAAATTCCATTCCAAATGTTTTCTTATAGACTTGGAGAGAAGGAGAAGACTCCAGGCCTGATCAGGGTCAAGAGTTTGGATTCTGTGACTGAAACAGCTTCGATGTGTAAGGCTGAAGCAGAAGCCAATATTTCCATGAGTGACAGTCTCACCGTTATAGATGATACAAGGTCTTTCTGCTCTGATGCAAGGTCACTTTATGATGTGAGATCAGTTTTCTCCGAGGCGCCAACCAAGAGAAGTCTAGCTGCAAAGAAGCGTTCAGAAGTGAGGGGTAGAAAAGCCTCAGGTTCATCAGATGCACAAACAACAGCAAGAATTTCATTGCCTAAACTTCCAACACCAAAGCTCTCATCACCAAGACTCTATTCCCCGAAACTTCCATCAGCAAAACCCCAACCCTCGAAAGCATCCTATGAATATGAATATGAGCCCGAGGCATCCCATGACTCTGAATATGAATACGAGTCCGATGCCTATGAATACGACTCGAAGCAACGTTCTCCCTTGCAGAAACATTTTGGTGTGGAAAACTTCAATCAACATAAGCTTGATATGAAAAATCCTTCTATTTCTACTCCAGTAAAAGAAAAGCTGTCTTCTGCTTTGTATCTCCAAATCAGCACCATCAGCCAGGCGTTGATATTTGTTACTCGGTCCAGGGGATGGTCATTCACAGAAAGACCAGGGCTTTTGCTTGTAGTAGCCTTCATCATTGCTCAACTGACTCCATTTACTAGCAAAAAGGATTTTGGGAAGAAATCTCGTGAGGTTGCATGGGCAGCTGAACAACGAACACTACATGGACTGCAATCTACTGAAGGAAAATTGTTTGCAGAGAAGAGCGCTTTTAGGGACATTAATCTCATGGCAGAAGAAGCTAAAGGGCACACAAAGATTGCAAGGTTGAGGGAGCTTCACACTCTCAAAGGAAAGGTGGAATCATTTGCAAAGCTGAAGGGTTTAGATATTGATGCCATGAACCAAAACTACACCATCTAA

>F_vesca_2

ATGGAGCCATTCCGAAGGCTTGGGGAGGCTTTAGGAAGTATTAAGTCTTTGATGGTGTTCCGAGACAATATCCATATCAATTACCGGCAATGCCTTTTGCTGCTCGACATCTTTGGCTCAGCGTTTGACTTAATAGCCGAGGAGATGAAACACAACCTGAGATTTGATGAGAAGCACACTAAATGGAAAGTTCTTGAGCAGCCATTGAGAGACATGCACAGGATATTTAAAGAAGGAGAGCTCTACATTAAGCATTGCTTGGACAAAAAGGAATGGTGGGCGAAAGCCATCTCACTCTATCAGAACACCGATTGTGTTGAGTTTCACATCCACAACTTGCTTTCCTGCATCCCAATTGTCATTGAAGCAATAGAAATTGCAGGGGAAGTGTCAGGGTCGGATTTGAGTGAGATGCAAAGGAGGAAAAGCGTGTATGCGGAAAAGTACAAAGAAACATATAGAGATTGGAAGCTTTTCCAATGGAGATTTGGGAAGCAGTACCTGATCACTCAAGATTTCTGCAATAGATTTGACACAGTATGGAAGGAAGATAGATGGACTCTTCTGAATCAAATCAGAGAGAGAAAACTGAGCTCGGGTTCAACAAAGTACGAACGACGCCTCACAGATCAACTTCTTAAAGCCCTAGATGGGCCAGAGCCTTGGAATGGGAAGCTCTTGCCTAGCTCAATCCTGGTGGGATCTACGGAGTACCAAGTGAGGCGAAGACTAGGCAGTGGAAGTCAGTACAAAGAGATCCTATGGTTGGGTGAAAGCTTTGCCTTGAGGCAGTTCAATGGAGAAATTGAACCCTTGCTGCCAGAGATCTCTTCATTGTTATCACTTTCCCACCCTAACCTTGTTCACTGCCTTTGTGGGTTTACAGATGAGGAAAAGAAAGAATGTTTCCTCATTATGGAGCTGATGAGTAGAGACCTTGGAAGCTATATCAAGGAGATCTGTGGCCCGAAGAAGCGAATTCCATTTACTCTTCATGTTGCTATTGATCTGATGCTTCAAATTGCAAGAGGAATGGAATATCTCCACTCGAAGAAAATCTACCACGGAGAATTGAATACTTGTAACATACTGGTTAAAGCAAGAGGGATATCCACAGAAGGTTTCTTGCAAGCAAAGATTTCAGATTTTGGTCTAACTTCTGTTAGAAACTCCACCCAGAAAAACCCTTCAAATCAGAATGGAAGCCCACCTGTTATATGGTATTCTCCTGAAGTTCTGGAAGCGCAAGAACATAGTGAAAGTAGTGAGAAAAAGTACACAGAAAAGTCTGATGTGTATAGCTTTGGGATGATTTGCTTTCAGCTTCTAACTGGGAAAGTGCCTTTTGAGGATAGTCATCTTCAAGGGGACAAAATGAGCCGAAACATTAGAGCAGGGGAGAGGCCATTGTTCCCCTTCTACTCTCTAAGATATGTCACCAACCTCACAAAGAGATGTTGGCATGCGGAACCAAATCAAAGGCCAAGCTTCTCATCCATCTGTAGGATTCTTCGCTATGTTAAAAGGTTCCTTGCCATGAATCCTGACTATGACAGCCAGCAAGACCCACCAGTGCCTATAGTAGATTACTGTGACATCGAGTCAGGGGTTCTGAGGAAGTTCCCTTCTTGGGGGAGCTCTGATCAACTAGCATTATCACAGATCCCATTTCATATGCTTGTTTATAGGATAGCGGAAAAAGACAGAAACTCAGGACTGAAAGATACTTCAGAATCAGGAAGCGACGCTTCGTTATGTGGGGATGGAATTGTCACCCCACCAGACGATCCATTCCCAGCTACACCTGAAAGGAAGTGTTCACCTGATACTATGCACAAGAAACTTCCATTGTTGAGGAGATCTTCAGACGTGAAACTCAACAGACAACCAGGAACACCAAGAGGACGGTCTGCCAGACCTCCACAGGGAACGACCCCTCCTTATCGTAGTTTTAGCATGAGAACAAGTTCAGAAAGCCTGCTAACGATGTCTCCAAAAATGCGGAGGACATCAATGGCTTCAGATATGGGGTCGAGTCCAAGAATACGGAGAACCAAATCCGGTCATGCCTCAGACTCTGAGCTCTCCTAG

>G_raimondii _1

ATGGAGCAATTTCGGCAGATAGGGGAGGCTTTAGGAAGTTTGAAGGCATTAATGGTTTTTCGTGACAACATCCAAATTAACCAGAGGCAATGCATTTTGTTGCTAGATATTTTTTATTCTGCATACAAATCGATAGCAGATGAAATGAAAGATAACCTGAAATTCGAAGAGAGGAACTTGAAATGGAAAGTTCTTGAAATGCCATTGAGAGAGCTCCACAGGATTTTCAAAGAAGGGGAAGCCTACATCAAGGAAAGCTTGGAATCTAAAGACTGGTGGGTTAAAGCCATAACTCTCTATCAGAATACTGACTGCGTCGAGTTGCATATCAGTAACTTGCTTTCCTGCATTCCGGTTGTCATTGAAGCAATCGAATCTGCAGCCCAACTGTCCGGTTGGGAACAAGATGAAATGCAAAAGAAAAAACGGGTGTACTCCAATAAGTACCATAAAGAATGGATTGATCCTGAACTTTTCCAATGGAAATTTGCAAAGCAGTATCTTGTTACTCAAGATTTTTGCAATAGGATTGACAATGTCTGGAAAGAAGATAGATGGATCCTTAAAAACAAAATCCAAGAAAAGGAAAACTCAAGGTTGAGAAAGCAGGAGAGAAAACTTGCAGATTTGCTTTTGAGAAACTTAGATAGTTCAAAATCTTTAAAAGATAAGCTTTTACCGAGTTCGATTCTATTGGGGTGGAAGGACTATCAGGTTAGACGGCGGCTCGGTAATGGGAGTCAATACAAGGAGGTTTACTGGTTAGGTGAAAGCTTTATTCTGAGACACATAATTGGAGATGTAGAAGCAGTAGCTTCTGATATTTCTTCATTATTATCTCTTTCCCACCCAAACATACTGCATTTCCTTTGTGGATTCACTGATGAAGAAAAGAAGGAATGCCTTTTGGTCATGGAACTAGCGCATAAAAGCCTCTGCGACTGCATAAAAGAAACTTGCGGCCCAAGAAAGCGAACATCGTTTTGTCTTCCGGTCACTGTTGATCTAATGCTTCAGATTGCAAGGGGAATGGAATATCTACACTCAAATAAAATCTACCATGGTGACTTAAATCCTTCCAGCATTTATGTTAAGCTGAGAGGAACATCTTCGGAAGGGTACATGCAGGTGAAAGTTTCAGGATTTGGCTTGTCTTCAATACCTCAGAGGGGAGCAGCGAACCAGAATGAAACACAGTCATTCATTTGGCATGCACCAGAGGTTCTAGAGGAGCAAGAACAGTCAGGAAGTAAAGTGAAATTAAAGTTCACTGGAAAAGCAGATGTGTATAGCTTTGGAATGATTTGCTTTCAGCTTCTTACTGGAAAAGTCCCATTCGAAGATGGCCATCTTCAAGGGGACAAAATGAGCCGAAACATAAGAGCAGGGGAGAGGCCACTGTTTCCTTTCAAGCCACCAAAATCTATAACAAGCTTGATCAAGAGATGTTGGCATGCTGATCCGGATCTAAGGCCTAGCTTCCTATCCATCTGCAGAATTCTTCGGTATATAAAACGGTCACTTCTAATGAACCCTGATTATAACAGTCAGTCGGAGTTACCGCAGCCGCTTGTGGATTACTGTGACATTGATATGAGGCTTCAACGGATATTCCCCACATGGGAAACTCCCAATTCATTATCCACATCACAAATCCCTTTTCAGATGTTTGTTTATAGAGTATTGGAAAAGGACAAAACAGGAGTTCCTCTCAAAGACACTTCGGAATCGGGAAGCGACAGAAATTCGGCTAGTGGCGATGAAAATGTCACTACTGATGAGCTATACACATCAGCAACTGACAGGTCCATGCCTTCACCTGAACCACTGCCTAGGATAAATACTACAATGAAGAAATCTGCAGATATTAGAACAAAACATCCAGTGACACCTAAAGCAAGATCAACAAGACCTCCAATGAATCAGCGTGTCCGCAGTTTCAGAATGAGTTCAGAAAGCCAGCTGCTCCTAGTGAGCCCCAGACTACGGAGATCATCATCCGGCCATGTCTCAGACTCTGAGCTCTCCTAG

>G_raimondii_2

ATGGAGGAATTCAGACAAATTGGTGAGGTTTTGGGAAGTCTAAGAACTCTTATGGTTTTACAAGGCGAGTTTCAAGTCAATCGACGACAATGTTGCTTGTTGTTTGATATATTTTGTTTGGCTTTCAATGTAATAGCCGAAGAAATAAGGTTGAACCTGAAAATCGAAGAAAAGAACATAAAATGGAGTCCCATTGATAACCCTTTGCGAGAGCTTCAAAAGATCTTTAAAGAAGGTGAATTATATGTAAAACAATGTATGGACAAGAAGGACTGGTGGGTTAAGGCAATAAACCTCCATCAAAACAAGGATTCTGTTGAGCTTCACATTCACAATTTGCTTTGCCACTTTCCGGCCTTCATTGAGGCCATTGAGATGGCCGGAGAGATCGCAGGGCTCGACCAGTATGAGATTCAAAGGAGGAGAGTCACACTCACGAGGAAGTACGACGCCGAGTGGAACGATCCGAAGCTCTTCCAGTTTCGATTCGGAAATCAGTACTTGATCCCTCGAGAGATTTGTAGCCGGTTCGAAAGTGCGTGGAGGGAAGATCGATGGAACCTTGTTGAAGCTCTGAGGGAGAAGAAGGCTTCGGAATCGGTAACGAAGAACCAACAACGGCTAGCTAGCTTGTTGATTCGTAAAATTATTGGATCAGAGGCTTCTAACGGTGAACTTTTCCCTAGTTCAATCTTGTATTGCGGGGATTACCAAGTAAGGCGACGATTATGGGGACAATATAAGGAAATACAATGGTTGGGAGATAACTTTGTGTTAAGAAACTTCTTTGGGGATGTTGAACCATTATATTCTGAAATCTCTAAATTACTTTCACTCACTCATCCAAACATACTGCAATACCTTTGTGGGTTTTACGATGAAGAAAAGAAAGAAGTGTTGCTTGTTCTCGAGTTAATGAACAAGGATCTCGGTAGTTACATGAAGGAGAATTACGGCTCGAGAAGGCGGATCTTGTTCCCTCTTCATGTTGTGGTTGATCTTATGCTTCAAATTGCAAGAGGAATGGAATATCTGCATTCAAAGAAGATTTATCATGGAGAGTTGAGCGCTAGTAACATCTTTCTAAAAGCTAGGAACAATGTTGAAGGTTGTTTCCATTTGAAGATCTCGGGTTACGGTTTATCTGATGTAAAGCCTCGTTCCTCCCCGAACTCGTCACCGAGGATGTGCGAGCCAAAACCTTTCATTTGGTATGCTCCAGAAGTTATGATAGAGCAAGAGCAGTCATCCAATGGATCTTCCATTTTGAAGTACTCAGAGAAGGCAGATGTGTACAGCTTTGGGATGCTTTGCTTTGAGTTATTGGCTGGGAAACAACCGTTTGAAGGACATGTAGAGAAGATGAGTAGAAATATCTTGGCAGGGGAGAGGCCTCTTTTTCCCTACACAGTCCCAAAATATCTTGTCAATTTAACGAAAAAGTGTTGGCATACCGATCCGAATCAACGCCCGAGTTTCTCATCAATTTGTCGGATTTTGCGCTACATCAAGAAGTTCCTCGTGATGAACCCTGACCATGACTATGATCAGCCTGAAGTGCAATGTCCTATTGAAGATTATTGTGAGATAGAGACATGGTTTGCAAGAAAGTTCACTGCAAATGAGACTTTCAATCCACTTTCAGTTGCACAAATCCCTTTTCAAATGTTTGCTTATCGGCTTGTCGAGAAAGATCGAACTATTATGAACACGAAGGATAAGAATGGCGAGTTAACCATCGAAGGGGCCTCGACTTGCAGAGATGATATTGTTTCTATAATAGAGGATCCATTGACAGCAACTAGTGACACAAAGTCTGTTGGTTCTGATGTGAAATCTCGTGGTTCAGATACCAAGTCAGTTTATTCGGATGTAAGATCAGTTTACAGTGAGGTTCCGGAAAGGAGAACAATACGGTTTCGCTCGCCCCCAAAGAGATTGGTTTCCACAAAGACTCCAGAGAAGAGAGTTGTAATGACAAAGAAAAACATTAACGTGAAGGCCAAGAAAAGTTCAGGGGCAATAAATGGACAATCTACTCGATCATCAACCTTGAACCGTGTTCACAGTACAGGGGTAATCCGAGAAAATCGATCATCATTCAGTACAGGTTCCTTCAACAGAGGTCGGCAACAAACATCAGGTCATACCTCGGATTAG

>G_raimondii_3

ATGGAGCAATTCAGGCAGATTGGTGAGGTTTTGGGAAGTTTGGGTGCTCTCATGGTTTTACAAGATGATACTCAAATCAATCGACGCCAGTGCTGTTTATTGTTTGATATCTTCAGTTCGGCTTTCAATACAATCGCCGAGGAGATAAAGCTGAACCTCAAACTAGACGAGAAGAACACCAAATGGAACGCCCTCGAACAACCATTCAAAGAGCTTCAAAGGATCTTCAAAGAGGGTGAAGTTTATGTCCGACAAAGTATGGACAAAAGAGACTGGTGGGTTAAAGCAATCAACCTCCACCAGAACAAGGATTGCGTCCATAACCACATCCACAATTTGCTTTCCCACTTCCCCGTTGTCATCGAGGCCATCGAGACGGCAGGAGAGACAGCAGGGCGCGACCGAGGCGAGATGCAAAGGCGGGGAATTGCTCTCAAAATGAAGTACGACAAGGAATGGAATGACCCCAAACTTTTCCAGTTCCGATTTGGGAAGCAATACTTGATCCCTCAAGACATTTGTAGTCGTTTCGAGAGTGCTTGGAGAGAAGATAGATGGAATCTTGTGGAAACTCTAAGGGAGAAGAGCGGTTCGGAATCCACAACCAAAACTCAGCAGCGTCTGGCTGACTTGTTGATCAAGAAAATCATTGGATCAGAGGGTTGTGTTGGCAAACTCTTCCCAAGTTCAATCTTGAATGGAAGGGATTACGTAGTGAGGCGACGAGTTGGGGGACAGTACAAGGAAATCCAGTGGCAGGGTGATAGCTTTGTATTGAGAAACTTCTTCGGGGATGTTGAAACATCAGCCTCTGAAATCTCCACCCTTCTTTCACTTTCACATCCCAATATATTGCAATACCTTTGCGGGTTCTACGATGAGGAAAAAAAAGAAGTATTGCTGGTTCAAGAGCTGATGAACAAGGATCTTACTTATTACATGAGTGGCTCCAAACGGAGGGTCTCGTTTTGTCTTCCTGTCGTGGTTGATCTCATGTTTCAGATTGCAAGAGGAATGGAATATCTTCACTCACAGAAGATATACCATGGAGATTTAAGCCCTTCTAATATCTTTCTCAAAGCTAGAAACAGTACTGGCGATTATTACCAGTTGAAAATCTCGGGTTATGGTTTATCCCCTGTCAAAACTACCACCGGTTCTTCTCTGAAACCGAATGAGACCAAGCCTTGCATTTGGTATGCTCCGGAAGTTTTGCAAGAGCAAGAGCAATGCCTGCCAGGGAATGCCGCCAGCTTTAAGTACACGGAGAAAGCAGATGTTTACAGTTTTGGGATGCTTTGCTTTGAGCTCTTGACTGGGAAAGTGCCATTCGAAGATGGACATCTTCAAGGGGATAAGGTGAGTCGAAATATTAGAGCAGGGGAGAGGCCTTTGTTTCCCTACACTGCGCCTAAATACCTTGCCAACTTAACCAAAAGATGCTGGCACTCCCGTCCGAATCAACGTCCAAGCTTCTCATCGATTTGTCGGATTTTACGATATGTAAAGAAGTTTGTCGTGATGAACCCTGACCATGATGAGCCCGATGCTCGATCACCGGTCTCAGATTATTGTGAGATAGAGTCATGGTTCTTGAAAGCGTTTGCTGCCAATGGGAGTTTCAATTCATTATCAGTGGCACAAATACCGTTTCAAATGTTTGCTTATAGGCTTGCAGAGAAAGACAAAACTATTTTGAACAGCATGGATAAGAATGGAGAGGGAGCAGCCTCAACTTCCAGAGAAGACATGAATTCAACAGTTGATGATCCATTGATAACAGCTAGTGATGCAAAATCTGCGGACTCTGATGGAAAATCGGTGTATTCTGAGATTCAAGAACAGAGATCAATCCATTTGGATTCAACGCCACAAAGGAGATCAGTTTGCAGCCGGATTCCAGAGAAGAAAATTTTACAGATGAAGAGAAACAGTAATGTGAAGGCGGCGAGAAAGAGTACAGGCCATAAAGATGGGATACCAAATGGAAAACCTACACAACCGCCATTACCTCGTGTTCAGAGTGTAAAGAATGTGAGAGTAAGTCGGTTAAAATCAATGACGAGGTCGTTGAGCACTGATCGGCTAAGATTAGCAGCTGGTGATACCTCGGATTAG

>L_usitatissimum_1

ATGGAGCAGTTCAGGCATGTAGGGGAAGTCCTGGGGAGCCTAAAGGCATTAATGGTGTTACAAGACAACATCCCGATCAATCCTCGACAATGCGGGCTGCTGCTCCAAATGTTCACCTCCGCATTCGACGTGATAGCCGAGGAGATCAAGCACAACCTCACGCTTGAAGAGAAGAACACAAAGTGGAAGCCATTGGAGGAGCCGTTGAGGGAGCTTCATAGGGTGTTCAAAGAAGGGGAGCTCTACGTCAGGCATTGCATGGACAACAAAGATTGGTGGGGGAAAGCTCTTTCTCTCCATCAGAACAAGGACTCCGTCGAGTTCCACATCCATAACTTGATCGCCTGTTTCCCCGCTGTGATCGAGGCGATCGAGACTGCCGGCGAGATCTCGGGGCTAGATGAAGCTGACATGAAGAGGAAGAGGGTGATGCTGTCGAAGAAGTACGATAGGAGCTGGTACGATCCGAAGCTGTTCCAATGGAGATTCGGGAGGCAGTATTTGATTCCCAGAGAGATTTGCTGTCAGTTCAAGGCTGCTGTGAGGGAGGATAAATGGCACTTGATTGAGAAGTTGAAGGAGAAGAAAGCTTCGGGGTTGGTCTCGAAGAACGAGCATCGACTAGTGGATTTGCTCTTGAAGAAGCTTAACTCTACTGAGAGGGTTAACGGGAAGTTACCTCCGAGTTCGATCTTGTTTGGAGGGGAGGATTATAAAGTGAGGAGGAGGCTAGGGGAAGGAAGCCAGTACAAAGAGATTCAATGGCTAGGAGAAGTATATGCAATGAGGCATTTGTTTCATGAGATTGAGCCGTTGAGTTCCGAGATTTATGACCTGATGTCGCTTGCTCATCCCAACATTGTTCAGTACTACTGCGGATTCTACGACGAGGATAAAAAGGAGTGTTTCCTTGTGATGGAGTTGATGAGCAAGGATCTATATTCTTACATGAAAGAGAACAGCAGTCCAAGGAAAAGGGTTCTTTTGCCTCTCCCTATTGTTGTGGATATCATGCTTCAGGTTGCTAGAGGAATGGAGTTTCTCCATTCCAGGAAGATCTACCATGGTGATTTGAACATCACTAACATCTTTCTGAAACCAAGGAAATCCACAGAAGGCTACTTCTCGGTTAAAGTTTCGGGCTTTGGTCTGACAAGAGTTGAAAACCAACCTTTGAAGCATGCTTCTCCGAACCCGGATCCAATCGATCCGACTATTTGGTATGCACCTGAAGTTCTGGCAGAACAAGAGAAAGGAGGAAACTTGGCGTGTAAGTACACGGAAAAGGCAGATGTTTACAGCTTCGGAATGCTGAGTTTCGGGCTCCTAACTGGTAAGCTTCCTTTCGAGGATGGGCATCTCCATGGAGATCAAATGATCAAGAACATTAGAGCTGGAGAAAGACCATTGTTTCCCTACCTATCACCAAAGTACCTTGTGAATTTGACTAAAAAGTGCTGGCATACGGATCCAAACTCTCGTCCTAGCTTCACCTCCATTTGCAGGATTCTTCGATACGTAAAGAAGTTCTTGATCATGAATCCAAGCGATGGCCAGCCGGATATGCACTTGCCACCTGTCGACTACACTGACCTTGAATCAGGATTCCTGAAGATGTTCCCTGCAGACACAACCTGTGATTCGAGCTCGGTTTCTCAGATTCCATTTCACATGTTTGCTTACAGACTGGTTGAGAGAGAAAAGACATCATTAGGTGTCAAGTTCAAGCAAAGTGAACCACCCACTGAGGCAGTTCCTAATGGTCCCGACGAGCCACATCATATCCTTGCAACTGAGGATCCAGCTGAAGAGTCCAAAATCGTTCTTTTCGATGCGAAGTCGGTTCTCGTGGCGAGTCCTGAGAAGCAATTCCCATCAGATATGAGGTCAGTTATTTCGGAGCCACCTAATAAAAATGCAAGTCCTGTCAGGAAGAAGCCAGCCAACGTTAAGGGTCTCTATTGGTGCTTTGCAGTAATTGAAGCAGCAAAACCAAAGCTGAAAACGCCAGTGAGGTCACCAGCACCAGTGAGATTATCAGCAACTTGTAGCCCAACTGGCCGCAATTCGAGGGCGAACACCAAAGTCAGTCCGAAAAGAACAACACCATCAATGGCTTCTCAGAATCCAGGGAAAGGGATATTATATAGACAAAGCTCTCCATAG

>L_usitatissimum_2

ATGAAGAGGAAGAGGGTGATGCTGTCGAAGAAGTACGATAGGAGCTGGTACGATCCGAAGCTGTTCCAATGGAGGTTCGGGAGGCAGTATTTGATTCCAAGGGAGATTTGCTGTCAGTTCAAGGCTGCTGTGAGGGAGGATAAATGGCACTTGATTGAGAAGTTGAAGGAGAAGAAAGCTTCGGGGTTGGTCTCGAAGAACGAGCATCGACTAGTGGATTTGCTCTTGAAGAAGCTTAACTCTACTGAGAGGGTTAACGGGAAGTTACCTCCGAGTTCGATCTTGTTTGGAGGGGAGGATTATAAAGTGAGGAGGAGGCTAGGGGAAGGAAGCCAGTACAAAGAGATTCAATGGCTAGGAGAGGTATATGCAATGAGGCATTTGTTTCATGAGATTGAGCCGTTGAGTTCCGAGATTTACGACTTGATGTCGCTTGCTCATCCCAACATTGTTCAGTACTACTGCGGATTCTACGACGAGGATAAAAAGGAGTGTTTCCTTGTGATGGAGTTGATGAGCAAGGATCTGTATTCTTACCTGAAGGAGAACAGCAGTCCAAGGAAAAGGATTCTTTTGCCTCTCCCTATTGTTGTGGATATCATGCTTCAGGTTGCTAGAGGAATGGAGTTTCTCCATTCCAGGAAGATATACCATGGGGATTTGAACATCACTAACATCTTTCTGAAACCAAGGAAATCCACAGAAGGCTACTTCTCGGTTAAAGTTTCTGGATTTGGTCTGACAAGAGTTGAAAACCAACCTCTGAAGCATGCTTCTCCGAACCCGGATCCAATCGATCCGACTATTTGGTATGCACCTGAAGTTCTGGCAGAACAAGAGAAAGGAGGAAACTTGGCGTGTAAGTACACGGAAAAGGCAGATGTTTACAGCTTCGGAATGTTGAGTTTCGGGCTCTTAACTGGTAAGCTTCCTTTCGAGGACGGGCATCTACATGGAGATCAAATGATCAAGAACATTAGAGCTGGAGAAAGACCATTGTTTCCTTACCTATCACCTAAGTACCTAGTGAATTTGACCAAGAAGTGCTGGCATACGGATCCAAACTCTCGTCCTAGCTTCACCTCCATTTGCAGGATTCTTCGATACGTAAAGAAGTTCCTGATCATGAATCAAAGCGATGGCCAGCCGGATATGCACTTGCCACCTGTCGACTACACCGACCTTGAATCAGGATTCCTGAAGATGTTCCCTGCAGACACAACCTGTGATTCGAGCTCGGTTTCTCAGATTCCATTTCACATGTTTGCTTACAGACTGGTTGAGAGAGAAAAGACATCATTAGGTGTCAAGTTCAAGCAAAGTGAGCCACCCAGTGAGGCAGTTCCTAATGGTCACGACGAGCCACATCATATCCTTCCAGCTGAGGATCCAGCTGAAGAGTCCAGAATCGTTCTCTTCGATGCGAAGTCAGTTCGTGTGGCGAGTCCTGAGAAGCAATTCCCATCAGATACGAGGTCAGTTATTTCGGAGCCACCTAACAAAAATGCAAGTCCTGTCAGGAAGAAGCCAGCCAACGTTAAGGGTCTACTGATGCTTGCAGTAATTGAAGCAGCAAAACCAAAGCTGAAAACGCCAGTAAGGTCACCAGCACCAGTAAGATTATCAGCAACTTGTAGCCCCACTGGCCGCAATTCGAGGACGAACACCAAAGTCAGTCCTAAAAGAACAACACCATCAATGGGTTCCCAGAATCCAGGGAAAGGGATATTATATAGACAAAGCTCTCCATAG

>L_usitatissimum_3

ATGGAGCAATTCAGGAGAATGGGGGAGCAGCTCGGCTGCCTGAAGGCATTAATGGTTTTTCGGGACAACATCGAGGTCAATCCGAGGCAATGCAAGCTGTTGCTAGATATCTGCAGCGCTGCATACGACGCGATCTCGGATCAAATCTTACAGAGTTTGAAATTCGAGGAGAGGCATCTGAAATGGGGAGTCCTCGAGCAGCCATTGATAGAGATTTGCAGGATTTTCAAGGAAGTGGAAGGTTACATCAGGCAGTGCCTCGAGACGACGACATCGTTTTGGGCAAAAGCTGTTGTGTTCCATCATAATCGAGACTCAGTGGAGTTTCTCGTCCACGGCCTGATCTCCACGATGCCTGCCGTGATCCAAGCTATTGAGACTGCGGGGGAGTTTGCGGGGAATGATCAGGAGGAGATTCGGATGAAGAGGGTGATCGTGTCCAATAAGTACCACAAGCAGTGGATCGGCGAACGGAAGATGTTCGAGATGAAGTATTCGAGACAGTATTTGGTTTCAAAAGATTTAGTCGCTCGGTTCGAGTCTGCTTGGTTGGAAGACGGATGGATCCTGATGAATAGACTCAGGGTTTTGAACTCCACGAAGCAGGAAGAGAAGCTGGCCGATTTTATGGCGAAGAGGATCGAAGGGAATTTGAAATTTTTACTTCCATGTTCGGTCCTGGTTGGTTCCAGAGATTATCAAGTGAGGCGGAGGCTCGGCGGAGGGAGCCAATTCAAGGAGATCCAATGGCTAGGTGGAGAGAGCTTTGCCGTGAGGCATTTCTTCGGAAACATCGAGCCGATGATTCCTGAGATTGTTCAGCTTTCGTCACTTTGCCATCCGAATGTTCTGCAATTGCTCTGTGGATTCACTGATGAAGACAAGAAAGAGTGTTATTTGGTATCAGAACTGATGAGTTGTAGAGACATTGGAAGCTACATTAAGGAAGCTTGCGGATCAAGAAACAGCAGGAGGCTACCGTTTTCACTCCCTGTTGCCGTCGATTTGATGCTTCAGATCGCACGGGGAATGGAGTATCTTCATTCCAGAAAGGTTTACCATGGAAATTTGAATCCTTCCAATATTTTGATCAGGCCGGGCGCTTCGGGCTCGAGTTTACTTGTTAAAGTGTCAGGATTCGGCTTGTCTTCGTTGCGGAATACGAACAACAACAACGTAGGGAATTCGTCGTCCTCATGTACTTCTTTGCCTTTCATCTGGACCGCTCCAGAAATTCTGGAAGATACGACAGACGATCGTATGTTAACGGAGAAGTCGGATGTGTACAGCTACGCGATGGTATGTTTTGAAATCCTGACAGGAAAAGTCCCATTTGAAGACAGTCATCTCCAGGGAGACAGGATGGCCCGCAATATTCTAGCAGGGGAACGGCCGTTGTTTCCATCTCATACACCGAAATTCGTGGCCAATTTCACGAAAAGATGCTGGCATCCTGACCCAAATTCCCGCCCAGGATTTTCGACCATCTGCAGGGTCCTTCGATACGCGAAGAAGTTCCTCCTTCTCACGAATCTTCCAAACAGTCAGGACACCACTCCCATGGTCGCGCCCCTGCTGGATTACTTGGAGATGGATTCCAAGCTTCGCCGGAGATTCCAGTCCTGGGAGAATCCTGACACCTCCGCTGTGTCGGAGGTGTCGTTTCAAATGTTTGCTTACAAGGTAATGGAGATGGAGAGGATGTCGTCGGCGGGGAAGGACTTTACCACTTCGGAATCGTCAGGAAGCGACCTGAGCGGAGATGAAGCTGCATTTGTGGACGTGATGGATCCGTTCAGGAGTCCTGACAGGAATAGTAGTAGTGCTAGTAGCAGTACCCATAATATGGTACGTTTGTCCTCGTCCCCTGAAGCGACAATGACCGCATCGCCGCCGGTTCTTAAGAGGTCGTCAACGGTTGGTTCCAAGAGAGCGACGTTTGATCCATGGAAGACCTCCCCCAAACATCCAGGGACACCGAGAGGAAGAGCAACGAGGCCGCCGCCGCTGGGACAATGTGTTGGAGGAGGGAGGAGCATGAGAATGAATTCAGAGAGTCAGTTGATGAACATGGCAAGTCCCAAGGTCACAAGGAGAACTACAACCGGCCATGCTTCCGATTCTGAGCTTTGA

>L_usitatissimum_4

ATGGAGCAATTCAGGAGAATGGGGGAGCAGCTCGGCGGCCTGAAGGCATTAATGGTGTTCCGGGACAACATCGAGGTCAATCCGAGGCAATGCAAGCTGTTGCTAGATATCTGCAGCGCTGCATACGACGTGATCGCGGATCAAATCTTACAGAGTTTGAAATTCGAGGAGCGGCATCTGAAATGGCGAGTCCTCGAGCAGCCATTGATAGAGATTTGCAGGATTTTCAAGGAAGTGGAAGGTTACATCAGACAGTGCCTCGAGACGACGACGTCGTTTTGGGCGAAAGCCATTGTGTTCCATCATAATCGAGACTCGGTGGAGTTTCTCGTCCACGGCCTGATCTACACGATGCCTGCCGTGATCGAAGCTATTGAGATTGCGGGGGAGTTTGCGGGGAATGATCGGGAGGAGATTCGGATGAAGAGGGTGATCGTGTCGAATAAGTACCACAAGCAATGGATTGGCGACCGGAAGATGTTCGAGATGAAGTATTCGAGGCAGTATTTGGTTTCAAAAGATTTAGTCGCTCGGTTCGAGTCTGCTTGGTTGGAAGACAGGTGGATCCTGACGAATAGACTCAGGGTTTCGAACTCCACGAAGCAGGAACGGAAGCTGGCCGATTTTATGGCAAAGAGGATCGAAGGGAATTTGAAATTTTTACTTCCATGTTCGGTCCTGGTTGGTTCCAGAGATTATCAAGTGAGGCGGAGGCTTGGCGGAGGGAGCCAATTGAAGGAGATCCAATGGCTAGGAGGAGAGAGCTTTGCCGTGAGGCATTTCTTCGGAAACATCGAGCCGATGATTCCTGAGATTGTTCAGCTTTCGTCGCTCTGCCATCCGAATGTTCTGCAATTGCTCTGTGGATTCACTGATGAAGACAAGAAAGAGTGTTTTTTGGTATCAGAACTGATGAGTTGTAGAGACATTGGAAGCTACATTAAGGAAGCTTGCGGGTCAAAAAACAGCAGGAGGCTACCATTTTCGCTCCCTGTTGCCGTCGATTTGATGCTTCAGATCGCACGGGGAATGGAGTATCTTTTTTCCAGAAAGGTTTACCATGGAAATTTGAATCCTTCCAATATTTTGATCAGGCCGGGCGCTTCGGGCTCGAGTTTACTTGTTAAAGTGTCAGGATTCGGCTTGTCTTCGTTGCGGAATACGAACAACAACAACGTAGGGAATTCGTCGTCCTCATGTACTTCTTTGCCTTTCATCTGGACCGCTCCAGAAATTCTGGAAGATACGACAGACGATCGTATGTTAACGGAGAAGTCGGATGTGTACAGCTACGCGATGGTATGTTTTGAAATCCTGACAGGAAAAGTCCCATTTGAAGACAGTCATCTCCAGGGAGACAGGATGGCCCGCAATATTCTAGCAGGGGAACGGCCGTTGTTTCCATCTCATAAACCGAAATTCGTGGCCAATTTCACGAAAAGATGCTGGCATCCTGACCCAAATTCCCGCCCAGGATTTTCGACCATCTGCAGGGTCCTTCGATACGCGAAGAAGTTCCTCCTTCTCACGAATCTTCCAAACAGTCAGGACACCACTCCCATGGTCGCGCCCCTGCTGGATTACTTGGAGATGGATTCCAAGCTTCGCCGGAGATTCCAGTCCTGGGAGAATCCTGACACCTCCGCTGTGTCGGAGGTGTCGTTTCAAATGTTTGCTTACAAGGTAATGGAGATGGAGAGGATGTCGTCGGCGGGGAAGGACTTTACCACTTCGGAATCGTCAGGAAGCGACCTGAGCGGAGATGAAGCTGCATTTGTGGACGTGATGGATCCGTTCAGGAGTCCTGACAGGAATAGTAGTAGTGCTAGTAGCAGTACCCATAATATGGTACGTTTGTCCTCGTCCCCTGAAGCGACAATGACCGCATCGCCGCCGGTTCTTAAGAGGTCGTCAACGGTTGGTTCCAAGAGAGCGACGTTTGATCCATGGAAGACCTCCCCCAAACATCCAGGGACACCGAGAGGAAGAGCAACGAGGCCGCCGCCGCTGGGACAATGTGTTGGAGGAGGGAGGAGCATGAGAATGAATTCAGAGAGTCAGTTGATGAACATGGCAAGTCCCAAGGTCACAAGGAGAACTACAACCGGCCATGCTTCCGATTCTGAGCTTTGA

>M_acuminata_1

ATGGAGCAATTCCGGCAGATCGGTGAAGTCCTGGGAAGCATCAAGGCGATGATGGCCGTGGCGGTGGAGATGAGAAACCATCTGAGGTTCGAGGAAAAGCTCATCAAGTGGAAGGCCCTGGAGCAACCGCTCAGGGAGCTGCACCGGGTGTTCCGGGAGGGGGAGCAGTACCTGAGGCAGTGCTTGGAGCCCCGGGATTGGTGGGGGAAGGCCATAGCCCTCGGTCAGAACACCGACTGCGTCGACTTCCACCTTCACGACTTGCTGTGGTGCATTCCGGTCGTGATGGAGGCGATCGAGAACGTGGGAGAGATCACCGGAACCGACCCGGAAGACATTTACCGGAAGAAGCTCGTGTTCTCGAAGAAGTACGAGAAGGAGTGGATGGAGCCGAAGCTCTTCCAGCACAAGCTGGGAAGCTTGTACTTGGCTTCCCAAGGCTTGTCCAGCAGAATGGACACATCGTCGGGGGAGGATCGATGGGTTCTCTCGGAAATGATTGCGGAGAAGAGGAGCCAAGGATCGAAGCCCCTGTCGAAGCAAGAGAACCGGCTCGCCGATCTCCTCCTGTGTCCCAAAGGGAAGCTCTTCCCCTGCTCGGTTCTCGTCGGATCCAGCGACTACCAAGTCAGAAGACGATTCGGGTCGGGAAACAACTACAAGGAAGTGCAGTGGATGGGCGAGAGCTTCGCGGTGAAGCATGTCATCGGAGAGATGGAGCCACTGATGCCTGAGATCTCCCTCCTGTCATCTCTTTCGCACACCAACGTGGTGCATTACATGTACTCGTTTGTGGACGAAGAGAAGAAAGAGTGCATGCTGGTGATGGACCTGATGAGCAAGGATCTCTCGAGCTACATCAAGGAGATCTGCTCCACGAGAAGAAAGGTCCCCTTCCCCTTGGTGGTGGCAGTGGACACCATGCTTCAGATTGCAAGAGGAATGGAGTATCTCCACTCCAAGAACATATACCACGGAGATCTGAACCCTTCAAACATACTGGTCAAGACGAGGAGTGCTTCACCGGACGGGCATCTGCACGCCAAAGTTACTGGATTTGGGCTGTCACCGGTGAAGCACTCCAAGCCCACAGCAACCCAAGCAGCTGCCACACAATCATGCATCTGGTACGCCCCGGAGGTGCTATTGGAGCAGGAGAGTAGCAGTGCCAAGTGCACGGAGAAGGCAGACGTGTACAGCTTCGGGATGATCTGCTTCGAGCTGTTGACAGGCAAACTCCCCTTCGAAGACAACCATCTCCAGGGGGACAAGATGAGCAAGAGCATAAGAGGAGGCGAGAGGCCTTCGTTCCCGGGTCAGTACCCCAAATATCTGATCAACCTGGCAAAGAGATGTTGGCACGGCGATCCATCACAGCGACCGGGCTTCAATTCCATCTGCAGAGCGCTTCGGTACATGAAGCGGTTCATGGTGATGAACCCTGATCACGGCCAGCCCGATGCGCCGATGCCACCGGTGGATTACTTTGAACTGGACATGAGTTTGTGCAAGAGATTCACAAACTGGGGGAGAAAGGACGTTCCTCGGGTCTCAGAAATCCCCTTTCAGATGTACGCTTACAGAGTGGTGGAGAGGGAGAGGACGAGTGCCAATGTGAAGGACAAGTGCTCGGATTCAGGGAGCGAAGGGGCTTCAGTTTGCGGCGATGAGAATGCATTTAGTATAACAGTGCCGGACGACGCGGTCTCTGCCTCTGTGGCTTCTGTGAGGTCACTGTATCCCATGGTATCTGAATCTAATAACAGGACCCCAACAAAGAAGGCTAGTAGTGGAAAGACCAATAATCCTTTAGGGAAGCTGCAGAAATCAAGAACCATGATACCCCCACACGTATCACCGGCAGGGCGCAACTTTAGAAGCAACTCTGAGAGCCGGCTGCAGCTGCAGCTGCAGCTCCAGCCAGTTATGATGAGCCCAAGAAGACGAAGGCCGTCGGGGCATGCCTCAGATTCAGAGCTGACATAG

>M_acuminata_2

ATGGAGCAATTCCGGCAGCTTGGCGAAGCCGTCGGAAGCCTCAAGGCGCTCATGGTGTTCCGCGACGAGATACGCGTCAACCGCCGGCAGTGCTACCTGCTCGTGGACGCCTTCGACCTGGCCTTCGACGCCGTCGCGGAGGAGATGCGTAGCCACTTGAGGTTCGAGGAGAAGCCGGGGAAGTGGCGCGCCCTCGAGCAACCTCTCAAGGAGCTCCACCGCGTCTTCCGGGAGGGAGAGCAGTACGTGAGGCGGTGCCTGGAGCCGGGGGAGTGGTGGGGGAAAGCCGTCGCCCTCAACCAGAACTCCGAATGCGCGATCGAGAACGTCGCCGAGGCCACCGCCGGGGGTGACCAGGAAGACATCCAAAAGAAGCGGCTCGTCTTCTCCAAGAAGTACGAGAGGGAGTGGATGGAGCCCGAGCTCTTCCAGCACAAGCTAGGGAAGCTCTACTTGTCCTCGCAAGAACTGCGCACCAGATTGGAGACGGCATGGAAGGAGGACCGATGGATTCTCTCCGAGACCATAGCCGAGAAGGCGAGCTCCGGGTCCAGGCCTCTGACGAAGCAGGAGAACAGGCTTGCCCAGCTCCTGGTTGGCCCGAAGGGGAAGCTTTGGCCGAGCTCAGTCCTCGTTGGCTCGCCGGACTACCAAGTAAGAAGAAGATTTGGGAGTGGGAGCGTCTACTACAAGGAGATCCAATGGATGGGCGAGACCTTCGTGGTGAAGCACGTTATCGGAGACACAGCGTCACTGACGAACGAGATCTCCATCCTCTCATCGATCGCGCACCCGAACGTAATGCAGTACATGTACTGCTTCACCGACGAAGAGAAGAAAGAGTGCTTGATGCTGATGGAGCTGATAAGCAAAGATCTCTCTTGTTACATACGAGAGGTTTGCTCCACAAGGAGAAAGGTGCTTCCTTTGCTGGTGGCAGTGGACACGATGCTTCAAATAGCCAGAGGAATGGAGTACCTTCACTCGAAGCAGATCTACCATGGCGACCTGAACCCTTCGAACATCTTCGTCAAGGCGAGGAATCCATCACCAGACGGCTATCTGCACGCGAAAGTTGGCGGCTTGGGGCTGTCGCCGGCGACGAACTCCAAGGCTTCATCATGCATTTGGCATTCTCCGGAGGTTCTACTGGAACAGGAGCAGACGGGGGATGGCAGCAGCAGCAGCAGCTCAAAGCGCACGGAGAAGGCGGACGTCTACAGCTTCGCCATGATATGCTTCGAACTCTTGACAGGAAAGGTACCCTTCGAGGACAACCATCTGCAGGGAGACAAGATGAGCAAGAACATCAGGGCCGGGGAGAGGCCATTGTTTCCGTCTCAGTCTCCCAAGTACCTCGTGACCCTAACGAGGAGATGTTGGCACTCTGATCCCTCGCAGCGCCCGAGCTTCTCCTCTGTTTGCAGGGTGCTCCGTTACCTCAAACGGTTCCTGGTGCTGAATCCAGATCACAGCCAGCCGGATCCGACGCCGCCGCCGGTGGATTACTTCGATCTGGAGAGCAGTCTGTCGAAGAGGTTCGCGAACTGGGCAAGGGGCGACCTTCTCCGAGCCTCGGAAGTGCCCTTCCAAATGTATGCATTCAGAGTGCTGGAAAGGGAGAGGACGAGCGCGAACGTGAAGGAGAGGAGCTCGGAGTCGGGCAGCGAGGGGGCATCAGTTTGCGGCGATGAGAATGCATGTAATGGTGTTCTTCCAGATGATGATGATACAATCGCGACCGTTGTTAGTGCCGATAAGTCGTTGCCTCAAAAGGTCACCCTTGACACGAACAAGAAAACAACATCGGCAAGGAAGGCCGATGGAAAGGCCAACAAGCAATCATCAGGGCAGAACCAGAAAGCAAGAATGGTCAGATCACCACAATTATCGTGTGGACGCAGCGTGAAGATGAACTCCGAAAGGCAGCTGCAACCAGTTGTGATGAGCCCAGGAAGGCAGAAGACATCAGGACATAACTCAGATACATAG

>M_acuminata_3

ATGGAGCAATTCCGGCAGATTGGTGAAGTTCTGGGAAGCATCGAGGCCCTGATGATCGTGGCAGAGGAGATGAGGAACCATCTGAGGTTCGACGAGAAGCTCATCAAGTGGAAAGCCCTCGAGCACCCTCTCAGGGAGCTCCACCGGATCTTCCGAGAAGGGGAGCAGTATCTCAGGCAGTGCTTGGAGCCCAGTGATTGGTGGGGAAAAGCCATAGCCCTCACTCACAACACTGACTGCGTCGAGTTCCACCTTCACGACTTGTTATGGTGTGTCCCCATCGTGATGGAGGCGATCGAGAATGTCGGGGAGATCACGGGAACAGACCAAGAAGACATTGGTAGGAAGAAGCTCGTTTTCTCGAAGAAGTACGAGAAGGAGTGGATGGAGCCTAAGCTCTTTCAGCTCAAGTTTGGTAAATCCTACTTGGCTTCTCAGGAACTACGCAACAAAATGGAAACAGCATGGAAGGAAGATCGATGGGTTCTCTCGGAAACGATAGCCGAGAGGAGAAGCCCAGGATCCAAGCCCCTGTCTAAGCAAGAGAACCGGCTTGCCGAGCTCCTGGTGTGCCCCAAAGGGAAGCTCTTCCCATCCTCAGCTCTTGTAGGATCGAATGACTACCAAGTCAGAAGAAGATTTGGGTCCGGAAACAACTACAAGGAAGTGCAATGGATGGGGGAGAGCTTTGCCGTGAAGCATGTCATCGGAGAGATTGAGCCACTGATGTCTGTGATTTCTCTCCTATCATCCGTCTCACACACAAACGTAGCGTATTACGTGTACTCGTTCGTCGATGAAGAGAAGAAGGAGTGCTTTCTGCTGATGGAGCTAATGACCAAGGATCTGTCCAGCTACATCAAAGAAATTTCCTCCACCAGAAGAAAGGTGTTATTCCCTTTGCTGGTGGCAGTAGACATAATGCTTCAGATTGCAAGAGGAATGGAGTATCTCCATTCCAAGAACATATACCATGGCGACTTGAACCCTTCAAACATATTGGTTAAGACGAGGAATTCTTCACCAGATGGATACTTGCATGTGAAAGTTACTGGATGTGGGCTTTCACCAATGAAGAACTCCAAGCCTTGGGCAAACCAGGCAGCTGCCACAAACCCATGTATCTGGTATGCTCCAGAAGTCCTCCTAGAGCAGGAGCGGTCAGGGGAGAGCAGTGGTGGCTCAAAGTGCACGGAGAAGGCAGATGTCTACAGCTTTGCAATGATATGCTTCGAGCTGCTGACAGGGAAAATTCCCTTTGAAGACGACCATCTTCAAGGAGATAAGATGAGCAAGAACATAAGAGCGGGTGTGAGGCCTTTGTTCGCATGTCAATCCCCCAAGTATCTGACCACCCTAACAAAAAGATGTTGGCAGACCGATCCGTCCCAGCGGCCCAGCTTCTCTTCCATTTGTAGAGTGCTCAGGTACATCAAGCGATTCTTGGTTATGAACCCAGATCATAGCCAACCTGATGCTCCAATGCCACCGGCGGATTACTATGACCTCGAAATTAGTCTGTGCAAGAGGTTCACAAACTGGGGAAGGAAAGATGTTCCCCGGGTCTCAGAAATCCCCTTTCAAATGTACGCTTATCGAGTTGCAGAAAGGGAGAAGACTAGCACCAACATCAAGGACAAGTGTTCGGATTCAGGGAGTGAAGGAGCCTCGGTTTGCAGCGATGAAAACGCGTTCATTATAACTCTCCCAGATGATGTGGTTTCTACCTCTGTGGGTTCTGTGAAGTCATTCTCTCCCACGATTTCTGATACTACCAACAAGACCTCCTCCACGGTGAAGGCTAGTGGGAAGAGCAATAAGCAATTAGGTAACTTGTGGTTTCATGCAGTGCTCGTTTGGATTTCAATTTGGCTCTCAAATTTCTTGGCATCCTTGATGTTAACCATTATACCCACACACCTGACCGGACGCACTGCAAGAACCAACGCTGAAAACCGGCTGCAGCTGCAACCAGTTATGATGAGCCCAAGAAGACGGAAGGCATCGGGGCATGCCTCAGATTCAGAACTAACATATATCTGA

>M_acuminata_4

ATGGAGCAATTCCGGCAGATCGGTGAAGCTCTAGGAAGCATCAAGGCTCTGATGGTGTTCCGCGACGAGCTGCGCGAGATGAGAAGCCATCTAAGGTTCGACGAGAAGCTCATCAAGTGGAAAGCCCTCGAGCCGCCTCTCAAGGAGCTCCACCGGATATTCCGAGAGGGGGAGCAATACATCAAACAGTGCTTGGAGGCCAGGGACTGGTGGGGGAAAGCCGTAGCCCTCGGCCAGAACGCCGACTGCGTCGAGTTCCACATCCACGACCTGCTATGGTGCATCCCCGTCGTGGTAGAGGCGATCGAGAACGTCGGGGAGATCACGGGAACCGACCAAGAAGAGATATACAGGAAGAAATTTGTCTTCTCCAAGAAGTACGAGAAGGACTGGTTGGAGCCCAAGCTGTTTCAGCTCAAGCTGGGAAAGACGTACTTGGTCTCGCAAGAACTGTGCAGCAGGATGGACACGGCATGGAAGGAAGATCGATGGATCCTATCGGAGAGGATAGCCGAGAAGCGAAGCCCTGGATCGAAGCCCCTGACGAAGCAAGAGAACCGGCTCGCTGAGCTCCTGGTGTCTCCCAACGGGAAACTCTTCCCCTGCTCGCTTCTCACCGGATCCAGCGACTACCAGGTCAGAAGAAGATTTGGGTCTGGAAACAACTACAAGGAAGTGCAGTGGTTGGGCGAGAGCTTCTCCGTGAAGCATGTCATCGGTGAGATTGAGCCACTGATGAGTGAGATATCTCTCCTCTCGTCCATCACGCACCCGAACGTCGTGCGCTACATGTACTCGTTTGTCGACGAAGAGAAGAAAGAGAGCTACATGGTGATGGAGTTGATGAGCAAGGATCTCTGCAGCTACATCAACGAGATCTCTTCCACGAGGAGAAAGGTCCCCTTCCCTCTGCTGGTGGCGGTGGACACGATGCACCAAATCGCGAGGGGAATGGAGTATCTTCACGGTAAGAACATATACCACGGCGACTTGAATCCTTCGAACATATTGGTCAAGACAAGAAACTCTTCGCCGGATGGGTATCTGCACGTAAAAGTGACCGGCTTTGGTCTGTCGTCGGTGAAGAACTCCAAACCTTCGGCAAACCAAGCACCTGCCACCAACCCATGCATCTGGTATGCCCCGGAAGTGCTCCTGGAGCAGGAGACGTCGGGGGAGAGCGGTAGCGCATCGAAGTGCACGGAGAAGGCAGACGTCTACAGCTTTTCGATGTTATGCTTCGAGCTGTTGACGGGGAAAATCCCCTTCGAGGACGATCATCTCCAAGGGGATAAGATGAGCAAGAACATAAGAGCCGGTGCAAGGCCGCTGTTCCCGTTTCACTCTCCCAAGTTCCTCACCAACCTCACAAAGAGATGTTGGCACGCCGATCCCTCTCAGCGACCCAGCTTCTCTTCCATCTGCAGAGTGCTTCGGTACATCAAAAGGTTCCTAGTCATGAACCCAGATCACAGCCAGCCTGATGCACCGATGCCACCGGTGGATTACTTCGACCTCGAAACCAGTCTGTCCAAGAAATTCGCAAGCTGGGCAAGGAAGGACACTCCCCGCGTCTCAGAAATCCCCTTTCAAATGTATGCCTACAGGGTCGTGGAAAGGGAGAAGACGAGTGCCAACGTCAAGGACAAGTGCTCGGATTCGGGGAGCGAAGTAGCGTCACTTTGCGGCGATGAGAACGTCACGAGACCACCGCACCTTTCAATGTTTGGGCGCAACCTCAGGACGAACCCTGGGAGCCGAATTCAACCAATCATGATGAGTCCAAGGAGGAGGGCAACAGGGCATGCATCGGATTCCGAGCTGACATAG

>M_domestica_1

ATGGATAACTCTGCAAACATGCATAATGATCATTTGGGTGTGAACAAAATGGGGAAAAATATAAGGAAGAGTCCTTTGCACCAACCCAAATTTGGTAATAATGACGCCAGGCAACAACCTCAACCTCAGGTGTACAATATAAGCAAGAATGACTTCCGGAACATTGTTCAGAAGCTTACTGGTTCGCCATCGCAAGAACCTTTGCCTAGACCTCCCCAGAATCCGCCAAAGCCCCAAAGTATGCGATTGCAGAGAATTCGACCTCCCCCATTAGCACCTATCAACAACAGACCCTTAAACACACCTCCAGCTCCTCTTCCTGCAGGCCCACCACAGGTTCCCTACAATAATAACTTTGTTAGGCCTCCTCAGTTTGGACATCCATCACCTACACCAATGCCACCATTTCCACATGGAGATTCAATGTGGCAAAATACAGCTGAATCTCCTATCTCAGCATATATGCGGTACCTTCAGAGTTCAATGTTAGATCCAACTCCAAGGGGCAACCAACCTCAGCCTCAACCACAAGTTCCAGGTCAAATGCAGTCTCAGCCACCATCAACCAGTTTACTTCCTAACCCATCCGTATCTGCTCACCCACCCCCGAGAATGAATGCCCCTGGGGCACATGCGCCTAATCTACAGCACCCACAGATGAATGGTCCTCCCCTCTTACCGTCACCAACTTCACAGTTTTTATTGCCATCCCCAACGGGTTACATGAACTTGTTGTCTCCACGGTCACCTTATCCATTGCTTTCACCTGGGATGCAGTTTCCTCCACCATGGACCCCCAATTTTCAGTTCTCTCCAATGGGCCAATCAGGGCTGTTAGGTCCAGGGCCTCAACCCCCACCTTCTCCTGGCGTTTTGTTTCCATTATCTCCTTCAGGGTTTTTCCCCATTTCAAGAGGCGGTTCGCATGGATGCTTGTCCGACCACGACTTCCTTCTTCCTTGGAGGGACAACGTTGAATCTTTTAGTTTGATTCCGACCAGACACCAGACAATACCAGAAACTGAAACGGAAATTTTTCGCCGTCTGAGCCGGAATGGAACAATTCCGGCATATTGGGGAGGTTTTGGGAAGTATGAAGGCTCTGATGATTGCTGTTTGTTGCATGATGTGTTCACGTTGGCGTTTGACACGATCGGAGAGGAGATCCGACTGAACTTGAAACTCGAAGAGAAGAAGACAAAATGGAAAGCGCTCGAGCAGCCTCTGAGGGAGCTGCACAGAGTTTTCAAGGAAGGGGAGCTGTACATTCGGCATTGCATGGATGCAAAAAACTGGTGGGGGAAAACAATCCTCCTCCATCAGAACAAGGACTGCGTCGAATTTCACATTCATAACTTGTTCTGCTACCATGCGGCCGTGATTGAGGCGATTGAGAATGCTGGAGAGATTGCGGGGCTCGATCAGGAGGAGATGCAGAAGAAGAGGATTCTGCTCGCAAGGAAGTATGATAGAGAGTGGAATGACCCCAAACTTTTCCAATGGAGATTTGGGAAGCAGTATTTGGTCCCGAAGGCGATCTGCAACAGGTTGGAGAGTGCTTGGAGGGAAGATAGATGGCGGCTTGTCGAAACTCTGAAGGTGAAAAAGATTTCAGGATCCTTTGGTTTGACGAGGAATGAGCAGCAACTTGGAGACTTGCTGCTCAAGAAACTTGGTGGGTCGGAAACCTTAAATGGGAAGCTCTTTCCGAGCTCAATCTTACTGGAATCCGACTATTACAGCGTTAGGCGGTGGTTAGGAGGAGGAACGCAGTACAAGGAGATCCAGTGGTTGGGGCAGAACTTTGCCATGAGACACTTCTTTGGGGATATTGAGCCGTTGAATTCTGAGATTTCAACTCTCCTCGCACTTTCCCACCCCAATGTGCTGCAGTACCTTTCGGGGTTTTATGACGAGGAGAAGAAGGAATGCTTTCTTGTTATGGAGTTGATGATCAAGGATCTGCGCTGCTACATGAAGGAAAACTGCGGCGCGAGAAGGCAGGTTTTGTTTTCAATCCCGGTAGTGGTTGATATCATGCTTCAGATTGCAAGAGGCATGGAATATCTGCACTCTAGGAAGATCTACCATGGGGAGTTGAACCCCTTGAATGTCTTTCTCAAGGCAAGGAGCTGCACAGAAGGCTATTTCCAAGTAAAAGTCTCGGGTTTCAGTTTATCATCCGTGCGCAAACCTACTTACAGAAAATCACAGCAGAAGAATGAAATCAACCCTTTGATTTGGTGTGCCCCGGAAGTCCTAGCTGAGCAAGAGCAACCAGGAAACAAAGGTCGTACCAAATACACGGAGAAAGCGGATGTATACAGCTTCGCAATGCTTTGCTTTGAGATCTTGACCGGGAAGGTTCCCTTCGAAGATTCACATCTCCAAGGGGACAAGATGAGCAACACTATAAGAGCAGGATGGAGGCCTCTCTTCCCATACCCTTCACCAAAATACCTCGTCAATTTAACCAAGAGATGCTGGCACAGTGACCCGTCTCAGCGCCTGAGCTTCTCGTCCATCTGTCGCATTTTGCGCTACATAAAGAAGTTCCTTACTCTGAACCCCTATGACGATCAGCCTTTACTGCAATCCCCTCATATGGATTACTGCGAGATAGAGTCATGGTTCCTGAAGAATAGTTCAGCTGCCGAATACGTTGATTTATCCTCAATATCGCAACTTCCATTCCAAATGTTCTCTTACCGGCTAGGGGAGAAGGAAAAGACTAGCCCGGACATCGTGAGGGTTAGGAGTCTGGATTCCTTAAGTGATACAGCATCAGCGATTTGCAAGCAAGAATCTCCGAACGGCAAAGTTGACATTGTGTCCATTGCAGAAGATCCATTCGTACCACTAAGCGACTCGAGGTCCGTTTGCTCTGATGTGAGGTCTGTTTATGATCTGAGGTCAGTTTGTTCCGAGGCTCCGATGAAGAGAACTCTCACTGCCAAGAAACATCAAGACGCGACAGCTAGAAAAGCCTCAGGTTACACCAGGACATCAAAAACACCAACCACACCAAGGACGTCAACACCAAGACTTGCGTCGCCAAGATTTCCACCGCCAAAACTTGCTTCACCGAAACTTCTGCCTGCAAAACCATGTCAACACAGTATGAAGACAAACCAAAGCCCTCCACTGCCGAGTCCTACGAGCACAAAGAGTAGTGCGAGCCGAAGCAGGCACAGAATGAGGGGGCACGTTTCGGACTCGGAGATACATTAG

>M_domestica_2

ATGGAGCAATTTCGAAAGATCGGGGAGGATTTAGGAAGTTTAAAGGCCTTAATGGTGTTCCAAGACAACATCCAAATCAATCAGCGGCAGTGCCTTCTGCTGCTCGATATCTTCAGCTCAGCGTACGAATCAATAGCGGAGGAGATGAGACATAATCTTCGGTTTGATGAGAAGCACATGAAATGGAGGGTTCTCGAGCAGCCGCTGAGAGAGCTCCACAGGATATTCAAAGAAGCAGAAGCTTACATTCGACAATGCTTGGAAACCAAGGATTGGTGGGCTAAGGCCATCACTCTCTATCAGAACTCCGATTGCGTTGAGTTTCACATCCATAACCTACTTGCCTGCATGCCAATCGTCATTGAAGCGATTGACATTGCTGGAGAAGTTTCAGGGTGGGATCAGGATGAGATACAGAGGAAGAAAACCGTGTTTGCGGATAAGTACAAAGAAGACTACAGAGATTGGAAACTTTTCAAGTGGAGGTTTGGGAAGCAGTATTTGATTACTCAAGACTTCTGCAATAGGTTTGAATCAGCAAGCAAAGAAGATAGATGGACTCTTCAGCATAAAATCCGCGAAAAGAAACTTTCGGGTTCAACAAAGTACGGGAAACGCCTCATAGATCTCCTTTTCAAAAGCATAGACGGATCAGAGACACAGAATTTGAATGGGAAGCTCTTACCTAGTTCAATCCTGGTGGGGTCTAAGGACTACCAGGTGAGGCGGCGACTTGGGGGTGGAAGTCAGTATAAGGAGATCCTATGGTTGGGTGAAAGCTTTGCCTCAAGACATTTAATTGGAGAAGTAGAAACCTTGTTGCCGGAGATTTCTTCGTTGTCATTGCTTTCCCACCCGAACATAGTGCACTTCCTTTGTGGGTTTACGGATGAGGAAAAGAAAGAGTGTTTTCTCATTATGGAACTCATGAGTAAAGACCTCTGCAGCTACATCAAAGAGATTTGTGGCCCGCGAAAACGTCTTCCATTTTCTCTTCCTGCTGCTATTGATCTGATGCTTCAAATTGCAAGAGGAATGGAATATCTCCACTCGAAGAAAATCTACCACGGAGATTTAAATCCTAGTAATATACTTGTTAAAGCAAGAGGCATCTCCACAGAAGGTTACTTACAAGCCAAGGTTTCAGGTTTCGGTTTAACTTCTGTCAAAAGCCCCACACAGAAAAGCACTTCAAATCAGAATGGGAGCCTACCTTTAATCTGGTATGCGCCGGAAGTTCTGGAAGAGCAAGAACATGGAAAAAGTACTGAGAAAAAGTACGCAGAAAAGTCCGATGTGTATAGCTTTGGGATGGTTTGCTTTGAGCTTCTGACAGGGAAAGTGCCCTTTGAGGACAGTCATCTCCAAGGGGACAAGATGAGCCGAAACATAAGGGCGGGGGAGAGGCCGCTGTTCCCATTCTACTCTCTTAGATATGTGACCAACCTCACAAAGAAATGTTGGCATAGTGATCCATACCAACGGCCAAGCTTCTCGTCCATCTGCAGGATTCTTCGCTATATCAAACGGTTCCTTGCAATGAATCCTGATTACAACAGCCAGCTAGACCCACCGGTTCCTATGGTAGATTACTGTGAGATTGAGTCAGTGCTTCTGAAAAAGATTCCTTCTTGGGGGACTTCTGAACCATCACCAWTATCACAAATCCCATTCCAAATGTTCGTTTATAGGATCGCAGAGAGAGAAAGAACATCTTCAAGCCTGAAGGATACTTCAGAATCGGGAAGYGACGGAGCTTCAATCTGTGGGGAYGAAATTGTGGTCCCTGCAGATGATTCGTTCCCATCAACACCTGAAAGGAAGAGTTTCTCTTCGCCTGATAGCGTGAAAAAGAAACTTCCATTCTTGAAGAAATCATCAGATGTGAAAGCCAACAGACTATCAGGAACACCAAGAGGACGGTCCAGACCTCCACAGATGACCCCTTGTGGCCGCAGTGTTAGTCTGAGATTGGGTTCAGAAAGCCAGCTGATGGAAATGAGTCCAAGAATACGGAGAACTAAATCCGGACATAGGAACGATCYATTCAAGGCATCCGTCGAGCATAGTGCAGGCGAGACTTCAACAGTGATGATATCTGACAAGATATTACCGCTTAACAGCAAGACAAAAGACAATCGGAAATCATTTCAGCCGGGRGCGAAAAAAGAGATTGTATATATCTTTGCAAAGCATTTAGAAGAGGCACARTTTCTATTAAAATTTAGAGTTCCAAGCATTTAG

>M_domestica_3

ATGGAACAATTCCGGCATATTGGAGAGGTTTTGGGAAGTATGAAGGCTCTGATGATGTTGAGAGATGAGATTCTGATCAATCAATGCCAGTGCTGTTTGTTGCATGATGTGTTCACGTTGGCGTTTGACACGATCGGAGAGGAGATCCAGTTGAACCTAAAACTCGAAGAGAAGAAGACGAAATGGAAAGCCCTCGAGCAGCCTCTGAGGGAGCTGCACAGGGTTTTCAAGGAAGGGGAGCTTTATATGCGGCATTGCATGGATGTGAAAAACTGGTGGGGCAAAACAATCCTCCTCCATCAGAACAAGGACTGCGTCGAATTTCACATCCACAACTTGTTCTGCCACTATGCAGCCGTGATTGAGGCGATTGAGAATGCTGGAGAGATTGCGGGGCTCGATCAGGAGGAGATGCACAAGAAGAGGGTTCTGCTCACGAGGAAGTATGACAGGGAGTGGAATGACCCCAAACTTTTCCAATGGAGATTTGGGAAGCAGTATTTGGTCCCGAAGTTGATTTGCAACAGGTTGGAAAGTGCTTGGAGGGAAGATAGATGGCGGCTTGTTGAAGCTCTTAAGGTGAAAAAGATTTCAGGATCCTTTGGTTCAACGAGGAATGAGCAGCAGCTTGGAGACTTGCTGCTCAAGAAACTGAGTGGGTCGGAAACCTTAAATGGGAAGCTCTTTCCGAGCTCAATCTTATTGGGATCAGCGGATTACCATGTTAGGCGGCGGTTAGGAGGGGGAACGCAGTACAAGGAGATCCAGTGGTTGGGCCAGAACTTTGCCATGAGACACTTCTTTGGGGATATTGAACCTTTGAATTCGGAGATTTCAACTCTTCTCACACTTTCCCACCCCAATGTGCTGCAGTACCTTTCCGGGTTTTATGACGAGGAAAAGAAGGAATGCTTTCTTGTTATGGAGTTGATGAACAAGGATCTGCGCTGCTACATGAAGGAAAACTGTGGCGCAAGAAGGCAGGTTTTGTTTTCACTCCCGGTAGTGGTTGATATCATGCTTCAGATTGCAAGAGGCATGGAATATCTCCACTCTAGGAAGATCTACCATGGGGAGTTGAACCCTTTGAATGTCTTTCTTAAGGCAAGGAGCTGCACAGAAGGCTATTTTCATGTAAAAGTCTCAGGTTTTGGTTTATCATGCGTGCGCAAACCTACTTATCGAAAATCACAGCAGCAGAACGAAATCAACCCTTTGATTTGGTGTGCCCCGGAAGTCCTAGCTGAGCAAGAGCAACCAGGAAACAAAGGTCGTTCCAAATACACGGAGAAAGCAGACGTATACAGCTTCGCGATGCTTTGCTTTGAGCTCTTGACTGGGAAAGTTCCTTTCGAAGATTCACATCTTCAAGGGGACAAGATGAGTAACACTATAAGAGCAGGAGGGAGGCCTCTCTTCCCATACCCTTCACCAAAATACCTYGTCAATTTAACCAAGAGATGCTGGCACAGTGACCCGTCTCAGCGCCTGAGCTTCTCRTCCATCTGTCGCRTTTTGCGCTACATAAAGAAGTTCCTTACTCTGAACCCCKAYGACGATCAGCCTWTACTGCAATCCCCTCMTAYGGATTACTGCGAGATAGAGTCATGGTTCCTGAAGAATAGTTCAGCTGCCGRRTACGYTGATTTATCCTCAATATCGCAACTTCCATTCCAAATGTTCTCTTACCGGCTTGGGGAGAAGGAGAAGACTAGCCCTGGCATCATGAGAATCAGGAGTCTGGATTCCATAAGTGATACAACATCCGCAAATTTCAAGCAAGAATCTCCGAATGGCAACGTTGACACTATGTCCATTGCAGAAGATCCATTCGTACCACTAAGCGAAACGAGGTCCGTCTGTTTTGATGTGAGGTCCTCTTATGATCTAAGGTCAGTTTGTTCCGAGGCTCCAATGAAGAGAACTCTCACTGCGAAGAAACGTCAAGAGGCGATGGCTAGAAAAGCCTCAGGTTACACACGGACATCAAAAACACCAACAACACCAAGACTCTCAACGCCAAGGCTTCCGTCGCCAAGATTTCCATCCCCAAAACTTCCGCCACCAAAACTCGCTTCACCGAAACTTCCACCAGCAAAGCTTTCGGTTCCAGCAAAACCATGTCTACGCGGTATGAAGACAAACCAAAGCCCTCCACAGCCGAGTCCTATGAGCACAAAGAGTAGTGCGAGCGGAAGCAGGCACAGAACGGGCGGACACGTCTCGGATTCGGAGATACATTAG

>M_domestica_4

ATGGAACAATTCCGAAGGATTGGGGAGGATGTAGGCAGCTTAAAGGCTTTAATGGTGTTCCAAGACAACATCCAAATCAATCAGCGGCAGTGCTTTTTGCTGCTCGATATCTTCAGCTCAGCATATGAATCAATAGCGGAGGAGATRAGACATAATCTTCGGTTTGATGAGAAGCACATGAAATGGAGGGTTCTCGAGCAGCCTCTGAGAGAGCTCCACAGGATATTCAAAGATGCAGAAGCCTACATTCGACAATGCTTGGATACCMAGGATTGGTGGGCTAAGGCCATCACACTCTATCAGAATTCCGATTGYGTCGAGTTTCACATCCATAACCTACTTGCCTGCGTGCCGATCGTCATCGAAGCAATCGACATTGCTGGAGAAATATCAGGGTTGGATCAGGATGAAATACAGAGGAAGAAAACCATGTTCGCAGACAAGTACAAACAAGACTACAGAGATTGGAAACTTTTCAAGTGGAGATTTGGGAAGCAGTATTTGATTACTCAAGACTTCTGCAATAGGTTTGAATCAGCAGGAAATGAAGATAGATGGACTCTTCTGCATAAAATCAGCGAAAAGAAACTTTCAGGTTCAACAAAGTATGGGAAACGACTCATAGATCTTTTCAAAAGCGCAGACGGATCAGAGTCACAGAATTTGAAAGGGAAGCTCTTACCTAGTTCAGTTCTGGTGGGATCTAAGGACTACCAGGTGAGGCGGCGCCTTGGGGCTGGAAGTCAGTATAAGGAGATCGTATGGTTGGGTGAAACCTTTGCCTCAAGACATTTCTTTGGGGAAATTGAACCCTTGTTGYCGGAGATTTCTTTGTTGTTATCACTTTCCCACCCAAACATAGTGCTCTTCCCTTGTGGCTTTACRGATGAAGAAAAGAAAGAGTGTTTTCTCATAATGGAACTGATGAGTAGAGACCTCTGCAGCTACATCAAAGCGATCTGTGGCCCCCGAAAACGTCTACCATTTTCTCTTCCTGTTGCTATTGATCTGATGCTTCAAATTGCAAGAGGAATGGAATATCTCCACTCAAAAAAAATCTACCACGGAGATTTGGATCCTTCTAAYGTACTTGTTAAAGCAAGAGGCATCTCCACAGAAGGTTACTTGCAAGCCAAGGTTTCAGGTTTTGGTTTAGCTTCTGTCAATACCTCCAACCACAAAAACCCTTCAAATCAGAATGGAAGCCTGTCTTTCATCTGGTATGCGCCTGAAGTTCTGGAAGAGCAAGAACAAGGAAAAAGTACWGAGAAAAAGTACGCAGAAAAGTCCGATGTGTATAGCTTTGGAATGGTTTGCTTTGAACTCCTGACAGGGAAAGTCCCTTTTGAGGATAGTCATCTTCAAGGGGACAAGATGAGCCGAAACATAAGGGCCGGGGAGAGGCCGCTGTTTCCRTTCTATYMTCTGAGATATGTGACCAACCTCACAAAGAAATGCTGGCATAGTGACCCCAATCAACGGCCAAGTTTCTCATCCATTTGTAGGATTCTTCGCTACATTAAACGGTTCCTTGCGATGAATCCTGATTACAACAGCCAGCTAGACCCGCCTGTTCCTATGGTAGATTACTGTGAGATTGAGTCTGGGCTTCTGAGAAAGATTCCTTCCTGGGGGAGTTCTGAACCATCACCAATATCACAAATCCCATTCCAAATGTTCGTTTATAGGATCGCAGAGAGAGAAAGAACGTCTTCAAGCCTGAAGGATACTTCAGAATCYGGAAGTGACGGAGCTTCAATCTGTGGCGATGAAATTGTGGTCCCTGCAGATGATCCGTTCTCAACAACACCTGAAAGGAAGAGTTTATCTTCGCCTGATAGTGTGAAAAAGAAACTTCCATTCTTGAAGAAATCGTCGGATGTGAAATCCAACAGACTACCAGGTTGCCGAACACCAAGAGGACGGTCCAGACCTCCACAGATGACCCCTTGCCGCAGTGTTAGTCTGAGATTGGGTTCAGAAAGMCAGCTGATGGCAATGAGTCCAAGAATACGGAGAACTAAATCCGGACATGCCTCAGATTCTGAGCTCTCCTAG

>M_domestica_5

ATGGAGCAATTTCGAAAGATCGGGGAGGATTTAGGAAGTTTAAAGGCCTTAATGGTGTTCCAAGACAACATCCAAATCAATCAGCGGCAGTGCCTTCTGCTGCTCGATATCTTCAGCTCAGCGTACGAATCAATAGCGGAGGAGATGAGACATAATCTTCGGTTTGATGAGAAGCACATGAAATGGAGGGTTCTCGAGCAGCCGCTGAGAGAGCTCCACAGGATATTCAAAGAAGCAGAAGCTTACATTCGACAATGCTTGGAAACCAAGGATTGGTGGGCTAAGGCCATCACTCTCTATCAGAACTCCGATTGCGTTGAGTTTCACATCCATAACCTACTTGCCTGCATGCCAATCGTCATTGAAGCGATTGACATTGCTGGAGAAGTTTCAGGGTGGGATCAGGATGAGATACAGAGGAAGAAAACCGTGTTTGCGGATAAGTACAAAGAAGACTACAGAGATTGGAAACTTTTCAAGTGGAGGTTTGGGAAGCAGTATTTGATTACTCAAGACTTCTGCAATAGGTTTGAATCAGCAAGCAAAGAAGATAGATGGACTCTTCAGCATAAAATCCGCGAAAAGAAACTTTCGGGTTCAACAAAGTACGGGAAACGCCTCATAGATCTCCTTTTCAAAAGCATAGACGGATCAGAGACACAGAATTTGAATGGGAAGCTCTTACCTAGTTCAATCCTGGTGGGGTCTAAGGACTACCAGGTGAGGCGGCGACTTGGGGGTGGAAGTCAGTATAAGGAGATCCTATGCTACATCAAAGAGATTTGTGGCCCGCGAAAACGTCTTCCATTTTCTCTTCCTGCTGCTATTGATCTGATGCTTCAAATTGCAAGAGGAATGGAATATCTCCACTCGAAGAAAATCTACCACGGAGATTTAAATCCTAGTAATATACTTGTTAAAGCAAGAGGCATCTCCACAGAAGGTTACTTACAAGCCAAGGTTTCAGGTTTCGGTTTAACTTCTGTCAAAAGCCCCACACAGAAAAGCACTTCAAATCAGAATGGGAGCCTACCTTTAATCTGGTATGCGCCGGAAGTTCTGGAAGAGCAAGAACATGGAAAAAGTACTGAGAAAAAGTACGCAGAAAAGTCCGATGTGTATAGCTTTGGGATGGTTTGCTTTGAGCTTCTGACAGGGAAAGTGCCCTTTGAGGACAGTCATCTCCAAGGGGACAAGATGAGCCGAAACATAAGGGCGGGGGAGAGGCCGCTGTTCCCATTCTACTCTCTTAGATATGTGACCAACCTCACAAAGAAATGTTGGCATAGTGATCCATACCAACGGCCAAGCTTCTCGTCCATCTGCAGGATTCTTCGCTATATCAAACGGTTCCTTGCAATGAATCCTGATTACAACAGCCAGCTAGACCCACCGGTTCCTATGGTAGATTACTGTGAGATTGAGTCAGTGCTTCTGAAAAAGATTCCTTCTTGGGGGACTTCTGAACCATCACCAWTATCACAAATCCCATTCCAAATGTTCGTTTATAGGATCGCAGAGAGAGAAAGAACATCTTCAAGCCTGAAGGATACTTCAGAATCGGGAAGYGACGGAGCTTCAATCTGTGGGGAYGAAATTGTGGTCCCTGCAGATGATTCGTTCCCATCAACACCTGAAAGGAAGAGTTTCTCTTCGCCTGATAGCGTGAAAAAGAAACTTCCATTCTTGAAGAAATCATCAGATGTGAAAGCCAACAGACTATCAGGAACACCAAGAGGACGGTCCAGACCTCCACAGATGACCCCTTGTGGCCGCAGTGTTAGTCTGAGATTGGGTTCAGAAAGCCAGCTGATGGMAATGAGTCCAAGAATACGGAGAACTAAATCCGGACATGCCTCAGATTCTGAGCTCTCCTAG

>E_guttata_1

ATGGATCAGTTCAGGCAGATTGGGGAGGTTTTGGGCAGTTTGAAGGCATTAATGGTAATGAAACACGAAATATCGATCAACCAGAGACAATGCTGCCTGCTTTTCGACATGTTGGAGTTAGCCTTCGAGACAATCTCCGACGAAATCCGACAGAATCTGAAGCTCGAAGACAAGAACACCAACAAGTGGAAGCCCCTCGAGTTTCCGATGAAGGAGCTCCACAAAGTGTTCAAAGAATGCGATCTCTACATACGACAATGCACCGACGTAAAAGAATGGTGGGGCCGAGTAATTAGCCTCCACATGAATAGGGACTGTTTGGATCTCCACATACATAACCTCCTCTCGTCTTTCCCCGTCGTGATCGAAGCGATCGAGACCGCGGCAGAAATCTCGGCCCTCGATCAAGAAGACATGCAGAAGAGACGAATCTCCCTCATGCAGAAGTACGACCAAGAGTGCGACGACCTAAAACACTTTTTGTGGAAATTCGGCAGAGAGTACTTAGTCCCGATAGAGATATGCGCCCGTTTAGAAAACGCTTGGAGCGAAGACCGTTGGCTCCTAGTCAAGAAAATCGAGGATAACAAAAAAACGGGGCCCATCGGTGAAATCTTGATCAGGAAAATGAAGAATGGGGATTCGTTGGATCGGAAGCTTCCGTCGAGCAACGTTTTGGTGGGGGCCCAAGATTATTTAGTGAAGCGGCGTTTGGCTTCGACGGGTGGGAATCTGAAGGAGATTCATTGGATGGGGGAGAATTATGCGTTGAGGACTTTTTACGGTGACGTGGGCCCGTTGTATTCCGAGATTTCTACGATTCTTTCGCTTTCGCATCCGAATGTGTTGCAGTACCTTTGCGCCTTTTACGATGAGGATCGGAAAGAAGGGTTTCTTGTGATGGAGCTTATGAGTAAGGATCTTGGTTCCTACATAAAAGAAAATTCCGGACAGAGAAACCGGGTCCCGTTTTCTATACCAATTGCTGTCGATATTATGCTGCAGATTGCTAGAGGGATGGAATATCTACACTCGAAGAAAATCTATCATGGAGTTTTGAATCCTTCGAGCGTACTTCTAAAGGCGAAAAACGCCTCGATTGGAGGGTTTCAAGCTAAAGTTGCGGGGTTCGGTTTGAGCTCGATCAAGAGCTACTACACTTCGAAAAGCCCTAAAAAAACGGGGGTCGATCTCGATATTTGGTTAGCACCGGAGATTTTAACCGAATTAGAAGGTAATAAAAACATTTCCCCAAACTACACCGAGAAAGCCGATGTTTATAGTTTCGGGATGCTCTGCTTCGAGCTTCTAACGGGGAAAGTTCCGTTCGAAGATGGCCATTTACAAGGCGAGAAGATGGCTCGTAATATACGAGCTGGCGAGAGGCCTCTTTTTTCGTACCCTTCTCCGAAATACTTGACAAATCTCACGAAAAAATGCTGGCAGACGAACCCTAATTTACGACCGAGTTTTTCCTCTATATGTAGGGTTTTACGGTACATCAAGAAAATCCTCGTCATAAATCCGGACCACGGCCACCCCGACTCTCCGCCGCCACTAGTGGACTACTGCGACATGGAGGCAGGGTACTTGAAGAAGTTTCCGGAAAAGGGGGGCGCGGGGGTTTTGGCCCCCCCGGTTCTGCAGATCCCGTTTCAGATGTTCACGTACAAGATAGTGGAAAAGGAGAAATTCTACTCCAAGAAATGGGAGATTCATAGGCCAGCTTCGATGTTTGACGAAGATCATAATATGGACGATTTCTTGCTTGCGCCGAGCGATCGGAGGTCCGTTTGTTCGGAGATTATCGACACCAAGAATTCCGGCATTGCTGTGGATATGAGGTCCGTTATTTCCGAGATTCCGCAACGGAAAATGCACCCGTTGGATCAGAGCTCGTTTGGTTTGGAGAGTCCTAGGAGGAAACTTTCGAGTAAATCGTTTAATCTGGAGAGGAGAACCGACCAGAATTTCACTAGTCGGGAGCTGAGAGAGGTTCTTGATAAAGCGATTATAGCATCCAAAGTAGCGGAATCTTCTCAGTTAGTGACGAAAGTGACGGAGCAGAAAGCGGAGAGTACCGAGATTCTTGAAAAGAAACAGTCTTTGAGACCAATAGTTGACCAAAAAACGAAATCTTTGGAGGGGGAAATGATACTGCGGCCGACGAGAAATCGGAAATTAAGCGAGATTCCGGAGAAGATAGTGAAGAATGGAGAATCTATAAAGAACACCGTTGAAAAAACCGAGGAAAAAACCGAGGAAAAAACTGGCAAGACAACCGTGATCATTAAACAGATGAAAGATGTAATAAAGACCAAGAAAATTGAAGAAACCCGAAACAGGAAAACGAGGCCCGTTTTCTCCGCAGCTTCATCTCCAGCGAGAGCTTTCGGCCAATCAGAAGCACCGAAAAACGAGCCGATCAGAATGAAGAAGAAAACTATGTCCGTTTCTTCGCCAGCTTCTTCTCCGTCTAGGGTTTTCAGAAAGTGCAGCCAATCAGAAGCATCACCGAAAGACGAGTCGCCGTCAAGAATGATGATGAAGAAGAAAAGTGCACCCGTTTCTTCGCCAGCTTCTTCGCCTTCCCGAACTTTCGCCACGTGCGCTTCTCCTTCGCCGAGACAGGTGTCGGCCTGTCATTCGTCGGTGGAGAATAATTCTTCTTCGTCGCCACTGGACCCGTTCGGGCGGTGCTCGAGACTCGGTAGACAGACCTCACTTCCGGACGTGACTGTGGGCCCACAAAGACCTAATCTAGATCGTGTTTCGTCTCATGTAGCATAG

>E_guttata_2

ATGGAGCAATTCAGGCAAATGGGAGAAGTTTTGGGAAGTTTAAAGGCACTAATGGTGTTCGAAAACCACGTATTCAACCCCCGACAATGCTGTCTGTTGGTAGACATGCTCACTTCCGGCTACGAAACGATCGCGCATCATATGAAACAGAATCTCCGATTCGAGGAGAAGAACACGAAATGGAAAATCATCGAGCATCCGTTGAAGGAGCTTCTCCGAGTATTCAAAGAAGGAGAATCGTACGTCAAGCAGTGCCTCGAGACGAAAGACTATTGGGCGAAAGCAATTGTCCTCTACCAAAACTCCGATTGCGTCGAGTTCCACATACACAACTTGCTCAGCACAATCCCGATAGTGATCGAAGCCATCGAAATGGCCGGACAAATATCGGGATCCGATCAAGACGAAATCCAACGCAAGAAATTCGTTTACTCGCTCAAATACCAGAAAGATTGGAGGGATCCAAGAATCTTCCAATGGATATTAGGCGACCACTACCTCGTTTCCCAAGACTTCTGCAATCGAATAGAGTCCGTTTGGGACGAAGACAGATGGTTCCTTATCAAGAAAATCAAAGACAAGATGCTAAACAACACCGGGAGTTCACACTCGAAACGTGACAGACGAATGGCCGACCTTCTTCTAAACTACTTAAATTCCTCCTCCTCCTCCTCCTCCTCAACATCATCGTCGCTATTACTACCGAGTTCGACTTTGGTGAACTCGAAGGATTATCAAGTGAGGAGAAGGCTCGGAAACGGGAGCCAGTTGAAGGAGATTCAATGGCTCGGCGAGAGCTTCGCCATACGCCATTTCTTCGGAGATGTGGAGCAATTAGTACCGGAGATTTCCGCAGACTTCACTCTTTCGCATCCGAATGTAATGCATTTTCTGTGCGGTTTCGCGGACGAGGAGAAGAAGGAGTGCTACTTAGTGATGGAGCTGATGAACAGAGATTTATCGAGTTACATTAAAGAGACGTGCGGGCCGAGGAAACGGATCCCGTTTCCGCTACCGGTTGCGGTTGATCTAATGCTGCAGATAGCCAGGGGAATGGAGTATCTGCATTCGAAGAAAATCTATCACGGTGAATTGAATCCTTCGAATATTTTAATCAAGAGCAGGAGCATTGCAGGTACAGAGGGTTACTTACATGCTAGGGTTTCGGGGTTCGGGTTATCTTCGTCGATTAGTAGCCTCACGGAGAAGACTCCGGTGAGCAGGCCGTTTATTTGGTACTCGCCGGAGGTGCTGGCGGAGCAGGACGAGTGGTTTGGCGGAGACGATAGAATCAGCTTGAAATACACAGAGAAATCAGATGTGTACAGTTTTGGGATGATTTGCTTCGAGGTTTTGACGGGGAAAGTTCCGTTTGAAGATACACATCTGCAGGGGGAGAAGATGAGCCGGAATATCAGGGCAGGGGAGCGGCCGCTTTTTCCGTTCCATTCGCCGAAGTATATCACGAATCTGACGAAGAAATGCTGGGCGACGGATCCGAATCAACGGCCGGGGTTTTCGTCGATTTGCCGGATTCTTCGCTACGCGAAGAGGTTTTTGGCGATGAATCCAGAGCACAGCCAGATCGACGCGCCGATGCCGCCGATTGATTTTGGGGAGATTGAATTGGGGATTGTGAGGGGGTTTCCGTTGATCGGGAACTCGAATTTGGTGTCGATTTGGCAGATTCCGTTTCAGATGTTTGCGTATAGGGTTACGGAGAGGGAGAGGGCTAACGTGACGAGCCAGAGGGAGAACTCTGAATCGGGAAGCGATGGGGCTTCCACTTGCGACGAGAGCGTCGTTCCGGTGGAGGATCCGTTCACTTCGCCGCCGGAGAGGAAATCGTCGCTGCCTTCGCCGGAGATGATGATCAGGAAACTCTCTATGTCGAAGAAATCGATGGAAGTCAGAACCCCCAAACAGCCAGGTACGCCGAAGGGAAGATCGAGACCGCCGCATACTCCGACGGGAAGAGCGATGAGGATGAATTCAGAGAGTCAGCTAACGATGATGACAATGAGTCCACGAACACGGAGAAGATCCGGCCACGCCTCCGATTCTGAACTGCCCTAG

>M_truncatula _1

ATGGAACAATTTAGACAAATTGGTGAGGCACTAGGAAGTTTGAAAGCTGTGATGGTGTTCAGAGAGAACATTCAAATCAATCAGAGACAATGTTGTCTTCTTCTAGATGTATTCAGCTTTGCATATGAATCTATTGCAGATGAGATCATACAGAATCTGAAATTTGAAGAGAAGAATGGAAAGTGGAAAGTTCTAGAACACCCCTTAAGAGAGATCCATAAGATTTTCAAAGAAGGAGAAAATTACATTAGGCATTGTTTGGAAATAAAGGATTTTTGGGCTAAAGCCATTACCTTGTGTCACAACACAGATTGTGTTGAGTTTTACATACACAATTTGCTATGTTGCATGCCTATTGTGATCGAAGCTATTGAATCGGCTTCCGAGACATCAGGTTGGGATCAAGATGAGATGCAAAGGAAGAGGCTTATCAATTCCAATAAGTATAGAAAAGAATTTAGAGACATGAAGCTTTTCAAGTGGAAATTTGGGAAGCAGTACCTCATTACTCAGGATTTGTGCAACCGCTATGATACGGCTTGGAAGGAAGATAGATGGCTTCTTTTCAACAAAATCCATGAAAAGAAAGTATCAGATGCTGCAACAAAGTATGAGAAGAAATTAATAGATTTACTTTCGAGGAATTCAGAAGTATCAGAATCACTCGAAGCTAAGCTTCTTTCAAGTTCAATATTGGTTTGTTCTAAGGACTATCAGGTGAGGAGAAGAATAGGGAATGGAAGTCAGTACAAAGAGATTCAATGGTTAGGTGAATATTTTGTTTTAAGACAATGTTCTGGTGACACTGATGCTTTGGAAAGTGAGATTAAAGAACTTTTATCTCTTTCACATCCAAATATAATGGACTGTCTTTGTGGTTTTACTGATGAGGAGAAGAAAGAATGTTTTCTGTTGATGGAACTGATGAGCAAAACACTTAGCAATCATATTAAAGAGGTTTATGGTCCGAGGAAGCGATTACCATTCTTGCTTCATGTAGCGGTTGATATTATGCTTCAGATTGCAAGAGGAATGGAATATATTCATTCAAAGAAAGTGTATCATGGAGAACTAAACCCTTCAAACATTCTTGTTAAGCCTAGAAGTACCTCTCCAGAAGGTTACTTGCATTGCAAGGTATCAGGTTTCGGTCTACCTTCTGTTAAGGACTTGAACCAGAAAGGGAATGCAAATCAAAATGGAACCCTCTCATTCATTTGGTACTCTCCAGAGGTACTTGAAGAGCAAGAACACTCAGGGGGTGTGTCGATATCCAAGTACACGGAGAAATCTGATGTGTACAGTTTTGGAATGGTTTGCTTTGAGCTTCTAACTGGAAAAGTTCCTTTTGAAGATAGTCATCTTCAAGGGGAGAAAATGAGCAGAAACATAAGGGCAGGAGAGAGGCCACTTTTTCCACTCAATTCACCGAAATATGTTATCAACTTAACAAAGAGATGTTGGCATACCGATCCAAATCAACGTCCGAATTTTTCGTCCATATGTAGAGTTCTTCGTTATGTAAAAAGATTTCTTGTATTGAATCCTGGTTACAATAGAGAGACAGACCCGCCAGTGCCACTTGTAGATTACTGTGATATAGAGTCAGCACTTTTGAGGAAGTTTCCTTCTTGGGGAAGTTCTGAATTGTCGCCAATATCTAATATTCCTTTTCAAATGTTTGCTTATCGAGTTATTGAGCGTGAAAAAGTAAGAACATGCTCTAGGGATTTCTCTGAATCTGGAAGTGATGCTTCGGCATGTGGTGATGAGCTTGTTACTTCTGGTGACGAGCCATTTCCATCCGTAACAGAAAAGAAATCTTCACTTGCACCTGAGATTGTAAACAACAGGAGGCTTACGACCAGGAAATCTCTAGATATGAGGATCAAACAACCAGTTACACCGAAAGGGAGACTAGATAGACCTCCACAAATGTCCCCTCGATTGAGAAGCATGCGGATTAATTCAGCTAAACATCTACTATCAACTCCAAGGATAATAAGAAGATCATCTTCTGGTCATGTCTCAGACTCTGAGCTAGCTTAG

>M_truncatula_2

ATGGAACAGCTCCGGCACATTGGTGAGGTTCTTGGAAGTCTAAAAGCGCTAATGGTTTTAAGCGACGAAATTCAAATCAATCAACGACAATGTTGTTTGCTCCACGACATATTTACTCTAGCATTCGATACAATCGCAGATGAGATAAGGCAGAATCTGAAGCTTGAAGACAGAACTACAAAATGGAAGCCTCTTGAATGTCCTCTTAGAGAGCTATCAAGGGTTTTCAAAGAAGGCGAACTATACATTAAGCATTGTTTGGATTCAAAAGATTGGTCAGGAAAAGCACTCACACTTTCTCAGAACAGAGAATGTGTTGAGTTTCACATTCACAACTTGCTTTGCTATTTCCCTGCGGTTATTGAAGCGATTGAAAATGCCGGGGAGATGTCAGGGTTGGATCAGGATGAGAATTCGAAGAAGAAATTGATGCTTGCAAGAAAATATGATGTGGAATGGAATGATCCAAAACTTTTTCAGTGGAGATTTGGCAAACAATATTTGGTTCCAAAAGAAATTTGCAAGCAATTGGAGAATGCTTGGAGAGAAGATAGGTGGAGGCTCATAGAAGCACTCAGAGAGAAAAAAGGTTCTTCCAAACTCACTTATTCAAAGAATGATCAACATTTTGCAGATATGTTGTTGAAGAAATTGATGAATGGTAATTCAGATAAAACTAATCACAAAAACAGTAATGATAAAGTATTGTGGCCAATTGGTGTTTTGTTAGGATCCAAAGATTACCAAGTGAGGAGAAGATTGGGAAGGGGGAAGGAATTTAAGGAAATTCAATGGTTAGGACAAAGTTTTGCTCTGAGGCATTTTCTTGGGGAGAGAGAAGCTTATGAAAATGAGATTTCCAATTTGTTATCACTTTCTCACCCAAACATATTGCAATATCTATGTGGTTTTTATGACGAAGAGAAGAAAGAGTTTTCTCTCGTGACGGAGTTGATGAACAAGGATTTGTGGACATACATGAGAGAGAATTGTGGTCCAAGGAGACAGATTTTGTTCTCTATACCTGTAGTGGTTGATCTCATGCTTCAAATGGCTAGAGGCATTGAATATCTCCACTCCAAAAACATTTACCATGGAAACCTTAATCCATGTAATATCCTACTAAGAGCTAGAAACTCACAAGAAGGTTATTTTCAAGCAAAAGTTGTTGGATTTGGGTTGTCATCTGTTGAAAATGGTGAAATTTATAATGCTTCTAGATCCTCACCGACACATAATCCAATTGGTGAAGAGATTAACCCTATAATTTGGTATGCACCTGAGGTTTTAACTGAGCTAGAACAAATGAAAAATGCTTTTACCTCTTGTAAGTACTCAGAGAAAGCCGATGCATATAGCTTTGGTATGATATGTTTTGAGTTGTTGACAGGGAAAATTCCTTTTGAGGACAATCATCTTCAAGGGATTAGAACCAACCAAAATATCAAAGCAGGAGAGAGACCTTTGTTTCCATATCGTTCGCCAAAATATCTTGTGAGTTTGATCAAAAAATGTTGGCAAACTGATCCGTCTCAGCGTCCTAGTTTCTCGTCTATTTGTAGGATTTTGCGCTACATCAAAAAGTTTCTTTCAATGAACACTGAGTATGTTTTGATCAATCCTGAACTAAATCAGCTTGAACTTCAGAACCCACCTGTAGATTGTTGTGACATTGAAGGAATTTTTCTCAAGAGTTTTCCAATGGAGAGGACATATAATATGTCATTTGTTTCACAAATACCTTATGAAATGTTTGCTTATAAAGTTGTTGAAAAAGGGAAGATTAATAATCAGAATATGAGTAATATTACTATTCCAACTATAGATAATAAGGGTAGTGAGTCTGAGCCAACAAAGGATGTGGCTACTGTTCGTAGCGAAGAAAGTGATGACCAAAGCACAGTTTTTGAAGATATTGATGCATCCATAATTAAAGATCTGGTAACCTATCCAAAATCAATTTCTGGTGACACACAGTCTATTTTTTATGATGCCCCATCAAAGAAGAAAGTTACAATAAAGAAACCATCACAAATGAAGCCAAAGAAAGACCAAGGAACGCCTAAACTGCAAGCAACAAAATCATTACCACCAACATTATCTGGTCGGGTTTCAAGGACAAATAAGGTGAACCAGTCATCATTGACATCAAGTTTTATAACCCCAGTTAAAAGAAGGCCTCCAAAAGTCTCAGAATCTGAGAGGAAATCCAATATTAACAAAGGGAATCAGTCAGCATCAGTGTCAACTTCAAGCTCAGTTAGAAAAAAAATACATGGAGGCAATGTCTCCGATTCTAAGCCTAGTTTAAATCTGAAAACAAGAGACCAATCACCATTTAACAGGTTAAAGACAAAAAAATCACAACTAACGATGAATTCTTCTTCGAGTCCAACACGAATAAGACGTGTCAATAGTACTAACAACAGCATTGTGAGGATGCAAAAAGGGTTCATTTCAACACCTCCAATGAGTCCTTCAAGGTCATACGCCAGAAGATGCTGCCATGCTTCGGATATGGATAGTTCTTTGAAGATACGAGGTAGATTGTCACCATTTCCATTGAGTCCTTTGAGTCCATATGTGAGCAATCAAAGGAATAGAAGAGGAAGCTTGTCACCATTAGCATTGAGCCCTTTGAGTCCTTATGTAAGAACAACATATGGTCATGTCTCAGATTAA

> O_sativa_1

ATGGAGCAGCTCCGGCAACTCGGCGAGGCGGTGGGTAGCATCAACGCGCTCATGGCGTTCGAGGACGACCTCCACATCAACCCGCGGCAGTGCCGCCTCCTCGCCGACGCCTGCGCGCGCGCGCTCGCCGCCGTCACGGGGCAGGTTCGCGCCCAGCTCCGCTTCGACGAGCGCGGCGCCAAGTGGCGCGCCATCGAGGCGCCGCTCCGCGAGCTCCACCGCGCCTTCCGCGACGCCGAGGCGTACGTCAGGCAGTGCCTGGACCCCCGCGGCAGCTGGTGGGCGCGCGCCGCGGCCATGGCGCACGGCACCGAGTGCGTGGAGCAGCACCTCCACAATGTCCTGTGGTGCGTCGCCGTCGCGCTCGAGGCCATCGACGCCGCCGGCGAGATCGCCGGCTCCGACCCGGACGAGCTCGCCAGGGGGCGGCTGGTGCTCGCCAGGAAGTACGACAGGGACATGCTCGATCCCAAGCTGTTCGAGCACGCGTTCGGCAAGCTATATCTGGTCTCCCAAGAGCTCGTTGCAAGGATGGACATGGCGTGGAAGGAGGACAGATGGGTGATATCGCAAATGTTCGACGAAATGAAAGGCCCTGCGGCGTCGAAACCTCTGAGCAAAAACGAGCACCGCCTCGCCGAGCTCCTCGCCGCGGCGATGGGGAAGCTGCACCCGGCATCTGTCCTTCTTGGCAGCGACTACAGCGTCCGGAGGCGGCTCGGCGGCCGGCTCAAGGAGGTGCATTGGATGGGGGAGAGCTTTGCGATGAAGCATTTCATCGGGGACACCGATGCCGCCGGCGCCGAGGTCGCGCTCCTGTGTTCGGTGGCGCACCCGAACGTCGCGCACGCCGCGTACTGCTTCCACGACGAGGAGAAGAAAGAGTACTTCGTGGTCATGGACCAGCTCATGGCCAAGGACCTCGGGAGCTACGTCAAGGAGGTGAGCTGCCCGCGGCGGCGGATCCCGTTCCCGCTCGTCGTCGCCGTCGACATTATGCTGCAGATCGCGCGCGGGATGGAGTATCTGCACGCGAAGAGGATAAACCACGGCGAGCTGAACCCTTCCAACGTGCTCGTCAAGCCGCGCCAACCCGATGGCGGCTACGTGCATGTCAAGGTCGCCGGATATGGGCAGCCAGCTGGCATCACAGCCGGCGGCGCAAAAGCTTCCGCCAATGGCAACGCCAACGGCAACGACAACTCCTGCATCTGGTACGCGCCGGAGGTGCTCAGGTCCGACGGCGTCGCGGACGCCGCGGCGGCGGGGAGGTGCACCGAGAAGGCCGACGTGTACAGCTTCGCCATGATCTGCTTCGAGCTGCTGACCGGCAAGGTGCCGTTCGAGGACAACCACCTGCAGGGCGACAAGACGAGCAAGAACATCTGCGCCGGCGAGCGGCCGCTGTTCCCGTTCCAGGCGCCCAAGTACCTCACCGCGCTGACGAAGCGGTGCTGGCACGCCGACCCGGCGCAGCGGCTGGCGTTCGCCTCCATCTGCCGCGTCCTCCGGTACGTCAAGCGGTTCCTGATCCTGAACCCGGAGCAGCAGCAGCAGCAGCAGCAGGGCCAGACCGACGACGCGCCCAAGCCGGCCGTCGACTACCTCGACATCGAGGCGCAGCTGCTGAAGAAGCTCCCGGCGTGGCAGCGTGGCGGCGAGGCGCCGCGCGTCGCCGACGTGCCGTTCCAGATGTTCGCGTACAGGGTCATGGAGAGGGAGAAGGCCGCCGGCGCCGTGCACGTCGCCAAGGACAGGGCCTCGGATTCGGGCAGCGACGGGAACTCGCTGTACGGCGACGAGAACGGGTTCGGTGCAATGTCGCCGGAGCACACCTTCTCCGCCGTGTCGAACGGCACCCTGCGGTCGCGGCCGGCCAGCAGCGACGGCAGGTTGCCGACGGCCAAGAAGGCGGACGGCAAGGCGCCCAGGCAAGCAGGGCCACAGCCGAAGGTGAAACCCGTGAACACGGCGGCGAGGACTCCGCACTCGGCAAGGCGGGCGCTCGGCGTGAAGCCCGACGATCACCTCCAGACCAACGGCGCTCCCACCGCCCGCCGGAGGACGCCCGAGATGGCCTCAGAATAG

> O_sativa_2

ATGGAGCAGCTCCGGCAGGTCGGCGAGGCCATCGGCGGCGTCAACGCCCTCATGGCCTTCCACGACGACCTCCGCTGCATCAACCCTCGCCAATGCGCCCTCCTCGCCCACGCCTACGCCCTCGCCTTCCGCGCCGTCGCCGGCGAGCTCCGCGCCCGCCTCCGCTTCCACGACCGCCTCACCAAATGGAAGCCCCTCGACGATCCCCTCCGCGAGCTCCACCGCGTCGTCCGCGACGGCGAGGCCTACATCCGCCACTGCCTCCTCCTCGACCCGGCCCACTGGTGGGCGCGCGCCGCCGCCGCCACGCACGGCACGGAATGCGTCGAGCACCACCTCCACAACCTCCTTTGGTGCGTCTCCGTCGTCGTCGAGGCGGTCGAGAACGTCGGCGAGGTCACCGGCTCCGACCCCGACGAGCTCGCCCGGCGGCGGCTGGCGCTGGCCAGGGACTACGACAAGGACCTGCTCGATCCAAAGCTCTTCCGGGAGAGGCTCGGCGAGACGTTCCTCGCCACGCGTGAGCTCGCCGCCCGGATGGACATGGCGTGGAAGGAGGACAGGTGGCTGCTGTCCCAGCTTCTCGACGAGAGGAAGGGCCCGACATCGTCGCCGGAGCCGCCATTGACGCGGCAGGAGCACCGGCTCGCCGACCTCCTCGCCGCGCCGCGCGGGAAGCTGCACCCGGCGAGCGTGCTCCTGATGAGCGACTTCCACATGCGGAGGCGCCTCGGGGGGAACGGGAACCTCAAGGAGGTGCAGTGGTTGGGGGAGGCGTTCGCTGTGAAGCACGTCGTCGGCGTCGACGCCGAGGCGGCGGCCGCGGAGGTGGCGGCGCTGGCGTCGGTGTCGCCGCACCCGAACGTGGCGCACTGCCGCTACTGCTTCCACGACGAGGAGAAGAGGGAGCTGTACATGGTGATGGACCAGCTGATGAGCAAGGACCTCGGCAGCTACGTCAAGGAGGTGAACTCCGCCAAGCGCCGGGCGCCGCTCCCGCTCGTCGTCGTCGTCGACACCATGCTGCAGATCGCGTGCGGCATGGCGCACCTGCACTCCAACAAGATGTACCACGGCAACCTCAACCCATCCAACGTGATCGTCAAGCCGCGCCATGGCGACGCCTACCTGCATGTCAAGGTCGCCGGCTTCGTCTCCGGCTCCGGCACGGCGAACGCGGCCAACCCATGCATCTGGTGCGCGCCGGAGGTGGTGGGGAACGAGGCGGCGGCGACGGAGAAGGGCGACGTGTACAGCTTCGGGATGATCTGCTTCGAGCTGATCACCGGGAAGATCCCGTTCGAGGACAACCACCTGCAGGGGGAGAACATGAGCAAGAACATCCGCGCCGGCGAGCGGCCGCTGTTCCCGTTCCAGTCGCCCAAGTACCTGACCAGCCTGACGCGGCGGTGCTGGCACGGCGAGGCGGCGCAGCGGCCGCCGTTCCACTCCATCTGCCGCGTGCTCCGCTACGTGAAGCGCTTCCTGGTGATGAATAACCCGGAGCAGGCGGCGGCGGACGCGGCTGGAGCCGGGCCGGCGGTGGACTACCTGGACATGGAGGCGCAGCTGCTGAGGAGGTTCCCGGAGTGGGAGGGGAACGGTGTCGCCGACGTGCCGTTCGAGATGTACGCGTACAGGGTGATGGAGAGGGACAAGATGAGCAACGCGTGCAGGGACAGGAGCTCCGACTCCGGCAGCGACGGCAACTCGCTGTGGGGAGATGACAGCGCCAGCGGCGGGTCCAGCACGACGGCGACGGACGCGTCGGCGTCGAGCCGGCCGCTGCTTGACCGGAGCGGCAGCACGAGGTCGTCGCCGCCGCGCAAGGTGGCCATCGCCGCCGCCAAGGCAGGCAAGTGTCGCAGCGGCATTGTTACTCGTCTAAAGCCTTCCTCCAAGATTACAGCTAGCTCCATGTCCGTGACGTGTGCAGGGCCGCCGCAGAAGTCGAGGTCGATGGGCACGGTGAGGCCGCCGCCGGTCGTGGCCCGCCGAACGCCGAGGATTAAGTCCGACGGCCACCTCAATAGGGCGGCAATTCCGCCTACCCGACGGCGTAAATCCGGCGGGAACGCTTCAGACTCGGAGTTAGCTTAG

>P_dactylifera_1

ATGGAGCAGTTTCGGCAGATTGGTGAAGTCCTGGGAAGCCTCAAAGCTCTGATGGTGTTCCGGGACGAGATACACATGAACCGGCGCCAGTGCTGCCTTCTGGTGGATGCCTTCAATGCTGCCTTTGAGGGGATTGGCGATGAGATCCGGCACCACCTGAAATTCGAGGAGAAGGTCACCAAGTGGAAGGCCCTCGAGCAACCCTTCAAGGAGCTCCACCGGATCTTCAGGGAGGGGGAGCAATACATCAGGCAGTGCCTCGAGCCCAAGGACTGGTGGGGGAAAGCAATTGCCCTCAGTCAGAGCACTGATTGTGTCGAATTCCATCTCCACAACTTGCTATGGTGCATCCCTGTTGTGCTCGAGGCGATCGAGAGAGTCGGAGAAATGTCAGGGTCAGATCATGAAGAGATCCACAAGAAGAGGGTGGTGTTCTCGAAGAAGTACGAGAGAGAGTGGATGGAGCCGAAGCTCTTTCAGCACAAGTTTGGAAAATTGTACCTGGTTTCACAGGACGTGTGTAGTAGGTTGGACAGTGCATGGAAAGAGGATAGGTGGATTTTATCTGAAATGATAGAGGAGAAGAGGAGTTCAGGGTTGAAACTGATGACAAAGCAAGAGCAGCGGCTCGCAGAGCTACTGATGGGTCCAAAAGGGAAGCTTTTCGCAACCTCGGTCCTTGTGGGATCGCAGGACTACTACCAAGTGAAGAGGCGATTCGGGTCAAGCAACTACAAGGAAGTCCAGTGGATGGGGGAGAGTTTTGCTGTGAAGCATGTCATTGGAGAGATTGAGCCACTAATGAATGAGATCTCTCTCTTGACATCGATCAACCACCCAAATGTGATGCATTGTATGTACTCATTCTACGATGAAGAGAAGAAAGAGGGATTTCTAGTTTTGGATCTCATGAGCAAGGATCTCTCCGATCATATCAAAGAGATTTGTAGCTCTAGGAGAAGAATCCCCTTCCCTCTGCTGGTGGCAGTGGACGTAATGCTTCAGATTGCAAGAGGAATGGAGTACCTGCATTCAAGAAAAATACACCATGGGGACTTGAACCCTTCCAATGTCTTAGTTAGGACAAGGAGTTACTCTCCTGATGGTTATTTGCATGTAAAGATTACTGGATTTGGTCTGTCACCTTTGAAGAAGTCAAAAGTTTCAGCAAACCAATCAGCTACCATGAATTCATGTATCTGGTGTGCCCCAGAGGTTCTCTTAGAGCAGGACCTGTCAGGGGAAGCTGGTAGTAGTAAGTACACAGAGAAAGCAGATGTTTATAGCTTTGGAATGATATGCTTCGAGTTGCTGACAGGAAAAATTCCTTTTGAAGACAATCATCTTCAGGGGGATAAGATGAGCAAGAACATAAGGGCGGGTGAAAGGCCACTGTTTCCATTTCAATGTCCAAAGTATCTCACCAGCCTAACAAAAAGATGCTGGCACGCCAATCCCTCTCAACGGCCAAGCTTCTCTTCTCTTTGTAGAGTGCTCCGTTACACCAAGCGCTTTCTAATTATGAACCCCAATCATGACCAGCCTGATCCAGTGGTACCTCCAGTGGATTACTTTGATATTGAAACGAGTCTATCAAAGAGATTCACAAACTGGGCAAGAAAAGAAACCATCCAGGTCTCAGAAATTCCTTTTCAGATGTATGCCTATAGGGTTATAGAGAGGGAGAAAACTAGTCTTAATGTCAAGGATAAAAGTTCGGAATCGGGAAGCGAGGAGGCATCGATTTGTGAAGATGAGAATGCATTCGGACAGCACCAAAAAGGGAAAATAGTGAAGCCACCGCAACCAAGTTGTGGGCGCGGTTTGCGGATGAGCTCTGAGAGCCAGATGCCAACGATAGCAATTAGCCCCGGACGGAGGACATCCGGGCATGTCTCAGATTGA

> P_dactylifera_2

ATGGAGCAGTTCCGTCAGGTAGGAGGGGTTCTGGGAAGCCTCAAGGCTCAGATGGTGTTTAAAGATGAGATACAAATCAATCAGCGCCAGTGCTGTTTGTTGGTGGATGCTTTCAGCCTGGCCTTTGAAGTGGTCGCAGAGGAGATGAAACAGTATCTAAGATTCGAAGAGAGATTCACAAAGTGGAACGCACTGGAGAGCCCCCTCAGAGAACTTTACAGGGTATTCAAGGAAGGAGAGCACTATGTCCGACAATGCTTGGAGACAAAGGATTGGTGGGCTAAAGCCATTTGCCTCAATCAGAACAGTGATTGCATCGATTTCCATCTGCACAATTTGTTATGGTGCATCTATGTCGTGCTTGAGGCAATCGAGAATGTCGGAGAGATATCAGGTTGTGATCAAGAAGTGATCCAGAAGAAAAGGCTTGTCTTCATGAAGAAGTATGAGCGAGAATGGATGGATCCACACCTTTTTCAGCACAGGTTCGGAAAGAAGTACTTGGTTTCGCAGGACATATGCAGCAGACTGGGTAGTGTCCGGAAGGAGGATCGGTGGATTCTCTCAGAAATGATGTCTGAGCAGAGAAACTCGGGTTCCACGCCTCTAGCAAAGCAAGAGAGACAGCTTGCTGAGCTTCTCTTGGCTCCGAAAGGGAAGCTCTTCCCGAGTTCAGTCCTGGTGAGGTCGAAGGACTACCAGGTAAGAAGACGATTGGGGAGAAGAGGCCCTTACAAGGAGGTCCAATGGATGGGGGAGAGCTTCGCCGTCGGGCATTTCTTCGGGGACATCGAGCCACTGAAGCAAGAGATCTCGCTCGTGTCATCGCTCAGGCACCCACATATAATGCACATCATGTATGCTTTCTCTGACGAGGAGCGGAAGGAATTGTTTCTGGTTATGGAGCTCATGAGCAAGGATCTCTCTAGTTATATCAAAGAGATATGCTCCACAAAGAGAAGGCTCCCCTTTCCTCTGCTGGTGGCAGTGGACATAATGCTACAAATTGCAAGAGGAATGGAGTATCTCCACTCTCGTGGGATATATCATGGAGATTTAAACCCGTCCAACATCTTAGTAAAGCCGAGAAATTCTAATCCAGATGGCCACCTGCATAAGCCGAGAGACTCGACAAATCAAACAGCTACAGATTCATGCATATGGTATGCACCCGAGGTTCTTTCGGAGCAAGAACAGTCCGCAGAGCCTAGTGTTTCCAAATGCACAGAGAAGGCAGACGTCTACAGTTTTGCAATGATTTGTTTTGAGGTGCTAACTGGAAAAATACCTTTCGAGGATGATCACCTTCAAGGAGATAAGATGAGCAGAAACATACGGGCAGGCGAGAGGCCGCTATTCCCATTGCCATCTCCAAAATACCTCACTAACTTGACCAAAAGATGCTGGCAGGCCGACCCCTCCCAACGACCGAGCTTCTCTTCTATCTGTAGAGTTCTTCGTTACATCAAGAGGTTTTTAATTATGAATCCGGATCGTGGCCAGCCTGACCTACCAGCTCCACCGCCTGACCTACCAGCTCCACCGGTAGACTACTTTGATCTCGAAGCGACACTGTCCAAGGGATTCACAACATGGGCAACTGCAGAGGCACTAACAGTCTCAGAAGTCCCCTTTCAGATGTTTGCTTATAGAGTTATGGAGAGGGAGAGGACTAATCTGAACTTTAGGGAGAGGAGTTCAGAGTCAGGGAGTGAGGGGGCAACATCGATATGGCAGCAGCTAAAAGGAGCAGTAAGACCACCACAAAAAACAGTACATGGATACAATATGAGGATGCAATCGGAAACCCACTTGCCGACAGTCGTCATGAGCCCGAGACGGAGGTTTTCAGGGCATGCATCAGATTCAGAGTTAGCATAG

> P_dactylifera_3

ATGGAGCAGTTTCGGCAGATTGGAGAAGTTCTGGGGAGCCTCAAGGCTCTGATGGTGTTCCGGGAGGAGATTCGCATGAACCAGCGCCAGTGCTGCCTGCTGGTGGATGCTTTCAACGCCGCCTTCGAGGGGATCGCCGACGAGATCCGACACCACCTGAAATTCGAGGAGAAGCTCACTAAGTGGAAGGGCCTCGAGCAACCCTTCAAGGAGCTCCACCGCATCTTCCGGGAGGGGGAGCAGTACATCAAGCAGTGCCTCGAGCCAAAGGACTGGTGGGGGAAAGCCATTGCCCTCAGTCAGAGCACAGACTGCGTCGAATTCCATCTCCACAACCTTCTGTGGTGCATCCCTGTCGTGCTAGAGGCGATCGAGAATGTCGGAGAAATGACGGGGTGCGATCAAGAAGAGATCCATCAGAAGAGGATTGTGTTCTCAAAGAAGTATGAGAAAGAGTGGGTGGAACCGAAGCTCTTTCAGCACAAGTTTGGTAAATTATACCTAGTTTCACAAGACGTTTGCAGTAGGATGGACGGCGCTTGGAAGGAGGATAGGTGGATTTTATCTGAAATGATAGCAGAGAAGAGAAGTGCAGGGTCGAGACCGATGGCAAAGCAAGAGAATCGGCTTGCAGAGCTACTGATGGGTCCCAAAGGGAAGCTTTTCGCAGCTTCGATCCTTGTGGGATCACAGGACTACCAAGTGAAAAGGCGATTCGGGTCGAGCAACTACAAGGAAGTCCAGTGGATGGGGGAGAGCTTTGCTGTGAAGCATGTCATTGGAGAGATTGAGCCACTGATGAATGAAATCTCTCTGCTATCGTCGATCAATCACCCGAACGTGATGCATTATATGTATTCATTCTATGATGAAGAGAAGAAGGAGGCTTTTCTGGTTACAGATCTCATGAGCAAGGATCTCTCCAGTCATATCAAAGAGATTTGTAGCTCGAGGAGAAGAATCCCCTTCCCTCTGCTGGTGGCAGTGGACATAATGCTTCAGATTGCAAGAGGAATGGAGTATCTCCATTCAAGGAAGATATATCATGGGGACTTGAACCTCTCCAATGTCTTGGTCAGGGCAAGGAATTCTTCTCCTGATGGCTATTTGCATGTAAAGATTGCTGGCTTTGGGCTGTCACCTTTGAAGAACTCAAAGGCTTCAGGAAACCAAGCAGCTGCCATGAATTCATGTATCTGGTATGCCCCAGAGGTTCTCTTAGAGCAGGAGCAGTCGGGGGATGCTGGTGGTAGTAAGTACACAGAGAAAGCAGATGTTTATAGCTTTGCCATGATATGCTTCGAGGTGCTGACAGGAAAAGTTCCCTTCGAAGACAATCACCTTCAGGGGGATAAGATGAGCAAGAACATAAGGGCTGGTGAGAGGCCATTGTTTCCATTTCAATCTCCGAAGTATCTCACCAACCTAACAAAAAGATGCTGGCATGCCGATCCCTCCCAGCGGCCAAGCTTCTCTTCTCTTTGTAGAGTGCTCCGTTATGTCAAGCGATTTCTAATCTTGAATCCTGATCATGGCCAGCCTGATTCAGTGGTGGTACCTCCAGTGGATTACTTTGATATTGACACGAGTCTCTCTAAGAGATTCACAAACTGGGCAAGAAAAGAAACCATCCAGGTCTCCGAAATTCCTTTTCAGATGTATGCTTACAGGGTTGTAGACAGGGAGAAAACTAGTGCTAATGTCAAGGATAAAAGTTCAGAGTCGGGAAGACAGCATCAAAAAGGGAGAACATTGAAACCACCGCAGCAAAATTGTGGGCGCAGTTTGCGGATGAACTCTGAGAGCCAGCTGCCTACAGTCGCAATGAGCCGAAGGCGAAGGATGTCTGGCCATGTCTCAGATTGA

>P_persica_1

ATGGAGCAATTTCGAAAGATTGGGGAGGATTTGGGAAGTTTAAAGGCCTTGATGGTGTTCCAAGACAACATCCAAATCAATCAGCGGCAGTGTTTTTTGCTGCTTGATATCTTCAGCTCAGCATATGAATCAATAGCAGAGGAGATGAAATATAATCTGAGATTTGAAGAGAAGCACACTAAATGGAGGGTTCTCGAGCAGCCGTTGAGAGAGCTCCACCGGATATTCAAAGAAGGAGAAGCCTACATCAGGCACTGCTTGGAAACAAAGGATTGGTGGGCTAAAGCCATTACTCTCTATCAGAACTCCGATTGCATTGAGTTTCACATCCATAACTTACTTTCCTGCATGCCGATTGTCATTGAAGCAATTGACATTGCTGGAGAAATATCAGGGTGGGGTCAGGATGAGATACAAAGGAAGAAAACCGTGTTCGCGGATAAGTACAAAGATGACTATAGAGATTGGAAACTTTTCAAGTGGAGATTTGGGAAGCAGTATTTGATTACTCAAGATTTCTGCAATAGGTTTGACACAACATGGAAAGAAGATAGATGGACTCTTCTCCATAAAATCAGAGAAAAGAAACTCTCGGGTTCGACAAAGTACGGGAAACGCCTCATAGATCTTCTTTTCAAAAGCTTAGATGGATCAGAGTCACAGCCTTTGAATGGAAAACTCTTACCTAGTTCAATCCTGGTGGGGTCGAAAGACTACCAGGTGAGGCGACGGCTAGGGGGTGGAAGTCAGTATAAAGAGATCCTATGGCTGGGTGAAAGCTTTGCCTCAAGACACTTCTTCGGGGATATTGAACCCCTGCTGCCAGAGATTTCTTCATTGTTATCTCTGTCCCACCCCAACATAGTGCACTTCCTTTGCGGGTTTACGGATGAGGAAAAGAAAGAGTGTTTTCTCATTATGGAACTGATGAGCAGAGACCTCTGCAGCCACATCAAAGAGATTTGTGGCCCACGAAAACGACTTCCATTTTCTCTTCCTGTTGCAGTTGATCTGATGCTTCAAATTGCAAGAGGAATGGAATATCTCCACTCGAAGAAAATCTACCACGGAGAATTGAATCCTTGTAACATACTTGTTAAAGCAAGAGGCATCTCCACAGATGGGTACTTGCAAGCCAAGGTTTCAGGTTTTGGTTTAACTTCTGCGAAAAGCCCCAGCCAAAAAAACCTTTCAAATCAGAATGGAAGCCTACCTTTCATATGGTATGCTCCGGAAGTTCTGGAAGAGCACGAACAGACAAAAAGTAGTGAGAAAAAGTACACAGAAAAGTCTGATGTGTATAGCTTTGGCATGGTTTGCTTTGAGCTTCTGACAGGCAAAGTGCCTTTTGAGGATAGTCATCTTCAAGGAGAGAAGATGAGCCGAAACATAAGGGCAGGAGAGAGGCCGTTGTTCCCATTCTATTCTCTGAGATATGTGACCAACCTCACAAAGAAATGTTGGCACAGTGACCCTAATCTACGGCCAAGCTTCTCATCCATCTGTCGGATTCTAAGATATATTAAAAGGTTCCTTGCAATGAATCCTGATTATAACAGCCAGCTAGACCCACCAGTGCCTATGGTAGATTACTGTGAGATTGAGTCAGGGATTCTGAGAAAGATTCCTTCTTGGAAGAGTTGTGAGCAACCACCAGTATCACAAATCCCATTTCAGATGTTTGTTTATAGAATTGCGGAGAGAGAAAGAACAACGTTGAAAGATACTTCAGAATCAGGAAGCGATGGAGCTTCAATCTGTGGAGATGAAATGGTGACCACTCCAGACGAACCATTCCCGCCCGCGCCTGAGAGGAAGTGTTTAACATCGCCTGATAGCATGAAAAAGAAACTTCCATTCTTGAAGAAATCTTCAGATGTGAAAGCCAACAGACTACCAGGCTTTGGTTACACAGGAACACCAAGAGGACGGTCTGTCAGACCACCACAGATGAGCCCTTGTGGCCGTAGTATTAGTCTGAGAATGAGTTCAGAAAGCCAGCTAATGGCGATGACTCCAAGAATACGGAGAACATCATCTGGACATGCCTCAGATTCTGAGCTCTCCTAG

>P_persica_2

ATGGAACAATTCCGGCATATTGGAGAGGTTGTGGGAAGCATGAAGGCTCTGTTGATGGTACAAGATGAGACTCTGATCAATCAATGTCAGTGCTCATTGTTGCATGATGTATTCAGTATGGCCTTTGACACAATTGGAGAGGAGATCAGGCTGAACCTAAAGCTCGAAGAGAAAAAGACCAAATGGAAAGTTCTTGAGAAGCCGTTGAGGGAGCTTCACACAGTTTTCAAAGAAGGGGAGCTCTACATTAGGCACTGCATGGATGCCAGTAATTGGTGGAGCAAAACAATCCTCCTCCATCAGAACAAGGACTCTGTTGAATTTCATGTTCATAATTTGTTCTGCTACTACGCGGCTGTGATTGAGGCAATTGAGAACGCTGGAGAGATTGCTGGACTCGATCAGGATGTGATGCACAAGAAGAGGATTTTGCTCACAAGGAAGTATGATAGGGAGTGGAATGACCCCAAGCTTTTCCAGTGGAGATTTGGGAAGCAGTATTTGGTCCCAAAGGCCATCTGCAAAAGGTTGGAGAGTGCTTGGAGGGAAGATAGATGGCGGCTGATTGAAACGCTCAAGGAGAACAAAATTTCAGGATCCCTTGGTTCGACAAGGAATGAGCAGCAGCTTGGAGACTTGCTGCTGAAGAAGCTACATGGGTCAGATTTCAGTGGGAAGCTGTTTCCAAGCACAATCTTATTAGCATCCAAGGACTACCAAATTAGGCGGCGGTTAGGAGGGGGAAGCCAGTACAAGGAAATCCAGTGGTTGGGGAAAAACTTTGCCTTGAGACACTTCTTCGGGGATCTTGAACCATTGAACTCGGAGATTTCAACTCTCCTCTCACTTTCCCACCCCCATGTACTGCAGTACCTTTGTGGGTTTTATGATGAGGAGAAGAAGGAGTGCTTTCTTGTTATGGAGTTGATGAGCAAGGATCTGCGCTGCTACATGAAGGAGAACCGTGGTGCAAGAAGGCAGGTTTTGTTTTCAATCCCAGTGGTGGTTGATATCATGCTTCAGATTGCAAGAGGCATGGAATATCTCCACTCTAGGAAGATCTACCATGGAGAGTTGAACCCTTGTAATGTCTTTCTCAAGGCAAGGAGCTGCACAGAAGGTTATTTTCAAGTAAAAGTATCAGGTTTTGGTTTATCATCCGTACACAAACCTACTTCACGATACTCACAGCAGCAAAATGAGATCAACCCTTTGATTTGGTGTGCCCCAGAAGTCCTGGCTGAGCAAGAACAACCAGGGAATAACCGTCGTACCAAATACACAGAGAAAGCAGACGTATACAGCTTTGCGATGCTTTGCTTTGAGCTCTTGACTGGGAAGGTTCCTTTTGAAGATTCACATCTCCAAGGGGACAAGATGAGTCGCACTATAAAAGCAGGAGGGAGGCCTCTGTTCCCTTTCCCTTCACCAAAGTACCTTGTCAACTTAACCAAGAGATGCTGGCACACTGACCCATCTCAGCGCCTGACTTTCTCATCCATCTGTCGAATTCTACGCTACATAAAGAATTTTCTTACCTTGAACCCCGACGACGATCAGCCTATACTGCAATCCCCTCCTATGGATTATTGTGAGATAGAGACATGGTTCCTGAAGAATAGTTCAGCAGCAGGGTATGCTGATTTATCCTCAATATCACAACTTCCATTTCAAATGTTTTCTTATAGACTTGGGGAGAAGGAGAAGACTAGGCCCGGCCTCATCAGGATTAAGAGTCTGGATTCTGTAAGTGATACAACTTCGATGAGCAGGCAAGAAGCCCCAAATTGCAGAAATGACAGTGTGTCTGTCGTCGAAGATCCATTTGTACCATTAAGTGATACGAGGTCTGTTTGCTCTGATTTGAGGTCTGTTTTTGATTTGAGGTCAGTATGTTCCGAGGCTCCAACCAAGAAAACTCTCACTGCAAAGAAACGCCCCGGAGTGAGCGCTAGAAAAGGCTTAGGGACATCAAGATCACCAGCAACACCAAGATTTTCGACACCAAGACTTTCGTCACCAAGGCTTCCTCCACCAAAACTTCCATTACCAAAAATTCCACCACCAAAAGTTTCGACGCCAATAAAACCATGTCGTCGCAGCAGCACGAAGATGAACAATGGGAGCAGTCCACTACCGAGTCCTATGAGTACAAAGAGTACTGCAAGCGGCAGACACAGACCGTGTGGTCATGTCTCGGACTCGGAGATACATTAG

>P_richocarpa_1

ATGGTATTGCAAGATGATATTCAGATCAATCAGAAGCAGTGCTGTTTGCTGGTTAATATCTTTTGTTTGGCCTTTAAAACGATAGCGGAGGAGATCAAGCAGAACCTAAAACTAGAAGAGAAGAACACCAAGTGGAGGCCTCTTGAAGAGCCTTTGAAAGAGGTGTATAGGGTATTCAAAGAAGGGGAACTTTATGTTAGGCGTTGCTTGGATAACAAAGATTGGTGGGGAAAAGCTATTAGCCTTCATCAGAATAAGGACTCTGTTGAGTTTCACATCCACAACTTGCTTAGCTGCTTTCCTGCTGTTATCGAGGCGGTTGAGATTGCTGGAGAAATCTCTGGACTTGACCAGGAAGAGATGACAAAGAAGAGGGCAATGCTGGTAAAAAAGTATGATAGGGGTTGGAGTGATCCGAAACTTTTCCAATGGAATTTTGGGAAGCAGTATTTGGTCCCTCGGGAAATTTGTAGGCAGATGGAGAGGGCTATGAGAGAAGATAGATGGCTTCTTATTGATACAATCAAGGAGAAGAGAAGGGCATTGCCACCAGGAAAGAGTGAGCATCGACTTGGAGAACTTTTGCTGAAGAAACTAAATGTACTAGCACCAGCTAATGGCAAACTCCCACCAAGTTCAATTTTGTTAGAAGCAGAAGATTACCAAGTGAGGAGAAGGTTAGGTGGAAATCAGCACAAGGAAATTCAATGGCTAGGGGAGAATTTTGCTTTGAGACATTTCTTTCACGATTTTGAACCGTTGAATTCCGAAATTTCTATGCTTTTGTCCCTTTCCCATCCCAACATAGTGCAGTACCTCTGTGGCTTTTATGATGAAAACAAGAAGGAATGTTTCCTTGTAATGGAATTGATGACCAAAGATTTCTATTCTTACATAAAAGAGAACAGCAGTCCGAAGAAGCGGGTTTTGTTCCCTCTTCCAACTGTAGTTGATATTATGCTTCAGATTGCAAGAGGAATGGAATTTCTCCACTCACGGAAGATCTACGTGGGAGATTTTAATCCTAGTAACGTCCTTCTCAAACCCAGGAAATCTACAGAAGGTTATTTTCATGTGAAAGTCTCAGGGTTTGGCTTAACATCTGTCAAAAATCATTCTTCTCGACACTCATCGCCAGAGCCAAGTCCAGTCGATACTTGCATTTGGCATGCCCCAGAAGTTCTGGCAGAGCAAGAACAGGCAAGAAATGCCAGCAGTAAAAAGTACACAGAGAAAGCAGATGTTTATAGCTTTGGGATGCTTTGTTTCCAGCTCTTGACAGGGAAACTTCCCTTTGAGGATGGGCATCTTCAAGGAGACCAAATGATCAATAATATAAGAGCAGGAGAACGGCCATTATTCCCATCCCTTTCACCAAAATACCTTGTAAGCTTAACCAAGAAATGCTGGCATACGGAGCCAAGTTATCGCCCTACTTTCTTGTCTATTTGTAGGGTTCTGAGATACATCAAGAAGTTCCTTGTGATGAACCCTAACGACGGTCAGCCTTATATACAGTCGCCGCCTGTAGATTTTTATGACTTGGAGGCAGGGTTTCTAAAGAAGTGTCAAGGGGAGGTGACTTGTGATCTGCCTTCAGTGTCACAAATTCCATTTCAAATGTTTTCTTATAGGCTTATAGAGAAAGAAAAAACTTGTGTACAAATTAAATTTAAGAATTCTGAAGCAGCAAGTGAAGCAGCCTCAAATGGCTGGGATGAGAGTAACTCTGTAGTGGAGGATCATCACGTACCTTCAATTGATGCCAGGTCGTTTACATCTGACGTGAAGTCTGTTGGTTTTGATATGAAGTCGACTTTCACTGAAGTTCCAGACAGGAAAATTCCATCTGACTTGAGGTCAGTTCGTTCCGAGCCCGCTGATAAGAAACTACTGTTGATAAAGAAAACTACCAGCGTGAAGGTCAGAAAAGTTCCAGGAAAGCCAAAACCCCTACCACCATCGAAATCATTGCCATGGACGCCACCAGGGCACAGCATGAAGATGAGATGCGAGAGTCCGAAACCATTATCCAAGAGCAGCATGAGCCCAATCAGACGCAGATCACCTGGACAAGCTTCAAACCCTTAG

> P_richocarpa_2

ATGGAGCAGTTTAGGCAGATTGGAGAGGTACTCGGAAGTCTGAAGGCATTGATGGTGTTTCGCGACAATATACAAATCAATCCACGACAATGCTGTCTGTTGCTTGATGTATTCAGTTTTGCATATGATTCAATAGCTGAAGAGATGAGACAAAACCTGAAATTTGAAGAGAAGAACGAGAAATGGAGAATTCTTGAACAGCCATTGAGAGAGATTTATAGAATATTCAAAGAAGGAGAAGGTTACATCAAGCAGTGCCTGGAAACCAAGGATTGGTGGGCCAAAGCCATTACCCTCTATCAGAACTCGTATTGCGTTGAGTTTTACATCCATAACTTATTATCATGTATCCCTGTTGTCATTGAATCAATCGAAATTGCTGGAGAATTTTCTGGATTAGATCAGGATGAGATACAGAAGAAGAGACTTGTGTACTCGAACAAATATCAGAAGGAATGGAAAGATCCCCGACTTTTCCAGTGGAAGTTTGCAAAGCAGTATCTTATTTCTCAGGAACTTTGCAATCGGTATAATACAGTTTGGAAAGAAGATAGGTGGGTGCTCCTAAACAAAATCCTGGAAAAGAAGATGTCAGGCTCGACAAAGCAAGAGCGGCAACTTACAGATATTCTATTGAAAAACTTGGAAGGATCGGAGCCAGTAAATGGGAAACTCTTACCATGTTCAATCTTAGTACGGTCCAAGGACTACAGTGTAAGGAGACGCCTTGGGAGTGGGAGTCAGTACAAGGAGATACTATGGCTGGGTGAAAGCCTTGCCCTGAGACACTTTTTTGGAGACATTGAGCCCCTATTTCCCGAGATTTCTTCATTGTTATCCCTTTCCCACCCGAATATATTGCAATTCTTTTGTGGGTTCACTGACGAGGAAAAGAAAGAGTGTTTTCTAGTCATGGAACTAATGACTAGGGACCTCTGCAGCTGCATCAGAGAAACCTGTGGCCCAAGAAAGCGCATTCCATTCTCTCTTCCCATTGCGGTTGATCTTATGCTTCAAATTTCCAGAGGAATGGAATATCTCCACTCAAAGGAAATCTACCATGGTAATTTGAACCCTTCTAACATTCTTGTTAAACCAAGAAACATCACCTCAGAAGGGTACCTGCACGCTAAGGTTTCGGGTTTCGGGCTCTCTTCAATCAAGAACTTCACTCCAAAAAACTCATCAAACCAGAACGAAACCCTCTCGTTCATCTGGTATGCTCCAGAAATTTTGGAAGAGAAAGAACAGACAGGAAGTGAAAAGAATTCTAAATACACAGAAAAAGCTGATGTCTACAGTTTTGGAATGGTTTGCTTTCAACTTTTGACTGGGAAAGTTCCATTTGAGGATAGCCATCTTCAAGGGGACAACATGAGCCGAAACATCCTAGCTGGAGAGAGGCCTCTATTTCCATTTTATTCACCAAAATATGTTACTAACTTGACAAAGAGATGTTGGCATACGGACCCAAATCAACGTCCAAGCTTCTCATCCATATGTAGGATCCTTCGCTATGTAAAACGTTTCCTTATTATGAACCCTGATTATAACAGAGAACCAGAGCCGCCAATGCCAGTGATAGACTACGGAGACATGGAGACGAAGCTTTTAAGGAAATTCCCTTCTTGGGATACAGCAGAATCATCAATGGTAGCACAAATTCCATTTCAAATGTTTGTTTACAGAGTTGTTGAAAAAGAGAAAGCAAGAACAACTCAGAAAGAAACTTCAGAATCAGGGAGCGATAAAGCCTCATTTAGAACACCAAAAGGCCGTTCAAGACCTCCCCAACTGCAGTGCACGCGTAGTTTAAGGACAGGTTCAGAAAGCCAGTTGATGATGATCAGTCCAAGACCCAGAAGAACATCTTCTGGTCATGCATCAGATTCAGAGATCTCCTAG

>P_vulgaris_1

ATGGAACAGTTCCGGCATATCGGAGAGGTGTTAGGGAGTCTGAAAGCGCTTATGGTTCTGCGCGACGAAATCCAAATCAACCAACGGCAATGCTGTTTGATCCTGGAAATATTCAGTTTGGCGATGGACACGATCGCGGAGGAGATACGGCAGAACCTGAAGCTGGAAGAGAGGAACAGCAAATGGAAGGTTCTGGAATTTCCTCTGAGTGAGTTGTGCAGGGTTTTCAAGGAGGGAGAGCATTACATAAAGCAGTGTTTGGATTCCAAAGGTTGGTTGGGGAAAGCCGTCACCCTCTCTCACCACCGAGACTGCGTGGAGTTCCACATCCACAACCTCCTCTGCTATTTCCCGGCGGTGATCGAGGCCATCGAGATTGCAGGGGAGGTTTCAGGTTTGCACCCGGAGGAGAAGGCAAAGAAGAAGGTGATACTGGATCGGAAGTACGACGTGGAGTGGAACGACCCGGAGCTGTTCCAATGGAGGTTCGGGAAGCAGTACCTGGTCCCGCGCGGCATTTGCAAGAAGTTGGAGAACGCGTGGAGGGAGGACAGGTGGAGGCTCATCGAGGCCATCAGGGACAAGACTCCGAAGGAGGGTGGTTCTGCTGCGTTCAGGAAGAGCGAACGTGGCCTCGCTAATATGCTCCTCAAGAAGCTTCTTAACGGGTCGGAGATCAGCCCGATTGCGGTGCTTGTCGGGTCCAAAGATTACCAGGTGAGACGCCGGTTGGCCCACCTGAGGGACTTCAAGGAAATTCAATGGCTGGGGCAGAGTTTCGCGTTGAGGAACTTTCAAGGGGAGAGAGAAGCGCACCAGGGTGAGGTCTCTTCTCTGCTCTCGCTTTCGCACCCCAACATAGTGCAATACCTTTGCGGGTTTTACGACGAGGTGAAGCAAGAGTTCTCTCTTGTGATGGAGTTGATGAAGAAGGATTTGTGGACTTACATGAGAGAGAATTGTGGTCCCAGGAGGCAGATTCTGTTCTCTGTCCCTGTTGTGGTGGATCTCATGCTTCAGATGGCCAGAGGCATGGAGTACCTTCATTCCCAAAACATTTCTCATGGACAACTTAACCCATTGAATATTCACCTCAAGTCAAGGAACTCCCAAGAGGGTTATTTTCAGGCAAAGGTTGCAGGCTTTGCCTTGTCTTCTGTCAACAATGGTAGTGCTGACACTGGTGCCATTCATGACTTTAGCCCTTTCACTTGGTATGCTCCTGAAGTTCTAACCCGGTTGGAACAAACGCAGAATGTCCTCTGTTCTTCTACAGAGAAAGCCGATGCCTACAGCTTTGGGATGATATGCTTCGAGTTGTTGACAGGGAAAGTGCCTTTTGAAGAAAACCATTTTCAAGGAGACAGAATGGTCCAAAACATCGAAGTCGGTGAGAGACCTTTGTTTCCTTACCATTCCCCGAAATACCTTGTGAGTTTGATCAAGAAGTGCTGGCAAACTGACCCTGCTCAGCGTCCCAGCTTCTCCTCCATCTGTAGGATTTTGCGGTACACCAAAAAGTTCCTTACCATGAACTCAGAGTCCCAAGTCACCAACCCTGAGCTCAACGTTCTTGAGGCTCCTCCTGTTGATTGCTGCGAGATAGAGACAATGTTCCTTAAGAGCTTTTCCACCGAGAGGTCGTCTTGTGTCTCCGCTGTGTCGCAGATACCCTATGAAATGTTTGCTTATAAGGTTGTGGAGAAAGAGAAAATGAATCCCAAAAACAGCGGTACTAAAGATGAGTGCTGCGAGCACCACCGAAAGGATGGGTATAGGAGTGAAAGTGATGAGCCAAGAACACTCTGTGAAGATGATAACGCATTCATAGTGGAAGATCCTCTCCCTCGAACAACCTACCCCACATCAGTTTGTGACGCACGAACTGTTTCCTTTCAAGTGCCCTACAAGAAAACTGTTAGAATAAAGAAGCAAACACATCTAAAGGAAAAAAAGGACCAAGGAACGCCAAAATTGCGGCCGACAAAATCATTACCATCTACATCGAGTCGAGTTTCAAGGACGAGAAAGGTGAATACACCATCACCGGTTCCAAGCAGTTTGAGCCCCAGTAGAAGACGATTTACCCTGGGGTCTGAATCGGATCGCAGCTCTAAAGGGAATCGATCACCATCACCGGCTCAAACGAGGTTGAGCCCCAGTAAAAGACGGCTTACCCTGGGGTCTGAATCGGATCGCAGCTCCAAAGGGAACCGATCACCATCACCATTGCCCATTCCTTCAAGGAAAAAACACGAGGGTAATTTGTCGGATCATAAGGGCAGTTTAAGGTCGAAGAAAGTGGATCAGTCAGCATTAACTAGATATGGTAGGATGACAGATAGTTCAAGGATGAAGTTGAACAAATCTTTGAGCCCGGGAAGAATGAGACGTGCCAACACTGAGACCAATTTGAAGGGGCTCACGCCACCCCGTCCATTGAGCAATGCAGGAACATCCACAAGGAGCAGAAAAAGTGATAATGTCTCAGTTTCGGACAACGGTTCAAGGATGTCAACAGGTAGGTTTTCGCCGTTGGCAGTCACTCCCCCAAGCCCCTTTGAGAAGAAAACAGACAAGCGGTCATCTTTCTGA

>P_vulgaris_2

ATGGAACAATTTAGGCAAATTGGGGAGGCACTTGGGGGTTTGAAATCTGTGATGGTATTCCGGGAGAACATTCAAATCAATCAGAGGCAATGTTCTCTGCTGCATGATGTGTTCAGCTTTACATATGAGTGTATTGCAGATGAGATTATACAGAACCTGCAATTTGAAGAGAAGAATGTGAAGTGGAAGATTCTAGAGCAGCCCTTGAGAGAGATCCATAAGATTTTCAGAGAAGGTGAAACTTACATTAGGCAATGCATGGAAACAAAGGACTGGTGGGCAAAAGCCATTACCTTATGCCACAACACAGATTGTGTTGAGTTTCATATACACAATTTGCTTTGTTGCATGCCAGTTGTAATTGAAGCTATAGAGTCTGCAGGAGAAACATCAGGGTTGGATCAAGAGGAGATGCAAAGGAAGAAGAATATAAATTCTAACAAATATAGGAAAGAGTATAGAGACATGAAACTTTTCAGGTGGAAATTTGGGAAGCAGTACCTCATCACTCAGGACTTATGCCACCGTTATGATACAGTTTGGAAGGAAGATAGATGGTTCCTTTACAACAAACTCTATGAAAAGAAATTAGAAGGTGTAACAAAGTATGAGAAGAAATTGATAGATTTGCTTTTGAGGAATTTGGAAATATCAGAATCAGTAGCAGGGAGGCTCCTTCCAAGCACAATATTGGTTGGGTCTAAGGACTTCCAGGTGAGAAGAAGAATGGGGAATGCTAGTCAGTACAAGGAGATTTCATGGTTAGGTGAAAGCTTTGTGATAAGACATTTTACTGGAGACATTGAAGCTTTGCAACCTGAGATCATAGAACTTTTATCTCTTTCCCATCCAAATATAATGGACTGCCTTTGTGGTTTTGCTGATGAGGAGAAGAAAGAATGTTTTTTGCTAATGGAACTCATGAGCAAAACCCTTTCCACCCACATCAAAGAGATTCATGGTCCACGGAGGCGAATACCGTTCTTGCTTCATGTGGCAGTTGATCTCATGCTTCAGATTGGCAGGGGAATGGAGTATTTGCATTCAAAGAAATTATATCATGGAGAACTAAACCCTTCAAGCATTCTTGTTAAGCCTAGAGGTACATCCCCAGAGGGTTACTTGCATGCCAAGGTAACTGGATTTGGACTAACTTCTGTCAAGGATTTGAATCAAAAAGGGACCACAAATCAGAATGGAACTCCCCCATTCATCTGGTACTCTCCTGAAGTACTTGAGCAGGAAAACTCAGGGGGTACATCAAGCTCCAAGTACACAGAGAAATCTGATGTGTATAGTTTTGGAATGGTTTGCTTTGAGCTTCTAACTGGCAAAGTCCCTTTTGAAGATAGTCATCTCCAAGGGGAGAAAATGAGCAGAAACATAAGGGCAGGAGAGAGGCCACTTTTCCCACTTAATTCACCAAAATATGTTATTAACTTGACAAAGAGATGTTGGCATATTGACCCAAATCAACGTCCAAGCTTCGCTACCATATGTAGAGTTCTTCGTTACATAAAAAGATTTCTTGCTATGAACCCTGGTTACAGTAGTCAGCCAGAACCACCACTGCCACCAGTAGATTATTGTGATGTAGAGTCTGTGCTTCTGAGGAGGTTTCCTTCTTGGGGAAGTTCCGAATCAGCACCAGTTTCAAAAATTCCATTTCAAATGTTTGCCTACCAAGTTATAGAGCGAGAAAAAATAACTACAGGTTGTAAGGAAAACTCTGAGTCTGGGAGTGATGCTTCAGCCTGTGGTGATGAACTTGTTACTTCAGGGGATGAACCACTTCCATCCACAGCTGAAAGGAAGACTTTTCTCGGAAATGAAATTATGAGCAGAAAAATTAATTTGACCAGGAAATCTCTTGATTTGAAGCTCGCCAAACAACCAGGTACACCGAAAGCGAGATCAGCCAAACCTCCACAAATGTCCCCTCGTGCAAGAAGTATGAGGATGTCTTCAGAGAATATATCAAGTCCAAGGGGAATAAGAAGAGTAGCTTCTGGTCATGTCTCAGACTCTGAGCTTTCTTAG

>R_communis_1

ATGGAAAGGTTTAGACAGATCGGAGAGGTACTGGGAAGTTTGAAGGCATTGATGGTGTTTCATGACAATATCCAAATCAATAAGAGACAATGCTGTTTGTTGGTTGATATATTCATTTTTGCTTACGACACAATTGCTGAAGAGATGAAGCAGAACCTGAGGTTCGAAGAAAAGCACACGAAATGGAGAATACTGGAACAGCCATTAAGAGAGATTCTTAGAACATTCAAAGATGGGGAGGGTTACATCAAGCAATGTTTGGAAACAAAGGACTGGTGGGCAAAAGCTGTTACACTTTATCAGAATACAGATTGTGTCCAGTTGTACATACATAACTTACTTACCTGCATTCCGATTGTAATTGAAGCAATCGAAACTACTGGAGAATTTTCAGGTTGGGATCAGGATGAGATACAGAAGAAGAGGCTTGTGTACTCGAACAAGTATCAAAAGCAATGGAAAGACCCTCAGCTATTTCACTGGAAGTTTGCAAAGCAGCATCTTATTACTCAGGATTTCTGCGAAAGATTCGATGTGGTTTGGAAAGAGGACAGGTGGATTCTCCTTAATAAAATCAAAGAAAAGACGGTTTTATGCTCCAGAAAATATGAGAGACAACTAAGAGACCTTCTGGTAAAGAATTTGGAAGAACAAGATACTGAGAATGCGAAACTCTTACCAAGCTCAATTCTAGTGGGGTCGAAAGATTATCATGTTAGAAGACGACTTGGGAATGGGAGTCAATACAAGGAGATACTCTGGTTAGGTGAAAGTCTTGCCATTAGACATTTCTTTGGTGATATTCAACCACTTGTTCCTGAAATTTCTTCCTTGTTATCTCTTTCACATCCAAATATATCATACTTCTTTTGTGGGTTCACTGATGAGGAAAAGAAAGAATGTTTTTTAGTTATGGAACTAATGAGTAGAGACATGTGCAGCTATATCAAAGAAATATGTGGACCAAGAAAGAGAAGTCTTCCATTTTCTCTTCCTGTTGCAGTTGATATTATGCTACAAATTGCCAGGGGAATGGAATATCTCCACTCAAAGAAAATATATCATGGTGATTTAAACCCTTCTAACATTCTTGTTAAACCAAGAAGCACCACAGAAGGGTACGTGCATGTTAAGGTTTCTGGGTTCGGGCTCTCGTCCTTTAAGAAAAACCCTTCAAAACAGAATGGAACTCTCTCATTCATTTGGTATGCACCAGAAGTTTTAGAAGAGCAAGAACAGACAGGAAGTGCCCCAAATTCAAAATACACAGAAAAATCTGACGTGTATAGCTTTGGAATGGTTTGCTTTGGAATTTTGACAGGGAAAGTTCCATTCGATGATAGCCATCTTCAAGGAGAGAAAATGAGCCGAAACATTCGAGCGGGAGAGAGGCCGCTGTTCCCATTAAATTCACCAAAATATGTCACTAACTTGACAAAGAGATGTTGGCAAGCTGACCCAAATCAACGCCCAAGTTTCTCATCAATCTGTAGAATTCTTCGCTATACAAAAAGGTTTCTCGCCATGAACCCTGATTACAATAGAGAGCTAGACCCACTGATGCCAGCAGTAGACTATGTTGACATGGACTCAAAGCTTCTAAGGAAATACCCGTCTTGGGATGCAGCTGATTCCTCACAAATAGCACAGATTCCATTTCAGATGTTTGTTTGCAGAGTTATTGAGAAAGAAAAAACCAGAGTAAACCAGAAAGAGACTTCAGAATCAGGAAGCGATAAAGCTTCTGTTAGTGGAGATGAAAACATGAACATAACCGATGATCCATTCCCGTCGCCCACATTACCATCACCCACAGGAAGGAAGTTCTTAACATCACCTGATACTCTGAACAGGAAACCTTCCATGTCAAAGAGATCGCCAGATGTGATAAGATCAAGTAGACAAGCAGGGACACCAAAAGGACGGTCAAGACCCCCACAACTGGCACCTTTCGGGCGTGGTTTAAGAATGAATTCAGAAAGCCAGCTAATGATGATGATTAGTCCAAGAATGAGGAGAACATCTTCGGGTCATGCATCAGATTCCGAGATTTCGTAA

>R_communis_2

ATGGAGCAGTTCCGTCAGATAGGAGAGGCATTGGGAAGTTTGAAGGCTCTAATGGTGTTGCAGGATGATATTCAGATCAATAGGAGACAGTGCTGTTTGTTGCTTAATATCTTCACTTCAGCTTTCGACACAATTGCGGAGGAGATCAAGCAGAATTTGAAGCTTGATGAAAAGAATACCAAGTGGAAACCCCTTGAAGAGCCTCTAAAAGAGCTTTACAGGGTCTTCAAGGAAGGGGAGCTCTATGTTAGGCGATGTTTGGATAGTAAAGATTGGTGGGGTAAAGCTGTTGTCCTCCATCAAAACAAGGACTCAATTGAGTTCCATATTCATAATTTGGTTAGCTACTTCCCCGCTGTTATCGAGGCAATCGAGACTGCAGGAGAGATCTCAGGACTTGACCAGGATGAGATGCAAAAGAAAAGGGTTATGCTTTTCAAGAAGTATGACAGAAGTTGGCATGATCAGAAACTTTTTCAATGGAGATTTGGGAAGCAATATTTGGTCTCCAGGGAAATTTGTAATCAAATTGAGACTGCAGTGAAAGAAGACGGATGGCTTCTCGTTGGAGCAATCAAAGAAAAGATGAAAGCAGGGTCACTAACAAAGAACGAGCAGCGACTTGGGGATTTATTACAGAAGAAGCTGAATGGACAGCTGCTAATTAATGGCAAACTCCCTCCAAGTTCAATTCTGTTAGGAGCAGAAGATTACCAAGTCAGGCGAAGATTAGGTGGAGGAAGCCAGTACAAAGAAATTCAATGGCTAGGAGAAAGCTTCGCTTTGAGACATTTCTTTGAGGATATTGAGCCATTAAACTCTGAGATATCTGTGCTGCTATCACTTTCCCATCCTAATATAGTGCAGTATCTCTGTGGATTTTACGATGAAGAAAAAAAGGAATGTTTTCTTGTAATGGAGTTGATGAGCAAAGATCTTTATGCATACATGAAGGAGAATAGCAGTTCAAGGAGGCAGGTTTTGTTCCCTCTTCCTATCGTTGTTGATATCATGCTTCAGGTAGCAAGAGGTATGGAATTTCTCCACTCACGAAAGATCTATGTTGGAGATCTGAATCCTACTAACATCTTTCTCAAACCAAGAAAATCCACGGAAGGTTATTTTCATGTGAAAGTCTCAGGGTTTGGTTTAACTTATATCGAAAATCCATCTTCTCGACACTCATCATCAAACCAAAATGCATTTGATCCTTGCATTTGGCATGCACCGGAAGTTTTAGCTGAAAGAGAACAGACCGGAAGCCCCTCTACTCGAAAACACTCAGAGAAAGCAGATGTTTACAGTTTTGGAATGCTTTGTTTCCAGCTTTTGACAGGTAAACTTCCTTTTGAGGATGGGCATCTTCAAGGAGACCAGATGGCGAAAAACATAAGAGCAGGGGAGAGACCATTATTCCCATCCCTTTCACCAAAATACCTTGTGAACTTAACCAAAAAATGTTGGCATACTGATCCAAACTATAGGCCTAGTTTCTCATCTATTTCTAGGGTTCTAAGGTACATCAAGAAGTATCTTGTGATGAATCCTGTCGATGGTCAGCTTGTTATGCAGTCACCACCAGTCGATTACTCAGAACTGGAAGCAGGGTTCTTGAAGAGGTATCCAGGGGAGATGAATGGTGGTCTAGCCTCAGTATCACAAATCCCATTTCAAATGTTTGTTTACAGGATTTCCGAGAAAGAAAGGACTTCTCTGAGCTTCAGATTTAAGCAGTCAGAAACAAGTGAAGCCGCCTCAAATGGCTGGGATGAGAATATTTCTGTAGTAGAGGATCCAGTGGTACCAGCAAGTGACGCAAGGTCAATTAGCTCCGACATGAAGACAGTTTGTTTTGATTTAAGTTCTATCTACATTGAGAATCCAGATGGGAAAATCCCATCTGATTTGAGGTCAGTTCGTTCAGAGCCTCCAGACAAGAAAATAATGTTGAGAAAAAAAAGTTCCAATGTCAAAGTCAGAAAAACGCCAGTTAAACCGAAGGCACCAATACCAGCAAGAACATCGCCATGGAATCCACCAGGACAGAGTCCAAAGGTCAGCCGAGAAAAGCCGTTCTCTTCAAGTCCTCTCAGTCCTGCAAGGCGCAAAGCAAGTGGGCAAGCCCTAAGTCCTGAGAAGTTATAG

>S_bicolor_1

ATGGAGCAGCTCCGGCAGCTCGGCGAGGCGTTGGGTGCCATCAACGCGCTGATGGTGTTCGAGCCCGAGCTCCGCGTGAACCCGCGGCAGTGCCGCCTGCTGGCCGACGCGTGCGCGCACGCGCTCGCCGCCGTCACGGGGGAGGTGCGCGCGCACCTCCGCTTCGAGGAGCGCGGCGCCAAGTGGCGCGCCGTCGAGCCGCCGCTCCGCGAGCTCCACCGCGCGTTCCGCGACGCCGAGGGCTACGTCCGCCACTGCCTGTGCCTGGACCCGCGCGGCGGCGGCGGGGGCAGCGGCTGGTGGGCGCGCGCCGCGGCCGCGGCGCACGGCACCGAGTGCGTCGAGCAGCACCTCCACGGTATCCTCTGGTGCGTCGCCGTCGCGGTCGAGGCCGTCGAGGCCGCCGCGGAGATCGCCGGCGACGACGCCGATGAGATCGCGCGGAGGCGGGTTGTGCTCGCCAACAAGTACGACGGTCGGAGCATGCTCGAGCCCAGGATGTTCCAGCACGCCTACGGCAAGCTGTACCTGGTGTCCCAGGAGTTCGTCGCGCGGATGGACACGGCGTGGAAGGAGGACAGGTGGCTGCTGTCGCAGCTGCTCGACGAGATGAAGTCGCCGGCGGCGCCGAAGCCGCTGACCAAGAGCGAACGCCGGCTCGCCGACGTCCTGGCCGCGCCGCGGGGAAAGCTGCACCCGGCGTCTGTCCTGCTCAACGGGGACTACAGCGTGCGGAGGCGCCTCGGTGGCAACCTGAAGGAGGCGCATTGGATGGGGGACAGCTTCGCCGTGAAGCATTTCATCGGGGACGCCGACGCCGAGGTCTCGATGCTCTCGTCGGTGGCGCACCCGAACGTGGCGCACGCCGCCTACTGCTTCCACGACGAGGAGAGGAAGGAGTACTTCGTGATCATGGACCAGCTCATGGCCAAGGACCTCGGCAGTTACGTCAAGGAGATGAGCTCCCCGCGGCGGCGGATCCCGTTCCCCCTCGTCGTCGCTGTCCACATCATGCTGCAGATCGCGCGCGGGATGGAGTACCTGCACGCCAACAAGATCTACCACGGCGAGCTGAACCCGTCCAACGTGCTCGTCAAGCCGCGGCAGCAGCCTGACGGGTACGTGCACGTCAAGGTCGCCGGGTTTGAGCGGCCGGCGGGCACCGTCACAAACGGCGCAAAGGCGTCTGCTAATGGCAATGCCAACGCAGCCGGCGCCGGCGGCGACGACACCTGCATCTGGTACGCGCCGGAGGTTCTCGAGCATCAGGGGAGCCGCGACAGGCACACCGAGAAGGCGGACGTGTACAGCTTCGCGATGATCTGCTTCGAGCTGCTGACGGGCAAGGTCCCGTTCGAGGACAACCACCTGCAGGGCGACAAGACGAGCAAGAACATCCGCGCCGGCGAGCGGCCGCTGTTCCCGTTCCAGGCGCCCAAGTACCTGGTCGCCCTGACCAAGCGGTGCTGGCACGCCGACCCGGCGCAACGCCCGCCGTTCGCCTCCGTCTGCCGCGTCCTCCGGTACGTGAAGCGGTTCCTGGTCATGAACCCGGAGCCGCAGCAGGCCGACGCGCCGCCAGCCGCGCCGGCCGCCGACTACCTCGACATCGAGGCGCAGCTGCTGAGGCGGATCCCGGCGTGGCAGCGGGGCCAGGGCGCGCCCCCGCGCGTGGCGGACGTGCCGTTCCAGATGTTCGCGTACAGGGCCGTGGAGAGGGAGAAGACCGCCGCCGGCGCGCACGCCAGCAGGGACAGCCGGGCCACGGATTCCGGCAGCGAGGAGAACTCGCTGTGCGGCGACGAGAACGGGGTCGGGGCGACCACCCCAGACGACGACGCGTCCACCGTGTCGGGCGGCACAGTGAGGTCGCGACCGGACAGCAGCGACGGCAAGAAGACTACTCCGGTCAGGAAGGCGGACGGCAAGACGCCGCCCAGACAAGCAGGATCTCAGCAGAAGGTGAAACCGGTGAGCGCGGTGAAGCCTCCGCCGGCGACGAGGAAAACGGTGGGCGTGAAGCCGGAGCTTCCGGCGCGCCGGCCGACGTCCGGGCACACCTCTGACTAG

>S_bicolor_2

ATGGACCAGCTCCGCCAGGTGGGCGAGGCCCTCGGCGGCGTGCAAGCGCTCATGGCCTTCGCCGACGATCTCCGCATCAACCCGCGGCAATGCAGCCTCCTCGCCGACGCCTGCGCGCTGGCTTTCGCCGCCGTAGCCGCCGAGGTGCGCGCCCACCTCCGCTTCAGCGAGCGCCTGGCCAAGTGGAAGCCCCTGGAGGCGCCGCTCCGGGAGCTCCACCGCGCCGTCCGCGACGCCGAGGGCTACGTCCGCCACTGCCTGGAGCCGCGGGACAGCTGGTGGGCGCGCGCCGCCGCGGCCACGCACGGCGCCGACTGCGTGGAGCAGCACATGCACAGCCTGCTGTGGAGCGTCGCCCTGGTGCTGGAGGCCGTCGAGCTCGTCTCGGAGGTGACGGGGTCCGACCCGGACGAGCTCGCCCGGCGGCGGCTGCTGTTCGCCAAGGACTACGACAGGGACATGCTGGAGCCGAGGCTGTTCCGGCAGAGGCTCGGCGCGCGGTACCTGGCCACGCGTGAGCTCGCCGCCAGGATGGACGCGGCGTGGAAGGAGGACCGGTGGCTCCTGTTGCAGTACCTCGAGGAACGGAGGAGCCCGGGCTCGCCGAAGCCGCTGACGCGGAACGAGCACCGGCTGGCCGACCTCCTGACGGCGCCGCGCGGGAAGGTGCACCCGGCGTCCGTGCTCCTGCAGGGCGACTTCCACGTGAGGAGGCGCCTGGTGGGCAACCTCAAGGAGGTGCAGTGGATGGGGGAGTCCTTCGCCGTGAAGCACTTCGTCGGCGCCGACGCCGACGCCGTGGGCGCCGAGGCCGCGCTGCTCACCTCCGTGGCGCACCCGAACGTAGCGCACTGCCGGTACTGCTTCCACGACGAGGACAAGCGGGAGTTCTTCCTCGTCATGGACCAGCTCATGACCAAGGACCTGGCCAGCCACGTGAAGGAGGTGAACAACGCCAAGCGCCGGGTGCCGTTCCCGCTCGCCGTCGTCGTCGACGTCATGCTGCAGATCGCGCGGGGAATGGAGTACCTGCACTCGAGGAAGATCTACCACGGCGACCTGAACCCGTCCAACGTGCTCGTCAGGACGCGGCACGCGGACGCGCACCTCCACGTCAAGGTCGCCGGGTTCGGGCAGTCCGCGGCGACCGTGGCCGCCGCCAACCCCAGGCCCAGCCCTAGGGCGTCGGCGAAAGCAGCAAACGCCACCGTCGCCGCCTCCAACCCTTGCATCTGGTACGCCCCGGAGGTGTTCGAGCAGGAGGCGGCCAAGTGCACGGAGAAGGCGGACGTGTACAGCTTCGGGATGGTCTGCTTCGAGCTGCTGACGGGGAAGATCCCGTTCGAGGACAACCACCTGCAGGGGGAGCACATGAGCAAGAACATCCGCACCGGCGAGCGGCCGCTGTTCCCGTTCCAGGCGCCCAAGTACCTGACCAGCCTCACCAAGCGGTGCTGGCACGGGGACCCGGCGCAGCGGCCGGCGTTCGCGTCCATCTGCCGCGTCCTCCGCTACGTGAAGCGGTTCGTGGTCATGAACCCGGCCCCCGCGGAGCAGCCGGACGCGACGCCGCCGCCGCCGCCCGTGCCGCCCGTGGACTACCTCGACGTCGAGGCGAGCCTGCAGAGGAGGTTCCCGGCGTGGCAGTCCGGAAACGCGGCGCCGCGCGTCTCCGACGTGCCGTTCCAGATGTTCGCGTACAGGGTGATGGAGAAGGAGAGGAACAGGGCGGCCATCCTGCACATCGGCGGCGGCGGCAGGGACAAGGCCTCCGACTCCAGCAGCGAAGGCAACTCGCTGTGCGGGGACGAGAGCGGCAGCGCGACGCTGTCGGACGCCGATGCTCTGTCCGTGTCGAGCCGCGGCACCACCATCACCACCACCACTACGCGGTCGCTGCCGGACCGCGCTGGCAGCAGGAAGGCGTCGCCGAGCGACCGCGCCAGCAGCAGGAAGGCGTCGCCGAGGAAGGTGGATAGCAGCAGGGTCGCAACCCGGCTAGCAGGGCGACCGGAAAAATCCAAGTCGATGGAGAAGTCCAAGTCGATGGGCGTGGTGAGGCCGCCGCAGATCATCAGGCGGACGCAGAGGATCAAGTCGGACGGCCACCTGAATTCGGACGTGATACCGTCTCCCCGGCGGCGCGGTTCCGGCGGGGGGCACGCTTCAGACTCTGAAATCTGA

>S_italica_1

ATGGAGCAGCTCCGCCAGGTGGGCGAGGCCCTCGGCGGCGTCACCGCGCTTATGTCCTTCGCCGACGACCTCCGCATCAACCCGCGCCAGTGCCGCCTCCTCGCCGACGCTTGCGCGCTCGCCTTCGCGTCCGTCGCCGCCGAGGTGCGCGCCCACCTCCGCTTCCGCGAGCGCGCGGCGAAGTGGAGGCCCCTGGAGGGGCCCCTACGTGAGCTCCACCGTGCCGTCCGCGACGCCGAGGGCTACGTCCGCCACAGCCTGGAGCCGCGCGACAGCTGGTGGGCGCGCGCCGCCGGGGCCACGCACGGCGCCGACTGCGTCGAGCAGCACCTGCACAGCCTGCTCTGGAGCGTCGCCGTCGTGATCGAGGCCGTCGAGGCCGTCTCGGAGGTGACCGGGTCCGACCCCGACGAGCTCGCCCGGAGGCGGCTGCTGTTCGCCAAGGACTACGACAGGGACATGCTCGACCCCAGGCTGTTCCGGCAGAGGCTCGGCGGGCGGTATCTGGCCACCCGCGAGCTCGCCGCCAGGATGGACACGGCGTGGAAGGAGGACAGGTGGCTCCTGTCCCAGCTCCTCGAGGAACGGAAGGACCCGGCGTCGACGGAAACGCTGACGCGGAACGAGCACCGGCTCGCCGACCTCCTGACGGCGCCGCGGGGGAAGGTGCACCCGGCGTCCCTGCTCCTGCATGGCGACTTCCACGTGCGAAGGCGCCTTGCGGGCAACCTCAAGGAGGTGCAGTGGATGGGGGAGGCCTTCGCGGTGAAGCACTTCGTCGGCGCCGACGCCGACGCCGTGGGCGCCGAGGTCCCGCTGCTCACACTGGTGTCGCACCCCAACGTGGCGCACTGCCGGTACTGCTTCCACGACGAGGACAAGAGGGAATTCTTCCTCCTCATGGACGAGCTCATGACCAAGGACCTGGCCTCCCACGTCAAGGAGGTGAACAGCGCCAAACGACGGGTGCCGCTCCCGCTCGTCGTCGTCGTCGACGCCATGCTGCAGATCGCGCGCGGCATGGAGTACCTGCACTCCAAGAAGATCTACCACGGGGACCTCAACCCGACCAACGTGCTCGTCAAGGCGCGGCACGCCGACGCGCACCTGCACGTCAAGGTCACCGGGTTCGGGCAGTCCGTCGTTGCCGCGGCCAGCCCAAGGCCGAGCCCGAGGGCATCGGCGAACGCGAACGCCAATAACGCCAGCGCCGCCGCCAACCCCTGCATCTGGTACGCGCCGGAGGTGCTGGAGCAGGAGGCGGCCAGGTGCAGCGAGAAGGCGGACGTGTACAGCTTCGCCATGGTCTGCTTCGAGCTCCTGACCGGCAAGATCCCGTTCGAGGACAACCACCTGCAGGGCGAGCACATGAGCAAGAACATCCGCGCCGGCGAGCGGCCGCTGTTCCCGTTCCAGGCGCCCAAGTACCTGACCAGCCTCACCAAGCGGTGCTGGCACGGCGACCCGGCGCAGCGCCCGGCGTTCGCGTCCATCTGCCGCGTCCTCCGCTACGTGAAGCGGTTCCTGGTCCTGAACCCCGCACCGGCGGACCAGCCCGACGCGCCGCCGCCCTTGCCACCGGTGGACTACCTCGAGGTGGAGGCGAGCCTGCTGAGGAGGTTCCCGGCGTGGCAGGCCGGGAGCGCGGCGCCGCGCGTCTCCGACGTGCCGTTCCAGATGTTCGCGTACAGGGTCGTGGAGAAGGAGAGGACCAGGGCGGCCATCCTGCACATCGCCAGGGACAAGGCTTCCGACTCCAGCAGCGACTGCAACTCGCTGTGCGGGGACGAGAGCGGCGGCAGCCTCGGCGCGGTGCTGTCGGACCCGGAGGCGCTGTCCGTGTCCAGCCGCGGCACGGTGCGGTCGCTGTCGGACCGGAGCGGCAGCAGGAAGGCGTCGCCGAGGAAGCTAGATCGCAGGATCACCGCCAGGCTAGCAGGCAAGCTTTCTGTCCATGTTGCTCCTTTCGCTGATGAGAGCTTCACCTCCACGTACGATGATGATGATATATGTGAGAATGATGGTGCAGGGCCGCCGCAGAAGTCCAAGTCGATGGGTGTGGTTAGGCAGCCGCCGCAGGTCATCAGGCGGACGCAGAGGATCAAGTCCGACGGCCATCTCAATTCTGCCGTGGTTCCGGCCAGACGGCGGCTCGTGTCCGTGGCCGCTTCAGGCGGGGGGCACCTTTCAGACTCCGAACTAGCCTAA

>S_italica_2

ATGGAGCAGCTCCGGCAGCTCGGCGAGGCGGTGGGCAGCATAAACGCGCTCATGGCGTTCGAGCCGGAGCTCCGCGTCAACCCGCGCCAGTGCCGCCTCCTCGCCGACGCGTGCGCGCACGCTCTCGCCGCCGTCACGGGCGAGGTGCGCGCGAGCCTCCGCTTCGAGGAGCGCGGCACCAAGTGGCGCGCCATCGAGGCCCCGCTCCGCGAGCTCCACCGCGCGTTCCGCGACGCCGAAGGCTACGTCCGCCAGTGCCTGGACCTGCGCGGCGACGGCAGCTGGTGGGCGCGCGCCGCGGCGGTGGCGCACGGCACCGAGTGCGTGGAGCAGCACCTCCACGCCATCCTCTGGTGCGTCGCCGTCGCGGTTGAGGCCGTCGAGGCCGCTGCGGAGATCGCCGGCTCCGATGCCGACGAGATCGCGCGGAGGCGGATGGTGCTCGCCAAGAAGTACGACAGGGACATGGTCGAGCCCAGGCTGTTCCAGAACGCGCACGGCAAGGTATACCTGGTACCCCAGGAGCTCGTGGCGCGGATGGACATGGCGTGGAAGGAGGACAGGTGGTTGCTGTCACAGCTCCTCGAAGAGATGAAGTCCCCGACAGCGCCGAAGCCCCTGACGAAGAGCGAGCAGCGGCTCGCCGACGTCCTGGCCGCGCCGCGGGGGAAGCTGCACCCGGCGTCCATCCTGCTCAGCGGCGACTACAGCGTGCGGAGGCGCCTCGGCGGTCGGCTCAAGGAGGCGCAGTGGATGGGGGAGAGCTTCGCGGTGAAGCATTTCATCGGGGACGCCGAGGCCGAGGCCGCGCTGCTCTCGTCGGTGGCGCACCCCAACGTGGCGCACGCCGCGTACTGCTTCCGCGACGAGGACAGGAAGGAGTACTTCGTGGTCATGGACCAGCTCATGGCCAAGGACCTCGGGAGCTACGTCAAGGAGGTGAGCTGCCCGCGGCGGCGGGTCCCGTTCCCGCTCGTCGTCGCCGTCGACATCATGCTGCAGATTGCGCGCGGGATGGAGCACCTGCACGCCAAGAAGATCTACCACGGCGAGCTGAACCCGTCCAACGTGCTCGTCAAGCCGCGGCAGCCCGACGGGTACGTGCACGTCAAGGTCGCCGGGTTTGAGTGGTCAGGCACCGTCACAACCGGCGGCAAGGCATCTGCCAACGGCAGCGCCAATGCCAACGCAACCGGCGGCGGCGACTACACCTGCATCTGGTACGCGCCGGAGGTGCTCGAGAAGGAGAGCGGCGATCCCTCCGCCAGGCGCACCGAGAAGGCGGACGTGTACAGCTTCGGGATGATCTGCTTCGAGCTGCTGACGGGCAAGGTCCCGTTCGAGGACAACCACCTGCAGGGCGACAAGACGAGCAAGAACATCCGCGCCGGCGAGCGGCCGCTATTCCCGTTCCAGGCGCCCAAGTACCTGGTCTCCCTGACGAAGCGGTGCTGGCACGCCGACCCGGAGCAGCGCCCACCGTTCGCCTCCGTCTGCCGCGTGCTCCGGTACGTGAAGCGGTTCCTGGTCATGAACCCGGACCAGCAGCAGGGCCAGACCGACGCGCCGCCGGCCGCGCCGCCCGCCGACTACCTCGACATCGAGGCGCAGCTGCTGAGGAGGATCCCGGCGTGGCAGCGCGGCGAGGGCGCCGCCGCCGCCGCGCGCGTCGCGGACGTGCCGTTCCAGATGTTCGCGTACAGGGCGGTGGAGAGGGAGAAGGCCGCCGGCGCGCACGCCGGGGGCAGGGACAGGGCCTCGGACTCCGGCAGCGAGGGGAACTCTCTGTGCGGCGAGGAGAACGGGACCGGGGCGGCCACGCCGGACGACGCGTCCACGGTGTCCGGCGGCACCGTGCGGTCGCGCCCGGAGAGCAGCGACGGCAAGAAGACGCCGGTCAGGAAGGCGGACGGCAAGGTGCCACCCAGACAAACAGGGTCTCGGCAGAAGGTGAAACCGGCGAGCGCGGTGAAGCCACCGACGACGGCAAGGAAAACGCTAGGCTTGAAGCCCGAGGTTCCAGCGCAGCGGCCGACGTCCGGGCACGCCTCGGACTAG

>S_lycopersicum_1

ATGGTTATAACCAGATCCATCTCTGGTTTATGTCCAAGCATGCCGGCGTTAGAGGAATTCAGGCAGATAGGAGAGGTAATTGGGAGTCTTAAAGCTCTGATGGTATTCCAAGAGGACATTCAAATAAATAAAAAACAATGTTGTTTGTTGGTGGATATGCTCAAATGTGCCTACAAGACACTTGCAGAAACAATGAAACAGAACTTGAGATTTGAAGAGAAGAATATCAAATGGAAAATTCTCGAGAACCCCTTAAGAGAGCTTCTCAGGGTGTTTAAAGAAGCAGAACAGTACATCAAGCAATCCCTGGAAAATAAGGATTTTTGGGCAAAAGCTGTAGTTCTATATAAGAATACAGATTGTATTGAATTTCACATCCACAATTTGCTCTCTTGTGTGCCAATTGTCATAGAGGCCATAGAAATAGCAGGAGAGATTTCAGGTAGTGACCATGACGAGATACAGAAGAAGAGATTCATTTACTCCATGAAGTATCAGAAAGAGTGTAAGGACCCACGAATCTTCCAATGGAAATTCGGAGAACAGTACATGGTTTCTCAGAAGTTCTGCGAAAGGGTATACTCAGTTTGGAATGAAGACAAGTGGATTTTGAAAAATAAAATTCGAGAGAAGAAGAATTCAGGTGCATGTACTTTGACAAAGCATGAGAAACGACTTGCTGATCTCCTCTTGAAGAACTTAAACGAGATGGAGATGGAGATGGAGAGTGATCCTAAGCTTTCACCTAGCTCAGTTCTTGTAAATTCTAAGGATTACCAAGTAAGAAGACGATTAGGCAGTGGAAGTCAGTACAAGGAGATCCAATGGTTGGGGGAGACCTTTTGTTTAAGGCATTTGTTTGGGGATATTAAGCCCCTGATTCCAGATATTTCGCGAGAACTGCATCTCTCTCATCCAAATATAATGCACATCTCCTGTGGTTTTACTGATGAAGAAAAGAGAGAATGTTTCTTAATCATGGAACTTATGAGCAAAGATTTATCTAGTTACATCAAAGAAATCTGTGGACCAAGAAAGCGTGTACCCTTTTCTCTTCCCGTTGCAATCGATTTACTCCTTCAGATTGCAAGAGGCATGGAATACCTACACTCAAAGAAAATCTATCATGGTGAATTGAATCCTTCCAACGTCCTTATCAAGGCGAGGAACGTATCTGCAGAAGGGTATTTACATGCAAAGGTTTGTGGGTTTGGCTCATCGTGTTCTATCAACCTTCCTCAGAAAGCTAATGTGAATCAAAATAATGGGACACTCCCATTCATTTGGTTTTCCCCAGAAGTCCTAGCTGAACAAGAGCAGTCAGGGAATGGAGGGAACATTAAGTATACCGAGAAATCTGATGTGTACAGTTTTGGGATGATATGTTTTGAGGTTTTAACAGGGAAAGTTCCATTTGAAGATAGCCATCTACAAGGAGACAAAATGAGTAGAAATATAAGGGCAGGAGAGAGGCCATTATTCCCATTTCACTCACCAAAATATGTGACTAGCTTGACAAAGAGGTGTTGGCATACTGATCCATATCAACGGCCAAGTTTTTCATCCATCTGCAGGGTTCTACGCTATGTGAAGAGATTCTTGGTGATGAATCCTGAACACAGCCAACAAGACTCACCATTGCCACCAGTAGACTACGGCGAGATTGAAGCTGTCATCTTTAGGAGCATCCCCTTAGGAAATTGTGACTCTGATCCTCTGCCTGTCACACAAATACCTTTCCACATGTTTGCATACAGGGTGACCGAGAAAGAAAAGTCAAGCACAATTCACAGAGACGTAAATTCTGAATCAGGAAGTGATGGAACTTCTGCATGTGGGGATGATCCTGTAACGGCAGATGATGCACTTCCATCACCAACTGAAAAGAAGAATATTGCATCTCCCGAGATATTGATCAAAAGACTTTCAATTAGGAAACCTGCCGATATCAAAGTCGGCAAGCAACCAGGAACACCAAGGGGAAGGACAGTAAGACCTCCAAGCCTACGTACTATAAGGCAAAATTCTGAAACTCAGTTAATGATGATGAACAGCCCAAGCCCAAGAACACGGAGATCATCTGGCCATGCATCAGATTCGGAGCTCCCTTAA

>S_lycopersicum_2

ATGGTATTGAAACATGACATTCAAATAAACCAGAAGCAATGTTGTCTTCTGTTTGACATGTATGTACAAGCTTTCGACACGATTTCCGAGGGGATCAAACATAATTTAAGGTTGAATGAGAGGAACACAAAGTGGAGAGCACTTGAACATCCAATGAAAGAGCTTCATAGGATCTTTAAAGAAGGTGAAATGTACATAAAAAGTTGTTTGGATGTGAAAGATTTGTGGGGCAAAGCTATAAGTTTACATCTCAATAAGGACTGTGTTGAGTTTCATGTTCATAACTTGCTCTGTTGTTTCCCTGTTGTGATTGAGGCTATTGAGACAGTTGCAGAAGTCTCAGGATTCGATGAGGAAGATATGCAGAAGAGGAGGACTGCACTCGCTAAAAAGTACGAGGGAGAGTCAACGTGTGATCCAAGATTCTTTCAATGGATGTTTGGGAAACAATACTTGGTCACTAGAGAGATATGTAGTGAATTGGAGAGTTGTTGGAAAGAAGATAGATGGTATCTTGTTGAAATGATCAGTCAAAAGATGAATGTTGCAGAAAACTTGGCGATAAACGAGCATAGGCTAGCTGAGATATTGCTTAAGAAGCTAGAGGGATCAAAACAACTCAAACAGATTAAGCTCTTGCCTAGTTCAATCTTGATTGGAGCAAGTGATTATCATGTGAAGAGACGTTTAGGCTCGCGAGGAGGACATGTTAAGGAGATTCAATGGTTAGGAGAAACGTTTGCGTTGAGGAACTTTTTTGGTGAGCTGATTGAATCAGTAGTTGCTGAGATTTCTTTAGTTTATTCACTTTCACATCCCAACATTCTGCAATATCATTGTGGTTTTTACGACGAAGAAAAGAAAGAAGGATACCTTGTTATGGAGCTAATGAATAAAAGTTTAGCAACTTACATAAAGGAGCATTCGTGTCAGAGGAAGAAAGGACCTTTTTCTATCCAAGTTGCTGTTGATATTATGCTTCAGATTGCTCGAGGGATGGAGTATCTGCACTCGAGAAAGATCTATCACGGAGAGCTGAATCCATCTAACGTGCTCCTTAAACCAAGAAATCCTTCAGCGGAGAGTTATTTTCATGCGAAAGTTAAAGGATTTGGTTTAACATCTGTTAAGAGTAGTTATAAGGCTGCTGATGAAAATTCTGCTGAGTCTTTCATATGGTATGCACCAGAAATTCTCACTGAAAAGGAAAAACCAGAAAGCAAATGCACTTACAAGTATACAGAGAAGGCTGATGTTTATAGTTATGGAATGATATGCTTTCAACTTTTAACGGGGAAAGCTCCGTTTGATGAACATCTCCAAGGGGAGAAAATGGCGCAGAATATTAGAACAGGGGAGAGACCTCTCTTCCCGCATCCTTCACCTAAATATCTCGTTAACTTGACAAGAAAATGCTGGCAAATGAATCCAGACCTTCGCCCAAGTTTCTCTTCTTTGTGTAGGATTTTGCATTATATCAAGAAAGTACTTGTCATAAATCCAGGACATGGCCAACCTGAATGTCCTCCTCCACTCGTAGACTACTGTGAAATCGAGGCTGGCTACTCAAAGAAGTTCGCGGGAGAGGATAGCACTGGTTTGGCCCCGGTATCACAAATTCCATTCCAGATGTTTGCTTATAGACTTGTTGAGAAAGAGAGAATATCTGGAAACTCTAAAGACAAGCACTGGGATTCATCACATTATGGTCATTCGAGTCACAGGACAGCATCAATGCAGAGCGACAATGAGCATATGGATGCAATGGATGATCTCTTTTTCGGTCCAAGTGATCGAAGGTCAGTTTGCTCAGAGATAATAGAGAGTAAAGGTCCAAGATTCTGGGATCAGAGGACAGTCATCTCTGAGACACCACTTCGAAAAGTTTTCTCATTCGAGCAATTGTCAGTTAGTTCTGAGAGTCCAAAAGAGAAAATCCCAGCAGCAGCAACAAATGAAAAGCCAATCTATGATGATATTCTGGAAAGGAAAGAAATACTAATACCATCAGGTGATCAGTCTTCACCGCATGTTGATACAACACCACGGAAGGCCATTTCATCAATGAGAAAGAACAAAAAGCTGAATGTTATGGAAATCAAGGAGAATGTAATGTCTTCGAAAACAGATGACAGAAAACCTTCATCGAGTGAGCAGAAAATAGTGACTCCTGTGATTTCAGCGAAGAAAAATTCATCAGAGGAGAAAAACTCATCTGACAACCAACAACCAAAAGATTCTCAAAGTCAAGAAAAGAAGCAGTCATCAACAGCTGCCAATGTCAAGAATATTCGTGGTGATCCTACGCAAAGGAAAGCTGTGTCAAGAAGAACAGAATCCAAGAATCCAGAGAAGAAAGTTCTATCAAGTCCGGAAGCCAATCAAAACTCAATTTCATCTGAGATTACAGCATCGAAACCAAGCAATTCAAAAGTAACTGCAAAGAAAGCACCATTTCACAAAAAACTCCAGGACAAAGAGAGTAGCACTACACCAGACCCATCACCAAGATCTTCACCAGCTAGAGCTAAGAGAATACAGTCATCAACTATGTCTTCTCCAACACGAAGCCTAAAAGGTTCATCTCCAAGATCACCTGCAATCACTACATATGGAAATGGATATCAATCCCCATGTGCATCACCATTAAATCCGTGTTCTCGTTATGCCAGAGTAAGCCGAGAGCCTCACTTGTCCTCAGCTATGAGCCCAAGTAGACCAAGGAAACTTCAGAGCATTAGCAATAACCAATCTGTGCACAACAATTAA

>S_lycopersicum_3

ATGGATCAATTTAGGCAAATTGGTGAAGTAGTTGGGAGTTTGAAGGCTCTTATGGTGTTGAAAAATGATATCCAAATCAATCAAAGGCAATGTTGTTTTCTATTTGATATGTTTGTTCATTCTTTTGATACAATTTCACAAGAAATCAAACATAACTTAAGGTTAAATGAGAGTAGTATAAAGTGGAAAGCTCTTGAGCTACCTATGAAAGAGCTTCATAGGATCTTTAAAGAAGGCGAATCGTACGTAAAATATTGTTTGGATGTTAAAGATTTGTGGGGGAAAGCCATAAGTCTCCATATGAACAGAGATTGTGTTGAGTTTCATATTCATAACTTGCTGTGTTGCTTCCCTGTTGTTATTGAAGCAATTGAGACTGCTGCAGAAATCTCGGGGTTCGATGAGGAGGAAATGCAGCAGAGGAGAACCGCGTTAATGAGGAAATACGATAGGGAATTGATTGATCCGAGGTTTTTTCAGTGGATGTCTGGGAAGCAGTATTTGGTAACTCGCGAAATATGTAGCCGGTTGGAGAGTTGTTGGAAAGAGGATAGATGGTTGCTTATCGATATGATAAGGCAAAAGAAAACAGAAACTTTGTTCAAGTATGAACAAAGGCTTGGAGACTTATTGTTAAGGAAATTAGATGGTGTTGAACCAATTAACAAGATATTGTTGCCTAGTTCAATCTTAGTCGGATCAAACGATTACCATGTGAAGAGACGTTTAGGGTCTTGGGGCGGACATGTTAAGGAGATACAATGGTTAGGTGAGAGTTTTGCTCTGAGGAACTTTTTCGGAGAGGTTGAACCGTTACATGATGAGATTTCTTTGGTGTTTTCTTTATCCCATCCCAACATTCTGCAATATCACTGTGGTTTTTATGATGAAGAGAGGAAAGAAGGGTATCTTGTTATGGAGCTGATGAATGATACTCTTACGACGTACATAAAAGATCATTCTGGTCAGAGGAAGAGACCGCCTTTTTCAACGTCTGCTGCAGTTGATATTATGCTTCAGATTGCAAAAGGGATGGAGTATCTGCATTCGAAAAAGATTTATCACGGAGAATTGAACCCCTCTCATGTACTTCTTAGAGCAAGGAATTCCTCAGCAGAGAGCTATTTTCACGCGAAACTCAAAGGTTTCGGATTAACTTCAATCAAAAGTACTTATAAGACTGCCAGCCACAATGCAGCTGATTCCGTTATATGGTTTGCCCCGGAAGTTCTTGCTGAACAAGAAAAACCAGGAAGCAAGTGTGCCTTCAAGTACACAGAACAAGCTGATGTTTATAGCTTCGGAATGATTTGTTTCCAAATTTTGACGGGGAAAGTTCCGTTTGATGAAGGACATTTGCAAGGGGAGAAAGTCGTGCGGAATATCAGAGCAGGGGAGAGGCCTCTCTTCCCATATCCTTCACCTAAATACCTTGTGAACTTAACAAGAAGGTGTTGGCATACCAATCCAAATCTTCGTCCACGTTTCTCGTCTCTATGCAGGATTTTACGTTACATCAAGAAGGTCCTTGTCATAAATCCGGAACATGGCCAGCCTGAAACTGCCCCTCCACTTGTAGACTACTGTGATATTGAGGCTAGCTACTCAAAGAAGTTTGCTGAAGAGGAGAGCACGAGTTTGATTCCCGTATCACAGATTCCTTTTCAAATGTTTGCTTATCGGCTCATTGAGAAAGAAAAGATTATTGGAAAGAACTGGGATCCATCACGTGATGGTTTTTCGGTCCACAGGAGAGCATCATTGCTGAGTGATGACGGGCAATTGTCTGCGATGGATGATCTCTTTCTTGCACCGAGTGATGGAAGGTCAGTTTGCTCTGAGATAATTGACAGGAAAGATTCAAGATTCTTTGATCAGAGGTCCGCTATCTCCGAGATACCACATAGAAGGCTTCTCTTATTCGATCAAACATCAGTTGGTTCTGAGAGTCCGGAGAGGAGATTCTCGTCTGTGGCAACAGCAGATGAAACACTTTTCTTTGCTGATAGTCCGGACAGGAAAGGAGCAGTATCATCACCATTAGTTAATAGGTCACCACGCATTGACTCATTAGAGAAAAAGATCGTTTCATCGATGGGCAAGAACCAAAGACTCAAACTTTTGGCCAACCAAGAGAAGGCAATGTCTCCAAAAGCAGGTGATGAAAAACCTTCATCAAGCGTAGCAGCTCCGCAAAAATTAGCATCTCCTCGGACTTCAGAGAAGATGAAATCAGCTGAACAAAATATAGTATCTCCTCCGACTTCAGAGGAAATGAAACCAGCTGCAAAAAATTTAGTATCTCCACGGACTACAGAGACGAAGAAATCAGCTGAGCCAAATTTAGTATCTCCACGGACTACAGAGATGAAGAAATCAGCTGAGCTAAATTTAGTATCTCCACGGACAACAGAAACGAAGAAATCAGCTGAGCAAAATGTAGTATCTCCTCCAATTTCAGAGAAGAAAATTTTATCTTATGATCAGAAACCGACTAGTTCTGAGACTCAAGAAAAGAAAAATGAATCAGATGATCAGAAACTAGTTAGTTCAGAGGCTCATGAGAAGAAACATTCATCTGATGATCAGAAATTACGACGTTCAAAGACTTTAGATCGGAAAAAGTCATTAGCTGCCAATGACAAACCACTTCATGGAGAGGAGATGCAAAGAACAAGTTCGAGAACCAGGAGACTAAACCAAAAGTTATTCAACATTGCAGACAAGAAAGTTCCTGAACCTAATCAAATTTCAATGTCATCCGAAAATCACGAGAAAATTGCATCCAAGCCCATCAACCAAAAGACAGCAGTCAAGAAAGATTCATCACGTAACAAAGTGCAGGACAGAAAGGCTAGCACAATACCAGAGAAACTGATCGACGACTCACCAATATCCTCACCAGCTAGAGGTAATAGAATACATTCATCGCCAATGTCTTCTCCAACACGATCCCCAAAGACATATTCAAGATCACCAGCAATCGGTTCAAACGGATATGGATATCAATCCCCAAGTTCATCACCATTAAATCCATGTTCACGATGTTCAAGAGTAAACCGAGAGTACCAAACTTCAGGTATGAGCCCACATACACAGAGGAAAGCTCATACCTGTCATTCGGAGATAGCTTAG

>S_polyrhiza_1

ATGGAGCAACTCCGGAAGATCGGAGAAGCTCTTGGGAGCCTCAAGTGTTTAATGGTGTTTCGAGACGAGATCCACATCAACACGCGTCAGTGTTCTCTTCTCCTCGATACCTTTAACCTCGCCTTCGAAGTAATCATGGAGGAAATCAGGACGAATCTGAGCTTCGACGAAAGGCTCACCAAATGGAAAGCCCTGGAGAATCCCTTGAGGGAATTCCACAGAATCCTGAGAGAGGGGGAGCAGTACATCCGACAGTCCATGGAGAGCGGCGACTGGTGGGGGAAAGCCATCTCCCGCGGGAGGAACACATACTGCGTCGAGTTCCACCTCCATGATCTGTTATGGTGCGTTGCCGTCGTGCTGGAGGCGATCGAGATCGCCGGCGACGTCGCAGGATGCAGCCAGGACGGCATTCGTCGGAACAGGATAATCCACTCCAAGAAATACGAGAGCGAATGGATGGATCCGAAGATATTCCAGCAGAGATTCGGCAGGGAGTACCTGATTTCTCCTAGCATTTGCAGTAGAATGGACTTCGCGTGGAAGGAGGACCAGTGGATGCTCCTGGAAAAGGCTTCAGAGAAGAGAAGGTCCTCGAAACAAGAAAATCGGCTGGCGGAGATTCTCGAAGGATCGAAGGGGAGAATCTTGCCGAGTTCAGTCCTGGTGGCCTCCAAAGATTATCAGGTGAGAAGAAGGGTAGGAAGCGAAAGCCATTACAGAGAAGTCCAATGGATGGGCGAGAATTTCGCTGTTCGCCATTTCTTCGGGGAGATCGAGCCACTGGTTTCGGAGATATCAATCCTCTCTTCACTCTCCCACCCGAACGTGATGCCATTATTGTGCGCCTTCTTCGACGAGGATCGGAAGGAGTGCCTCCTCCTCACGGAGCTGATGCAGAAGAGCCTCTGCAACTACATGAAGGAGATCTCCAGCCCGAGAAGGAGAGTTGCCTTCTGCCTCCCCGTTGCCGTTGACATCATGCTTCAGATTGCGAGAGGAATGGAGTACCTCCACTCGCGCAGGATCTACCACGGGAACCTAAATCCTTCCACCATCCTCGTCAAAACAAGGAATTCTTCTTCTTCTTCTTGTTCATCTGAAGGCCACCTACTGGTAAAAATTGCAGAGATTGGACTTGCATCCTCGAAAAATCTCAAGAACTCCACAAATCAGGCCTGTTCAAATCCGTGCATATGGTTCGCCCCAGAAGTTCTTCTGGAGCAAGAGCAGAAGGGAGATGCTTGCGAGTCCAAGCTCACGGAGAAGGCAGACGTCTACAGCTTCGGTATGATCTGCTTCGAACTGTTAACAGGGAAGGTGCCCTTCGAAGACGGGCATCTCCAAGGAGACAAGATGAGCAGGAACATCCGAGCTGGTGAGAGGCCATTGTTTCCATTCTCTTCTCCAAAGTACCTCACCAGCCTGACCAAGAGATGCTGGCAGACCGACCCATCACAGCGTCCGAGCTTCTCGTCGATTTGCAGGGTGCTCCACTACATCAAACGCTTTCTCCTCCTGAACCCAGATCACAGCCAGCCTGATGCACCTTCTCCACCCGTAGATATCTTCGATCTCGAAACGAGCTTCTCCAGGATATTCCCCACATGGGCGCTACCAGAACCCACGCCCGTTTCACAGGTTCCCTTCCAAATGTTTGCTTACAGGGTGATGGAGAGGGAGAAGAGCTCAGAGTCCGGAAGCGAAGGAGCATCCCTCTGCGGGGACGAGAACTCCTTCAGCTCTGGTTTCTCCGAGGATCAGTTCTCCCCGCCTTTGGCTCCCTCGAAGTCGTCGGCGCAGTTGAATCTCGAAGCGACCTGCAAGAAGGTGCCGCCGCAGAAGAAGACGGTCAACGGGAAGCCAGCCAAGGAATCAGGTCAAATTCTCATCTCCCCTCGTGAACGTTCCGCTCTTGAGGACGAACTCTGA

>S_polyrhiza_2

ATGGAGGAGCTACGGCGAATCGGGGAAGTGCTGGGCTCTCTCAAAACCCTAATGGCCTTCCAAGACGACATCGCCATTAACCGGCGGCAGTGCTTCCTCCTGGCTGACGCCCTCAACCTCGCCTTCGAGGTGGTCTCAGAGGAGATCCGGGAGAACCTGAGCTTCGATGAGAGGCTGACCAAATGGGGAGCTCTGCAACACCCGCTGCGGGAGCTACACAGGATCTGCGTAGAGGCCGAGCTCTACGTCCGCCATTGCATGGAGGGCAGAGATTGGTGGGCCAGCGCCATCGTCTCCCTCGGCCACACCGCCGACTGCGTCGAGCTCCACCTCCACAATTTGCTTTCCTGCCTCCCGGTGGTTGTGGAGGCCGTGGAAATCGCCGGCGAGATCGCCGGCAGCGACCCGGAGGAGATCCACCGGAGGAGGTTCATCTTCTCCAAGAGGTGCGAGAGGGAGTGGATGGAGCCCTCCCTGTTCCAGCGCCAGTTCGGGCGGCGCCGCCTGGTTTCTCCGAGCTTCTGCGCAGCCATGACCGCCGCCGGGAGAGAGGACCGGTGGGCCCTCCTCGAGACCGCCGCCGCCAAGAGGGGCGGCAGCCGGCTGGTGGCGCTGGTGGAAGGGTCGAAGGAGAAGCTCTTCCCCAGCTCGATCCTCGTGGCTGCCAAGGACTATCAGGTGAGGCGCAGGCTGGGGAGCGGCAGCCATTACAAGGAGGTCCACTGGATGGGGGAGAACTTCGCCGTGCGGCACTTCTTCGGCGACATAGCGCCGCTGGTGCCGGAGATCCTCCGCCTCTGCTCGCTGGCGCACCCCAACGTGATGCCCGTCCTATGCGCCTTCTCCGACGAGGAGAGGAAGGAGTGCTTCCTCCTTATGGAGCTCATGCAGAAGAGCCTCTGCAGCCACATCAAGGAGCTCTCCGGCCCGCGGCGGCACCGCCCGTTCCCCCTACCGGCGGCCGTCGACGTCATGCTGCAGATCGCCAGAGGCATGGAGTACCTCCACTCCAAGGGGATGCCCCACGGGAATCTGAGCCCCTCCCTCGTCCTCGTGAAGGGTAGCTCCGACGGCCACCTGCGGGTGAAGGTCACTGGCGTGGGGCTCTCCTCGGTGAAGAACCCCAAGTCCGCCGCCGCGGGCGGCGACCGCTGCATCTGGCTGGCCCCGGAGGTGCTGCTCGAGGAAGAAGAGCACGGAGGGAGGCTGCCGACGGGGGGCTCCACGAGGTACACGGAGAAGGCCGACGTGTACAGCTTCGCCATGATATGCTTCGAGCTCCTGACGGGGAAGGTACCCTTCGAGGACGGCCACCTACGGGGGGAGAAGATGAGCAGGAACATTAGGGCAGGAGAGAGGCCGCTCTTCCCCTTCCCCTCCCCCAAGCACCTCACCGGACTCACCAAGAAGTGCTGGCAGGCCGACCCGTCGCAGCGCCCGAGCTTCTCCGCCATCTGCAGAGTCCTCCGCTGGGTCAAGCGCTTCCTCCTCCTCAATCCCGACGCGGCGGCGCCGGGAGCGGCGGCGGACTACTTGGATATGGACGCCGCCCTGGCCAGGAGATTCCCCGGGTGGGCCCACCCGGAGACGACTCAGATCCCGTTCCAGATGTTCGCCTACAGGGTCATGGAGAGGGAGAAGGAGAGGAGCTCTGAGTCTGGGAGCGAAGGGGCCTCCGTCTGCGGGGACGATAACCTCTCCGAGGAGCTCCTCTCTTCTACGCCGTCGTCTCTCCCACTCTGCTCCGAGCTCGGGAAGGCGGGGCCGGGGAAGGAGGACCGAAGGGCCAGCATGGGATCGTCGTCGTCGGTGGGGCAGAAGCTGAAGGCGAAGACGGCGAGGCCGCCGCAGCTGGGATCATGCGCCGCCATACGGGCCCGCTCGGAGAACCAACTGCAGCCGATCAAGGCGAGCCCGGGGCGGCGGCGGTCCTCCGGACACGCCTCTGATTCAGATATAGCTTAA

>S_tuberosum_1

ATGGATCAATTTAGGCAAATTGGTGAAGTAGTTGGGAGTTTGAAGGCTCTTATGGTGTTGAAAAATGATATCCAAATCAATCAAAGGCAATGTTGTTTTCTATTTGATATGTTTGTTCATTCTTTTGATACAATTTCTGAAGAAATCAAACATAATTTAAGGTTAAATGAGAGTAATATAAAGTGGAAAGCTCTTGAGCTGCCTATGAAAGAGCTTCATAGGATCTTTAAAGAAGGCGAATCCTACGTAAAATATTGTTTGGATGTTAAAGATTTGTGGGGGAAAGCCATAAGTCTCCATATGAACAGAGATTGTGTTGAGTTTCATATTCATAACTTGCTGTGTTGCTTCCCTGTTGTTATTGAAGCTATTGAGGCTGCTGCAGAAATCTCGGGGTTCGATGAGGAGGAAATGCATCAGAGGAGAACCGCGTTAATGAGGAAATACGACAGGGAATTGATTGATCCGAGGTTTTTCCAGTGGATGTCTGGGAAGCAGTATTTGGTAACTCGCGAAATATGTAGCCGGTTGGAGTGTTGTTGGAAAGAGGATAGATGGTTGCTTATCGATGTGATAAGGCAAAAGAAAACAGAAACTTTGTTGAAGTATGAACAAAGGCTTGGAGACTTATTGTTAAGGAAATTAGATGGTGTTGAACCAATTAACAAGATATTGTTGCCTAGTTCAATCTTAGTCGGATCAAACGATTATCATGTAAAGAGACGTTTAGGGTCGTGGGGCGGACATGTTAAGGAGATACAGTGGTTAGGAGAGAGTTTTGCTCTGAGGAACTTTTTCGGAGAGGTTGAACCGTTACATGATGAGATTTCTTTGGTGTTTTCTTTATCCCATTCCAACATTCTGCAATATCACTGTGGTTTTTATGATGAAGAGAGGAAAGAAGGGTACCTTGTTATGGAGCTGATGAATGATACTCTTGCGACATACATAAAAGATCATTCTGGTCAGAGGAAGAGACCGCCTTTTTCAACGTCTGCTGCAGTTGATATTATGCTTCAGATTGCAAAAGGGATGGAGTATCTACATTCGAGAAAGATTTATCACGGAGAATTGAACCCGTCTCATGTACTTCTTAGAGCAAGGAATTCCTCAGCAGAGAGCTATTTTCACGCGAAACTCAAAGGCTTCGGCTTAACTTCAATCAAAAGTACTTATAAGACTGCCAGCCACAATGCAGCTGATTCCGTTATATGGTTTGCCCCGGAAGTTCTTGCTGAACAAGAAAAACCAGGAAGCAAGTGTGCCTACAAGTACACAGAGAAAGCTGATGTCTATAGCTTCGGAATGATTTGTTTCCAAATTTTGACAGGGAAAGTTCCGTTTGACGAAGGACATTTGCAAGGGGAGAAAGTCGTGCGGAATATCAGAGCAGGGGAGAGGCCTCTCTTCCCATATCCTTCACCTAAATACCTTGTGAACTTAACAAGAAGATGTTGGCATGCCAATCCAAATCTTCGTCCACGTTTCTCGTCTCTATGCAGGATTTTACGTTACATCAAGAAGGTCCTTGTCATAAATCCAGAACATGGCCAGCCTGAAACTGCCCCTCCACTTGTAGACTACTGTGATATTGAGGCTAGCTACTCGAAGAAGTTTGCTGAAGAGGAGAGCACGAGTTTGACCCCGGTATCACAGATTCCTTTTCAAATGTTTGCTTACCGGCTCATTGAGAAAGAGAAGATTTTTGGAAAGAACTGGGATCCATCAAATGATGGTTTTTCAGTCCATAGGAGAGCATCATTGCTGAGTGATGACGGGCAATTGTCTGCGATGGATGATCTCTTTCTTGCACCGAGTGATGGAAGGTCAGTTTGCTCTGAGATAATTGACAGGAAAGATTCAAGATTGTTTGATCAGAGGTCAGCTATCTCCGAGATACCACATAGAAGGCTTCTCTTATTCGATCAAACATCAGTTGGTTCTGAGAGTCCGGAGAGGAGATTCTCGTCTGTGGCAACAGCAGATGAAACACTTTTCTTTGCTGATAGTCCGGACAGGAAAGGAGCAGTATCATCACCAATAGTTAATAGGTCACCACGCATTGACACACTAGAGAAAAAGATCATTTCATCGATTAGCAAGAACCAAAGACTCAAATTTTTAGCCAACCAAGAGAAGGCAATGTCTCCAAAAGCAGCTGATGAAAAACCTTCATCAAGCATAGCTACTGCACAAAATTTAGCATCTCCACCGACTTCAGAGGAGATGAAACCAGCTGAACAAAATTTAGTATCTCCACGGACTACAGAGACGAAGAAATCAGCTGAGCCAAATTTAGTATCTCCACGGACTACAGAGACGAAGAAATCAGCTGAGCCAAATTTAGTATCTCCACGGACTACAGAGACGAAGAAATCAGCTGAGCCAAATTTAGTATCTCCACGGACTACAGAGACGAAGAAATCAGCTGAGCCAAATTTAGTATCTCCACGGACTACAGAGACGAAGAAATCAGCTGAGCCAAATTTAGTATCTCCACGGACTACAGAGACGAAGAAATCAGCTGAGCCAAATTTAGTATCTCCACGGACTACAGAGACGAAGAAATCAGCTGAGCCAAATTTAGTATCTCCACGGACTACAGAGACGAAGAAATCAGCTGAGCCAAATTTAGTATCTCCACGGACTACAGAGACGAAGAAATCAGCTGAGCCAAATTTAGTATCTCCACGGACTACAGAGACGAAGAAATCAGCTGAGCCAAATTTAGTATCTCCACGGACTACAGAGACGAAGAAATCAGCTGAGCCAAATTTAGTATCTCCACGGACTACAGAGACGAAGAAATCAGCTGAGCCAAATTTAGTATCTCCACGGACTACAGAGACGAAGAAATCAGCTGAGCAAAATGTGGTATCTCCTCCGATTTCAGAGAAGAAAAATTTATCTTATGATCAGAAACCGACTAGTTCTGAGACTCAAGAAAAGAAAAATGAATCAGTTGATCAGAAACTAATCAGTTCAGAGGCTCATGAGAAGAAACATTCATCTGATGATCAGAAATTACGACGTTCAAAGACTTTAGATCGGAAAAAGTCATTAGCTGCCAATGACAAACCACTTCATGGCGAGGAGATGCAAAGGAAAAGTTTGAGAACCAGGAGAATAAACCAAAAGTTATTCAACATTGTAGATAAGAAAGTTCCTGATTCTAATCAAATTTCAATGTCATCCGAGAATCACGAGAAAACTGCATCCAAGTCCATCAACCAAAAGACAGCAGTCAAGAAAGATTCATCGCGTAACAAAGTGCAGGACACAAAGACTAGCACTATACCAGAGAAACTGATCGACGACTCACCAAGATCTTCACCAGCTAGAGTTAATAGAATACATTCATCGCCAATGTCTTCTCCAACACGATCCCCAAAGACATATTCAAGATCACCAGCAATCGTCTCAAAGGGATACGAATATCAATCCCCAAGTTCATCACCATTAAATCCATGTTCGCGATGTTCACGAGTAAACCGAGAGTGCCAAACTTCAGGTATGAGCCCACATATACAGAGGAAAGCTCATACCTCTCATTCAGAGATAGCTTAG

>S_tuberosum_2

ATGGATCAATTCAGGCAGATTGGTGAGGTAGTTGGGAGTTTAACTGCTCTAATGATATTGAAACATGACATTCAAATAAACCAGAAGCAATGTTGTCTTCTGTTTGACATGTATGTACAAGCTTTCGACACGATTTCCGAGGGGATCAAACATAATTTAAGGTTGAATGAGAGGAACACAAAGTGGAAAGCACTTGAACATCCAATGAAAGAGCTTCATAGGATCTTTAAAGAAGGTGAAATGTACATAAAAAGTTGTTTGGATGTGAAAGATTTGTGGGGCAAAGCTATAAGTTTACATCTCAATAAGGACTGTGTTGAGTTTCATGTTCATAATTTGCTCTGTTGTTTCCCTGTTGTGATTGAGGCTATTGAGACAGTTGCAGAAATCTCGGGATTCGATGAGGAAGATATGCAGAAGAGGAGGACTGCACTCGCTAAAAAGTACGAGGGAGAGTCAATGTGTGATCCAAGATTCTTTCAATGGATGTTTGGGAAACAATACTTAGTTACTAGAGAGATATGTAGTGAATTGGAGAGTTGTTGGAAAGAAGATAGATGGTATCTTGTTGAAATGATAAGTCAAAAGATGAATGTTGCAGAAAATTTGGCGATAAACGAGCATAGTCTAGCTGAGATGTTGCTCAAGAAACTAGATGGATCAAAGCAACTCAGACAAATTATGCTCTTGCCTAGTTCAATATTGATTGGAGCAAGTGATTATCATGTGAAGAGACGTTTAGGCTCGCGAGGAGGACATGTTAAGGAGATTCAATGGTTAGGAGAAACGTTTGCGTTGAGGAACTTTTTTGGTGAGCTGATTGAACCAGTAGTTGCTGAGATTTCTCTAGTGTTTTCACTTTCACATCCCAACATTCTGCAATATCATTGTGGTTTTTACGACGAAGAAAAGAAAGAAGGATACCTTGTTATGGAGCTAATGAGTAAAAGTTTAGCAACTTACATAAAAGAGCATTCGAGTCAGAGGAAGAAAGGACCGTTTTCTGTACAAGTTGCTGTCGATATTATGCTTCAGATTGCTCGAGGGATGGAGTATCTGCACTCGATAAAGATCTATCACGGAGAGCTGAATCCATCTAACGTGCTCCTTAAACCAAGAAATCCCTCTGCGGAGAGTTATTTTCATGCAAAAGTTAAAGGATTTGGTTTAAAATCTATGAAGAGTAGTTATAAGGCTGCTGATGAAAATTCTGCTGAGTCTTTCATATGGTATGCACCAGAAATTCTCACTGAAAAGGAAAAACCAGAAAGCAAATGCACTTACAAGTATACAGAGAAGGCTGATGTTTATAGTTATGGAATGATATGCTTTCAACTTTTAACGGGGAAAGCTCCGTTTGATGAACATCTCCAAGGGGAGAAAATGGCGCAGAATATTAGAACAGGGGAGAGACCTCTCTTCCCGCATCCTTCACCTAAATATCTCGTAAACTTGACAAGAAAATGCTGGCAAACGAATCCAGACCTTCGCCCAAGTTTCTCTTCTTTGTGTAGGATTTTGCATTATATCAAGAAAGTACTTGTCATAAATCCAGGACATGGCCAACCTGAATGTCCTCCTCCACTCGTAGACTACTGTGAAATCGAGGCTGGCTACTCAAAGAAGTTCCCAGGAGAGGATAGCACTGGTTTGGCCCCGGTATCACAAATTCCTTTCCAGATGTTTGCTTATAGACTTGTTGAGAAAGAGAGAATATCTGGAAACTCTAAAGACAAGCACTGGGATTCATCACATTATGGTTATTCGAGCCACAGGACAGCATCAATGCAGAGCGACAATGAGCATATGGATGCAATGGATGATCTCTTTTTCGGTCCAAGTGATCGAAGGTCAATTTGCTCAGAGATAATAGAGAGTAAAGATCCAAGATTCTGGGATCAGAGGACAGTCATCTCTGAGACACCACTTCGAAAAGTTTCTTCATTCGAGCAAATGTCAGTTAGTTCTGAGAGTCCAAAAGAGAAATTCCCTGCAGCAGCAGCAAATGAAAAGTCAATCTATGATGATATTCTGGAAAGGAAGGAAATACTAACACCATCAGGTGATCAGTCCTCACCGCGTGTTGATACAACACCACGGAAGGCCATTTCATCAATGAGAAAGAACAAAAAACTGAATCTTTTGGAAATCCAGGAGAATGTAATGTCTACGAAAACAGATGACAGAAAACCTTCATCGAGTGAGCAGAAAACAGCGTCTCCTGTGATTTCAGAGAAGAAAAATTCATCAGGTGATCAGATAATAACTCATTCCGAGACTCAAGAAAAGAAACATGCACCAGATGATCAGAAACTAATAAGTTCTGCAACTCCAGAGGAGAAAAACTCATCTGATAACCAACAACCAAAAGATTCTCAAAGTCCAGAAAAGAAGCAGTCATCAACAGCTGCCAATGTCAAGACCATTCGTGGTGATCCTACGCAAAGGAAAGCTTTGTCAAGAAGAACAAAATCCAAGAATCCAGAGAAGAAAGTTCTATCAAGTCCGGAAGCCAATCAAAACTCAATGTCATCTGAGATAACAGCATCGAAACCAAGCAATTCAAAGGTAACTGCAAAAAAAACGCCATTTCACAAAAAACTACAGGACAAAGAGAGTAGCACTACACCAGAAAAACTGACAGACCCATCACCAAGATCTTCACCAGCTAGAGCTAAGAGAATACAGTCATCAACTATGTCTTCTCCAACACGAAGCCTAAAAGGTTCATCTCCAAGATCACCTGCGATCACTACACATGGAAATGGATATCAATCACCATGTGCATCACCATTAAATCCGTGTTCTCGTTATGCCAGAGTAAGCCGAGAGCCCTACTTGTCCTCAGCTATGAGCCCAAGTAGACCAAGGAAACCTCAGAGCATTAGCAATAGCCAATCTGTGCACAACAGTTAA

>S_tuberosum_3

ATGCCGGCGTTAGAGGAATTCAGGCAGATAGGAGAGGTAATTGGGAGTCTTAAAGCTCTAATGGTATTCCAAGATGACATACAAATAAATCAAAAACAATGTTGTTTGTTAGTGGATATGCTCAAATGTGCCTACAAGACAATTGCAGAAACAATGAAACAGAACTTGAGATTTGAAGAGAAGAATATTAAATGGAAAATTCTTGAGAACCCCTTAAGAGAGCTTCTCAGGGTGTTCAAAGAAGCAGAACAGTACATCAAGCAATCCCTGGAAAATAAGGATTTTTGGGCAAAAGCCATAGTTCTATATAAGAATACAGATTGTGTTGAATTTCACATCCACAATTTGCTCTCTTGTGTGCCAATTGTCATCGAGGCCATAGAAATAGCAGGAGAGATTTCAGGTAGTGACCATGACGAGATACAGAAGAAGAGATTCATTTACTCGATGAAGTATCAGAAAGAGTGTAAGGACCCACGAATCTTCCAATGGAAATTCGGAGAACAGTACATGGTTTCTCAGAAGTTCTGCGAAAGGGTATGCTCAGTTTGGAATGAAGACAAGTGGATTTTGCAAAATAAAATTCGAGAGAAGAAGAATTTAGGTGCATGTACTTTGACAAAGCATGAGAAACGACTTGCTGATCTCCTCTTGAAAAACTTAAACGAGATGGAGATGGAGAGTGATCATAAGCTTTCACCTAGCTCACTTCTTGTAAATTCTAAGGATTACCAGGTAAGAAGACGATTAGGTAGTGGAAGTAAGTACAAGGAGATCCAATGGTTGGGGGAGACCTTTTGTTTAAGGCATTTGTTTGGGGATATTAAGCCCCTCATTCCAGATATTTCGCAAGAACTGCATCTCTCTCATCCAAATATAATGCACATCTCCTGTGGTTTTACTGACGAAGAAAAGAGAGAATGTTTCTTAATCATGGAACTTATGAGCAAAGATTTATCTAGTTGCATCAAAGAGATCTGTGGACCAAGAAAGCGTGTACCGTTTTCTCTTCCCGTTGCAATCGATTTACTCCTTCAGATTGCAAGAGGCATGGAATACCTACACTCAAAGAAAATCTATCATGGTGAATTGAGTCCTTCCAACGTCCTTGTCAAGGCGAGGAACGTATCTACAGAAGGGTATTTACATGCAAAGGTTTGTGGATTTGGCTCATCGTGTTCTATCAACCTTCCTCAGAAAGCTAATGTGAATCAAAATAATGGGACACTCCCATTCATTTGGTTTTCCCCAGAAGTCCTAGCTGAACAAGAGCAGTCGGGGAATGGAGGAAACATCAAGTATACTGAGAAATCTGATGTGTACAGTTTTGGGATGATATGTTTTGAGGTTTTAACAGGGAAAGTTCCATTTGAAGATAGTCATCTACAAGGAGATAAAATGAGTAGAAATATAAGGGCAGGAGAGAGACCATTATTCCCATTTCACTCACCAAAATATGTGACTAGCTTGACAAAGAGGTGTTGGCATACTGATCCATATCAACGACCAAGTTTTTCATCCATCTGCAGGGTTCTACGCTATGTGAAGAGATTCTTGGTGATGAATCCTGAACACAGCCAACAAGACTCACCATTGCCACCAGTAGACTATGGAGAGATTGAAACTGTCATCTTAAGGAGCTTCCCCTTCTTAGGAAATTCTGAATCTGATCCCCTGCCTGTCACACAAATACCTTTCCACATGTTTGCATACAGGGTGACCGAGAAAGAAAAGTCAAGCACAATTCACAGAGACGTAAATTCTGAATCAGGAAGTGATGGAACTTCAGCATGTGGGGATGATCCTGTAACGGCAGATGATGCACTTCCATCACCAACCGAAAAGAAGAATATTGCATCTCCCGAGATTTTGACCAAGAGACTTTCAATTAGGAAACCTGCTGATATCAAAGTCGGCAAGCAACCAGGAACACCAAGGGGAAGGACGGTAAGACCTCCAAACATACGTACTATAAGGCAAAATTCTGAAAGTCAGTTAATGATGATGAACAGCCCAAGAACACGGAGATCATCTGGCCATGCATCAGATTCGGAGCTTCCTTAA

>T_cacao_1

ATGGAGCAATTCAGGCAGATTGGTGAGGTTTTGGGAAGTTTGAGAGCTCTCATGGTTTTACAAGATGAGGTCCAAATCAATAGACGCCAGTGCTGTTTGTTGTTTGATATCTTCTGTTTGGCTTTCAATACGATAGCCGAGGAGATCAGGCTGAACCTGAAACTAGAAGAAAAGAACACGAAATGGCATGCCCTTGACCATCCCTTGAGAGAGCTTCACAAGATCTTTAAAGAAGGAGAACTCTATGTTAAACAATGTATGGACAAGAAAGACTGGCGGGTGAAGGCAATCTACCTCCACCAGAACAAGGATTGCGTTGATAATCACATTCACAATTTGCTTTGCCACTTCCCGGTTGTCATTGAGGCCATCGAGACTGCAGGAGAGATAGCAGGGCTTGACCAGGATGAGATGCAAAGGAGGAGAGTTGCACTTGCAAGGAAGTACGACAAGGAATGGAATGACCCAAAACTCTTTCAGTTTAGATTCGGAAAGCAGTATTTGGTCCCTCGAGATATTTGTAGCCGGTTTGAGAGTGCTTGGAGAGAAGACAGGTGGAATCTTGTTGAAGCTCTCAAGGACAAGAAAAATTTGGATACTGCAACCAAAAACGAGCAGCGGCTTGCTGGCTCGTTGATTAAGAAAATTATTGGATCAGAGGCTTATAAGGGTAAACTTTTCCCGAGTTCAATTTTGTATGGAGGCGATTACCTAGTTAGGCGACGATTAGGGGCACAGTATAAGGAAATCCAATGGTTGGGAGATAGCTTTGTATTGAGACACTTCTTTGGGGAGGTTGAGCCTTCATGTTCTGAAATCTCTACCCTACTTTCACTTTCTCATCCTAACATATTGCAATACCTTTGCGGGTTTTATGATGAGGAAAAGAAGGAGGTCATGCTTGTTCTAGAGTTGATGAACAAGGATCTCAGTTGTTACATGAAGGAGAATTGTGGCTCGAGAAGGAGAATCTTGTTCTCTTTTCCTGTTGTGGTTGATCTTATGCTTCAGATTGCAAGAGGAATGGAATATCTCCACTCACAGAAGATCCGTCATGGAGAGCTAAACCCTTCTAATATCTTTCTGAAAGCTAGGAACAACACAGAAGGTTTTTTCCATTTAAAAATCTCGGGCTATGGTTTATCCCCTGTCAAAACTCGTTCTTCTCCGAACTCATCACCAAAGCCAATTGAGCCAAACCCTTTTATTTGGTACGCTCCGGAAGTTTTGCTAGAGCAAGAACAGCCAGGAAATATTACCATTTCCAAGTACACAGAGAAAGCAGATGTCTACAGCTTTGGGATGCTTTGCTTTCAGCTCTTGACGGGGAAAGTTCCTTTCGAAGATGGACATCTCCAAGGAGAGAAGATGAGTAGAAATATTAGAGCAGGGGAGAGGCCTCTCTTTCCATACACAGCACCTAAATACCTCGTCAACTTAACCAAAAGATGCTGGCATACTGATCCAAATCAGCGTCCAAGTTTCTCATCCATTTCTAGGATCTTGCGCTACATCAAGAAGTACCTTGTGATGAATCCTGACCGTGATCAGCCTGAAATGCAATCCCCAGTTGCAGATTACTGTGAGATAGAGGCATGGTTCCTGAAAAAGTTCACCGCAAATGGGACTTTCAGTCCATTATCAGTAGCACAAATCCCATTTCAAATGTTTGCTTATAGGCTTGCTGAGAAAGATAAATCTATTTTGAACACCCAGGATAAGAATGGGGAGTTAGCCAGCGAAGCAGCCTCAATTTGCAGAGATGAGAATAATTCTACAGTGGAGGATCCCTTGATAGCAGCAAGTGAAACAAAGTCTGTTTCTTCTGATGTAAAATCGGTTTGTTCAGATATCAAGTCAGTCTATTCAGATATCAGGTCAGTCTGTTCTGAAATTCCAGAAAGGAGATCAATCAATTTCGATACACCACAAAAGAGATCGATTTCCACCAAGATTCCAGAGAAGAAAAAAATGTTACTGATGAAGAAAAACACTAACGTGAAGGCGAAAAACTTTACAGGGACACCAAATGCACAATCTCCTCGACCACCAACTCTGAATCGTGGTCACAGTGTAAGGATAAACCGAGAAAGTCGCTCACCATTAATAACGAGTTCCGTGAGCAGAGGCCGGCAAAGAGCAGCAGCAGGTCATACCTCGGATTAG

>T_cacao_2

ATGCCTCCTGGTTTTAGACCAAAAATGGAGCAATTTCGACAGATCGGGGAGGTGCTAGGGAGTTTGAAGGCTTTAATGGTATTTCGTGACAGCATCCAGATCAACCAGAGGCAATGCATTTTGTTACACGATATGTTTAGTTTCGCATACAGATCGATAGCAGATGAAATGAGAGAGAACCTGAAATTTGAAGAAAGGAACATCAAATGGAAAGTTCTTGAAATGCCTTTCAGAGAGCTCCACAGGGTTTTCAAAGAAGGGGAAGCCTACATCAGGCAGAGCTTGGAATCCAGAGACTGGTGGGCTAAAGCCATTACTCTCTATCAGAATTCGGATTGCGTCGAGTTACATATCCATAACTTGCTTTCCTGCATCCCTGTTGTCATCGAAGCAATCGAAACTGCAGCAGAATTATGTGGTTGGGAACAAGATGAAATGCAAAAGAAGAGGCGGGTGTACTCCAACAAGTACCATAAAGAATGGATTGATCCTCAGCTTTTCCAGTGGAGATTTGCAAAGCAGTACCTTATTACACAAGATTTTTGCAATAGGATTGACACTGTTTGGAAAGAAGATAGATGGATCCTTCTAAACAAAATCCTGGAAAAGAAAAGCATGGGTTCGAGAAAGCAGGAGCGCAAACTTGCAGATTTGCTTTTGAGAAACTTAGATAGTTCAGAGTCTTTGAATGGGAGGCTCTTACCGAGTACAATGCTATTAGGGTCTAAGGACTACCAGGTTAGACGGCGGCTTGGAAACGGGAGTCAATATAAGGAGGTCTACTGGTTAGGTGAAAGCTTTGCTTTGAGACATTTTTTTGGAGACGTAGAAGCAGTAGCTCCTGACATTTCTTCTTTATTATCTCTTTCCCACCCAAACATACTGCATTTCCTCTGTGGATTTACTGATGATGAGAAGAAGGAATGTCATTTGGTCATGGAACTAATGAATAAAAGCCTCCGCAACTACGTAAAAGAGATTTGTGGCCCAAGAAAGCGAGTACCATTTCCTCTTGCGGTTGCTGTTGATCTAATGCTTCAAATAGCGAGAGGAATGGAATATCTCCACTCAAATAAAATCTACCACGGTGATTTAAATCCTTCCAACATTCACGTCAAGTCAAGAGGAATGTTCACGGAAGGGTACATGCAGGCAAAGGTTTCAGGATTTGGCTTATCTTCCATAGTTCATTTACCACAAAAGAACACATTGATGAACCAGAATGAAACCCTGCCATTCATCTGGCATGCTCCAGAGGTTCTAGAAGAGCAGGAGCAGACAGGAAGTAAAGGGAATTCAAAGTACACAGAAAAAGCTGATGTTTACAGCTTCGGAATGATTTGCTTTCAGCTTCTAACTGGAAAAGTCCCATTTGAAGATAGCCATCTTCAAGGGGACAAAATGAGCCCAAACATCAGGGCAGGGGAGAGGCCATTATTTCCTTTCCAGTCACCAAAATGTGTAACCAACCTGACCAAGAAATGTTGGCATGCTGATCCAAATCTACGGCCTAGCTTCTTATCCATCTGCAGAATTCTTCGGTATGTAAAACGGTCCCTTCTAATGAACCCTGATTATTATAACAACCAGTCAGAATTACCAATGCCAGTGGTGGATTACTGTGACATTGAGTCAAAGCTTCAACGAAAATTCCCCACATGGGAGGCTTCCAATCCGTTACCTATATCACAAATCCCTTTTCAGATGTTTGTTTATAGAGTACTAGAAAAGGAGAAAATAGGTCATTCTCTCAAAGACACTTCAGAATCAGGAAGTGACAGAACTTCAGTTAGTGGGGATGAAAATGCCACTGCTGATGACCCATTGTCATCAACAACTGAGAGGAGGTCCTTACCTTCGCCTGAATCGACTCCCAGGAGACTTTCAGCATTGAAAAAATCTCCTGATATTAGAGCAAAACATCCAGTGACACCAAAAGGAAGAGCAGTAAGACCTCCTCAACTGAGCCGTTGTGGGCGCAGTTTAAGAATGAATTCAGAAAGCCAGCTACTTCTAACAAGTCCAAGAATACGGAGGACAGCATCCGGTCATGCCTCAGATTCCGAGCTCTCCTAG

>S_parvula_1

ATGGAGCAATTCAGACAAATCGGAGAGGTTTTGGGAAGTCTAAATGCACTAATGGTATTACAAGACGATATCTTGATCAACCAAAGACAGTGTTGTTTGTTGTTAGAGATCTTCAGCTTAGCTTTCAACACTGTCGCAGAAGAGATCAGACAGAATCTGAAGCTTGAAGAGAAGCATACTAAATGGAGAGCCCTTGAGCAACCCTTGAGAGAGCTTTACAGAGTGTTTAAAGAAGGAGAGTTGTATGTTAAGCATTGTATGGACAATAGTGATTGGTGGGGCAAAGTTATCAATTTTCATCAGAACAAAGATTGTGTTGAGTTTCATATACACAACCTGTTCTGTTATTTCCCGGCGGTTGTCGAAGCGATTGAGTCTGCCGGGGAGATCTCGGGTCTTGACCCTTCTGAGATGGAGAGAAGAAGAGTTGTCATCTCAAGAAAGTACGATAAAGAGTGGAATGATCCTAAGCTATTTCAATGGAGGTTTGGGAAGCAATATCTTGTACCGAGAGATATTTGCGTCCGGTTTGAGCATTCATGGAGAGAAGATAGATGGAATCTTGTAGAGGCATTGCAAGAGAAGAGGAAATCGAACAGCGATGATATCGGAAAGACCGAGAAGCGTTTAGCTGATCTGCTTCTGAAGAAACTAACCGGCTTGGAGCAGTTTAACGGGAAACTGTTCCCGAGTTCGATACTTCTTGGCTCAAAGGATTATCAGGTGAAAAGGAGGTTAGATGCAGATGGACAGTACAAGGAGATTCAATGGCTAGGTGATAGTTTTGCAGTGAGACACTTCTTCAGCGATCTTGAGCCGTTAAGTTCTGAGATTTCATCTCTTTTAGCTCTTTGTCATTCGAATATACTTCAGTACCTATGTGGATTCTATGAAGAAGAACGGAGAGAATGCTTTTTGGTTATGGAGTTGATGCACAAAGATTTGCAGAGCTACATGAAGGAGAATTGTGGACCAAGAAGAAGATATCTCTTCTCTGTCTCCGTCGTGATTGATATCATGCTGCAGATTGCAAGAGGAATGGAGTATCTCCATGGGAATGACATCTTCCATGGAGATTTAAACCCAATGAACATTCTTTTGAAAGAGAGGAGTCACACCGAAGGTTATTTCCACGCGAAAATCTCTGGATTTGGTTTATCTTCTGTGAAAACTCAGTCTTCTTCTTCTTCTTCTTCTCGGTCTTCCTCGAGACCAAGCACTCCTGATCCTGTGATTTGGTATGCACCTGAAGTTCTAGCAGAGATGGAACAAGACTTGAAAGGCACAACAACTCCAAAATCGAAGATGATGCACAAAGCAGACGTGTATAGCTTTGCAATGGTGTGTTTTGAGCTTATAACAGGTAAAGTTCCGTTTGAAGATAGTCATCTACAAGGTGAGCAAATGGCTATCAACATTAGGATGGGAGAGAGACCTCTTTTCCCTTTCCCTTCACCGAAACACCTCGTTAGTCTGATTAAACGGTGTTGGCACTCAGAACCGAGCCAGCGTCCAAACTTCTCTTCCATCTGCCGGATTCTACGGTACATCAAGAAGTTCCTAGTTGTGAATCCAGATCATGGTCACCCTCAAATTCAAACGCCACTAGTTGACTGTTGGGATTTAGAAGCAAGGTTCTTGAGAAAGTTTCCAGGTGACTCAGGATCTCACACAGCGTCCGTGACTCAAATCCCTTTCCAACTATACTCATACCGTGTTTTCGAAAGAGAGAAGATGAATCCAAACTCACAAGAAAGTAGTAACTCAGAGGCAAGTGAATCAGAAAGCGTTTCAGTGGTTGAAGATCCACCTAATGCCATGACAATAAGAGATACAAAGTCTTTGTGTTTAGATACAATGTCTGAATACTCAGATACAAGATCAGTCCACTCAGAGGCTTCCACCAAAAAGGTCTCCGTTTTAAAGAAAACGGGTGAATTGGTAAAGCTAAGAAAAAGTCCAAGCTTAGGTTCAGAAAAGTTGAGATCTACAGGGACTTCGCCGGTAAAAGCAAGATCATCGCCAAAAGCTTCACCGTTGAGTCCATTTGGAAGAACCATTAAAGCGAGGAAGGATAATCGATTGCCTTTGAGTCCAATGAGTCCTCTAAGTCCAGGAATACGTAGACAACGAACTGGTCATGCTTCAGACTCTGAGCTTACTTAG

>S_parvula_2

ATGGAGCAATTCAGGCAAATCGGCGAGGTTCTAGGAAGCTTAAACGCGCTCATGGTATTACAAGACGATATCTTGATCAACCAAAGACAATGCTGTCTCTTGTTAGAGATTTTCAGTTTGGGTTTCAACACGGTAGCCGAAGAGATCAGACAAAACTTGAAGCTCGAAGAGAAGCATACCAAATGGAGAGCCCTTGAACAGCCTTTGAGAGAGCTCTACAGAGTGTTTAAAGAAGGCGAAATGTATGTTCGAAACTGTATGTCCAATAAAGATTGGTGGGGCAAAGTTATCAACTTTCATCAGAACAAAGATTGTGTTGAGTTTCACATACACAACTTGCTCTGTTACTTTCCGGCCGTGATCGAAGCCATCGAGACAGCCGGAGAGATTTCTGGTCTTGACCCTTCTGAGATGGAGAGAAGGAGAGTTGTGTTTTCGAGAAAGTACGATAGAGAGTGGAATGATCCGAAGCTATTTCAATGGAGGTTTGGGAAACAGTATCTGGTTCCGAGGGACTTTTGCATCCGGTTTGAGCATTCGTGGAGAGAGGACAGATGGAATTTAGTGGAGGCATTACAAGAGAAGAGAAAGTCGAAGAGCGATGAGATTGGGAAGACCGAGAAGCGTTTAGCTGATTTTCTGTTGAAGAAACTAACCGGATTGGAACAGTTTAACGGTAAGCTCTTCCCGAGTTCGATTCTTCTTGGTTCTAAAGATTACCAAGTGAGGCGGAGACTAGGTGGTGGTGGTCAGTACAAAGAGATTCAATGGTTAGGTGATTCTTTCGTGTTGAGACATTTTTTCGGGGAGCTTGAGCCGTTAAATGCAGAGATCTCTTCTCTGTTGTCGCTTTGTCACTCGAATATACTTCAGTACCTATGTGGATTCTATGATGAAGAAAGGAAAGAATGTTCTCTGGTTATGGAGTTAATGCACAAGGACTTGAAAAGCTACATGAAGGAGAATTGTGGACCAAGAAGAAGATATCTCTTCTCTGTTCCCGTTGTGATCGATATAATGCTTCAGATCGCAAGAGGGATGGAGTATCTTCATTCAAATGAGATCTTCCATGGAGACTTGAATCCCATGAATATTCTATTGAAAGAGAGGAGTCACACCGAAGGTTACTTCCACGCGAAGATCTCAGGTTTCGGTTTGATCTCAGTCAAAAGTCAGTCTTTTACTCGGGCGTCCTCTAGACCAACCACTCCTGATCCTGTGATTTGGTATGCACCTGAGGTTCTAGCAGAGATGGAACAAGATCTCAAAGGTATAGCTCCGAGATCCAAGTTTACGCATAAAGCTGATGTTTATAGCTTTGCTATGGTGTGTTTTGAGCTTATAACCGGTAAAGTTCCATTTGAAGATAGTCATCTTCAAGGTGATACGATGGCTAAGAACATTAGAATGGGAGAGAGACCTCTCTTTCCATTCCCTTCTCCTAAATACCTTGTTAGTCTAATCAAACGGTGTTGGCATTCAGAACCGAGCCAGCGTCCGACTTTCTCTTCGATTTGCCGGATACTACGCTATATCAAGAAGTTCCTAGTTGTGAATCCGGATCACGGTCACATTCAAATCCAAAACCCACTTGTTGATTGCTGGGACTTGGAAGCAAGATTCTTAAGAAAGTTCTCACTCGAGTCAGGGTCTCACGCGGAATCTGTAACACAAATACCGTTCCAGCTTTACTCATACAGGGTTGCAGAGAAAGAGAAGATGAGTCCAAACTTCAACAAGGAAGAAAATTCGGAAACAGGTGAAACTGCTTCTGAAAACGTTTCGGTGGTTGAAGATCCACCAACAACAACGCCAATGTATACAAAGTCATTGTGCTTAGATGCAATATCTGAATACTCAGAGTCAGATACAAGATCAGTTCACTCAGAAGCTCCCATGAAAAAGATCTCAGCTTTAAAGAAAAGCGGCGACATGACGAAACTCAGAAGAAATTCAAGCGCAGGTTTACGGTCTTCAGGATCATCGCCAGTAAAGCCAAGATCAACACCGAAAGTGTCATCACCATTAAGTCCATTTGGAAGAAACAGCAAAGCAAGGAAGGATACAAGATTGCCTTTAAGTCCTATGAGTCCTTTGAGCCATGGAAGACGCAGACATCTCTCTGGTCCTGCTTCAGACTCTGAGCTAACATAG

>S_parvula_3

ATGGAGCAATTTCGAGAGATAGGAGAGGTATTAGGAAGCATAAGAGCGTTAATGGTGTTCAAGGACAACATTCAAATCAACCAACGTCAATGCACTCTATTACTCGACCTCTTCACCGCGACATACGATTCAGTTTCCGAATCTTTGATATCAAATCTACGTTTCGGAGAGAAGAACACCAAATGGAGGATTCTAGAACAACCCTTGAGGGAGCTTCTATGGGTGGTACGAGAAGGGGAGGCTTACGTTAGAATGTCTTTAGAGCCTAAACTTGGGTTTTGGGCTAAAGCTATTGTGTTACAACACAACAGAGATTGCACAGAGCTTCACATTCATAACTTGCTGTCTTGTGTACCGATTATCATCGAGGCTATTGAGATGGCAAGTGAGGTTTCAGGGTGGGATGAAGAAGAGATGAACAAGAAGAGATTGGTGCATTCTAACAAGTATATGAAGCAATGGAATGACTCTCAGATGTTTACCTGGAAGTTCGGGAGAGAGTATTTGGTGACCGCAGACTTTTGCAGTCGGTATGAGAGTGCTTGGAGAGAGGACAGGTGGATTCTAATGAAGGAGCTTCAGGAGAAAAGGCGCCCCGGTTCAAGCAAACACGACAGGAAGATGGCTGATTTCCTTTTGAAAAATTTAGGAGATGGAAACGAGAGTCCCAAGCTATTTCCGTCTTCTATTCTGGTCAGCACGAAAGATTACCAAGTGAAAAAGAGATTAGGCAATGGAAGTCAGTATAAGGAGATTACCTGGTTAGGCGAAAGCTTTGCGCTTAGGCATTTCTTTGGGGATGTTGATGCTTTGCTTCCTCAAATTACTCCATTGCTGTCTCTTTCACACCCAAACATTGTATATAACCTCTGCGGATTCGCGGATGAGGAGAAGAAAGAGTGTTTCTTGGTCATGGAACTGATGAGTAAAAGCCTCGGGACTCACATCAAAGAGGTATGTGGTCCGAGGAAGAAGAACACTCTCTCCCTCCCAGTTGCAGTTGATCTGATGCTTCAGATAGCTCGGGGTATGGAATATCTTCACTCGAAAGGAATATACCACGGGGAGTTGAATCCATCTAACATTCTTGTCAAACCAAGAAGAAATAACCAATCCGGAGATGGGTATCTCCATGGGAAGATTTTTGGGTTCGGGTTGAATTCTGTCAAGGGCTTCTCCTGTAAACGTGTTTCTTCAATGGGACAGAATGAGAATTTTCCATTCATATGGTATTCCCCGGAGGTGCTAGACGAACATGAACAGAGCGGAACTGCAGGAAGCCTTAAGTATACTATTAAATCTGACGTGTATAGCTTTGGGATGGTTTGCTTTGAGCTTCTGACAGGTAAAGTACCGTTTGAAGATAGCCATCTGCAAGGAGATAAAATGAGTAGAAACATTAGGGCAGGGGAGAGACCGCTTTTCCCATTTCATTCACCTAAATTCATAACAAACCTGACAAAAAGATGCTGGCATGCTGATCCGAATCAGCGTCCAACATTCTCGTCGATAAGCAGAATTCTCCGATACATCAAACGGTTTCTAGCTTTAAATCCGGAATGTTACAGTAGTAGCCAACAAGACCCGCCCGTTGCCCATCCTGTGGATTACTGTGAGATTGATGCTAAGCTGTTGCAGAAGCTTTCATGGGAATCGACAGAGTTACTTCAAGTTTCACAGGTTCCTTTTCAAATGTTTGCATATAGGGTTGTGGAACAGGCAAAGACTTGTAAGAAAAACAATCACCGAGACACATCTGAATCGGGTAGTGAATGGGCTTCGTGTAGTGAGGATGAGGGAACTCCCAGAGGACGATCAAGGCATCCCCCACTGAGCCCTTGTGGTCAAAGAATGAGAACGAACTCTGAGAGTCAACTGACACTTCTGAGTCCAAAGATACAGCGATCAAACTCTGGTCATGTCTCTGACTCTGAGCTTTCCTAG

>V_vinifera_1

ATGGAGCAGTTTCGGCAGATTGGAGAGGTTTGGGGGAGTTTGAAGGCTCTAATGGTTTCAGCCTTCAAAACAATTGTAGAGGAGATCAGGCAGAACCTGAGATTAGAAGAGAAGAACACCAAATGGAAAGCTCTTGAACAGCCTTTAAGAGAGCTTCACAGGGCTTTTAAAGACGCAGAACTATACATCAGGAACTGCTTGGACACCAAGGACTGGTGGGGCAAAGCAATGAGCCTCCACCAGAACAATGACTCTGTTGAGTTTCACATCCATAACTTGCTCTGCACTTTTCCTATTCTTTTCCAGTGGAAATTTGGAAAGCAGTACCTGGTTACCCGAGAAATTTGCAGTAGGTTGGAGAGTGCTTGGAAGGAAGATAGGTGGCTTCTTCTTGAAATGATTAGGGAGAAGAAAAGTTCAGGGTCAGTTGGGAAGAATGAGCAGAAGCTTGGAGACCTGCTGCTTAAGAAACTAAATGGATCAGAGAAGTTCAACGGGAAGCTTTCCCCGGGTTCAATTCAGTGGTTGGGGGAGGGTTTTGCTTTGAGGCAATTTTTTGGGGAGATTGAGCCATTGAATTCAGAGATTTCTTCTCTGTTGTCACTTTCCCACCCCAACATAATGCAGTACCTTTGTGGATTTTATGATGAGGAAAAGAAAGAGTGCTTACTTGTGATGGAGATGATGAACAAGGATCTTCACAGTCAGATTAAAGAAAATTGTGGACAAAGGAGGCGTATTCTGTTTCCTCTCCCGGTCATGGTTGATCTCATGCTTCAGATTGCCAGAGGCATGGAGTATCTTCACTCTGTGAAGGTCTATCATGGAGATTTAAACCCTTCTAATATTTTTCTGAAGGCAAGGAACTCCTCCACTGAAGGTTACTTCCATGCAAAAGTTTCGGGTTTTGGGCTATCGTCCATCAAGAATCACACTTTTTCTCGGAGCTCCCCAGGCCAGAATGGAACTGACCCTCTCATTTGGCATGCACCAGAGGTTCTGGCTGAGCAAGAACAGCTAGGAAGTAGTTGTAGTTCCAAGTTCTCAGAGAAAGCTGATGTTTACAGTTTTGGAATGCTTTGTTTTGAGCTTCTGACTGGGAAAAGGCCTCTTTTCCCATTCCCTTCACACAAGTACCTTGCAAACCTAGCCAAAAGATGCTGGCACACTGACCCAGTTCAGCGCCCAAGTTTCTCTTCCATATGTAGAATTCTTCGCTACATCAAGAGGTTCCTTGTGCAGAACCCTGATCACAGTCAACCGGAACTGACACAGCCACCTGTAGATTACTTTGAACTGGAGGCAGGGTTTGTGAAGAAGCTTACCGTGGAGGGGCAACTTGATCCAATGCCTGTATCACAAATTCCTTTTCAGATGTTTGCTTATAGACTGGTAGAAAAAGAGAAGACCTCACTGGACTATAAAGACAAGAGCTCGGAATCAGCAAGTGAGGCAGCAGCTTCAGGTTCTGGGGATGAGAATGTTGCGCTAGTAGATGATCCATTTCTGCAAGCAAGCGAAGGTAGGTTAGTTTGTTCAGAGACTCCAGAAAAGAAAATATCACCCGCAAAGATATCTGCAGAATCAAAAACCCGAAGAAGTCTAGGAACACCAAAAGCTCAAGCTCAGAAACCGACACCATTGAAACCATGTGGCGGCATGAAACTGAATCGAGGGGCACAGATGGGACTGGGAAGAACATCCGGGCATGCCTTAGATTCAGATGTATCATAG

>V_vinifera_2

ATGGAGCAATTCAGGCAAATGGGAGAGGTTTTGGGGAGTTTGAAGGCATTTATGGTGTTCCGGGAAGATATTCAGATCAACCAGAGGCAATGCTGTTTGTTGCTGGATGTTCTCGACTTGGCTTATCACACAGTAGCAGAAGAGATGAAGCAGAATCTTCGATTAGACGAGAAGCATACAAAATGGAAAATTCTTGAACAGCCCCTGAGGGAGTTGCACGGGGTATTCAAAGAAGGAGAACACTACGTAAGACAGTGTTTGGACACGAAAGAATGGTGGGCTAAAGCCATTATATTCTATCAGAACACGGACAGCGTTGAGTTTCACATCCATAACTTGCTCTGCTGCTTTCCAACCCGAAACTCTTTCAATGGAGATTCGGGAAGCATGAGCTTAACAAAGTATGAACAACGACTCAGGGATCTCCTTTTCAAGAACTTGGATGTGTCACAGCCTTTGAATGGGAAGCTCCTACCGAGTTCAATCCTACTGGGCTCTAAGGACTACCAGGTGAGGCGGCGGCTAGGGAGCGGGAGCCAGTACAAGGAGATCATGTGGTTGGGTGAAAGCTTTGCTTTAAGACACTTCTACGGGGAGATTGAGCCTCTTGTCCCTGAAATATCCCAACTCTTGTCCCTCTCCCACCCGAATATAATGCGCTTACTTTGTGGGTTCATTGATGAGGAAAAGAGAGAGTGTTTTTTGGTTATGGAACTAATGTATAAAGATCTCTGCAGCCATCTCAAGGAGATATGTGGACCAAGGAGGAGACTTCCATTTTCTCTTCCTGTTGCAGTTGATCTTATGCTTCAAGTTGCAAGAGGAATGGAGTATCTCCACTCAAAGAAAATCTACCATGGGGATCTAAATCCTTCTAACATTCTTGTTAAAGCAAGGAGCATCTCCACAGAAGGGTATTTGCATGCCAAAGTTTCAGGGTTCGGTTTATCTTCCACTAAAAACTTAAACCAGAAAACTTCTCCAAATCAAGCTGTGAATCTCCCGTTTATATGGCATGCTCCAGAAGTTCTAGCAGATCAAGAACAGCTGGGTAAAACTGGGAATTTCAAGTACACAGAGAAGGCAGATGTGTACAGTTTTGGAATGGTTTGTTTTGAACTCTTGACTGGGAAAGTCCCTTTTGAGGATAGCCATCTTCAAGGAGATAAGATGAGTCGAAATATTAGGGCAGGGGAGAGGCCCCTTTTCCCATCCAATTCACCAAAATACATAACAAACTTGACAAAGAAATGTTGGCATACTGACCCGAATCAGCGGCCAAGCTTCTCATCCATCTGTAGGATTCTTAGGTATACGAAATGGTTCCTAGCTATGAACCCTGATCACAGCCAGCCAGATGCCCCAGTGCCACTTGTTGATTTCTGTGACATTGAGGCAGGGCTTCTGAAGACGATACCCTCATGGAGAAGTTCTGATCCACTGCCAATATCAGACATCCCATTTCAGATTTTTGTCTATAGAGCTGTGGAGAAGGAGAAATTACATGCAATTCCTAAACAAAAGAGTTCAGAATCAGGAAGTGATGGATATTCAACCTGTGGAGATGAGCCTCTGACCATAATTGATGATCCACCCCCACCCATGACTGAAAAGAAGCCTTCAGTTTCTTCTGAAACTACGAGCAAAAAAATTTCAGTCTTGAAGAAATCTTCCAGTGTGAAAGTCATTAAGCAAACAGGAACACCTAAAGGACAATTGCTCAGACCTCCACTGATGAGCCCATGTGGGCGTACTATCAGACTGAGCTCCGAGGGGCATCTAGTGGCAATGATCCCAAAACCACGAAGAACACGTGGGCATGCCTCAGATTCTGAGCTCTCCTAG

>Z_mays_1

ATGGACCAGCTCCGCCAGGTGGGCGAGGCCCTCGGCGGCATCCACACGCTCATGGCCTTCGCCGACGATCTCCGCATCAACCCGCGACAATGCCGCCTCCTCGCCGACGCCTGCGCGATGGCCTTCGCCGCCGTCGCCGCCGAGGTGCGCGCCCACCTCCGCTTCAACGAGCGCCTGTCCAAGTGGAAGCCCCTGGAGGCGCCGCTCCGGGAGCTCCACCGCGCCGTCCGCGACGCCGAGGGCTACGTCCGCCACTGCATGGAGCCGCGGGGCAGCTGGTGGGGGCGCGCCGCCGCGGCCACGCACGGCGCCGACTGCGTCGAGCAGCACATGCACAGCCTGCTGTGGAGCGTCGCCGTGGCGCTCGAGGCCGTCGAGCTCGTCTCGGAGGTCACGGGGTCCGACCCGGACGAGCTCGCCCGGCGGCGGCTGCTCTTCGCCAAGGACTACGACAGGGACATGCTGGAGCCGGCGCTGTTCGTGCAGAGGCTCGGCGCGCGGTACCTGGCCACGCGCGAGCTCGCCGCCAGGATGGACGCGGCGTGGAAGGAGGACCGGTGGCTCCTGTCGCAGTACCTCGAGGAGCGGATGAGCCCGGGCTCGCCGGAGCCGGTGCCGCGGAGCAAGCACCGGCTGGCCGACCTCCTGACCGCGCCGCGCGGGCAGGTGCACCCGGCGTCCGTGCTCCTGCAGGGCGACTTCCACGTCCGCAAGCGCCTCATGGGCAACCTCAAGGAGGTGCAGTGGATGGGGGAGGCCTTCGCGGTGAAGCACCTCGTGGGCGCCGGCGCCGACGCCGACGCCGCGTGCGCCGAGGCCGCGCTGCTCACCTCCGTGGCGCACCCCAACGTGGCGCACTGCCGGTACTGCTTCCACGACGAGGACAAGAAGGAGTTCTTCCTGGTCATGGACCAGGTCATGACCAAGGACCTGGCCACCCACGTCAAGGAGGCGAACAGCGGCAAGCGACGGGTACCCTTCCCGCTCGTTGTCGTCGTCGACGTGATGCTGCAGATCGCGCGCGGCATGGAGTACCTGCACTCGAGGAACATCTACCACGGCGATCTGACCCCGTCCAACGTGCTCGTCAGGACGCGGCATGCCGACGCGCACCTGCACGTCAAGGTGGCCGGGTTCGGGCAGCACGCGGCGGCGGCGGCCGCGGCCGCGGCCGGCCACTGCCACAGGCCCAGCCCTAGGGCGTCAGCGAAAGCATCAAACGCCACCAGCGCCCCTTGCATCTGGTACGCGCCGGAGGTGCTGGAGCAGCAGGAGGCGGCCAAGCGCACGGAGAAGGCGGACGTGTACAGCTTCGGGATGGTCTGCTTCGAGCTGCTGACGGGGAAGGTCCCGTTCGAGGACAACCACCTGCAGGGGGAGCACATGAGCAAGAACATCCGCGCCGGCGAGCGGCCGCTGTTCCCGTTCCAGGCGCCCAAGTACCTGACCAGCCTGACCAGGCGGTGCTGGCACGGGGACCCGGCGCAGCGGCCGTCGTTCGCGTCCGTCTGCCGCGTCCTCCGCTACGTGAAGCGGTTCGTGGTCATGAACGTGAACCCCGCCCCCGCGGAGCAGCCAGACGCGGCGGTGCCGCCGCCCGTGCCGGCGGTGGACTACCTCGACATCGAGGCGAGCCTGCAGAGGAGGTTCCCGGCGTGGCAGCCCGGGTCTGGGAACGCGGCGCCGCGGGTCTCCGACGTGCCGTTCCAGATGTTCGCGTACAGGGTGGCGGAGAAGGAGAGGAGCCGGGCGGCCATCCTGCACATCGGCGGCGGCAAGGGCGCCTCCGACTCCAGCAGCGACGGAAACTCGCTGTGCGGGGACGAGAGCGGCGGCACGGCGCTGTCTGAAGCCGAGGCTCTGACCGTGTCGAGCCGCGGCGCCACCACCACGCGGTCGCTGCCGGACCGCGCTGGCAACAGGAAGGTGGATGGCAGCAGGGTCGCCTCCAGGCTAGCAGGCAAGCCTTATGTCCATAGCTAG

>Z_mays_2

ATGGAGCAGCTCCGGCAGCTCGGCGAGGTGGTGGGCGCCATCGACGCGCTCATGGCGTTCGAGCCCGAGCTCCGCGTAAACCCGCGCCAGTGCCGCCTCCTGGCCGACGCGTGCGCCCGCGCGCTCGCCGCCGTCACGGGCGAGGTGCGCGCGCACCTCCGCTTCGAGGAGCGCGGCGCCAAGTGGCGCGCCGTGGAACCCGCGCTCCGCGAGCTCCACCGCGCGTTCCGCGACGCCGAGGGCTACGTCCGTCAGTGCCTGGACCCGCGCGGCGGGGGCAGCTGGTGGGGGCGCGCCGCGGCGGCGGCGCACTGCACTGACTGCGTGGAGCAGCACCTCCACGGCATCCTCTGGTGCGTCGCCATCGCGGTCGAGGCCGTCGAGGCCGCCGCGGAGATCGCAGGTCACGACGCCGACGAGACCGCGCGGAGGCGTCTGGTGCTCGCCAACAAGTACGACAGCAGGAGCATGCTCGAGCCCAGGCTGTTCCAGCACGCGTACGGCAAGCTGTACCTGGTGTCCCAGGAGCTCGTTGCGCGGATGGACACGGTGTGGAAGGAGGACAGGTGGCTGCTATTGCAGCTGCTCGACGAGATGAAGTCGCCGGCGGCGCCGAAGCCGCTGACGAAGAGCGAGCAGCGGCTCGCCGACGTCCTGGCAGCGCCGCGGGGAAAGCTGCACTCGGCGTCTGTCCTGCTCAACGGCGACTACAGCGTGCGGAGGCGCCTCGGCGGCAACCTCAAGGAGGCGCATTGGATGGGGGAGAGCTTCGCGGTGAAGCATTTCATCGGGGACTCCGAGGCCGAGGTCTCGATGCTGTCGTCGGTGGCGCACCCGAACGTGGCGCACGCCGCGTACTGCTTCCACGACGAGGAGAGGAAGGAGTACTTCGTGGTCATGGACCAGCTCATGGTCAAGGACCTCGCCAGCTACGTCAAGGAGATGAGCAGCCCGCGGCGGCGAATCCCGTTTCCCCTCGTCGTCGCTGTCGACATCATGCTGCAGATTGCGCGCGGGATGGAGCACCTGCACTCCAAGAAGATCTACCACGGCGAGCTGAACCCGTCCAACGTGCTCGTGAAGCCGCGGCAGCCCGACGGGTACGTGCACGTCAAGGTCGCCGGGTTTGAGCGGCGGGGCGCCGTCACAAATGGCGCAAAGGCGTCTGCTAATGGCAGTGGCAATGCCAGCGCAGCCGGCAGCGGCGGCGACGACACCTGCATCTGGTACGCGCCGGAGGTTCTCGAGCATCAGGAGAGCCGCGACAGGCACACCGAGAAGGCGGACGTGTACAGCTTCGCGATGATCTGCTTCGAGCTGCTGACGGGCAAGGTCCCGTTCGAGGACAACCACCTGCAGGGCGACAAGACGAGCAAGAACATCCGCGCCGGCGAGCGGCCGCTGTTCCCGTTCCAGGCGCCCAAGTACCTGGTCGCCCTGACGAAGCGCTGCTGGCACGCCGACCCGGCGCAGCGCCCGCCGTTCGCCTCCGTCTGCCGCGTCCTCAGGTACGTGAAGCGGTTCCTGGTCATGAACCCGGAGCAGCAGCAGGGCCAGGCCGACGCTCCGCCGGCCGTGCCGGCCGCTGACTACCTCGACATCGAGGCGCAGCTGCTGAGGCGGATCCCGGCGTGGCAGCGTGGCCAGGGCGCACCACCGCGCGTGGTGGACGTGCCGTTCCAGATGTTCGCGTACAGGGCCGTGGAGAGGGAGAAGACCGCCGGCGCGCACGCCAGCAGGGACAGGGTCACGGATTCCGGCAGCGAGGAGGACTCGCTGTGCGGCGACGAGAACGGGGTCGGCGCCACCACGCCAGACGACGACGCGTCCACCGTGTCAAACGGCACTGTGAGGTCGCGCCCGGACAGCTGCGACGGCAAGAAGACTCCGGTCAGGAAGGCAGACGTCAAGGCGCCGCCCAGACAAGGAGGCGCGTTGCATCCTGTCGCCACATCTTCTTCGAGAAAACTTGAGAGGACCGATTCTGAACATGGTACTTCTGGATTCTCTGCAGGGTCTCAACAGAAGGTGAAACCGGCGAGCGCGGTGAAGCCTCCGCCGGCGACGAGGAAAACGGTGGGCGTGAAGCCCGAGCATCCGGCGCGGCGGCCGAAGTCCGGGCACGCCTCGGACTAG

>A_coerulea_1

ATGGAGCAATTTCGGATGGTTGGAAAGGTACTGGGAAGTTTAAAGGCACTTATGGCTTTCAAAGGTGATATTCAAATCAATCAGCGACAGTGCTGTTTGTTGGTTGATATATTCGAAAACGCGTTTGAATGCATATCAGAAGAGATGAGAAACAATCTCATATTTGAAGAAAGGCACTTAAAATGGAGGCCTCTTGAACAGCCTTTAAGGGAGCTTCAAAGGATCTTTAAAGAGGCTGAATTTTATGTTCATCAATGTTTAGATGTGAAGGACTGGTGGGGCAGGGTTATTTTACTTGCTAACAATACAGATTGTGTTGAGATTCACATTCATAACATGCTCTGCTGCATTCCTACTATCGTTGAAGCTATCGAGATGATTGGAGAGATCACTGGCTCTGACCAGGATGAGATGCAAAAGAAAAAGCTTATACTCTCCAAGAAGTACGACAGGGAATGGCAAGACCCAAAGCTTTTCGAGTGGAAATTTGGGAAGCAGTACCTGATTACTCAGGATATATGCAACAAGCTGCAGAGGGTTTGGAAGGAGGATAGATGGGATCTTCTTGAAACCATTACCAAAAAGAGGAACTCCGGGAGTTTAACCAAACATGAGCAACGACTTGCAGAATTCCTGCTAGACAAATTAAATGAGTCGGAGCCCACAAATGGAAAGCTCTTCTCGAGTTCGATCTTAGTAGGTTGGAAGGATTACCAAGTACGGAGACGGTTAGGGAACGGAAGCCACTACAAGGAAATCCAGTGGTTGGGTGAAAGCTTCGCTGTAAGACATTTTTTCGGAGAGATAGAATTAATAGTCCCGGAAATATCTCTTCTCTCGTCCCTTGTTCACCCCAATGTGTCACAGTTCCTCGGCAGTTTTTGTGATGAGGAGAAGAAAGAATGTTTTTTGGTCATGGAACTGATGAATAAGGATCTTTCTACTTACATAAGAGAGACCGGCTCTCCAAGGAAGCGGCTCCCATTCACTATACCAGTGGCAGTTGATCTAATGCTTCAGATTGCCAGAGGGATGGAGTATCTTCATTCTCGGAAGATCTATCATGGAGATTTAAATCCTTCTAATATCCTTGTCAAAACAAGAAATTCATCTTCAGAACTTTACTTCCATGCAAAGATCTCAGGTTTTAATCTGTCGTCCATTAAGAATTTCACCCCTAGAAGTACTTCTTCAACACAAAGTGGCACTCAGACATTTATATGGTACGCCCCAGAAGTTCTATCAGAGCAAGAAAATCCAGGGAACATCTATTTTGCCAAGCACACCGAGAAGGCTGACATCTACAGCTTTGGGATGATATGCTTCCAGCTTCTGACAGGAAAAGTGCCATTTGAGGATAGCCATCTTCAAGGAGACAAGATGAGCCGGAACATAAGGGCAGGGGAGAGGCCACTCTTTCCACATTCCTCACCCAAATACCTTGTTACCTTAACAAAGAAATGCTGGCAGACTGATCCAACTTTACGCCCTAGCTTCTCGTCCATCTGCAGGATTCTCCGTTACGTCAAGAAATTCCTGCTGATGAATCCTGATCACAGCCAGTCCGATGTACCATCATCACCGCTGATAGATTATTGTGAATTGGAAGTGGGTCTTTCAAAGAAATTCCCCACATGGGAGAGCACAAACTTAACACCAGTATCACAAATCCCATTTCAAATGTTTGCCTATAAAGTTTCAGAAAAGGAAAAGATTCATTTAAACATCAAAGAGAAGGGCTCAGAATCAGGCAGTGATGGAGCTTCTGTCTGTGGGGATGATAATATAGGTACTGTGGAAGATCCGTTTCCACCAACAACAGAGAAGAAGCATGCAGTTAGTGCTGAATTTATCATTAAAAAGACTTCAACAAAGAGGATTACTGATGGGAAAACATCCAAACCAGATGCAATTCAACAAGGAGCAGCCCCTGTTCGGGAGCAGTGTGATCAAAAGAAAACCAATCGGATCACTTTCTGGGAGCGTTGGGATCAAAAGAAGGCCAACTGGAGGAGTATAGCTTCAAACAAAAATACTATTTTGACTGTAGCTGCTAATAGTGCAGATAAAATATCAAAACGACCTGCTCAACGATTTGTACATTAG

>A_trichopoda_1

ATGGAACAACTCAGACAGATTGGAGAAGTTTTGGGAAGTCTCAAAGTCTTAATGGTTTTCAAGGATGATATTCGGATCAATAGGCACCAGTGCTCTCTCTTATATGAGATTTTCAAGCTTGGGCTCCAAGGAATTGCGGAAGAAATCAGAGACAACCTAAAATTCGATGAGAGATTGAAGAAATGGAAGGCTTTGGAACAGCCACTGAAGGAACTCTTTAGGACTGTAAAAGAAGGCGAAAGCTACATCAAACATTGCTTAGAACCCAAGGATTCATTGGGTAAGGCAATATCTCTGCCTCAAAACAACGACTGCGTCGAATTCCACCTACACAATTTCATATGGTGCGTTTTTGTGGTGTTTGAGTCCATAGAAATTGCAGGAGAGATAGCAGGCTATGATCAAGAGGAAACCCAAAAGAAAAAGCTCCTCTTTTCTAAGAGGTATGAAAGAGAATGGCTTGACAAAAGTCTTTTCCAATGGAATTTTGGAAAGAAATACTTGGTTTCAAGAGAAATATGCAACAGATTGAATACTGTATGGAAAGAAGACAGGAAAATGTTTCTAGAGATATTGGAAGAGAAGCGAAGAACCAAATCAGAAAGTCGAATCCTTGACCTCTTGCATAAGAATTTCGAGGGCTCTGAGGTTTCAAAGGGAGACTTATTCCCTTGCTCCATATTAACGGATCCAAAAGACTATCAGGTGAGGAGGAGACTAGGGAATGGTGGCCAGCACAAAGAGATCCATTGGATGGGAGAAAGTTTTGCAGTGCGCCATTTTCACGCAGAGGTCGAACCCCTGATCCCTGAAATATCTTTATTATCTTCACTCATGCATCCAAACATTATGCATATCCTTTGTGGGTTTGTTGACAAGGAGAGAAAGGAATGCTTCCTTGTAATGGAATTAATGAACAGGGACCTTTCCAACTACATAAAGGAAACATGTGGTCCTAGAAAGAGGGTCCCTTTCACTCTACCAGTGACTGTTGATGTCATGCTTCAAATAGCCAGAGGAATGGAGTACCTTCACTCTCGAAAGATATATCATGGGAATCTCAATCCTTCAAACATTCTAGTTAGAGTGAGAAACCCATCGTCAGAGGGACATCTGCATGTGAAAATCACCGGTTTTGGGCTCACCTCTGTAAAGAATTCAACCTCTAAGAACTCTGCAAATCAACCCACAACAGATCCAGTTATCTGGTATGCACCTGAAGTTCTTCAGGAGCAAGAACAAGGGAACCCTACGAGTCCATCAAAGTACACAGAGAAAGCAGACGTTTATAGCTTTTCAATGATATGCTTTGAGTTGTTGACTGGTAAGTTGCCTTTTGAAGAGGGGCACTTACAAGGAGACAAGATGAGCCGGAATCTAAGAGCAGGTGAGAGACCATTGTTTCCTTTTCCTTCTCCAAAGTATCTCACTGCCTTAACCAAGAAATGCTGGCAAGACGACCCAAACCAGAGGCCAAACTTCTCATCAATATGCAGGACTCTTCGATACATAAAGAGATTCTTGGTCATGAACCAAGATCATGATCAGCCTGATGCCCCTCTTCCCCTGATCGATTATATGGACTTGGAGGCAAACTTCTCAAAACGCTTATTGGGTTGTGGTGAAGAATTTACAGAGTCAGTTTCTCTCATCCCATTTCAAATGTTCGCTTATCGGGTTATAGAGAGACAGAAAACCATGGTGAATTTGAAGGAAAAGAACTCAGAATCAGGGAGTGATGGAGCATCTGTCGGGGGGGACGAGAACCAGGGGAATGTGGAAGAGACATTCTCCAATCTCAAGCCTACATCTCGGTCCTCTCAGAGTAGTTCAGAGTCCACCAAGAAACCAGCCTTTGCAAAGAAGACCACAAATGGAAAATCAAGAAAATCATCAGATCAAATTGTTCAGTACAATGAAAACAGAGGCCAAGTGTGGTTATAA

>C_canephora_1

ATGGAACAGTTCCGCCAGATTGGGGAGGTTGTGGGAGGTTTGAAAGCTCTAATGGTTTTGAAACATGATATTCCAATCAACCAGAGACAATGCTGTTTGCTTTTCGATATGTTTGCGTTGGCTTTTGAAACTATTTCGGATGAAATCAGGCAAAACTTAAGGCTAGATGAGAGGAATGTCAAGTGGAAAGCTCTCGAGTATCCTCTGAAAGAGCTTCATAGGGTATTCAAGGAAGGTGAACAGTACATAAGATTGTGCCTGGATGTCAGGGATTGGTGGGGGAAGGCAATCAGTCTCCATCTCAACAGGGACTGTGTTGAGTTTCACATTCACAACTTGCTTAGCAGCTTTCCCGTTGTCGTTGAGGCTGTTGAAGCTGTTGCAGAATTCTCGGGTGCAGACCTGGAAGATATGCAAAAAAGGAGAGTTGCACTAATGAGAAAATATGAAGCAGATTGCAGTGATCCAAAATTCTTCCAGTGGATGCATGGTAAGCAGTATTTGGTGCCTCGAGAGATATGCAGCCGCCTACAATCTGCTGGGAAAGAAGATAGATGGTTTCTCCTCGAAGCAATAAGAGAAAAGAAAAGTGCAGTTCCATCTACGTTGGCAAAGCATGAGCACAGGCTTGGTGACCTGTTGCTGAAAAAATTAAGTGGATCGGAGCCACCAAAACTGAAGCTCCTGCCAACTTCAGTTCTAGTGGGAGCACATGACTTCTACGTGAAGCGGAGGTTAGGATTAGGTGAGGGACACATGAAAGAAATCCAGTGGCTGGGGGAAAGTTTTGCTTTGAGGACCTTCTTTGGAGAGATAGAACCATTACACGAAGAGATTTCTCTAGTTCTTTCACTCTCCCATCCCAACATTTTGCAGCACCTCTGCGCTTTCTATGACGAGGAGAGGAAAGAGGGCTTTTTAGTGATGGAGCTTATGCACAAGAATCTTAAAAATTACATAAAAGAGAATTGTGGGCAAAGAAAGCGAATTCCTTTCTCTGTTCCAGTTGCGGTTGATATCATGCTTCAGATTGCTAGAGGACTGGAGTATTTGCACTCAAGGAAGATTTATCATGGGGAACTGAACCCATCTAACATCCTTCTAAGAGCTAGACATTCATCCACTGAGAGTTACTTTCAAGCAAAACTCACTGGATTTGGATTGGCCTCTATCAAGAGTTTTGCTCGATCTCAACATTCGAGTGTTGACCCTGTTATATGGTATGCCCCAGAAGTTCTCGCTGAACAAGAGCAGCCAGGAAAAAAGGGCAACGCCAAGTATTCTGAAAAAGCTGATGTCTACAGTTTTGGGATGCTCTTTTTTGAACTTTTGACAGGGAAAGTTCCATTTGAAGACAGCCACCTCCAAGGGGACAAGATGGTTCGAAATATTAGAGCAGGAGAGAGGCCACTTTTCCCATATCCAGCGCCCAAATACCTGTCAAACTTAACCAGAAAATGTTGGCATCCAAATCCATTTATCCGTCCGAGCTTTTCTTCCATATGTAGGATTCTACGCTACATTAAAAAGGACCTCATAATTAATCCTGAACATGGTCAACCTGAATCACCACCTCCACTTGTGGACTACTGCGACATAGAAGCAGCGTACTCAAAGAAATTCCCGGGGGAGGAGAGCCCTGATTTGAGCCCGGTAACCGATATTCCCTTCCAGTTGAATGCTTATAGACTTGTTCAGAAAGAAAAGAGTTGCGGAAGCTCTAAAGACAAGAACTGCGATTTGGGAAGCGAAGAACTACCTCAAAGGCCAGCATCAATATATGGTGATGAGCATGTGGCCGCAATAGATGATCTCTTTTTGGTACCAAGTGATCGAAGGTCAGTTTGTTCAGAAATAATTGAAAGCAAGAACATTCGGGCGGCATATGATGAAAGGTCGGTTATCTCTGAGATTCCACACAAACTATTCTTCTCTGATCAGAGATCAATTGGTTCTGAGAGTCCGGGGAGGAAATCTCTATTAGCAGCTCCAACTGAGCAAAGGCCAAATGTTGCTGATACTCCACAAAGGAAAGTCTCAGTGGCAGTAGCAGTCGATCAAATCTCCTCTCTTTCTCGAACTCCAGAAAGAAAAGTTCCTCCAACAGCAACAATCAATCACAAGACAAAGCATTCTGAGAACCTGGGGAAGGATATGGGGTCAATAGCAGATCAGACGACCGCTGATTCTAAGTCTCAAGATGGAAATTCCAAGTCAAAAACTGCAGCTAAGCTGCATGAACAAAGTAATGGACGTTCCAATGTACCAGAAAAAGAAATCAAATCGAAAACAGCTGCTGACCCTGTTCCAGTTACTTCTGGAGCTTTATCAAGGAAAGTTTCGTCAAGGAAAATAGCTCATCAAAGAAAACTTTCTGAGATTCCAGAAAAGTCTGTCTCAGCTTCCTTATTAAATCCGAAGTCAAATTCTTCCAGTAAGAAAAGTGAACCAACCAGACCTGCCGCCAAGCCCTCTCCTCCCAAGAAATCGGACAAGAAGCATTCCATGAACAAGAAAATGAAAGATGTAAACTCAGAAAAAAATCCAGGAAATTCAAAAGATAACTTACCAAGGTCTTCGTCTGCAAGAATATCACAAACGATTGCATCATCATCAGCAGCAGCAGCAGCATCACCAACAAGAGGAACAAGGTTTTCATCAACATTTGCGTTGAAAGGGTATATGTCACCACAAGCATCACCACTGCATCCATGTGCTCGTTGCTGCCGGACAAGCAGAGAGGCCATCTTGTCTTCACCAGCAATGAGCCCATGCAGAGCTAGACTAACGCATGTATCGGAGACGGAGATTGCATAA

>C_canephora_2

ATGGAGCAATTCAGGCAGATTGGAGAGGTGATTGGAAGTCTCAAGGCATTGATGGTCTTTCAGGATGACATACAAATTAATCAGCGACAGTGTTGCTTGCTGGTTGACATGATGAATCGTGCTTATAAAACAATTGCCAAGGAGATGCAACAGAACCTGAGATTTGAAGAGAAGAGTACGAAATGGAAGGTGCTTGAGAGTCCCTTAAGAGAGCTCCTCAGGATAATTAGGGAAGGAGAAATTTACATCAGGCAGTGCCTAGAAACGAAGGACTGGTGGGTTAAAGCTATAAGGCTTTATCAGAATTCAGACTGTGTTGAGTTCCATATACATAACCTGCTCTGCTGCATCCCCATTATACAAATTTCTGGTTCAGTAAGTTCAACAAAGCAGGAGCAGCGACTCGCTGATCTTCTTCTGAAGAACTTAAGTGTGTCAGGACCAGTTGAAGAGAAGCTCCTGCCTTCCTCCACCTTAGTGAGCTCTAAGGATTACCAGGTCAGGCGGCGCTTAGGAAGTGGAAGCCAGTACAAAGAGATTCAATGGTTGGGTGAAAGCTTTGCTCTGAGACACTTTTTTGGGGACATTGAACCATTGTTTCCAGCAATCTCTCAAGAATTAGCACTTACTCATCCAAATATTATGCATATCTTTTGTGCATTTACTGATGAGGAAAGAAAGGAATGCTTCTTAGTTATGGAACTCATGAACAGAGATCTTTCCAGTCATATAAAAGAAATTTGTGGACCAAGGAAGCGAATTCCATTTTCTCTTCCTGTTGCAGTTGACCTTATGCTTCAGATTGCAAGAGGAATGGAATATCTCCACTCAAAAAAAATCTATCACGGAGATTTGAATCCTTCAAACATTCTTGTAAAATCCAGAAATATCAATACAGATGGTTACTTGCATGCAAAAGTTTGCGCATTTGGACTTCCCTCCTCTGTCAGCCTTCCTCAGAAATCCAATTCCAATCAGAATGGGCAGCTTCCAGTCATCTGGTACGCTCCAGAGGTGCTAGCAGATCAAGAACAATCAGGCAATGCTGATAACTCCAAGTACACGGAGAAGGCAGATGTGTATAGTTTTGGGATGATTTGCTTTGAGCTTTTAACAGGGAAAGTCCCTTTTGAAGATGGCCATCTCCAAGGAGATAAAATGAGCAGAAACATTAGAGCAGGGGAAAGACCACTATTCCCATTTCACTTGCCAAAATATTTGACGAATTTGACAAAAAAGTGCTGGCACGATGACCCAAACCAACGTCCAAGTTTTGCATCCATCTGCAGAATTCTCCGGTACATAAAGAGATTCTTGTTGATGAACCCTGATCATAGCCAGCCAGAAACCCCAGTGCCACTCGTGGATTACAGTGAAATTGAGACAGTGATCTTGAGGAGCTTTCCTTCTTGGGCAGAATCCAATATATTGCCCATATCAGAGATTCCCTTTCAGATGTTTGTATACAGGGTAGCAGAGAAGGAAAAATCAAATATAAGCCAGAGAGAAAATTCTGAATCAGGAAGTGATGCTTCAGCATGTGGGGATGAAAGTGCAACTGTGGATGATCCATTACCATCACCCTCTGAAAGGAGGTCTAGTGCCTCTCCAGAAACTATGAACAGGAGAATGCTATCGTCCAAAAAATCAGCAGATGTCAAACCCATTAAACAGCCAGGGACACCAAAGGGTCGAGTAAGGCCCCCTCCTATGTCTTCTTGTGGACGGACTATGAGGATGAATTCAGAAAGCCAGTTAATGGTAATGAGTCCAAGACGGAGAGCATCTGGTCACGTATCAGATTCAGAGCTCTCATAG

>P_taeda_1

ATGGATTACTTGCGTCTCTCAGGACAAGTTATTGGAGGCCTTGACATTTTGTTGTTGCATTTTGAAGATGCTATTCTTCTTCACAGAAAGCAATGCTCTGTTCTGGCCAAAGTTTTCTCTCATGGAGTCAACCACATTTCAAATCTGATAAAAGAGACATTGAGAACGGATCAGGAAGTGATGCAGGAAAAATGGAAATTGTTGGAACCAGCACTGAAAGAGCTTTACAACACAGTAAGAGAAGGGGAGCTATTCATCAGACAATGTTTCAGTTCAGGAGGGAAAGATTGGTGGGTGAAGGCCATTACTCTGTATCAGAGCACTGAAGCAATAAACCTCCATCTCCACAATTTCTTATGGTGCATCTGCAGTCTTGTTGATTGCATTGAACTTGCAGCTGAGGAAACTACAGGAGATGAGGAGGAAGAAACCCAATTGAGAAAGCTCTTGCACTGCAAGAAATATGATAAGGATTGGCTTCATCCAAAGCTTTTCGAGTGGCAATTTGGAAAACAATATCTTGTTTGTGAGGAGATCAAGAGTTTTTCCAGTCTCTGGGAACAAGATATGAAGGCATTGCAGGGAGCCATGAAGAGCAAGATCATGAATTCCTGCAAAGACTTGCATGTGAAGAGAAGAATTGGAAGTGGAAGCTTTGGTGAAGTCAAAGAGATTATCTGGATGGGAGAGAATTTTGCAATGAGGCATATGTTTACCAATGTGGAAATATTTCTTCATGAGGTCTCTGCAATGTCTAAGCTTTTGCACCCAAATCTTTTGCAGCCATTAAGCTCGTTTGTAGATAAAGACAGAAGTGAGTGCTGTCTGGTGACAGAATTGATGCATAAGGATCTTCGTAGCTTCATCAACGAGAATTCTTCTCCTCGCAAGAGAATCCCATTTTCACTTCCAAGGGCAATAGATATTGTGCTTCAAATAGCCAGAGGGGTGGAGTATTTGCATGCCCATGGTATCATCCATGGGGCCTTAAAATCATCCAATGTACTCGTCAAGGTAGGACCAATTGAAGGCCAAATCCATCTGAAAATTTCAGACTTTGGACTCATTAAATCAAATCTGTTGAGTTCAAAATACTCATCTAGTTCAGGAGATACCCCTTCCTGTTGGTGGCGCGCACCAGAAGTTCAGAGCATTAGGCCTGAAGATGAAGCAGGGGCCAGGGTTTACCCTACCAAATATACAGATAAAGCAGATGTCTACAGTTTTGGCATGGTCTGCTATGAGATTTTGACAGGGAAGATACCGTTTGATGATGGTCGTGTGAAATCAGATATTGCCAAGATGAAATCTGGAGAAAGGCCTCTGATTCCTTCAACTTTTCCCAGGTATCTGAGTAGCTTCATTAAAAGATGCTGGCATGCAGATCCCAAATTTCGCCCCTCCTTTTCTTCCATATGCAGGATTCTCAGACACATTAAACGCAAAATATTTCTGGTTCCTGATCCCAATTCTGCCCAAGATTTGGTCCCTTCGCCGGATCTCTGTGACATTGAAACAAGCCTCGCCAGGCACTTGCCTCAGTGGGGGACTAGAGAATTTGGGGATCAAGTTGCCAATATTCCGCTTCAGATGTTTGCATTCAGAGTGATGGAAAGGGAAAAGATTATGAAATACAGTGCCGCGGGAGCAGTGGTTGCAGGAATCAAGAGTGTTGATAAAATCGTAGCAGCAGCAGAAGGCGCTGGTGGGAGTAAGAGTGCTGATAAAATGGAGAATTATGAAAATCATGAGAGCGTGGAGGGGTTGACAAATTCTGTATCGAATTCGATAACGAAATTGAAAGTACCCAGAAGTCCCAGCTATGGCTCTTCTTCGCCATTGCATCGTTCCAACAAAATTAAGGTTCTAGCTAAGAAGACATTCATCGATTCCAAAATATACAGATCTCCAGGAGAATCCAAAATGGGGGCATTGCAATCTGGAATAAGCCCAGATCGAAGGAGATCAGCTGCCGCAGTGTCAGCATACTACCTGAGCAGTTGA
